# Discovery of cephalotaxinone enzymes reveals a whole plant model for homoharringtonine biosynthesis

**DOI:** 10.1101/2025.08.26.672243

**Authors:** Yaereen Dho, Kevin Smith, Elizabeth S. Sattely

## Abstract

Plants produce diverse molecules that inhibit protein translation. A lead example is homoharringtonine (HHT), a key tool for ribosomal profiling and an FDA-approved treatment for chronic myeloid leukemia. HHT is commercially produced through semi-synthesis from the alkaloid core cephalotaxine (CET) extracted from endangered *Cephalotaxus* species. Despite its significance, the CET/HHT biosynthetic pathway remains unresolved. Here, we use paired untargeted metabolomics (stable-isotope precursor feeding) and transcriptomics to elucidate a near-complete biosynthesis to CET without prior knowledge of intermediates and biosynthetic genes. We show that while the CET core is biosynthesized only in growing root tips, CET and HHT accumulate throughout the plant. We discovered seven pathway intermediates and six novel enzymes that produce cephalotaxinone, the likely direct precursor of CET. Included are non-canonical cytochrome P450s, an atypical short-chain dehydrogenase, and a 2-oxoglutarate-dependent dioxygenase that together result in carbon excision and CET/HHT pentacyclic backbone formation. This study establishes a metabolic route to the HHT core scaffold and suggests a whole-plant coordination model in *Cephalotaxus*, where cephalotaxinone is produced in root tips and distributed throughout the plant for subsequent elaboration to HHT.

## MAIN TEXT

*Cephalotaxus* (plum yew) is a slow-growing coniferous tree from the *Cephalotaxaceae* family that produces over 70 structurally diverse *Cephalotaxus* alkaloids with the most prominent being cephalotaxine (CET)—the core structure of *Cephalotaxus* alkaloids^1–3^. Although no unique bioactivity for CET itself has been reported, its ester derivatives—such as harringtonine, isoharringtonine, deoxyharringtonine, and homoharringtonine (HHT)—have demonstrated potent antileukemic activity *in vivo*, each significantly inhibiting P388 lymphoid leukemia in mice at doses as low as 1 mg/kg^2,4–6^. HHT, in particular, exhibited nanomolar inhibitory activity against myeloid and lymphocytic leukemia cell lines^7^ and was approved by the U.S. Food and Drug Administration (FDA) in 2012 for the treatment of chronic myeloid leukemia (CML)^8,9^, especially in patients resistant to tyrosine kinase inhibitors (comprising 30% of CML cases)^10^. Also known as omacetaxine mepesuccinate (Synribo®), the HHT drug acts as a protein translation inhibitor by binding to the A-site of the eukaryotic large ribosomal subunit, preventing the initial elongation step in protein synthesis^11–14^. The specificity of harringtonine alkaloids has also made it a critical research tool in ribosomal profiling^15^.

Despite its biomedical value, HHT remains difficult to access. Natural levels of HHT in *Cephalotaxus* species are extremely low (∼0.06-3.2 mg/g dry weight)^16,17^, and its current commercial production primarily relies on a semi-synthetic route involving the extraction of CET from *Cephalotaxus* needles, followed by chemical esterification with a synthetic side chain^1,18–20^. While total syntheses of CET and HHT have been reported^18,21,22^, these approaches are limited by multi-step procedures and low yields, making them commercially less favorable. Furthermore, *Cephalotaxus* species have been classified as endangered due to extensive harvesting, inherently slow growth rate, and long seed development time^1,2,23^. Since late 2024, although still listed in FDA-approved drug, HHT has been discontinued in the U.S. in part due to manufacturing limitations^24,25^, underscoring the need for alternative production platforms. Metabolic engineering of *Cephalotaxus* alkaloid biosynthesis into tractable heterologous hosts could provide a sustainable, scalable production of CET and HHT; however, the development of such solutions has been impeded by a fundamental lack of knowledge of their biosynthetic pathways. Furthermore, knowledge of biosynthetic genes would enable studies to determine the specific biological role of these alkaloids in the producing species.

Early radioisotope labeling studies in actively growing *Cephalotaxus harringtonia* demonstrated that CET is derived from the amino acids tyrosine^26,27^ and phenylalanine^27,28^ (Figure 1). Based on these findings—and the minor presence of homoerythrina-type alkaloids in *Cephalotaxus* species—it was hypothesized that a phenethylisoquinoline scaffold is an intermediate of CET biosynthesis, formed from tyrosine and phenylalanine via a pathway analogous to those described for *Colchicum* and *Schelhammera* alkaloids^2,26–28^. However, the failure of cinnamic acid to incorporate into CET has questioned the validity of this hypothesis^2,27^. More recently, a phenethylisoquinoline scaffold (**1**) was reported in *C. hainanensis*, along with upstream biosynthetic enzymes capable of converting tyrosine and phenylalanine into **1**^29^, resembling the early steps in colchicine biosynthesis^30^. These findings support a biosynthetic pathway model that hinges on this Pictet-Spengler metabolite, but to date, there is no direct evidence that this compound **1** is a precursor to CET or HHT alkaloids (**Fig. 1**) and no additional pathway intermediates have been identified. Moreover, CET and HHT are found distributed throughout the plant body, and thus a site of active biosynthesis has remained elusive^2,16^. Collectively, these gaps have made elucidating the biosynthetic pathways of *Cephalotaxus* alkaloids particularly challenging.

**Figure 1.**
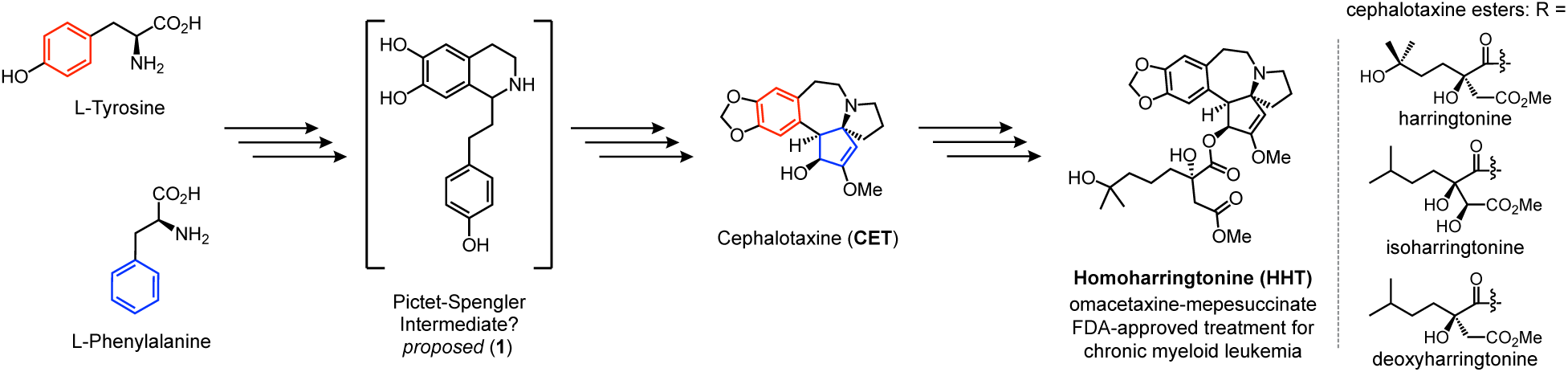
Proposed biosynthetic pathway of CET, the core structure of HHT, and its esterified derivatives. The proposed pathway for CET, the core structure of HHT, biosynthesis is based on radioisotope-labeling studies (shown in red and blue) and structural characterization of alkaloids from *Cephalotaxus* species. Over 20 distinct CET esters have been identified to date, four of which shown here exhibit notable biological activity. R denotes the variable ester side chain that can be attached to the CET core to form different CET esters (i.e. harringtonine, isoharringtonine, and deoxyharringtonine), whereas the HHT side chain is shown explicitly.

Here, we address these long-standing challenges by generating paired metabolomic (via isotope-labeled precursor feeding) and transcriptomic datasets across diverse tissues of *C. harringtonia* harvested during its actively growing season. This integrated approach revealed a site of active biosynthesis of CET, uncovered previously unknown key intermediates, and ultimately enabled the discovery of a set of highly co-expressed biosynthetic enzymes responsible for biosynthesis of the CET core.

## RESULTS

### Labeling studies identify growing root tips as a site of active biosynthesis of CET and specific phenethylisoquinolines as intermediates in the pathway

In studying biosynthetic pathways of metabolites of interest in non-model plants like *Cephalotaxus harringtonia*, where genomic resources and genetic manipulation tools are limited, transcriptomics with co-expression analyses has often been used when the spatial, temporal, or condition-specific production of the compounds is known^31–34^. However, sequencing a plant transcriptome without prior knowledge of when and where the pathway is expressed may not capture relevant biosynthetic genes. Moreover, knowledge of the correct biosynthetic intermediates is also crucial for the discovery of biosynthetic genes. In our prior work using a D_2_O labeling study, we proposed that HHT is actively biosynthesized in new growth needles of *Cephalotaxus*^35^. However, it was unclear whether the entire pathway or only a portion (i.e. the final steps) is active in these tissues. Given that our efforts to identify biosynthetic genes expressed in these tissues were unsuccessful^36^, we considered whether parts of the pathway were expressed in other plant tissues. Thus, we first attempted to identify a site of active biosynthesis for the alkaloid core, CET, specifically and elucidate potential intermediates in the pathway with stable-isotope precursor feeding experiment across different tissues of *C. harringtonia*.

To collect various tissues of *C. harringtonia*, we harvested plant tissues (young inner needles—YIN, young outer needles—YON, old needles—ON, young stem—YSt, old stem—OSt, and root tips—Rt) when the plant appears to be actively growing—this was typically spring^16,37,38^ (**Fig. 2a**). We then first compared natural accumulation of CET and HHT in different tissues using liquid chromatography-mass spectrometry (LC-MS). Both CET (**Fig. 2b**, left) and HHT (**Fig. 2b**, right) were detected at high levels in all different tissues collected in May 2022, with lowest in YSt. CET was about 50- to 200-fold higher in amount than HHT throughout the plant (**Supplementary Fig. 1a**). Tissue samples collected in May 2023 were also analyzed and compared (**Supplementary Fig. 1b and 1c**). In both sample sets, both CET and HHT are generally abundant in all tissues with no consistent pattern of accumulation.

**Figure 2.**
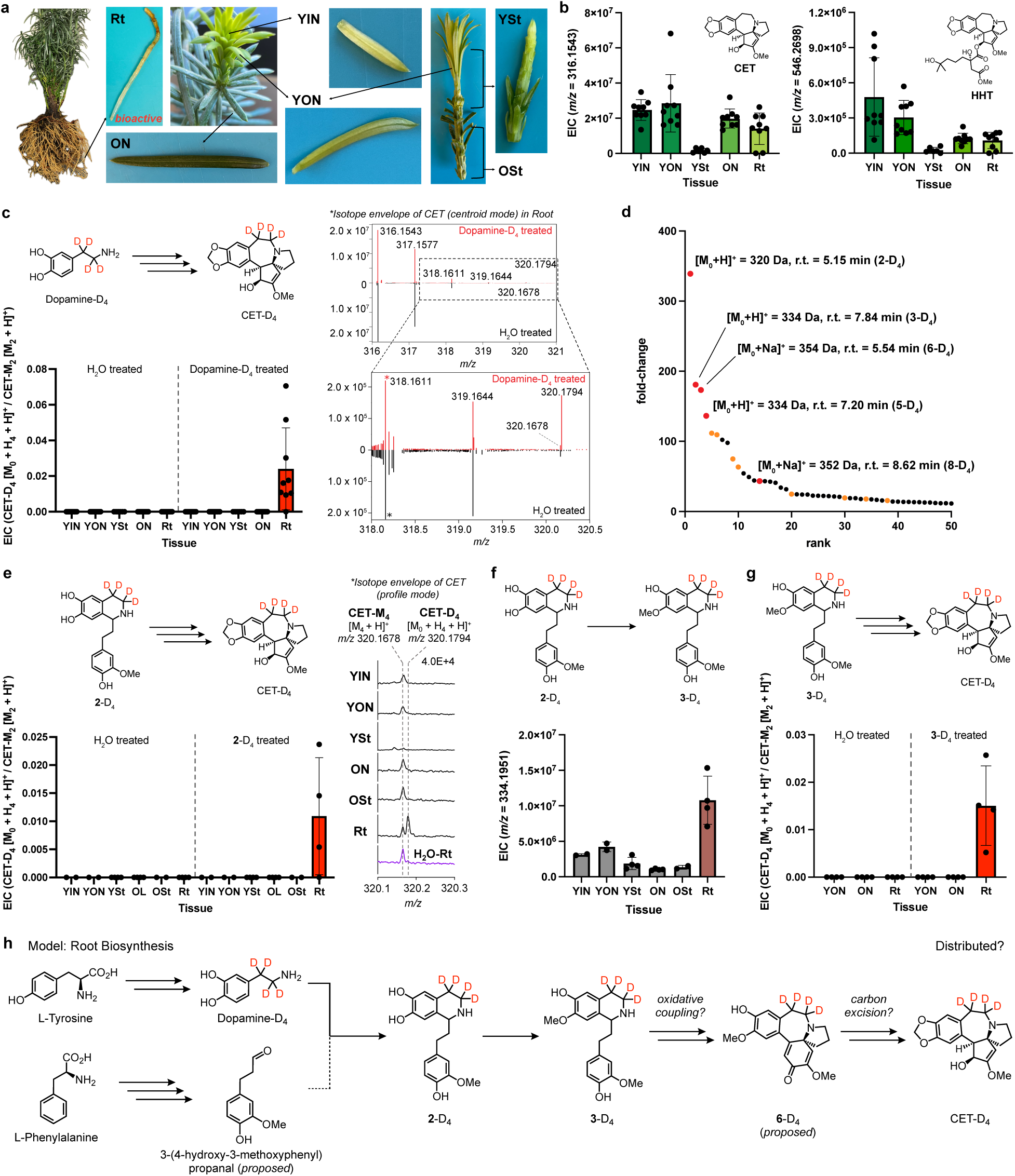
Stable-isotope precursor feeding study reveals that root tips (up to length of approximately 5-8 cm) of actively growing plant are a site of active biosynthesis of cephalotaxine and that specific phenethylisoquinolines serve as intermediates in this process. **(a)** Different tissues of *C. harringtonia* (young inner needles—YIN, young outer needles—YON, old needles—ON, young stem—YSt, old stem—OSt, and root tips—Rt) harvested during actively growing season of the plant (May or June) and dissected for paired feed-in experiments (metabolomics) and RNA-seq (transcriptomics). **(b)** Natural accumulation of CET ([M + H]^+^ = *m/z* 316.1543, r.t. = 2 min) and HHT ([M + H]^+^ = *m/z* 546.2698, r.t. = 10.8 min) throughout the native plant, *C. harringtonia* harvested in May 2022. **(c)** Relative isotope enrichment of CET-D_4_ in deuterated precursor dopamine-D_4_ feeding (right half of bar graph) compared to water treatment (left half of bar graph) in five different tissues indicates active biosynthesis of CET in the growing root tips (up to length of approximately 5-8 cm) of the native plant. On the right, isotope envelope of CET in centroid mode, comparing dopamine-D_4_ treated Rt (red) vs H_2_O treated Rt (black), is shown—with dopamine-D_4_ treated Rt having an extra peak of *m/z* 320.1794 corresponding to CET-D_4_ ([M_0_ + H_4_ + H]^+^). **(d)** Untargeted metabolite analysis (XCMS) comparing dopamine-D_4_ fed roots tips to water-fed root tips of *C. harringtonia* (*n* = 9 independent replicates for each experimental condition). The unique mass features (*P* < 0.2; fold change > 3.5; 200 < *m/z* < 600) are ranked based on their increasing fold change in abundance between the two conditions, with top 50 shown. Features confirmed to be above signal to noise and represent true mass signatures corresponding to potential intermediates (with four deuteriums, resulting from incorporation of dopamine-D4) in CET biosynthesis are shown in red and their isotopologues and/or adducts in orange. r.t., retention time. While not shown here, CET-D_4_ ranked 127^th^ out of 148 unique features identified by XCMS with increasing fold change. **(e)** Relative isotope enrichment of CET-D_4_ in **2**-D_4_ feeding (right half of bar graph) compared to water treatment (left half of bar graph) in six different tissues indicates active biosynthesis of CET in the growing root tips (up to length of approximately 5-8 cm) of the native plant and **2**-D_4_ as an intermediate in CET biosynthesis. On the right, isotope envelope of CET in **2**-D_4_ fed tissues (YIN, YON, YSt, ON, OSt, and Rt) and in H_2_O fed Rt in profile mode, focusing on the fourth isotope peak, M_4_ (M + 4), is shown. In the **2**-D_4_ fed Rt sample, an additional fourth isotope peak appears at *m/z* 320.1794, corresponding to CET-D_4_ ([M_0_ + H_4_ + H]^+^) due to four deuterium incorporation, which is distinct from the natural carbon isotope peak at *m/z* 320.1678, corresponding to CET-M_4_ ([M_4_ + H]^+^) observed in all different tissues fed with **2**-D_4_ or water. **(f)** Observation of **3**-D_4_ ([M + H]^+^ = *m/z* 334.1951, r.t. = 7.8 min) in all six different tissues treated with **2**-D_4_. O-methylation of **2**-D_4_ to **3**-D_4_ predominantly occurs in the actively growing root tips. **(g)** Relative isotope enrichment of CET-D_4_ in **3**-D_4_ treatment (right half of bar graph) compared to water treatment (left half of bar graph) in three different tissues indicates active biosynthesis of CET in the growing root tips (up to length of approximately 5-8 cm) of the native plant and **3**-D_4_ as an intermediate in CET biosynthesis. **(h)** Summary of updated proposed biosynthetic pathway for CET with stable-isotope precursor feeding experiments. Dotted line represents proposed aldehyde incorporation not yet supported by labeling studies. For all quantification, EIC of LC-MS was used. For CET-D_4_ enrichment studies **(c, e, g)**, CET-D_4_ was normalized by M_2_ (M + 2) isotopologue to avoid detector saturation effects, since CET ionized well in LC-MS and was highly abundant in our measurement conditions. Experiments in **(b-d), (e-f),** and **(g)** were conducted in May 2022 (*n* = 9 biological replicates for each tissue type, except YSt, *n* = 6), in May 2023 (*n* = 4 biological replicates for YSt, ON, and Rt, and *n* = 2 for YIN, YON, and OSt), and in June 2024 (*n* = 4 biological replicates for each tissue type) respectively. The values and error bars are mean and ± standard deviation (SD).

To determine whether CET and HHT are actively made in all different tissues, as detected, or synthesized in a specific tissue and transported throughout the plant, we next conducted a stable isotope precursor feeding experiment with samples from various parts of an actively growing plant. Sectioned tissues of *C. harringtonia* were incubated in a solution of dopamine-D_4_ (deuterium-labeled alkyl groups), a predicted early intermediate in CET biosynthesis based on the prior isotope labeling study that showed tyrosine serving as an upstream precursor^26,27^ and on the hypothesis that CET biosynthesis involves a phenethylisoquinoline scaffold^2,26,29^. Interestingly, enrichment of CET-D_4_ ([M_0_ + H_4_ + H]^+^ = *m/z* 320.1794) was observed only in the root tips (up to length of approximately 5-8 cm)—Rt, and not in other tissues (**Fig. 2c** and **Supplementary Fig. 2**). These results support a model where CET biosynthesis from dopamine occurs at the root tips of an actively growing plant, contrasting with the widespread accumulation pattern of CET, and that dopamine serves as a key pathway precursor.

We then performed an untargeted metabolite analysis (XCMS) comparing dopamine-D_4_-fed root tips with water-fed controls to search for pathway intermediates. A ranking using increasing fold change in abundance between the two conditions (fold change > 3.5 and *P* < 0.2), with *m/*z in between 200 and 600, revealed several deuterium (D_4_) labeled metabolites that may serve as potential intermediates in CET biosynthesis. Notably, these mass features each have an *m/z* close to CET and a large fold-change relative to others in the list of 148 unique mass features found (**Fig. 2d** and **Supplementary Fig. 1d**). We searched for both the deuterated and unlabeled isotopes of these molecules using extracted ion chromatograms (EICs) from both dopamine-D_4_ and water treated root tips and found that while the unlabeled isotopes were present in both sample sets, the deuterated isotopes accumulated only in dopamine-D_4_-treated samples. These results indicate that these molecules not only arise from dopamine-D_4_ but also are true metabolites of *C. harringtonia*. Interestingly, the first two mass signatures in the ranked list were found to correspond to specific phenethylisoquinoline scaffolds (**2**-D_4_ and **3**-D_4_) when compared to authentic synthesized standards (**Supplementary Fig. 3**). Unlike CET that was found throughout the plant body, **2** and **3** accumulated only in root tips (**Supplementary Fig. 1e-f**). Nonetheless, their D_4_-enrichment (**2**-D_4_ and **3**-D_4_) from dopamine-D_4_ feeding showed a similar pattern to that of CET-D_4_, which occurs exclusively in growing root tips and not in other tissues (**Supplementary Fig. 1g-h**). These results suggested that **2** and **3** could be intermediates of CET biosynthesis and that they could represent sequential metabolism since they differ only by an O-methylation. Meanwhile, untargeted analyses also identified CET-D_4_. Its relatively low fold change (3.8) compared to earlier potential intermediates could be attributed to its position further downstream in the pathway. Three additional mass signatures with large fold change and interesting *m/z* that are close to that of CET (**Fig. 2d**, red dots) were subsequently also found to be true intermediates of CET biosynthetic pathway (see **Fig. 4** and **Extended Data Figs. 9-10**), suggesting that feeding of early precursor (dopamine-D_4_ in this case) combined with untargeted analyses using LC-MS effectively revealed intermediates of interest.

To determine whether **2** is an intermediate of CET biosynthesis, different tissues of actively growing *C. harringtonia* were sectioned and soaked in a solution of **2**-D_4_ and analyzed with LC-MS in a manner similar to the dopamine-D_4_ feeding study. Relative isotope enrichment of CET-D_4_ in **2**-D_4_ feeding compared to water treatment in six different tissues further supports the growing root tips as a site of active biosynthesis of CET and that **2**-D_4_ serves as a pathway intermediate (**Fig. 2e**). Furthermore, unlike when dopamine-D_4_ was fed, **3**-D_4_ was observed in all different tissues when treated with **2**-D_4_. Nevertheless, O-methylation of **2**-D_4_ to **3**-D_4_ predominantly occurred in the growing root tips compared to other tissues (**Fig. 2f** and **Supplementary Fig. 1i**), following a similar pattern of high **3**-D_4_ enrichment in root tips from dopamine-D_4_ feeding (**Supplementary Fig. 1h**).

Since both dopamine and **2** are incorporated into **3** and appear to be involved in CET biosynthesis in root tips, we next sought to determine whether **3** is a direct intermediate in the CET biosynthetic pathway rather than a side-product or unrelated metabolite. Similar feed-in experiments and analyses done with dopamine-D_4_ and **2**-D_4_ were conducted with **3**-D_4_. Enrichment of CET-D_4_ from **3**-D_4_ feeding occurred only in growing root tips and not in other tissues sampled (**Fig. 2g** and **Supplementary Fig. 1j**), indicating that **3** serves as a true intermediate in CET biosynthesis and further supporting that a site of active biosynthesis of CET is growing root tips and not other tissues. Furthermore, our identification of **2** and **3** reveals direct evidence for CET biosynthetic intermediates beyond the tyrosine and phenylalanine precursors that had been identified in the 1970s^26–28^ and provides experimental evidence for the long-standing hypothesis that phenethylisoquinolines act as intermediates in CET biosynthesis^2,26^.

Because **2** and **3** were both found to be intermediates of CET, and phenethylisoquinolines—especially compound **1**—have long been hypothesized to be intermediates of CET^2,26^, we also searched for other possible phenethylisoquinolines such as **1**, **11**, **12**, and **13** to examine their biological relevance, and whether they could serve as potential intermediates in the CET pathway (**Supplementary Figs. 4-5** and **Supplementary Table 1**). Although natural accumulation of **1**^29^ and **11** were observed in root tips of *C. harringtonia*, we did not detect enrichment of **1**-D_4_ nor **11**-D_4_ following feeding of dopamine-D_4_ (**Supplementary Fig. 4**). Moreover, neither **12** nor **13** were detected anywhere in any tissues of *C. harringtonia* sampled (**Supplementary Fig. 5**). We also did not detect deuterium enrichment of **12** and **13** (**12**-D_4_ and **13**-D_4_) from feeding dopamine-D_4_. Feeding of **1**-D_4_ and **12**-D_4_ in different tissues sampled from *C. harringtonia* also did not result in enrichment of CET-D_4_ (**Supplementary Figs. 4-5**). These findings collectively suggested that CET is derived from **2**, which is formed through a Pictet-Spengler-like condensation of dopamine and 3-(4-hydroxy-3-methoxyphenyl)propanal (**Fig. 2h** and **Supplementary Table 1**). Compounds **1** and **11** are likely not on pathway to CET but represent additional alkaloids that accumulate in *Cephalotaxus* roots but were not actively made during the time of sampling.

Additionally, unlike CET, no enrichment of deuterated HHT (HHT-D_4_) was detected in any of the sectioned tissues of *C. harringtonia* fed with dopamine-D_4_, **2**-D_4_, or **3**-D_4_. This result may be interpreted in two ways: i) the feed-in assay failed to capture HHT biosynthesis despite root tips being the sole site of HHT production, or ii) the alkaloid core of HHT gets made only in roots and is transported to other parts of the plant, where side-chain biosynthesis and its attachment occur to produce HHT. The widespread availability of both CET and HHT in all different tissues likely support the latter model.

Collectively, the stable-isotope labeled precursor feeding studies with various tissues sectioned from an actively growing *C. harringtonia* revealed that the root tips are the primary site of CET alkaloid core biosynthesis, which does not align with the widespread accumulation pattern of CET and HHT observed throughout the plant. The biosynthetic pathway begins with tyrosine^26,27^ and phenylalanine^27,28^ and proceeds through the intermediates dopamine, **2**, and **3**. Compound **3** likely undergoes oxidative coupling to form compound **6** (a proposed intermediate with a mass signature identified via XCMS analyses; **Fig. 2d**), followed by a series of oxidations, including carbon excision, and reduction to ultimately produce CET (**Fig. 2h**). Our data suggest that the alkaloid core of HHT is biosynthesized and subsequently transported from root tips and distributed throughout the plant, where side-chains are attached when needed to produce the toxic ester-derivatives such as HHT^1,2,4^. The availability of downstream intermediates (**2** and **3**) exclusive to root tips further supports our model. Having identified **2** and **3** as intermediates in the pathway, we next sought to identify a candidate gene in biosynthesis that could be used as a bait gene to shortlist candidate CET biosynthetic enzymes.

### Paired transcriptomics and metabolomics analysis identifies *Ch*OMT-1 as the biosynthetic gene and reveals co-expressed candidate genes involved in CET biosynthesis

Transcriptomics with co-expression analyses has often been used in elucidating plant biosynthetic pathways when spatial, temporal, or condition-specific production of a compound of interest is known^31–34,39^. With the knowledge that a site of active CET biosynthesis is specific to root tips, we sought to use transcriptional analyses to identify CET biosynthetic enzymes. To do so, we first aimed to obtain a transcriptome of *C. harringtonia* that fully captures CET biosynthetic genes by generating our own *in-house* transcriptome paired with stable-isotope labeling studies on diverse tissues of actively growing *C. harringtonia*, harvested in May 2023. Each tissue sample was divided for parallel profiling (one for metabolomics and the other for transcriptomics), ensuring that the biosynthetically active root tip tissues were analyzed using RNA-seq. We also utilized PacBio Iso-Seq to avoid technical limitations of de novo transcriptome assembly, such as misassembled or fragmented contigs^39,40^, and obtain an accurate, full-length transcriptome that could be used as a reference for short-read RNA-seq (29,163 transcripts after processing). Illumina sequencing from multiple various plant tissues (a total of 36 biosynthetic and non-biosynthetic samples) was then used to quantify relative gene expression.

Using these paired metabolomic and transcriptomic datasets across diverse tissues of *C. harringtonia*, we sought to identify an initial enzyme in CET biosynthetic pathway that could be used as a bait gene for co-expression analyses to discover CET biosynthetic enzymes. With the knowledge that O-methylation of **2** to **3** is involved in CET biosynthesis in root tips, we proposed that the enzyme responsible for this transformation of **2** to **3** would be a good first candidate to be used as the bait gene.

Because O-methylation of **2** to **3** mainly occurs in the root tips of the plant and *S*-adenosylmethionine-dependent methyltransferase is a family of enzymes known for performing O-methylation^30,41^, we focused on transcripts that had expression in roots and their Pfam annotated as methyltransferase. We then prioritized our candidate gene list by conducting a Pearson’s correlation (*r* > 0.85) between transcript expression levels and **3**-D_4_ enrichment across all 15 tissues fed with **2**-D_4_. These analyses together resulted in a shortlist of 18 methyltransferase-annotated transcripts, whose expression strongly correlated with *in vivo* metabolic conversion of **2**-D_4_ to **3**-D_4_ (**Fig. 3a** and **Supplementary Fig. 6**).

**Figure 3.**
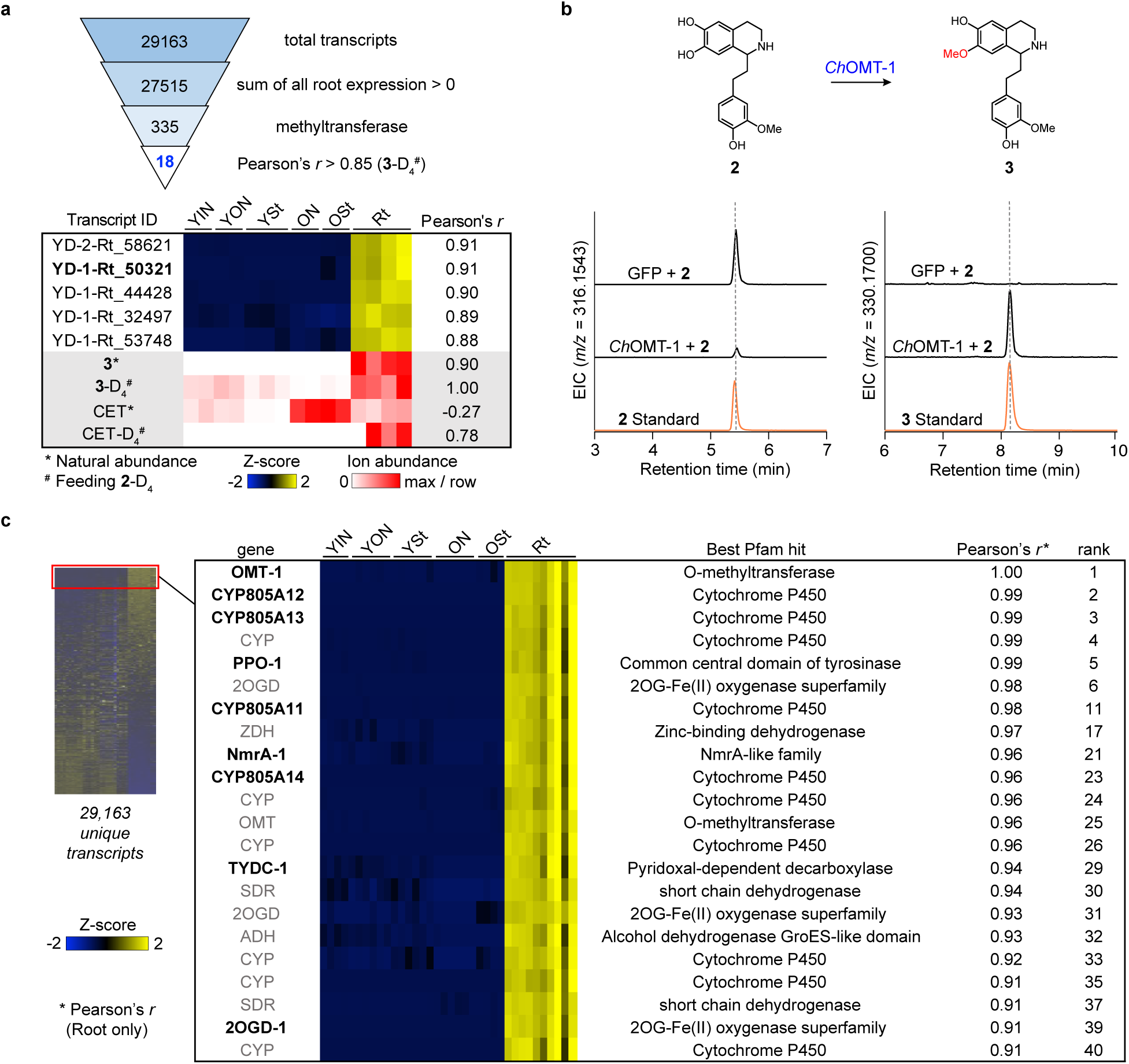
Combined transcriptomics and metabolomics analysis identifies *Ch*OMT-1 as the biosynthetic gene and reveals co-expressed candidate genes involved in CET biosynthesis. **(a)** The funnel plot shows the transcript selection process from 29,163 total transcripts to 18 candidate methyltransferases. The heatmap shows expression profiles (Z-score scaled) of the five (out of 18) tested methyltransferase-annotated transcripts in 15 samples of six different tissues of *C. harringtonia* from an independent experiment A conducted in May 2023, selected based on expression in root tips (site of active biosynthesis) and strong correlation with **3**-D_4_ enrichment following **2**-D_4_ feeding (Pearson’s *r* > 0.85). Transcripts are ordered by decreasing correlation, with YD-1-Rt_50321 (in bold) identified as *Ch*OMT-1. Below, a metabolomic heatmap (red scale) shows LC-MS ion abundances of **3** and CET at natural isotope abundance (*) and their four-deuterated analogs (^#^) after **2**-D_4_ feeding, measured in the same *C. harringtonia* tissues used for transcriptomic analysis (each tissue was divided for parallel profiling). Six different tissues (15 samples): 2 young inner needles (YIN), 2 young outer needles (YON), 3 young stems (YSt), 2 old needles (ON), 2 old stems (OSt), and 4 root tips (Rt). **(b)** Expression of *Ch*OMT-1 in *N. benthamiana* with co-infiltrated **2** leads to consumption of **2** (*m/*z 316.1543, r.t. of 5.4 min) and production of an O-methylated compound **3** (*m/z* 330.1700, r.t. of 8.1 min), as shown by the LC-MS chromatograms. This experiment was repeated more than three times, with similar results observed each time. Retention times of authentic standards (orange traces) are shown for comparison. Note: minimal retention time shifts (∼0.2-0.3 min) are often observed in LC-MS runs conducted at different times. **(c)** RNA-seq analyses across various tissue types—YIN, YON, YSt, ON, OSt, and Rt—combined with Pearson’s correlation using *Ch*OMT-1 as the bait gene, identify a set of strongly co-expressing biosynthetic gene candidates from 29,163 unique transcripts. Initial Pearson correlation analysis across all tissues using *Ch*OMT-1 as a query gene identified 316 transcripts (r > 0.98). As many of these showed expression only in root tips, a second round of correlation analysis was conducted using Rt samples alone, yielding 41 top candidates (r > 0.90). Out of 41, genes of biosynthetically relevant enzyme families such as oxidases, reductases, O-methyltransferases, and pyridoxal phosphate (PLP)-dependent enzymes are shown here, and those involved in CET biosynthesis are highlighted in bold. Expression values within the heat map are log_2_-transformed values of each transcript calculated as Trimmed mean of M (TMM)-normalized counts per million (CPM) in Z-score scale. Pearson’s *r* shown here are from the second round of correlation analysis done using only Rt samples. The whole heatmap is composed of 36 unique samples—YIN (4), YON (6), YSt (6), ON (6), OSt (4), and Rt (10)—harvested from the same plant but at two different times (one set, A, harvested in early May 2023 and the other set, B, two weeks after).

A batch of 5 out of the 18 that were initially cloned from *C. harringtonia* cDNA was tested for function using *Agrobacterium*-mediated transient expression in *N. benthamiana* leaves with direct co-infiltration of **2** as a substrate^30,42^. LC-MS analysis of infiltrated leaf extracts revealed that one candidate methyltransferase (YD-1-Rt_50321, highlighted in bold in **Fig. 3a**) consumed **2** and produced **3** (**Fig. 3b** and **Supplementary Fig. 7**). This O-methyltransferase responsible for O-methylation of **2** to **3** was referred to as *Ch*OMT-1.

*Ch*OMT-1 was then utilized as the bait gene to identify a set of strongly co-expressing CET biosynthetic gene candidates from 29,163 unique transcripts. Initial Pearson correlation analysis across all 36 tissue samples using *Ch*OMT-1 as the query gene identified 316 highly co-expressed transcripts (*r* > 0.98). As many of these accumulate primarily in root tips and are not detected in other tissues, a second round of correlation analysis (*r* > 0.90) was conducted using Rt samples alone, enabling prioritization of 41 top candidates (**Fig. 3c** and **Supplementary Fig. 8**). This top-ranking candidate list is rich in genes that are generally associated with secondary metabolism such as cytochrome P450s (CYPs)^30,32,43^, Fe(II)/2-oxoglutarate-dependent dioxygenases (2OGDs)^39,44,45^, and dehydrogenase/reductase enzymes^46,47^, suggesting that it contains CET biosynthetic enzymes responsible for oxidations and reductions that are required to form CET from **3**. Furthermore, the list contains early CET biosynthetic genes that encode enzymes, which have been previously shown to generate dopamine from tyrosine (*Ch*TyDC-1 and *Ch*PPO-1)^29^.

### Discovery and engineering of *Cephalotaxus* alkaloid biosynthesis

With a shortlist of candidate CET biosynthetic genes from co-expression analysis (**Fig. 3c**) and unique masses of the next top two potential intermediates (**6** and **5**) after **2** and **3** identified from untargeted metabolite analysis (XCMS) of stable-isotope precursor feeding study (**Fig. 2d**), we next sought to elucidate the downstream biosynthetic pathway of CET after **3**. To eventually produce CET, we hypothesized that **3** would go through a series of oxidations such as oxidative coupling of two methoxyphenols to form **6** and carbon excision from the lower six-membered ring C to form the five-membered ring C’ of CET (**Fig. 2h** and **Fig. 4b**).

**Figure 4.**
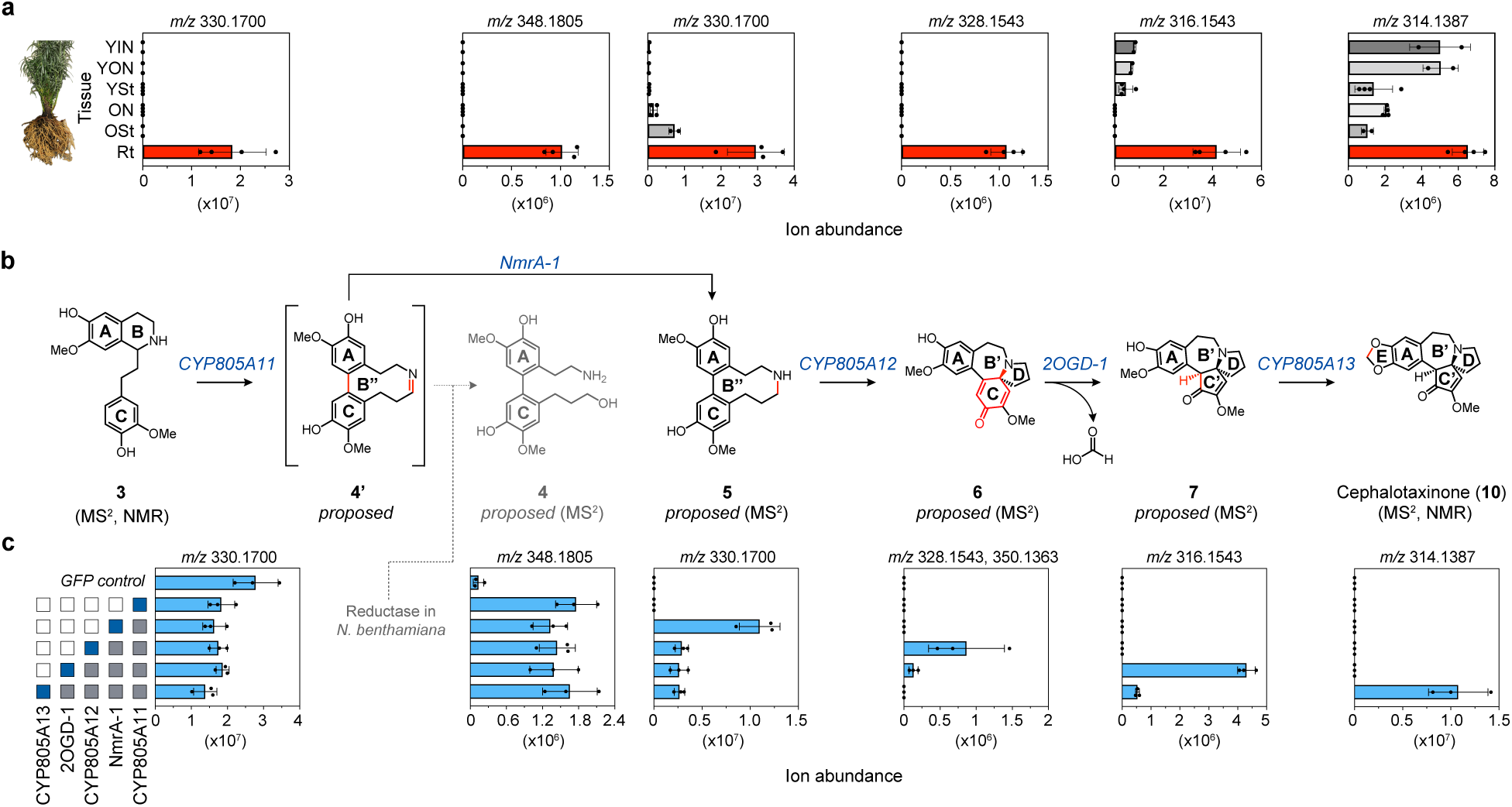
Discovery of a pathway for CET alkaloid biosynthesis, from phenethylisoquinoline (3) to cephalotaxinone. **(a)** Natural accumulation of CET biosynthetic intermediates in the native plant, *C. harringtonia*, harvested in May 2023. The data are reported as mean ± SD of the extracted ion abundance (*n* = 4 biological replicates for YSt, ON, and Rt, and *n* = 2 for YIN, YON, and OSt) for the exact ion mass [M + H]^+^ corresponding to each compound. For quantification of cephalotaxinone, a 200-fold dilution was performed to the extracted samples to avoid saturation of the parent isotope peaks in LC-MS. Young inner needles (YIN), young outer needles (YON), Young stems (YSt), old needles (ON), old stems (OSt), and root tips (Rt). **(b)** Proposed biosynthetic pathway of CET reconstituted in *N. benthamiana* through transient co-expression of five identified biosynthetic genes from *C. harringtonia*, illustrating the stepwise conversion from compound **3** to cephalotaxinone (**10**). All the structures are supported by LC-MS/MS analyses, with the first and final intermediates (**3** and cephalotaxinone) further supported by authentic synthesized and purified standards, including NMR analyses. **(c)** Gray boxes to the left of the graphs indicate biosynthetic genes that are co-expressed in *N. benthamiana* leaves; blue box represents the final acting enzyme. For each intermediate, the data are reported as mean ± SD of the extracted ion abundance (*n* = 3) for the exact ion mass [M + H]^+^ (for **6**, both [M + H]^+^ and [M + Na]^+^) corresponding to each compound. While the overall biosynthetic pathway of CET is linear as shown, the final steps involving *Ch*CYP805A13 and *Ch*2OGD-1 may occur in bifurcated sequence (Extended Data Fig. 8).

We speculated that **3** first undergoes oxidative transformation: hydroxylation, oxidative coupling, or both. All the oxidases (10 CYPs and 3 2OGDs)^32,45,48–50^ in the shortlist of candidate genes (**Fig. 3c**) were each cloned and tested by *Agrobacterium*-mediated transient transformation in *N. benthamiana* with **3** co-infiltrated as a substrate. However, LC-MS analysis of leaf extracts revealed that none of the candidates resulted in additional hydroxylation ([M + H]^+^ = *m/z* 346.1649), oxidative coupling ([M + H]^+^ = *m/z* 328.1543), or both ([M + H]^+^ = *m/z* 344.1492) of **3**. However, one of the tested enzymes (*Ch*CYP805A11) resulted in partial consumption of **3**. We therefore conducted untargeted metabolite analysis (XCMS, see Methods) comparing *Ch*CYP805A11-expressed leaves to GFP-expressed leaves (negative control), both co-infiltrated with **3**, to examine if any relevant unique mass with a significantly increasing fold change (Δ > 5) in abundance between the two conditions (*P* < 0.1) appeared (**Extended Data Fig. 1**). XCMS analysis reported 27 unique masses, of which 7 are above the signal to noise threshold and can be identified in the raw LC-MS data. These seven (**Extended Data Fig. 1d**, red dots) all represent isotopologues or ions of a single compound ([M + H]^+^ = *m/z* 348.1805, r.t. = 7.07 min) that corresponds to a single hydroxylation and reduction of **3**. Based on the MS/MS analyses of both the molecule and its four-deuterated form from **3**-D_4_ feeding in root tips of *C. harringtonia* experiment, the compound was assigned the structure of **4** (**Extended Data Fig. 1**). These results suggested that *Ch*CYP805A11 catalyzes oxidative coupling of two methoxyphenols of **3** (rings A and C), forming **4’** (**Fig. 4**). However, due to its unstable imine ring, **4’** likely undergoes imine hydrolysis to form **4’’** that contains a very reactive aldehyde moiety, which then gets reduced by a promiscuous reductase in *N. benthamiana* to form **4** (**Extended Data Fig. 1h**). The instability of **4’** could explain why this molecule was not detected in LC-MS but instead, a mass corresponding to a single hydroxylation and reduction of **3** (**4**, [M + H]^+^ = 348.1805 Da) was observed. This observation was consistent with metabolites observed in the native plant *C. harringtonia*, indicating that the promiscuous reductase is not specifically present only in *N. benthamiana* but also in other plants in general.

To form the fused bicyclic backbone of CET (rings B’ and D), it has been speculated that a precursor similar to **5**, formed through oxidative coupling of phenethylisoquinoline, undergoes an intramolecular cyclization via nucleophilic attack by the nitrogen atom^2,51,52^, indicating that the ring B’’ of **4’** needs to be preserved (**Fig. 4b**). We thus hypothesized that there exists a reductase in CET biosynthesis for reducing the imine bond of **4’** to form the stable compound (**5**) and that **4** is not a direct intermediate but a side-product of CET biosynthesis. To evaluate our hypothesis, all five dehydrogenases/reductases^46,47,53,54^ in the shortlist of candidate genes (**Fig. 3c**), including an atypical short-chain dehydrogenase (SDR)—nitrogen metabolite repression regulator (NmrA)-like family enzyme^55–59^ (*Ch*NmrA-1) (sequence analyses of homologs, **Supplementary Fig. 9**), were each cloned and co-expressed with *Ch*CYP805A11 in *N. benthamiana* leaves using *Agrobacterium*-mediated gene delivery, with **3** co-infiltrated as a substrate. LC-MS analysis of the *Ch*NmrA-1 infiltrated leaf extracts revealed partial reduction of **4** and production of a new peak ([M + H]^+^ = *m/z* 330.1700, r.t. slightly left to that of **3**) (**Extended Data Fig. 2**). This new peak corresponded to the compound with the fourth largest fold-change in earlier XCMS analysis comparing dopamine-D_4_-fed root tips with water-fed controls of *C. harringtonia* (**Fig. 2d**), suggesting that it is likely to be an intermediate of CET biosynthesis. MS/MS analyses of both the new molecule and its four-deuterated form were used to assign the structure as **5** (**Extended Data Fig. 2**). Collectively, the results support our hypothesis that **3** goes through a *Ch*CYP805A11 mediated oxidative coupling to form an unstable intermediate **4’**, which gets reduced by *Ch*NmrA-1 to produce **5**.

To further verify the functions of *Ch*CYP805A11 and *Ch*NmrA-1, *in vitro* assays with yeast microsomal protein fractions of *Ch*CYP805A11 and purified *Ch*NmrA-1 were conducted (**Extended Data Fig. 3**). Unlike in *N. benthamiana*, production of **4** was not detected in *in vitro* assays with purified yeast microsomes containing *Ch*CYP805A11 and incubated with **3**. Aside from trace amounts of **5**—potentially arising from non-enzymatic reduction in the presence of excess NADPH, we did not see any single new product based on untargeted metabolite analysis comparing purified yeast microsomes of *Ch*CYP805A11 to empty vector microsomes (negative control) in the absence of *Ch*NmrA-1, despite near-complete consumption of **3**. However, when both the microsomes of *Ch*CYP805A11 and purified *Ch*NmrA-1 were combined with **3** *in vitro*, production of **5** was observed by LC-MS. Moreover, treatment of *Ch*CYP805A11 microsomes incubated with **3** followed by NaBD_3_CN (an imine reducing agent)^60^, in place of active *Ch*NmrA-1, resulted in formation of **5**-D (**Extended Data Fig. 3h-k**). These findings suggest that *Ch*CYP805A11 acts on **3**, producing an unstable intermediate, proposed to be **4’**, which can be reduced enzymatically by *Ch*NmrA-1 to form **5** (see **Supplementary Fig. 10** for proposed mechanism) or chemically by NaBD_3_CN to form **5**-D. Thus, the likely direct product of *Ch*CYP805A11 and the substrate for *Ch*NmrA-1 is **4’** rather than **4** (a reduced form of the aldehyde derived from hydrolysis of **4’**, proposed to form through promiscuous plant reductase).

As discussed earlier, we next speculated that **5** goes through an oxidative intramolecular cyclization via nucleophilic attack by the nitrogen atom to produce the fused bicyclic backbone of CET^2,51,52^ (**Fig. 4b**). Therefore, the rest of 9 CYPs and 3 2OGDs in the shortlist of candidate genes (**Fig. 3c**) were each tested in combination with *Ch*CYP805A11 and *Ch*NmrA-1 in *N. benthamiana* leaves with co-infiltrating **3** as a substrate. Addition of *Ch*CYP805A12 resulted in consumption of **5** and production of a new peak, proposed to be **6** based on its MS/MS fragmentation pattern (**Extended Data Fig. 4**). This was the first intermediate identified containing a tertiary amine and a bicyclo[5.3.0]decane core (rings B’ and D) that are crucial to the structural backbone of CET. Compound **6** also corresponded to the molecule with third largest fold-change in earlier XCMS analysis comparing dopamine-D_4_-fed root tips with water-fed controls of *C. harringtonia* (**Fig. 2d**), indicating that LC-MS-based untargeted analyses of early precursor feeding studies in dissected tissues of a native plant could be used as an effective tool to identify biosynthetic intermediates.

To produce CET from **6**, a carbon excision from ring C and formation of methylenedioxy bridge in ring E are necessary (**Fig. 4b**). Because CYPs and 2OGDs have been known to catalyze such unusual oxidative transformations^39,48,49,61,62^, the remaining 8 CYPs and 3 2OGDs in the shortlist of candidate genes (**Fig. 3c**) were again each tested in the transient expression system, producing **6**, and analyzed using LC-MS/MS. Interestingly, each *Ch*2OGD-1 and *Ch*CYP805A13 resulted in consumption of **6** and formation of different compounds, **7** (**Extended Data Fig. 5**) and **8** as well as **9** (**Extended Data Fig. 6**) respectively based on their MS/MS analyses. Co-expression of both enzymes with *Ch*CYP805A11, *Ch*NmrA-1, and *Ch*CYP805A12 in *N. benthamiana* leaves co-infiltrated with **3**, led to consumption of **7** and **8** (including **9**) and production of a new peak ([H+M]^+^ = *m/z* 314.1387) that was confirmed to be cephalotaxinone (**10**), a *Cephalotaxus* alkaloid^1,17^, using LC-MS/MS comparison to a purified cephalotaxinone standard (**Extended Data Fig. 7**). The ability to convert **6** ([H+M]^+^ = *m/z* 328.1543) to **7** ([H+M]^+^ = *m/z* 316.1543), and **8** ([H+M]^+^ = *m/z* 326.1387) to cephalotaxinone ([H+M]^+^ = *m/z* 314.1387)—both involving a 12 Da loss corresponding to one carbon—demonstrates that *Ch*2OGD-1 is responsible for performing the remarkable carbon excision reaction that contracts ring C to C’, a key step in the biosynthesis of CET (**Extended Data Figs. 5 and 7**).

Moreover, conversion of **6** to **8** and **7** to cephalotaxinone illustrated that *Ch*CYP805A13 catalyzes methylenedioxy bridge formation to generate ring E of CET (**Extended Data Figs. 6-7**). When *Ch*CYP805A13 acted on **6**, it produced not only **8** but also **9**, a side-product likely produced by a promiscuous reductase in *N. benthamiana* through reduction of a transient intermediate **9’** that likely arises from reversible ring opening of the tertiary amine in **8** (**Extended Data Fig. 6m**). Formation of **9** is unlikely to result from direct oxidation of **5** by *Ch*CYP805A13, as **9** was not detected in the absence of *Ch*CYP805A12 (**Extended Data Fig. 8**).

The five enzymes in *C. harringtonia* identified through co-expression analysis using *Ch*OMT-1 as the bait gene establish a biosynthetic pathway of CET from **3** to cephalotaxinone (**Fig. 4** and **Extended Data Fig. 8**). The order of the pathway was validated through sequential addition of each enzyme (**Extended Data Fig. 8a-b**) and dropout experiments in which each individual enzyme was removed from the engineered pathway to cephalotaxinone (**Extended Data Fig. 8c**). The sequential accumulation of proposed pathway intermediates with no detection of alternative mass signatures also supported that the overall biosynthetic pathway of CET from **3** to cephalotaxinone is linear, with the final steps involving *Ch*2OGD-1 and *Ch*CYP805A13 may occur in bifurcated sequence (**Extended Data Fig. 8**).

Reconstitution of CET biosynthetic pathway from **2** to cephalotaxinone in *N. benthamiana* was also established with co-expression of all the six newly discovered biosynthetic genes in this study and co-infiltration of **2** (**Extended Data Fig. 9**). Furthermore, the identities of all the newly discovered intermediates (**2**, **3**, **5**, **6**, **7**, **8**, and cephalotaxinone) and side-products (**4** and **9**) produced heterologously in *N. benthamiana* were supported by co-elution with metabolites found in *C. harringtonia* root tips that exhibited identical MS/MS fragmentation patterns (**Extended Data Fig. 10**). This indicated that the compounds produced in the *N. benthamiana* expression system are biologically relevant pathway intermediates. In addition to the authentic synthetic or purified standards used to confirm the structures of **2**, **3**, and cephalotaxinone, structural assignments of other proposed intermediates were all supported by LC-MS/MS analyses of the molecules and comparison of their MS/MS fragmentation patterns to those of the corresponding four-deuterated forms from dopamine-D_4_, **2**-D_4_, or **3**-D_4_ feeding study or to those of the related CET biosynthetic intermediates (**Extended Data Figs. 1-10**).

Moreover, all the identified intermediates were only observed at high levels in growing root tips of *C. harringtonia* relative to other tissues (**Fig. 4a** and **Extended Data Fig. 9c**). The exception is cephalotaxinone, which is found throughout the plant body. This accumulation of CET intermediates in growing root tips aligned with our findings of the core biosynthesis of HHT (CET) occurring at the root tips. However, the wide distribution of not only CET and HHT (**Fig. 2b**) but also cephalotaxinone (**Fig. 4a**) throughout the native plant as well as reversible conversion between cephalotaxinone and CET from early radioisotope labeling studies^27^ suggests that it is cephalotaxinone that gets primarily made in the growing root tips and is transported throughout the plant. Thus, reduction of cephalotaxinone to CET most likely occurs not only in root tips but also in other tissues, including the aerial part of the plant (**Fig. 5**). Furthermore, in D_2_O labeling studies using excised branches of *C. harringtonia*, there was no detectable enrichment of deuterated cephalotaxinone in needles. In contrast, there was a clear enrichment of deuterated CET and HHT in young needles in these experiments (**Supplementary Fig. 11**).

**Figure 5.**
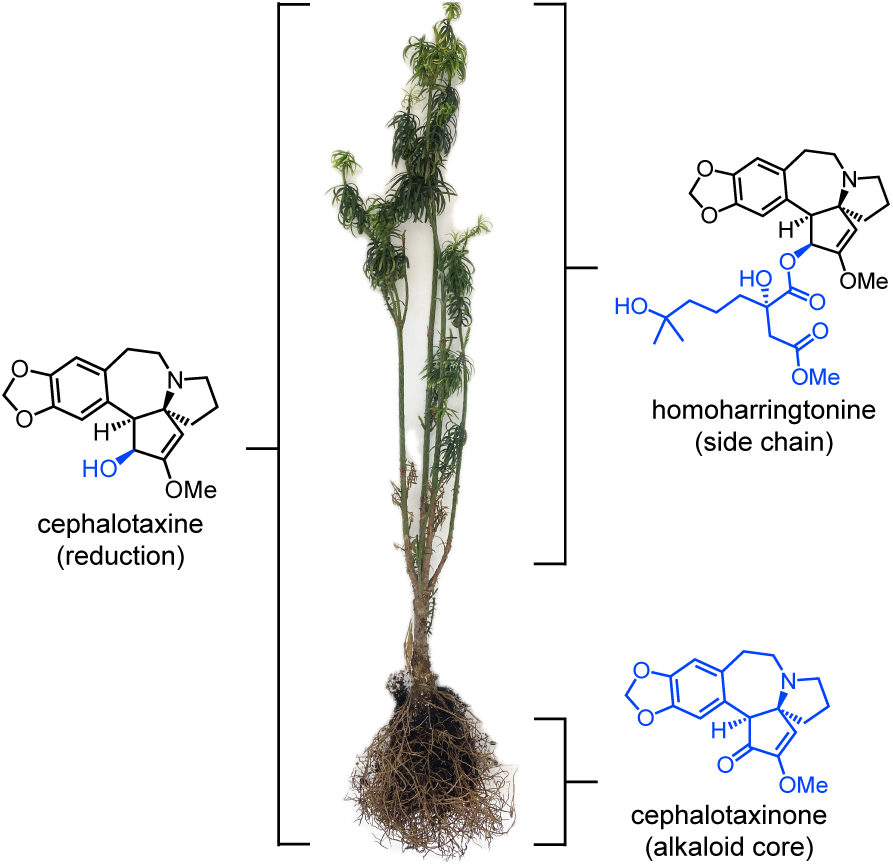
Proposed model illustrating sites of active metabolism for whole plant coordination of *Cephalotaxus* alkaloid biosynthesis. Proposed tissue specific metabolism is indicated in the parenthesis. Molecular components that are proposed to be actively produced, based on stable-isotope labeled precursor feeding and D_2_O labeling studies, are highlighted in blue.

Additionally, enzymes previously characterized in our laboratory that contribute to HHT side-chain biosynthesis, including *Ch*IPMS2 and *Ch*IPMI_SSU2^36^ (corresponding to YD-1-Rt_transcript/23102.p1 and YD-1-Rt_transcript/44947.p1, respectively), are expressed throughout the plant, with relatively high expression in young needles, young stem, and root tips. These data together support a model (**Fig. 5**) in which the cephalotaxinone alkaloid core is primarily biosynthesized in root tips and subsequently transported throughout the plant body for active reduction to CET (**Supplementary Table 2**). Subsequent side chain tailoring to complete HHT biosynthesis likely occurs in the aerial part of the plant, especially young needles (based on isotope labeling and transcript abundance), and potentially also in root tips (based on transcript abundance).

Collectively, we have elucidated a biosynthetic pathway to cephalotaxinone. We discovered six novel biosynthetic enzymes that perform unique transformations, seven pathway intermediates, and two side-products in CET biosynthesis from **2** to cephalotaxinone, which is primarily produced in growing root tips of *C. harringtonia* and then gets transported throughout the plant for conversion into CET and HHT.

Despite these advances, future investigations are still needed to fully understand the final step of cephalotaxinone to CET and clarify the Pictet-Spengler like reaction to produce the phenethylisoquinoline scaffold in CET biosynthesis. We tested a number of candidate enzymes, including dehydrogenases and reductases, prioritized based on co-expression with the six CET biosynthetic genes that are highly expressed only in root tips of *C. harringtonia* (**Supplementary Fig. 8**), for conversion of cephalotaxinone to CET. However, none of them resulted in production of CET. It is possible that the downstream enzyme(s) responsible for reducing cephalotaxinone to CET may have a significantly different expression pattern relative to early CET biosynthetic genes.

With respect to the formation of the phenethylisoquinoline scaffold in CET biosynthesis, we note that a previous report had described a Bet v 1-like protein (*Ch*PSS) that catalyzes a Pictet-Spengler like reaction in the biosynthesis of compound **1**^29^. Our data support that compound **2** is the likely pathway precursor (**Fig. 2**), and we also were curious about which enzyme(s) enable biosynthesis of this initial phenethylisoquinoline scaffold. We tested a series of putative Bet v 1-like proteins identified through co-expression analysis with *Ch*OMT-1 and a homolog of *Ch*PSS^29^ (corresponding to YD-1-Rt_transcript/53915.p1 in our transcriptome with two aa difference). However, these enzymes—heterologously expressed in *N. benthamiana* and assayed using crude leaf protein lysates *in vitro*—do not appear to promote condensation of dopamine with the requisite aldehydes to produce either compound **2** or **1** above slow background levels of condensation (**Supplementary Fig. 12**). In contrast, the previously characterized *Cj*NCS^63^ was active in this assay. It is possible that a different enzyme class has evolved in *C. harringtonia* to catalyze this scaffold forming step that remains to be discovered.

## DISCUSSION

Our study demonstrates careful regulation and coordination of biosynthesis of CET, the core scaffold of the ribosome inhibitor HHT, by the plant. *Cephalotaxus* species are slow-growing conifers that follow a seasonal growth cycle, with bud expansion beginning in late March, shoot emergence in late April, and rapid shoot elongation occurring throughout May, followed by winter dormancy^16,37,38^. Our stable-isotope labeled precursor feeding experiments with sectioned tissues of *C. harringtonia* showed that this seasonal cycle strongly influences CET biosynthesis. In tissues collected during active shoot growth in May, incorporation of dopamine-D_4_ and **2**-D_4_ into CET-D_4_ in root tips was observed, whereas sectioned tissues from dormant plants harvested in November and December showed no enrichment of CET-D_4_ (**Supplementary Table 3**). Furthermore, our *in-house* transcriptome generated from tissues of actively growing *C. harringtonia* collected in May 2023 captured all CET biosynthetic transcripts, whereas our *in-house* transcriptome generated from tissues harvested after the spring growth period (August 2018) had all the CET biosynthetic transcripts missing (**Supplementary Fig. 8**). Although potential limitations of feeding assays and technical errors in RNA sampling from August 2018 cannot be entirely excluded, the consistent detection of CET-D_4_ enrichment in root tips of an actively growing *C. harringtonia* harvested in May 2022, May 2023, and June 2024, together with transcriptome data from May 2023, supports that CET biosynthesis is seasonally activated during active spring growth, particularly in May and June.

Our findings further illustrate strategic coordination of *C. harringtonia* to produce *Cephalotaxus* alkaloids through whole plant coordination metabolism. The stable-isotope labeled precursor feeding studies and metabolite profiling of various tissues of actively growing *C. harringtonia* revealed that the root tips are the primary site of CET alkaloid core biosynthesis, contrasting with the widespread accumulation pattern of cephalotaxinone, CET, and HHT observed throughout the plant (**Fig. 2** and **Fig. 4a**). The non-toxic cephalotaxinone^1,64^ gets synthesized in the growing root tips and transported throughout the plant in an inert form, which can later be reduced to CET^27^—an apparent non-toxic precursor^2,4^ poised for esterification with different side-chains to generate toxic defense compounds such as HHT^1,2,4^ when and where needed, including young needles (**Fig. 5**). Together, our findings in *C. harringtonia* highlights a model in which a plant employs whole-organism metabolic coordination to carefully produce and store toxic defense metabolites, thereby ensuring effective defense while minimizing self-toxicity and unnecessary energy expenditure.

It is notable that HHT is one of the few known plant-derived small molecule NPs that inhibit eukaryotic protein translation, whereas most known plant-derived inhibitors of this process are proteins^65,66^. Compared to prokaryotes, far fewer eukaryotic translation inhibitors are known in plants^13,65^. Notable plant NPs other than HHT that act as eukaryotic protein translation inhibitors by binding to the large ribosomal subunit include ricin, a protein from castor beans^67^, as well as lycorine and narcisclasine, small molecules from *Amaryllidaceae* family^13,65^. Interestingly, *Amaryllidaceae* species have been shown to exhibit spatiotemporal regulation of the biosynthesis of ribosome inhibitor molecules like lycorine^32^, which may represent a strategy to mitigate potential autotoxicity—similar to the coordinated CET/HHT biosynthetic pathway observed in *Cephalotaxus*.

Our study elucidates the long-elusive biosynthetic pathway to cephalotaxinone—the core alkaloid scaffold of an FDA-approved anti-cancer drug HHT. Stable-isotope labeled precursor feeding, integrated with tissue-resolved metabolite profiling and transcriptomic co-expression analysis, enabled us to identify growing root tips as a site of active HHT alkaloid core biosynthesis, discover seven CET pathway intermediates, and functionally characterize six novel biosynthetic enzymes. Leveraging this paired, *in-house* generated metabolomic and transcriptomic dataset across diverse tissues of actively growing *C. harringtonia* was critical in fully capturing the spatiotemporally expressed CET biosynthetic genes and identifying co-expressed genes in biosynthetically active tissues, demonstrating a powerful framework for pathway discovery in genetically intractable, slow-growing non-model plants. Furthermore, our findings suggest that *C. harringtonia* employs a sophisticated whole-plant coordination strategy to produce the toxic ribosome inhibitor HHT, offering deeper insights into how certain plants regulate the biosynthesis of eukaryotic ribosomal toxins as a defense mechanism while minimizing autotoxicity. Overall, our work not only establishes a metabolic route to the core scaffold of HHT, enabling future sustainable, large-scale production of this clinically essential drug, but also expands our understanding of whole-plant coordination for biosynthesis of specialized metabolites and the unique biochemical transformations catalyzed by plant enzymes to generate these complex, bioactive molecules.

## METHODS

### Chemicals and reagents

All biological and chemical reagents were purchased from commercial vendors, unless stated otherwise. These include dopamine (MilliporeSigma), 3-hydroxy-4-methoxyphenethylamine (Combi-Blocks), 3-methoxy-4-hydroxyphenethylamine (Combi-Blocks), 3-(4-hydroxyphenyl)propanal (Chemscene), 3-(3,4-dihydroxyphenyl)propanal (Astatech), 3-(4-hydroxy-3-methoxyphenyl)propanal (Astatech), dopamine-D_4_ (1,1,2,2-D_4_; Cambridge Isotope Laboratories), cephalotaxine (Selleckchem), and homoharringtonine (MilliporeSigma).

### Plant growth

The *Cephalotaxus harringtonia* var. Fastigiata plants used to measure natural abundance of their secondary metabolites and to conduct stable isotope labeled precursor feeding experiments as well as D_2_O labeling studies were purchased from Forest Farms (forestfarm.com). They were grown in pots with soil, watered approximately once every week. Growth conditions consisted of ambient laboratory temperature and lighting.

The *N. benthamiana* plants used for heterologous gene expression were sown and grown in PRO MIX HP Mycorrhizae soil (Premier Tech Horticulture) on growth shelves with a 16/8-h light/dark cycle at ambient laboratory temperature. 4-5 weeks old plants after seeding were used for *Agrobacterium*-mediated transformation.

### Stable isotope labeled precursor feeding experiments to identify the site of active biosynthesis of CET

Various tissues from *C. harringtonia* var. Fastigiata were excised from the whole plant using ethanol-sterilized scissors. For young inner needles (YIN), actively growing inner young light-green yellowish needles of 0.5-2 cm in length were harvested. Young outer needles (YON) were actively growing outer light-green needles, 1.5-3 cm in length. For young stems (YSt), light-green stems with YIN and YON of 2-3 cm in length from the tip of the stem (meristem) was harvested; all the surrounding needles were removed using a sterile scalpel. Old needles (ON) were mature, dark green needles (≥1-2 years old), 4-7 cm in length. For old stems (OSt), at least 1-2 year-old mature dark green stems (4-5 cm in length) with ON removed were collected. Before harvesting root tips (Rt), they were thoroughly washed with water, quickly rinsed with 30% ethanol, and washed again with water as a final washing step. Then, roots of 5-8 cm in length from the root tip, including the root meristem, were collected. Needles and roots, due to their thin structure, were latitudinally split into equally sized sections for paired metabolomics (feeding experiments) and transcriptomics (RNA-seq). Stems were longitudinally sectioned into equal halves for the same paired analyses.

Tissues used for accumulated metabolite analysis and RNA extraction were immediately flash frozen in liquid nitrogen and stored at -80 °C until further processing. For feeding experiments, sterile 12-well plates were filled with 2 mL of an aqueous solution of 100 µM dopamine-D_4_, **1**-D_4_, **2**-D_4_, **3**-D_4_, **12**-D_4_, or merely deionized water (negative control); see **Supplementary Table 3** for details on when and which experiments were conducted. Each well contained 7-9 needles for YIN, 3 needles for YON, 1 stem for YSt, 3 needles for ON, 1 stem for OSt, and 4-5 roots for Rt, with all tissues cut into smaller pieces to maximize direct contact with the substrate solution. For Rt, the 12-well plates were wrapped in aluminum foil to simulate the darkness experienced by roots growing underground. The assay was conducted for five days, with the solution replaced by freshly prepared substrate solution halfway through (after ∼2.5 days). Following the remaining ∼2.5 days, sectioned tissues were collected into pre-weighed 2 mL Eppendorf Safe-Lock tubes, flash frozen in liquid nitrogen, and lyophilized to dryness. Samples were then processed for analysis using LC-MS (see *Metabolite extraction from plant material and LC-MS analysis of metabolites* in Methods). All feeding experiments were performed on tissues from the same plant.

### D_2_O labeling studies and deuterium enrichment analysis

D_2_O labeling study with *C. harringtonia* var. Fastigiata was conducted much as previously described^35^. Briefly, two branches (approximately 7 cm in length) were excised from the base of *C. harringtonia* var. Fastigiata using ethanol-sterilized scissors and placed cut end down in a Falcon tube containing 10 mL of either 100% deionized water (control) or 10% D_2_O in deionized water (prepared from 100% D_2_O; Sigma-Aldrich). Throughout the experiment, the respective deionized water was replaced every 3-4 days. The caps were placed on the tubes such that airflow could occur. Growth conditions consisted of ambient laboratory temperature and lighting. Both young needles (three needles per sample) at the top end of both branches, which were determined based upon having a lighter green hue, softer needles, and smaller size compared to the rest of the old mature needles, and old needles (two needles per samples) were harvested from both branches treated with either D_2_O or H_2_O after 5, 10, and 15 days of incubation. Three samples were collected at each time point, except the control for old needles that had two samples collected at each time point. All samples were collected into pre-weighed 2 mL Eppendorf Safe-Lock tubes, flash frozen in liquid nitrogen, and lyophilized to dryness. Samples were then processed for analysis using LC-MS (see *Metabolite extraction from plant material and LC-MS analysis of metabolites* in Methods).

The relative isotope enrichment of *Cephalotaxus* alkaloids, including cephalotaxinone, CET, and HHT, were calculated much as previously described^35^. Briefly, mass isotopologue distributions (MIDs) were obtained by summing MS signal intensities over retention time windows of interest using the **get_MID.py** script from https://github.com/Stanford-ChEMH-MCAC/d2o_metabolomics^35^. From MIDs, isotopic enrichment (IE) was quantified by the following formula:

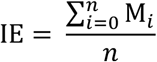

where IE is total heavy isotopic enrichment in the compound, *n* is the number of (integer) mass isotopologues that can be detected, and M_i_ is the relative abundance of the *i*th isotopologue^35^. Then, for the deuterium enrichment quantitation (the relative isotope enrichment), the IE value of a deuterium-labeled sample was subtracted from the IE value of an unlabeled control sample, correcting for the natural abundance of heavy isotopologues—mainly from ^13^C.

### RNA extraction

Flash-frozen *C. harringtonia* tissues (from May 2023 paired feeding experiments) were homogenized to a fine powder under liquid nitrogen using a chilled mortar and pestle that were pre-treated with RNAse Zap RNAse Decontamination Solution (Thermo Fisher Scientific). RNA from the powdered tissue samples were extracted using RNeasy Plant Mini Kit (Qiagen) with on-column DNase digestion. RLC buffer was used as the lysis buffer. For root samples, the extracted RNA was further cleaned up using the same kit. RNA concentration and purity were measured using a NanoDrop and Qubit (Qubit® 2.0 Fluorometer), and quality was further assessed using an Agilent 2100 Bioanalyzer.

### Preparation and sequencing of Illumina NovaSeq 6000 libraries and processing

Library preparation for Illumina NovaSeq 6000 (S4 Flow cell – PE150) sequencing was performed using the NEBNext® Ultra™ II RNA Library Prep Kit for Illumina® (non-directional; New England Biolabs), as per the manufacturer instructions. Individual libraries were barcoded with internal index sets at Novogene. For fragmentation, Diagenode Bioruptor Pico was used, and inserts were size selected with NEBNext beads to have an average insert size of 150-200 bp. The PCR products were purified (AMPure XP system). The quality and size distribution of each library were confirmed with analysis using a High Sensitivity DNA chip on a 2100 Bioanalyzer (Agilent). Each library was quantified using Qubit (Qubit® 2.0 Fluorometer) and real-time PCR (Kapa). Equimolar amounts of libraries for each sample (36 total, from May 2023 feeding experiments) were pooled for paired-end sequencing within a single Novaseq 6000 lane (S4 Flow cell – PE150).

The quality of raw Illumina sequencing reads was evaluated using FastQC (Babraham Bioinformatics, Babraham Institute). These reads were processed to remove adapters, barcodes, and low-quality sequences using Trimmomatic^68^, using the following parameters: <ILLUMINACLIP:TruSeq3-PE-2.fa:2:30:10 LEADING:5 TRAILING:5 SLIDINGWINDOW:4:5 MINLEN:50>. The trimmed reads were then re-assessed with FastQC to confirm quality.

While the processed Illumina sequencing reads were used to quantify each transcript abundance against PacBio Iso-Seq transcriptome, we also generated a *de novo* transcriptome assembly from Illumina reads using Trinity^69^ with default settings to leverage the higher sequencing depth and broader range of tissues provided through Illumina sequencing, resulting in generation of 541,990 contigs. Quality of the newly assembled transcriptome was assessed by mapping Illumina short reads to the transcriptome using bowtie2^70^ and through Benchmarking Universal Single-Copy Orthologs (BUSCO) analysis for completeness^71^. Then, peptide sequences, with longest open reading frame (ORF) identified for each transcript, were predicted using Transdecoder^69^, which resulted in 144,299 contigs. Redundant contigs were removed using CD-HIT-EST^72^ with 99% identity threshold, yielding a final Illumina *de novo* transcriptome of 102,958 contigs.

### Preparation and sequencing of PacBio Iso-Seq libraries and processing

Preparation of PacBio Iso-Seq libraries were performed as previously described^39^. For Iso-Seq, only a subset of tissues from the 36 total samples was sequenced: Rt1 (root tips), Rt6 (root tips), an equal combination of Rt4 and Rt8 (root tips), an equal combination of YIN3 and YIN4 (young inner needles), an equal combination of YSt3 and OSt3 (young and old stems), and an equal combination of ON1 and ON2 (old needles). Inclusion of Rt1, Rt,6 and Rt8—tissues with active CET biosynthesis—ensured that the transcriptome contained CET biosynthetic genes. Additional tissues were included to represent all other major tissue types beyond root tips, enabling capture of complete transcripts from *C. harringtonia*.

PacBio SMRT library preparation and sequencing were performed by QB3 Genomics, UC Berkeley, Berkeley, CA (RRID: <u>SCR_022170</u>) on the PacBio Sequel II System using the Sequel II 3.1 binding kit with a 30-hour movie time. RNA concentration was adjusted to <5 ng/μL using the Qubit HS RNA Kit, and RNA quality was assessed via capillary electrophoresis on an Agilent Bioanalyzer 2100 using the Eukaryotic Total RNA Pico Assay, as per the manufacturer instructions, with a 2 min denaturation at 70 °C followed by a 2 min cold shock on ice in a freezer. PacBio SMRT sequencing library was prepared as described in the Iso-Seq™ Template Preparation for Sequel® Systems manual using the NEBNext® Single Cell/Low Input cDNA Synthesis & Amplification Module and the Pacific Biosciences Iso-Seq Express Oligo Kit with the barcoded primers provided in the manufacturer’s protocol for first-strand cDNA synthesis (standard purification) and PCR amplification. Barcoded cDNAs from six representative samples (containing 10 unique tissue samples as discussed above) were pooled in equimolar amounts, and a single library was prepared from this pooled cDNA using the SMRTbell Prep Kit 3.0 following the standard protocol. The raw data were then processed using the Iso-Seq 3 pipeline in SMRT® Link software (PacBio) with default settings, resulting in generation of 357,877 high quality transcript isoforms. The quality of the Iso-seq transcriptome was assessed by mapping short read Ilumina short reads to the long read transcriptome assembly using bowtie2^70^ and through BUSCO analysis for completeness^71^. Peptide sequences, with longest ORF identified for each transcript, were then predicted using Transdecoder^69^, resulting in 349,712 contigs. Redundant contigs were removed using CD-HIT-EST^72^ with a cut-off at 99% identity, yielding a final PacBio Iso-Seq transcriptome of 29,163 unique transcripts.

For the old *in-house* PacBio Iso-Seq transcriptome—derived from tissues (young and old needles, old stem, and roots) harvested in August 2018 that lack CET biosynthetic genes and used as a comparison to the newly prepared *in-house* transcriptome from tissues harvested in May 2023 (see **Supplementary Fig. 8**)—library preparation and sequencing were performed by the Joint Genome Institute (JGI). The 2018 dataset was prepared and processed in the same manner as the May 2023 Iso-Seq transcriptome, except for this difference in sequencing facility and usage of CD-HIT-EST cut-off at 98% identity. The final processed transcriptome consisted of 48,657 unique transcripts.

### Analysis of RNA-seq data

Both the final processed PacBio Iso-seq and *de novo* Illumina-generated transcriptomes (from tissue samples in May 2023) were annotated with best-hit Pfam terms, followed by all subsequent (lower-ranked) Pfam hits^73^. Annotations also included the best Basic Local Alignment Search Tool (BLAST) hit from *Arabidopsis thaliana* proteomes^74^ (via UniProt), *Taxus chinensis* proteomes^75^, *Cephalotaxus hainanensis* proteomes^29^, and the *in-house* old Iso-seq transcriptome of *Cephalotaxus harringtonia* (harvested in August 2018), using BLASTP. Quantification of processed Illumina reads against both the PacBio and Illumina-generated transcriptomes was performed using Kallisto^76^ with default parameters.

For all downstream expression analysis, log_2_-transformed, TMM-normalized CPM expression values were used. Ultimately, the PacBio Iso-seq-generated transcriptome (from tissue samples in May 2023 paired feeding experiment) was used for all analyses and candidate gene selection to avoid technical limitations of *de novo* Illumina transcriptome assembly (i.e. misassembled or fragmented contigs)^39,40^. For example, *Ch*OMT-1, an essential CET biosynthetic gene we discovered, was detected in the PacBio Iso-seq transcriptome but absent from the *de novo* Illumina assembly.

### Cloning of candidate biosynthetic genes

cDNA was prepared from extracted RNA of *C. harringtonia* tissues harvested in May 2023 for paired metabolomics and transcriptomics using the SuperScript^TM^ IV First-Strand Synthesis System Kit. Because the candidate biosynthetic genes were highly expressed in root tips tissues, they were cloned mostly from cDNA generated from root tips of *C. harringtonia*. Coding sequences were PCR-amplified using Q5® High-Fidelity DNA Polymerase (New England Biolabs) following the manufacturer’s protocol. Oligonucleotide primers were designed to anneal to the 5′ and 3′ ends of the coding sequence for each target gene, incorporating flanking sequences homologous to the intended Gibson assembly sites of the appropriate expression plasmid. Following PCR amplification, the PCR products were assessed using gel electrophoresis on a 1% (w/v) agarose gel. PCR products were either purified using the DNA Clean & Concentrator-5 kit (Zymo Research) prior to transient expression construct assembly or, when necessary, were excised from the gel and purified using the Zymoclean Gel DNA Recovery Kit (Zymo Research).

For transient expression construct assembly, pEAQ-HT plasmid was digested with *AgeI and XhoI* restriction enzymes, and PCR amplicons were inserted into the digested pEAQ-HT vector using NEBuilder® HiFi DNA Assembly Mix (New England Biolabs), as per the manufacturer instructions. These Gibson assembly reactions were directly transformed into NEB 10β *Escherichia coli* cells (New England Biolabs) and grown on Luria–Bertani (LB) agar plates containing kanamycin (50 µg/mL) overnight at 37 °C. Positive colonies, with successful insertion of the target gene, were selected by screening with Colony PCR and were used to inoculate 4 mL LB liquid cultures with kanamycin (50 µg/mL), which were grown overnight on a rotary drum at 37 °C. Plasmid DNA was isolated using the ZR Plasmid Miniprep Kit (Zymo Research), and the sequences of the inserts were verified by either whole-plasmid sequencing (Plasmidsaurus) or Sanger sequencing (ELIM Biopharm Inc.).

pEAQ-HT expression constructs were transformed into *Agrobacterium tumefaciens* (GV3101:pMP90) via the freeze-thaw method. The transformed cultures were grown on LB agar plates containing kanamycin (50 µg/mL) and gentamycin (30 µg/mL) at 30 °C for two days to select positive transformants, which were confirmed by colony PCR. The verified positive colonies were then used to inoculate 2 mL LB liquid cultures (50 µg/mL kanamycin and 30 µg/mL gentamycin), which were grown for two days at 30 °C on a rotary drum and subsequently used to prepare 25% glycerol stocks that were snap-frozen in liquid nitrogen and stored at -80 °C for long-term use.

### Agrobacterium-mediated transient expression of Cephalotaxus genes in N. benthamiana

Transient expression in *Nicotiana benthamiana* was conducted much as previously described^30,47^. Glycerol stocks of *Agrobacterium tumefaciens* strains containing the pEAQ-HT expression construct of interest were streaked onto LB agar plates (kanamycin 50 µg/mL and gentamicin 30 µg/mL), which were then incubated at 30 °C for two days until a dense bacterial lawn developed. The bacterial cells were scraped off with a sterile pipette tip and resuspended in 1 mL of *Agrobacterium* induction buffer (10 mM MES, pH 5.6,10 mM MgCl_2_, 150 µM acetosyringone; Acros Organics), which were then incubated at room temperature for 1-3 hours. The optical density at 600 nm (OD_600_) was measured for each strain resuspension to determine relative cell density. Individual strain was diluted in induction buffer to an OD_600_ of 0.2-0.3 for infiltration. For transient co-expression, multiple strains of interest were combined in equal concentrations, at a final OD_600_ of 0.2-0.3 for each strain. These individual strains or strain mixtures were infiltrated into the abaxial side of 4-5 week-old *N. benthamiana* leaves using a 1 mL needleless syringe. For each transient expression reaction, three leaves—each from a different plant—were used as biological replicates to minimize any batch effects or biological variations among plants. After 3-4 days of infiltration, 150-250 µL of substrate (25-50 µM of compound **2** or **3** in water) was infiltrated into the abaxial side of previously *Agrobacterium*-infiltrated leaves using a needleless syringe. The area infiltrated with substrate was marked, excised one day after the substrate infiltration, and collected into pre-weighted 2 mL Safe-lock tubes (Eppendorf), which were flash frozen in liquid nitrogen and lyophilized to dryness for subsequent metabolite analysis.

### Preparation of crude protein lysates from N. benthamiana and their in vitro reactions for assessment of Pictet-Spengler condensation for phenethylisoquinoline scaffold formation

To evaluate the functionality of candidate enzymes for phenethylisoquinoline scaffold formation in a more controlled environment, cell-free protein lysates were prepared from *N. benthamiana* leaves transiently expressing genes of interest, much as previously described^77^. Briefly, leaves were harvested 4-5 days after infiltration with the *Agrobacterium* strain of interest (1-2 leaves for per sample) and snap frozen in liquid nitrogen. Enzymes were co-expressed for batch testing. The tissue was homogenized using a mortar and pestle with liquid nitrogen. Then, an ice-cold 2 mL Eppendorf tube was filled with the powdered tissue (∼1.5 mL), which was resuspended in 1.5 mL of ice-cold Tris buffer (100 mM Tris-HCl pH 7.4, 10% glycerol, 1 mM phenylmethylsulfonyl fluoride and 10 mM *β*-mercaptoethanol). The mixture was incubated at 4 °C for 30 min with gentle rocking to facilitate protein extraction. The mixture was then centrifuged at 12,000*g* and 4 °C for 10 min to pellet plant cell debris. The supernatant was removed, and this crude protein lysate was used as the bulk buffer for *in vitro* reactions.

Each reaction was carried out in a total reaction volume of 50 µL, containing 1 µL of 2.5 mM dopamine (final concentration, 50 µM), 1 µL of 2.5 mM requisite aldehyde (final concentration, 50 µM)—3-(4-hydroxyphenyl)propanal or 3-(4-hydroxy-3-methoxyphenyl)propanal to form **1** or **2** respectively, and 48 µL of the crude protein lysate. The reactions were incubated at 30 °C for 3 h, after which the reactions (50 µL) were quenched by an equal volume (1:1) of acetonitrile with 0.1% formic acid (50 µL) and diluted with 150 µL of methanol to a final volume of 250 µL (a final 5-fold dilution). Samples were then filtered through 0.45 µM hydrophilic PTFE filter plates (MilliporeSigma), and the filtrates were transferred into LC-MS vials for analysis. The remaining unused crude protein lysates were aliquoted (160 µL), snap frozen in liquid nitrogen, and stored at -80 °C.

### Heterologous expression of ChCYP805A11 in yeast

Expression of *Ch*CYP805A11 in *Saccharomyces cerevisiae* (yeast) was performed as much previously described^30,78^. The coding sequence of *Ch*CYP805A11 was PCR amplified from its pEAQ-HT construct and annealed into BamHI/EcoRI digested pYeDP60 plasmid (Carb^R^ for *E. coli* selection; *ADE2* for yeast selection) such that the resulting construct contained a C-terminal 6xHis tag through Gibson assembly, as discussed above in cloning of candidate biosynthetic genes. Similarly, the assembled plasmid construct was transformed into NEB 10β *E. coli cells, which were plated for selection on LB agar (100* µg/mL carbenicillin) and grown overnight at 37 °C. Positive colonies, verified through colony PCR, were grown on a rotary drum overnight in liquid LB media containing carbenicillin (*100* µg/mL) at 37 °C. The plasmid DNA was isolated using the Zymo Miniprep Kit and sequenced through whole-plasmid sequencing (Plasmidsaurus) to confirm the sequence of the inserted gene.

The assembled plasmid construct was transformed into chemically competent *S. cerevisiae* WAT11 (*ade*2), which contains a chromosomal copy of the *A. thaliana* cytochrome P450 reductase 1 gene (*ATR1*)^79^, using the Frozen-EZ Yeast Transformation II Kit (Zymo Research). Positive transformants were selected on synthetic drop-out media plates lacking adenine (6.7g/L yeast nitrogen base without amino acids, 20 g/L glucose, 2 g/L drop-out mix minus adenine [−Ade], 20 g/L agar)^80,81^ through growth for 2 days at 30 °C, which were confirmed through colony PCR. A single, positive colony was then used to inoculate a 4 mL of liquid cultures of synthetic drop-out medium, grown for 2 days at 30 °C with shaking at 250 rpm. The liquid cultures were then used to prepare 25% glycerol stocks and stored at -80 °C for long-term usage.

Freshly streaked colonies from *S. cerevisiae* WAT11 strain of interest (harboring pYeDP60 plasmid of *Ch*CYP805A11) were inoculated in 4 mL of liquid cultures of synthetic drop-out media (-Ade) and were grown at 30 °C for 2 days with shaking at 250 rpm. Then, 2 mL of the starter culture was used to inoculate 500 mL of the synthetic drop-out media (-Ade) and grown at 28 °C with 250 rpm shaking until reaching a cell density of approximately 5 x 10^7^ cells/mL, estimated through OD_600_ measurements. At this density, gene expression was induced by adding 50 mL of a sterile galactose solution (200 g/L) to a final concentration of 10% (v/v). The culture was grown for another 18 h at 28 °C and 250 rpm to reach a cell density of 5 x 10^8^ cell/mL, which was immediately used for microsomal protein isolation. Moreover, other than *Ch*CYP805A11 with C-terminal 6xHis tag, a negative control of empty vector and *Ch*CYP805A11 without His-tag (a negative control for immunoblotting) were prepared as well.

### Isolation of ChCYP805A11-enriched microsomes from yeast

Yeast microsomal proteins were isolated from yeast following a previously described protocol^78^. The galactose-induced yeast cultures were centrifuged at 5,000*g* for 5 min at 4° C to pellet cells. The cells were then resuspended in 1 mL of ice-cold TEK buffer (50 mM Tris-HCl, 1 mM EDTA, 100 mM KCl, pH 7.4) per 0.5 g of wet cell pellet mass and incubated at room temperature for 5 min The cells were centrifuged again as above and resuspended in 5 mL ice-cold TES B buffer (50 mM Tris-HCl, 1 mM EDTA, 600 mM sorbitol, pH 7.4). All subsequent steps were performed at 4 °C and/or on ice.

Approximately 5 mL (a volume equal to that of the cell resuspensions) of glass beads (0.5 mm diameter) was added to the cells, which were lysed by vigorous shaking for 10 min (30 s of shaking followed by 30 s on ice to avoid heating). Next, 10 mL of ice-cold TES B buffer was added to the lysate, which was then removed from the beads and saved. To extract any residual microsomal proteins on the beads, the beads were washed twice with 10 mL TES B buffer, and all washes were combined with the initial lysate (∼30 mL total). This pooled lysate was centrifuged at 23,000*g* for 10 min at 4 °C to pellet and remove large cell debris, and the resulting supernatant (∼30 mL) was diluted two-fold with ice-cold TES B buffer (to a final volume of ∼60 mL). Microsomes were precipitated by adding NaCl to a final concentration of 150 mM and polyethylene glycol (PEG)-4000 to a final concentration of 0.1 g/mL, followed by incubation on ice for about 1 h with periodic mixing to dissolve the solutes. Microsomal proteins were then pelleted by centrifugation at 10,000*g* for 10 min at 4 °C and were resuspended in 1 mL of ice-cold TEG storage buffer (50 mM Tris-HCl, 1 mM EDTA, 20% (v/v) glycerol, pH 7.4). The protein concentration of the microsomal fraction was measured using Bradford Assay, and 50-100 µL aliquots were snap-frozen in liquid nitrogen and stored at -80 °C until usage.

### Immunoblotting (Western blot)

Purified yeast microsomes expressing *Ch*CYP805A11 with a C-terminal 6-His tag, *Ch*CYP805A11 without a His-tag, or empty vector were each loaded into individual wells of a precast polyacrylamide gel (4–15% Mini-PROTEAN^®^ TGX™ Precast Protein Gels, 12-well, 20 µl; Bio-Rad 4561085). Proteins were separated for a total of 55 minutes (50 V for 10 min, followed by 150 V for 45 min) and transferred onto a polyvinylidene difluoride (PVDF) membrane using a Bio-Rad Trans-Blot Turbo Transfer System (Bio-Rad, 1704150). Membranes were incubated with horseradish peroxidase (HRP)-conjugated anti-His tag antibody (1:1,000 dilution; Biolegend) for 2 h at room temperature with gentle rocking. Blots were then washed and imaged using an iBright FL1500 Imaging System (Invitrogen, a44241). The microsomes were also analyzed for protein content by sodium dodecyl sulfate polyacrylamide gel (SDS-PAGE) with Coomassie Blue staining (see **Extended Data Fig. 3b** for both the immunoblot and the protein gel).

### Heterologous production and purification of ChNmrA-1

For heterologous production and purification of *Ch*NmrA-1, the gene was expressed in *E. coli*. The coding sequence of *Ch*NmrA-1 was first PCR amplified from its pEAQ-HT construct and inserted into NdeI/XhoI digested pET-24b (Km^R^) plasmid such that the resulting construct contained a C-terminal 6xHis tag through Gibson assembly, as discussed above in cloning of candidate biosynthetic genes. Similarly, the assembled plasmid construct was transformed into NEB 10β *E. coli cells, which were plated for selection on LB agar (50* µg/mL kanamycin) and grown overnight at 37 °C. Positive colonies, verified through colony PCR, were grown on a rotary drum overnight in liquid LB media containing kanamycin (*100* µg/mL) at 37 °C. The plasmid DNA was isolated using the Zymo Miniprep Kit and sequenced through whole-plasmid sequencing (Plasmidsaurus) to confirm the sequence of the inserted gene. The verified plasmid DNA was then transformed into *E. coli* BL21 (DE3) cells (NEB) as an expression host for heterologous protein production. A single positive colony confirmed via colony PCR was then inoculated in 3 mL of LB media (*50* µg/mL kanamycin) and grown overnight on a rotary drum at 37 °C, which was used to prepare 25% glycerol stocks that were snap-frozen in liquid nitrogen and stored at -80 °C for future use.

Freshly streaked colonies from *E. coli* BL21 strains of interest (harboring pET-24b of *Ch*NmrA-1) were inoculated in 2 mL of liquid LB cultures (*50* µg/mL kanamycin) and grown overnight on a rotary drum at 37 °C. A 500 µL of the starter culture was inoculated in 200 mL of liquid LB culture containing kanamycin (*50* µg/mL), which was grown on a culture shaker at 37 °C with 250 rpm until OD_600_ of ∼0.6 was reached. Once reached, the temperature was reduced to 18 °C and shaken for 0.5-1 h, followed by 0.5 mM IPTG induction. The cultures were shaken at 250 rpm for 18-20 h at 18°C and then centrifuged at 5,000 rpm for 10 min at 4 °C to pellet the cells. All subsequent steps were performed on ice and/or at 4 °C. The cell pellets (∼1-2 g) were resuspended in 5-8 mL of ice-cold binding buffer (50 mM potassium phosphate, pH 7.8, 100 mM NaCl, 10 mM imidazole) and lysed using EmulsiFlex^®^-B15 High-Pressure Homogenizer with pressure at 15,000 psi; samples were homogenized three times to ensure full lysis of the cells. Cell lysates were centrifuged at 10,000*g* for 30 min at 4 °C to pellet cell debris.

For protein purification, the clarified lysate was added to Ni-NTA resin (Ni-NTA Agarose, Invitrogen; R901-15) and incubated at 4 °C for 1 hour with gentle rocking. Bound proteins were eluted sequentially using potassium phosphate buffer (50 mM potassium phosphate, 100 mM NaCl, pH 7.8) with increasing concentrations of imidazole. Each eluted fraction was analyzed for protein content by SDS-PAGE with Coomassie Blue staining (see **Extended Data Fig. 3c** for the protein gel). Fractions containing pure enzyme were combined and concentrated using Amicon® Ultra-15 Centrifugal Filter Units (10 kDa molecular weight cut-off) by centrifugation at 4,000*g* for 20 min at 4 °C. The unfiltered protein-containing solution (concentrated to <1 mL) was diluted with 10-13 mL of ice-cold potassium phosphate buffer and reconcentrated as above; this dilution-concentration cycle was repeated 6-8 times to dilute out imidazole. The final concentrated protein solution was diluted to approximately 1 mL with ice-cold phosphate buffer, which had glycerol added to a final concentration of 10% (v/v). The purified protein concentration as measured using Bradford Assay, and 30 µL aliquots were snap-frozen in liquid nitrogen and stored at -80 °C for future use.

### In vitro reactions with purified ChNmrA-1 and ChCYP805A11-enriched microsomes

*In vitro* reactions with purified *Ch*NmrA-1 and *Ch*CYP805A11-enriched microsomes were carried out in 200 µL of potassium phosphate buffer (50 mM potassium phosphate, 100 mM NaCl, pH 7.8) with a final concentration of 50 µM of initial substrate (compound **3**) and 1 mM NADPH. Purified *Ch*NmrA-1 (6xHis-tagged) and yeast microsomal proteins containing *Ch*CYP805A11 (6xHis-tagged) were each added to a final concentration of 0.4 µg/µL in a total reaction volume of 200 µL. As negative controls, microsomal fractions containing empty vector (EV) and denatured (boiled) *Ch*NmrA-1 were used. Dependence of the reaction on both the substrate (**3**) and cofactor (NADPH) was also demonstrated through dropping out each component. The reactions were incubated at 30 °C for several time points (0.5, 2, and 4 h). The reaction aliquots (20 µL) were then quenched by an equal volume (1:1) of methanol (20 µL), followed by addition of an equal volume (1:1) of ice-cold acetonitrile with 0.1% formic acid (40 µL). Samples were further diluted with 120 µL of ice-cold methanol to a final volume of 200 µL (2:3, v/v). In parallel, to probe the structure of compound **4’**, proposed to be an unstable imine intermediate, a 20 µL aliquot taken from the same 200 µL *in vitro* reaction was not immediately quenched with methanol, but instead treated with an equal volume (1:1) of 100 mM NaBD_3_CN in acetate buffer (200 mM sodium acetate, pH 5.5) and incubated for 15 min at room temperature to chemically (non-enzymatically) reduce the imine of **4’** to the corresponding amine, yielding monodeuterated **5** (**5**-D)^60^. The resulting reactions (40 µL) were quenched with an equal volume (1:1) of ice-cold acetonitrile with 1% formic acid (40 µL), followed by dilution with 120 µL of ice-cold methanol to a final volume of 200 µL (2:3, v/v). All quenched samples were vortexed and centrifuged briefly. The resulting supernatants were filtered through 0.45 µM hydrophilic PTFE filter plates (MilliporeSigma), and the filtrates were transferred into LC-MS vials for analysis.

### Metabolite extraction from plant material

The overnight-lyophilized tissues (i.e. various tissues of *C. harringtonia* from feeding experiments and leaves from *N. benthamiana* transient expression experiments) were homogenized on a ball mill homogenizer (Retsch MM 400) using a 5 mm diameter steel bead for each sample tube, with shaking at 25 Hz for 2 min Steel beads were then removed, and each tissue sample was extracted with 40 µL and 20 µL of 80% methanol in water per milligram of dry plant tissue weight for *C. harringtonia* tissues and *N. benthamiana* leaves, respectively. The samples were briefly vortexed and sonicated, then incubated for 30 min at room temperature for metabolite extraction. For *C. harringtonia* tissues, additional 20 min of incubation at 4 °C was performed. The extracted samples were centrifuged at 16,000*g* for 1 min to pellet plant cell debris, and the supernatant was filtered through hydrophilic PTFE filter plates with 0.45 µM pore size (MilliporeSigma). The filtrates were transferred into LC-MS vials for analysis.

For quantification of natural abundance of cephalotaxinone, CET, and HHT, a 1000-fold and 200-fold dilution was performed for tissues harvested in May 2022 and May 2023 respectively to avoid saturation of the parent isotope peaks in LC-MS. For tissues harvested from D_2_O labeling assays (conducted in late April 2022), additional 100-fold dilution was also performed to avoid saturation of the parent isotope peaks in LC-MS.

### LC-MS analysis of metabolites

All metabolite samples were analyzed on the LC-MS instrument of an Agilent 1290 Infinity II UHPLC coupled to an Agilent 6546 Q-TOF mass spectrometer (6546 LC-MS). For separation of the metabolites, a reversed-phase liquid chromatography using a ZORBAX RRHD Eclipse Plus 95Å C18 column (Agilent, 2.1 x 50 mm, 1.8 μm) was performed with a mobile phase consisting of two buffers: water with 0.1% formic acid (buffer A) and acetonitrile with 0.1% formic acid (buffer B). An injection volume of 1 µL and a flow rate of 0.6 mL/min were used with the following 29 min gradient method: 0-0.5 min, 3% B; 0.5-7.5 min, 3-10% B; 7.5-15.5 min, 10-25% B; 15.5-20.5 min, 25-50% B; 20.5-21.5 min, 50-95% B; 21.5-23.0 min, 95% B; 23.0-24.5 min, 95-30% B; 24.5-26.5 min, 30-3% B; 26.5-29.0 min, 3% B. MS data were collected using a dual-inlet Agilent Jet Steam electrospray ionization (Dual AJS ESI) in positive ion mode with a mass range of 50-1,700 *m/z* and a rate of 1.00 spectrum per second (1000 ms per spectrum). The ionization source was set as follows: 325 °C gas temperature, 10 L/min drying gas, 35 psi nebulizer, 350 °C sheath gas temperature, 12 L/min sheath gas flow, 135 V fragmentor, 45 V skimmer, 750 V Oct 1 RF Vpp, 4000 V VCap, and 0 V nozzle voltage. For tandem mass spectrometry (MS/MS) analysis, the mass ions pertaining to individual metabolites were fragmented with collision energies of 15, 20, 30, or 35 V. All data were collected in centroid mode, but when needed, data were also stored in profile mode. Metabolite samples from *in vitro* reactions were analyzed using a shortened gradient method in which the first 7.5 min were identical to the standard long gradient method; all relevant metabolites in *in vitro* reactions (including compounds **4** and **5**) eluted within this window, with compound **3** eluting shortly thereafter (∼7.8 min)—resulting in the same elution behavior as observed with the long gradient method. The remainder of the shortened gradient was as follows: 7.5-9 min, 10-50% B; 9-10 min, 50-95% B; 10-11.5 min, 95% B; 11.5-12.5 min, 95-3% B; 12.5-15 min, 3% B.

### Metabolomics and MS data analyses

LC-MS data were analyzed using Agilent MassHunter Qualitative Analysis 10.0 and Agilent MassHunter Quantitative Analysis 11.0 software. The ion abundances for a given mass signature were calculated using the automated integration (‘Agile’ method with default settings) of extracted ion chromatograms (EICs) with a 20 ppm mass tolerance. Peaks integrated by the Quantitative Analysis software were manually inspected to ensure correct peak selection and integration.

Untargeted metabolomic analyses were conducted using XCMS Online^82^ with default parameters of UPLC / UHD Q-TOF (155). The output of this analysis contains a tabulated list of identified mass signatures at specific retention times, with associated *m/z* values, peak intensity fold change, *P* value (two-tailed unequal variance Welch’s *t*-test), retention times, and extracted peak intensities.

For XCMS analysis comparing dopamine-D_4_-fed root tips with water-fed controls of *C. harringtonia* from isotope labeled precursor feeding studies, we used the following filters to look for differential mass signatures: *P* value < 0.2, fold change > 3.5, and *m/z* range of in between 200 and 600.

For XCMS analysis to determine masses found at differential levels between two samples from *N. benthamiana* transient expression experiments, we used the following filters, unless stated otherwise: *P* value < 0.1, fold change > 5, average peak intensity > 5x10^4^, retention time < 18 min, and 250 < *m/z* < 600.

For XCMS analysis to determine masses found at differential levels between two samples from *in vitro* experiments, we used the same filters as above for *N. benthamiana* transient expression experiments, with the exception of the retention time window, which was modified to accommodate the shortened gradient (< 10 min).

### Synthesis of different phenethylisoquinolines (1, 1-D_4_, 2, 2-D_4_, 3, 11, 12, 12-D_4_, and 13)

The synthesis of compound **1** (1-(4-hydroxyphenethyl)-1,2,3,4-tetrahydroisoquinoline-6,7-diol) was adapted from a previously described protocol^83^. Dopamine (0.42 mmol) and of 3-(4-hydroxyphenyl)propanal (0.50 mmol) were combined in 4 mL of a 1:1 (v/v) mixture of acetonitrile and potassium phosphate buffer (0.1 M, pH 6.0). The reaction mixture was stirred at 50 °C for 16 h under N_2_ with reflux. The mixture was then frozen in liquid-nitrogen and lyophilized to dryness.

The above procedure was applied to the synthesis of all compounds listed in Table 1 below, substituting the amine and aldehyde starting materials as specified.

**Table 1.** Synthesis of phenethylisoquinolines.

| Compound | Amine (0.42 mmol) | Aldehyde (0.50 mmol) |
| --- | --- | --- |
| **1** | dopamine | 3-(4-hydroxyphenyl)propanal |
| **1-D<sub>4</sub>** | dopamine-D<sub>4</sub> | 3-(4-hydroxyphenyl)propanal |
| **2** | dopamine | 3-(4-hydroxy-3-methoxyphenyl)propanal |
| **2-D<sub>4</sub>** | dopamine-D<sub>4</sub> | 3-(4-hydroxy-3-methoxyphenyl)propanal |
| **3** | 3-hydroxy-4-methoxyphenethylamine | 3-(4-hydroxy-3-methoxyphenyl)propanal |
| **11** | 3-hydroxy-4-methoxyphenethylamine | 3-(4-hydroxyphenyl)propanal |
| **12** | dopamine | 3-(3,4-dihydroxyphenyl)propanal |
| **12-D<sub>4</sub>** | dopamine-D<sub>4</sub> | 3-(3,4-dihydroxyphenyl)propanal |
| **13** | 3-methoxy-4-hydroxyphenethylamine | 3-(3,4-dihydroxyphenyl)propanal |

For compounds **1**, **1**-D_4_, **11**, **12**, **12**-D_4_, and **13**, the lyophilized crude product was resuspended in ∼1-2 mL of DMSO and directly loaded onto a Biotage® Sfär C18 D Duo 100 Å 30 µm 6 g column. Purification was performed an automated Biotage Selekt system using water with 0.1% formic acid and acetonitrile with 0.1% formic acid as mobile phases. Eluting fractions containing the compound of interest (identified by LC-MS) were pooled, frozen in liquid nitrogen, and lyophilized to dryness.

For compounds **2**, **2**-D_4_, and **3**, the lyophilized crude product was resuspended in ∼2-3 mL of 80% MeOH and purified by preparative HPLC using an Agilent 1260 Infinity preparative-scale HPLC system with an Agilent 1100 diode array detector and a Clipeus C18 5-µm, 10-mm x 250-mm semi-prep column (Higgins Analytical). Water with 0.1% formic acid (A) and acetonitrile with 0.1% formic acid (B) were used as the mobile phases.

For compounds **2** and **2**-D_4_, an injection volume of 500 µL and a flow rate of 5 mL/min were used with the following gradient method: 0-5 min, 3% B; 5-8 min, 3-10% B; 8-23 min, 10-20% B; 23-24 min, 20-25% B; 24-24.1 min, 25-97% B; 24.1-29.1 min, 97% B; 29.1-29.2 min, 97-3% B; 29.2-34 min, 3% B. Fractions were collected from 1-34 min, and those containing the compound of interest (verified by LC-MS) were combined, frozen, and lyophilized to dryness.

For compound **3,** an injection volume of 500 µL and a flow rate of 5 mL/min were used with the following gradient method: 0-5 min, 3% B; 5-8 min, 3-16% B; 8-18 min, 16-24.5% B; 18-19 min, 24.5-30% B; 19-19.1 min, 30-97% B; 19.1-24.1 min, 97% B; 24.1-24.2 min, 97-3% B; 24.2-29 min, 3% B. Fractions were collected from 1-29 min, and those containing the desired compound (verified by LC-MS) were pooled, frozen, and lyophilized to dryness.

All the purified products **1**, **1**-D_4_, **2**, **2**-D_4_, **3**, **11**, **12**, and **12**-D_4_ were analyzed by both LC-MS/MS and NMR. For compound **13**, its structure was verified by LC-MS/MS.

### Heterologous production and purification of compound 3-D_4_ using ChOMT-1

Compound **3**-D_4_ was biosynthesized from chemically synthesized **2**-D_4_ by heterologous expression of *Ch*OMT-1 in *E. coli* BL21 (DE3). Cloning and expression of *Ch*OMT-1 were performed exactly as described for *Ch*NmrA-1 (see Methods: *Heterologous production and purification of ChNmrA-1*), except that the induction culture volume was increased to 900 mL. After 18 h of 0.5 mM IPTG induction of *Ch*OMT-1 at 18 °C with shaking at 250 rpm, compound **2**-D_4_ was added to a final concentration of 50 µM for O-methylation reaction of **2**-D_4_ to **3**-D_4_. The biosynthetic reaction was run for 5 h with continuous shaking of 250 rpm at 18 °C. The bacterial cells were pelleted, resuspended in 900 mL of fresh LB media, and subjected to two additional cycles of the O-methylation reaction (total of three 900 mL reactions). The combined supernatants containing the biosynthesized **3**-D_4_ were extracted with ethyl acetate (1:1, v/v) four times. Organic phases were combined, dried using rotary evaporation, and resuspended in ∼3 mL of 80% MeOH. Purification was performed using the same preparative HPLC system, column, and mobile phases as described for compounds **2**, **2**-D_4_, and **3**, but with a modified gradient method.

For compound **3**-D_4_, an injection volume of 200 µL and a flow rate of 5 mL/min were used with the following gradient method: 0-5 min, 3% B; 5-8 min, 3-9.5% B; 8-16 min, 9.5% B; 16-21 min, 9.5-11% B; 21-36 min, 11% B; 36-41 min, 11-13% B; 41-44 min, 13-16%; 44-46 min, 16-20 min; 46-47 min, 20-30%; 47-47.1 min, 30-97%; 47.1-52.1 min, 97% B; 52.1-52.2 min, 97-3% B; 52.2-57 min, 3% B.

Fractions were collected from 1-57 min, and those containing the compound of interest (verified by LC-MS) were combined, frozen, and lyophilized to dryness. The purified product **3**-D_4_ was then analyzed by LC-MS/MS and NMR.

### Extraction and purification of a cephalotaxinone (10) standard

Old leaves and stem of *C. harringtonia* were harvested, frozen at -80 °C, and lyophilized to dryness. The dried tissues (∼10 g) were then ground to a fine powder or small pieces using mortar and pestle, and extracted with 700 mL of methanol for 3-4 days at room temperature with constant stirring. Extracts were filtered using vacuum filtration and dried by rotary evaporation. Because the compound of interest (CET intermediate) is alkaloid, an acid-base extraction was performed to remove water-soluble and non-alkaloid impurities prior to two rounds of chromatography. The residue was resuspended in 450 mL of water with HCl (pH 2-3). This fraction was partitioned with 400 mL of diethyl ether or *tert*-butyl methyl ether. The aqueous fraction was collected, and the organic phase was washed twice with 100 mL of water (pH 2-3) to maximize extraction of the desired compound, after which each of these aqueous fractions was combined with the initial 450 mL aqueous fraction. Then, the pH of the aqueous phase was increased to pH 8-9 using saturated sodium bicarbonate. This solution was partitioned with 600 mL of ethyl acetate. The organic layer, which supposedly contains the target compound, was collected, combined with two additional 100 mL ethyl acetate fractions used to wash the aqueous phase, and dried using rotary evaporation.

The dried residue was resuspended in ∼1-2 mL of methanol and loaded onto a 3.5 cm-diameter column packed with 18 g of P60 silica gel (SiliCycle). Hexane (HPLC grade; VWR) and ethyl acetate (ACS reagent grade; J.T. Baker) were used as mobile phases. The following gradient was applied: 100 mL each of 50%, 60%, 70%, 80%, 90%, and 100% ethyl acetate. Each step was collected in three fractions, yielding a total of 18 fractions. Fractions containing the compound of interest (identified by LC-MS) were combined and dried by rotary evaporation. The partially purified product was resuspended in ∼3-4 mL of 80% MeOH for second round of chromatography.

For the second round, purification was performed using the same preparative HPLC system, column, and mobile phases as described for compounds **2**, **2**-D_4_, and **3**, but with a modified gradient method. An injection volume of 450 µL and a flow rate of 5 mL/min were used with the following gradient method: 0-5 min, 3% B; 5-8 min, 3-7% B; 8-10 min, 7% B; 10-13 min, 7-10% B; 13-16 min, 10% B; 16-20 min, 10-15% B; 20-24 min, 15%; 24-30 min, 15-25 min; 30-37 min, 25-40%; 37-37.1 min, 40-97%; 37.1-42.1 min, 97% B; 42.1-42.2 min, 97-3% B; 42.2-47 min, 3% B. Fractions were collected from 1-47 min, and those containing the compound of interest (confirmed by LC-MS) were combined, frozen, and lyophilized to dryness.

### NMR analysis of purified compound

DMSO-d_6_ (ThermoFisher Scientific) was used as the solvent for all NMR samples of phenethylisoquinolines (**1**, **1**-D_4_, **2**, **2**-D_4_, **3**, **3**-D_4_, **11**, **12**, and **12**-D_4_). For NMR sample of cephalotaxinone (**10**), CDCl_3_ (Acros Organics) was used as the solvent. ^1^H, ^13^C, and 2D-NMR spectra were mostly acquired on a Bruker NEO 500-MHz spectrometer, and on a Varian Inova 600-MHz (for ROESY for **10**) at room temperature using VNMRJ 4.2. The data were visualized, analyzed, and processed on MestReNova v.15.1.0. Chemical shifts were reported in ppm downfield from Me_4_Si by using the residual solvent peak as an internal standard: 7.26 ppm for ^1^H and 77.16 ppm for ^13^C ppm chemical shift of CDCl_3_ and 2.50 ppm for ^1^H and 39.52 ppm for ^13^C ppm chemical shift of DMSO-d_6_.

The NMR spectra for compounds **1**, **1**-D_4_, **2**, **2**-D_4_, **3**, **3**-D_4_, **11**, **12**, **12**-D_4_, and cephalotaxinone (**10**) are shown in **Supplementary Figs. 13-63**, and their ^1^H/^13^C assignments are detailed in **Supplementary Tables 6-15**.

### Sequence analysis of NmrA-like proteins and construction of their phylogenetic tree

Sequence analyses of NmrA-like proteins (atypical SDRs) and related enzymes were performed in Geneious (v2022.2.1). To generate protein alignments of NmrA-like proteins (atypical SDRs), an assortment of protein sequences with PFAM (PF05368), containing the NAD(P)^+^/NAD(P)H-binding motif, was downloaded from UniProt (https://www.uniprot.org/), Phytozome (https://phytozome-next.jgi.doe.gov/), and NCBI database (https://www.ncbi.nlm.nih.gov/). NmrA-like proteins (PF05368) from high-quality plant genomes, top 2 best BLASTP hits of *Ch*NmrA-1 in the NCBI database, enzymes with verified reductase activity in plant biosynthesis such as the pinoresinol-lariciresinol reductases (PLRs) in lignan biosynthesis (PF05368) and classical SDRs (PF00106), and NmrA-like proteins (PF05368) with verified regulatory functions such as transcription repressors or redox sensor proteins from animals, fungi, and bacteria were selected. Another NmrA-like protein from *C. harringtonia* (YD-1-Rt_transcript/51169.p1) was also included for analysis, which was identified using best reciprocal BLASTP analysis: *Ch*NmrA-1 was used as a query against the selected high-quality plant genomes, and the top hits were then queried back against our *C. harringtonia* transcriptome. *Ch*NmrA-1 was used as the reference for amino acid numbering in alignments, as it represents the focal NmrA-like protein investigated in this study. Classical SDRs, including SDR1 from *A. thaliana* (UniProt ID: Q9M2E2)^84,85^, were included as canonical enzymatic references for comparison of the NAD(P)H-binding motif^55,86^. All sequences were downloaded from UniProt, except the protein from *T. lanceolata* (accession number, XP_077234198.1) and *Pt*SDR-2 from *P. tetrastichus* (GenBank accession number, OR538109) which were downloaded from NCBI database. The downloaded sequences were aligned using MAFFT^87^, and the phylogenetic tree was constructed using FastTree^88^ with a rate of categories of sites as 20 and an optimization of the Gamma20 likelihood.

### General software use and graph generation

Routine data compilation was performed in Microsoft Excel (version 16.62), including calculation of Pearson’s correlation coefficient used for co-expression analysis. General LC-MS data analysis was performed with Agilent MassHunter Qualitative Analysis 10.0. Chromatograms and mass spectra were plotted using IGOR Pro (version 8.04). Bar graphs, lines graphs, and scatterplots were plotted using GraphPad Prism 10, including mean and standard deviations denoted in the graphs.

Geneious Prime (v2022.2.1) was used for bioinformatic analyses of nucleic acid and protein sequences. This software was also used for several sequence alignments (MAFFT)^87^ and phylogenetic tree generation (FastTree)^88^. NMR data were processed and visualized on MestReNova v15.1.0. Molecular weight calculation, and chemical structural visualization were conducted using ChemDraw Professional v25.0.2.

### Data Availability

The raw RNA-seq data analyzed and used for discovery (*C. harringtonia* tissues harvested in May 2023) have been deposited in the National Center for Biotechnology Information (NCBI) Sequence Read Archive (SRA) database under the BioProject ID PRJNA1395191. The processed RNA-seq data, including both the processed PacBio Iso-seq transcriptome and a sheet of expression values (in log_2_-transformed values of each transcript calculated as TMM-normalized CPM) for all 29,163 transcripts, analyzed and used for discovery have been deposited at the NCBI Gene Expression Omnibus (GEO) (accession GSE319274). Gene sequences for enzymes characterized in this study have been deposited in the NCBI GenBank under the following accessions: *Ch*OMT-1 (PX794901), *Ch*CYP805A11 (PX794902), *Ch*NmrA-1 (PX794903), *Ch*CYP805A12 (PX794904), *Ch*CYP805A13 (PX794905), and *Ch*2OGD-1 (PX794906). The UniProt database (https://www.uniprot.org/), Phytozome database (https://phytozome-next.jgi.doe.gov/), and NCBI database (https://www.ncbi.nlm.nih.gov/) were used for identifying and obtaining NmrA-like protein sequences that were used in phylogenetic analyses. All other raw/source data are available upon request.

## ACKNOWLEDGEMENTS

We thank Stephen Lynch (Stanford University) for his assistance and helpful discussion related to NMR analysis. We also thank Prof. David Nelson (University of Tennessee) for providing cytochrome P450 nomenclature. We acknowledge Prof. George Lomonossoff for providing us with the pEAQ-HT plasmid. We are grateful to Warren Lau and Catherine Liou for early contributions to this project. The author is also thankful to Diego Wengier and Sarah Niehs for feedback on this manuscript. An initial *C. harringtonia* transcriptome (August 2018) (proposal: 10.46936/10.25585/60001097 led by J. Bohlmann) was performed by the US Department of Energy (DoE) Joint Genome Institute (https://ror.org/04xm1d337), a DoE Office of Science User Facility, and is supported by the Office of Science of the US DoE operated under contract number DE-AC02-05CH11231. The research is supported by an NIH R01 GM121527 to E.S.S.

## AUTHOR CONTRIBUTIONS

**Y.D.** and **E.S.S.** led the project and conceived of experimental procedures. **Y.D.** performed the experiments, generated the new *C. harringtonia* transcriptome (May 2023) used for discovery, and analyzed the data. **K.S.** collected the samples for the initial *C. harringtonia* transcriptome (August 2018) for seasonal comparison and helped develop the hypothesis for a whole plant coordination of HHT biosynthesis. **Y.D.** and **E.S.S.** interpreted the data, and **Y.D.** wrote the manuscript with edits from **E.S.S.**

