## Extended Data and Supplementary Information for "Discovery of cephalotaxinone enzymes reveals a whole plant model for homoharringtonine biosynthesis"

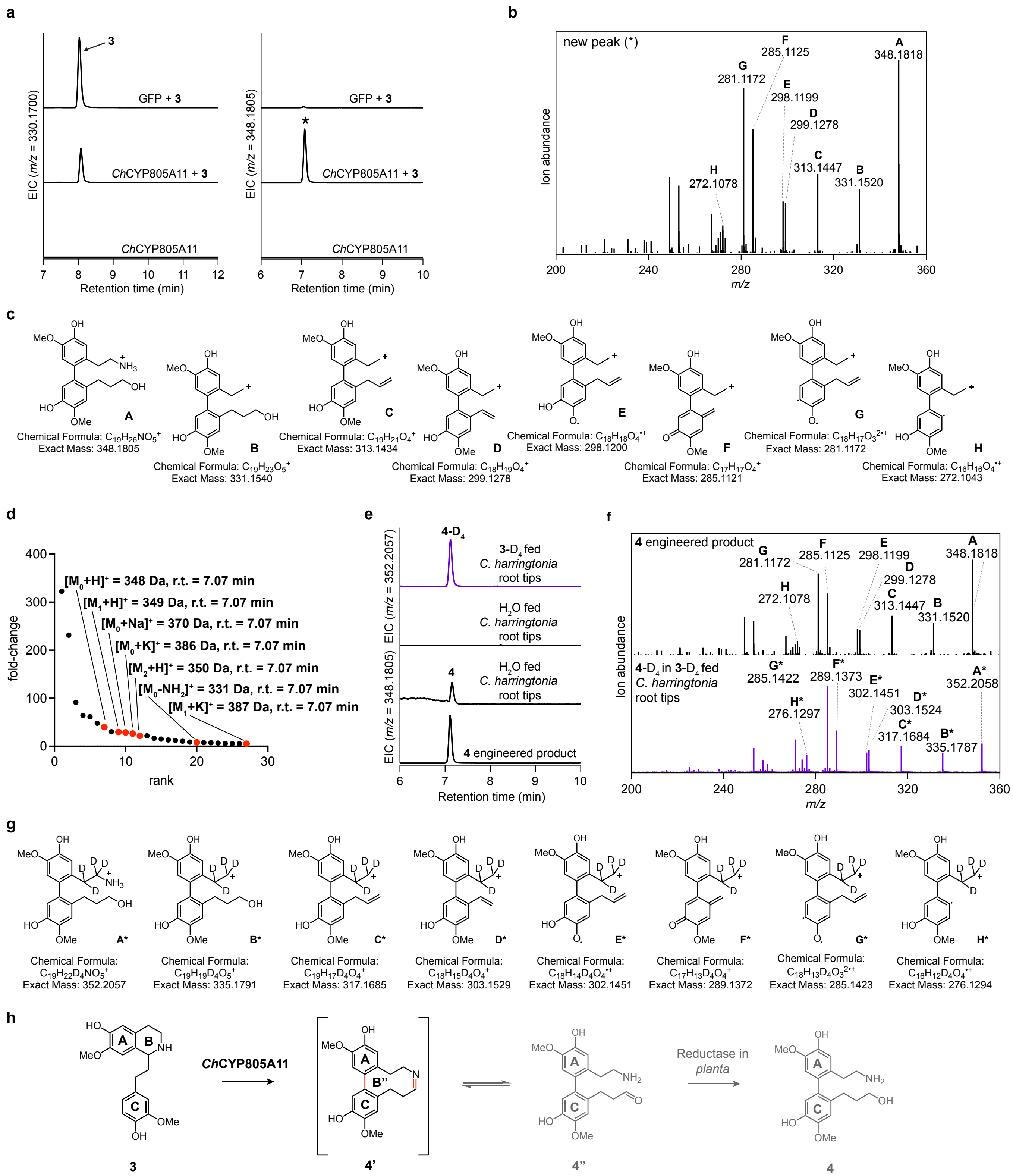


**Extended Data Figure 1. Characterization of *Ch*CYP805A11 and its resulting product (4). (a)** Expression of *Ch*CYP805A11 in *N. benthamiana* with co-infiltrated **3** leads to consumption of **3** ($[M + H]^+$ = *m/z* 330.1700, r.t. of 8.1 min) and production of a new compound (*) that corresponds to a hydroxylation and a reduction ($[M + H]^+$ = *m/z* 348.1805), as shown by the LC-MS chromatograms. The new compound is dependent on substrate (**3**)**.** This experiment was performed more than three times, with similar results observed each time. **(b)** MS/MS fragmentation spectrum of the generated *m/z* 348.1805 product (*) at a collision energy of 15 V, with key experimental fragment ions labeled. **(c)** Putative structures for ion fragments generated from MS/MS analysis of the new product (*m/z* 348.1805). The theoretical and experimental fragment ions support the proposed structure. **(d)** Untargeted metabolite analysis (XCMS) comparing transient expression of GFP (negative control) to that of *Ch*CYP805A11 with co-infiltrated substrate **3** (*n* = 3 independent replicates per condition). The unique *m/z* features ($P < 0.1$ and fold change > 5 between samples, see Methods) are shown in ranked order based on their increasing fold change in ion abundance between the two conditions. The mass isotopologues and adducts of the putative product (*m/z* 348.1805) are shown in red. The rest in black are below signal to noise threshold and cannot be identified in the raw LC-MS data. r.t., retention time. **(e)** The new *m/z* 348.1805 engineered product (**4**) corresponds to a metabolite found in the root tips of *C. harringtonia*, as shown by the LC-MS chromatograms. Deuterium-labeled analog of **4** (**4**-$D_4$) is observed in **3**-$D_4$ fed *C. harringtonia* root tips ($[M + H]^+$ = *m/z* 352.2057), shown in violet traces. **(f)** Comparison of the MS/MS spectrum of enzymatically produced **4** in *N. benthamiana* (black traces) with that of **4**-$D_4$ from **3**-$D_4$ fed *C. harringtonia* root tips (violet traces) at a collision energy of 15V. The two spectra have very similar fragmentation patterns with a major difference of four hydrogens in mass. **(g)** Putative structures for ion fragments generated from MS/MS analysis of **4**-$D_4$ ($[M + H]^+$ = *m/z* 352.2057). The theoretical and experimental fragment ions support the proposed structure. **(h)** Proposed reaction catalyzed by *Ch*CYP805A11, as supported by MS/MS fragmentation pattern of the new product (**4**) and its comparison to that of the corresponding four-deuterated form (**4**-$D_4$). Compounds **4'** and **4''** are not detectable due to their structural instability, leading to detection of **4** instead.

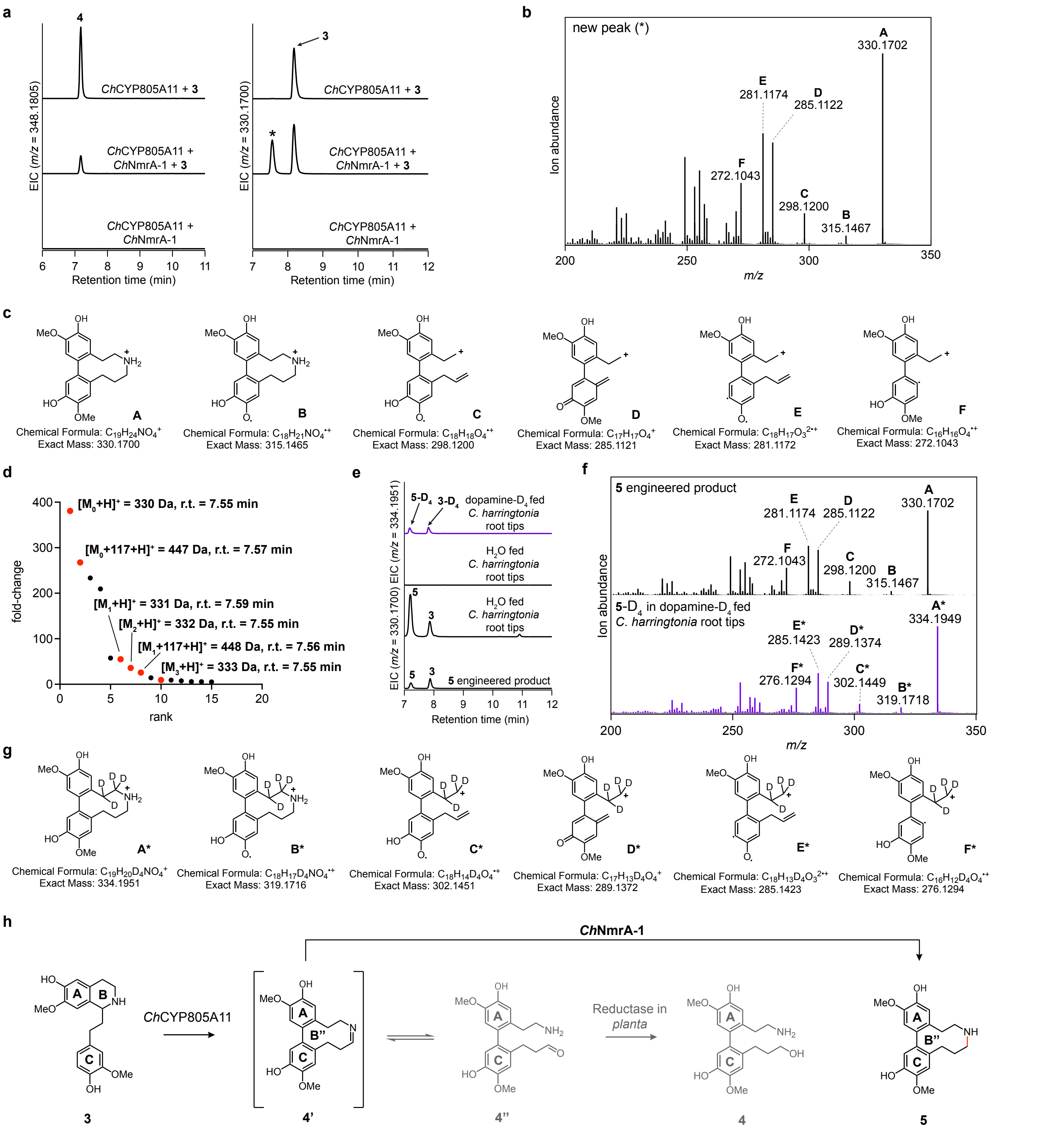


**Extended Data Figure 2. Characterization of *Ch*NmrA-1 and its resulting product (5). (a)** Addition of *Ch*NmrA-1 to the *N. benthamiana* transient expression system with co-infiltrated **3** leads to a decrease of **4** ($[M + H]^+$ = *m/z* 348.1805) and production of a new compound (*) that corresponds to a dehydroxylation and a loss of two hydrogens ($[M + H]^+$ = *m/z* 330.1700, r. t. of 7.6 min), as shown by the LC-MS chromatograms. The new compound is dependent on the initial substrate (**3**). This experiment was performed more than three times, with similar results observed each time. **(b)** MS/MS fragmentation spectrum of the generated *m/z* 330.1700 product (*) at a collision energy of 30 V, with key experimental fragment ions labeled. **(c)** Putative structures for ion fragments generated from MS/MS analysis of the new product (*m/z* 330.1700). The theoretical and experimental fragment ions support the proposed structure. **(d)** Untargeted metabolite analysis (XCMS) comparing the presence and absence of *Ch*NmrA-1 in the *N. benthamiana* transient co-expression system with co-infiltrated substrate **3** ($n$ = 3 independent replicates per condition). The unique *m/z* features ($P < 0.1$ and fold change > 5 between samples, see Methods) are shown in ranked order based on their increasing fold change in ion abundance between the two conditions. The mass isotopologues and adducts of the putative product (*m/z* 330.1700, r.t.

of 7.6 min) are shown in red. The rest in black are mostly below signal to noise threshold and cannot be identified in the raw LC-MS, independent of the initial substrate **3**, or non-native to *C. harringtonia*. r.t., retention time. **(e)** The new *m/z* 330.1700 engineered product (**5**) corresponds to a metabolite found in the root tips of *C. harringtonia*, as shown by the LC-MS chromatograms. Deuterium-labeled analog of **5** (**5**-$D_4$) is observed in dopamine-$D_4$ fed *C. harringtonia* root tips ($[M + H]^+$ = *m/z* 334.1951), shown in violet traces. Note: minimal retention time shifts (~0.2-0.3 min) are often observed in LC-MS runs conducted at different times. **(f)** Comparison of the MS/MS spectrum of enzymatically produced **5** in *N. benthamiana* (black traces) with that of **5**-$D_4$ from dopamine-$D_4$ fed *C. harringtonia* root tips (violet traces) at a collision energy of 30V. The two spectra have very similar fragmentation patterns with a major difference of four hydrogens in mass. **(g)** Putative structures for ion fragments generated from MS/MS analysis of **5**-$D_4$ ($[M + H]^+$ = *m/z* 334.1951). The theoretical and experimental fragment ions support the proposed structure. **(h)** Proposed reaction catalyzed by *Ch*NmrA-1, as supported by MS/MS fragmentation pattern of the new product (**5**) and its comparison to that of the corresponding four-deuterated form (**5**-$D_4$). Compounds **4'** and **4''** are not detectable due to their structural instability.

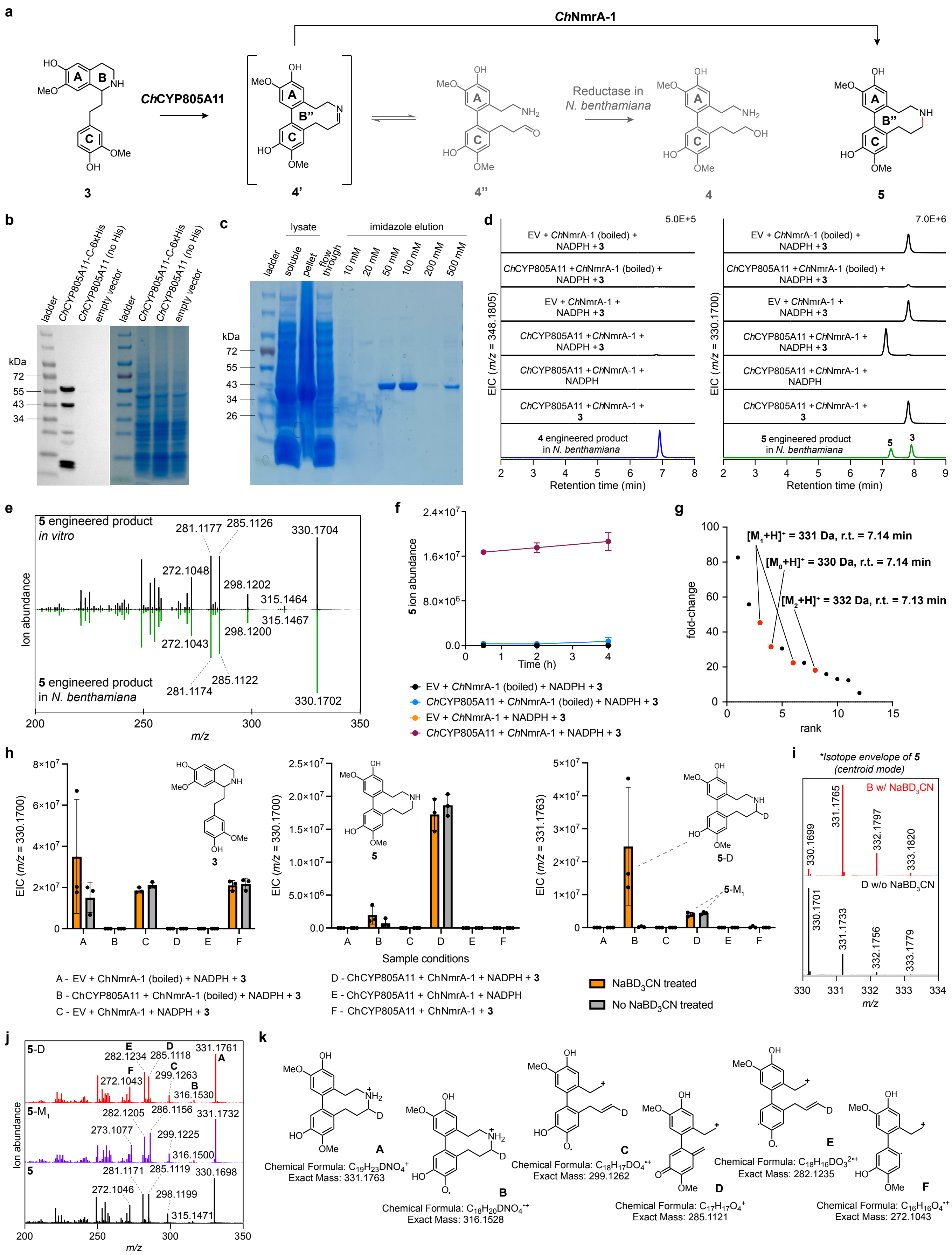


**Extended Data Figure 3. *In vitro* confirmation of *Ch*CYP805A11 and *Ch*NmrA-1 catalytic activity.** *Ch*CYP805A11 (with a C-terminal 6xHis tag) enriched microsomes were produced in and isolated from yeast. *Ch*NmrA-1 was expressed in *E. coli* BL21 cells with C-terminal 6xHis tag, which were used to purify the enzyme through Ni-NTA affinity chromatography. These enzymes were used to confirm their catalytic activities *in vitro*. **(a)** Proposed reactions catalyzed by *Ch*CYP805A11 and *Ch*NmrA-1 for biosynthesis of **5** from **3**, confirmed through *in vitro* characterization. **(b)** Immunoblot of *Ch*CYP805A11 with C-terminal 6xHis tag (MW = 60.20 kDa) enriched microsomes produced in and isolated from yeast (left). Yeast microsomes enriched with *Ch*CYP805A11 without a His-tag and yeast microsomal fractions containing empty vector (EV) are included as a negative reference. SDS-PAGE gel of the same fractions of yeast microsomes is included as another negative reference (right). **(c)** SDS-PAGE gel of fractions from purification of *Ch*NmrA-1 (MW = 36.48 kDa) with Coomassie blue staining. **(d)** LC-MS chromatograms from *in vitro* assays (4 h reaction) of *Ch*CYP805A11 and *Ch*NmrA-1 where **3** and NADPH serve as the substrate and the cofactor respectively. Shown are the EICs, illustrating no production of compound **4** ($[M + H]^+$ = *m/z* 348.1805) by *Ch*CYP805A11 and production of compound **5** ($[M + H]^+$ = *m/z* 330.1700) in the presence of both *Ch*CYP805A11 and *Ch*NmrA-1 *in vitro*. These are confirmed by comparison to compounds **4** (blue traces) and **5** (green traces) produced in *N. benthamiana*. EV, yeast microsomes with empty vector (negative control). This experiment was performed at least three times, with similar results observed each time. **(e)** Mirror plots compare the MS/MS fragmentation spectrum of the *in vitro* generated *m/z* 330.1700 product **5** (black traces) with that of compound **5** produced in *N. benthamiana* (green traces) at a collision energy of 30 V, with key experimental fragment ions labeled. Same fragmentation patterns demonstrate that the *in vitro* generated *m/z* 330.1700 product is truly compound **5**. **(f)** Time course of **5** production by *Ch*CYP805A11-enriched yeast microsomes and purified *Ch*NmrA-1 with **3** and NADPH as the substrate and the cofactor respectively, compared to reactions of excluding each enzyme or both enzymes by using EV instead of *Ch*CYP805A11 and boiled *Ch*NmrA-1. The data are reported as mean ± SD of the extracted ion abundance ($n$ = 3 reactions per condition). **(g)** Untargeted metabolite analysis (XCMS) comparing the presence and absence (denatured) of *Ch*NmrA-1 from 4 h *in vitro* assays (sample conditions D vs B, see the legend for panel h) with substrate **3** and cofactor NADPH ($n$ = 3 independent replicates per condition). The unique *m/z* features ($P < 0.1$ and fold change > 5 between samples, see Methods) are shown in ranked order based on their increasing fold change in ion abundance between the two conditions. The mass isotopologues and adducts of the putative product (*m/z* 330.1700, r.t. of 7.1 min) are shown in red. The rest in black are below signal to noise threshold and cannot be identified in the raw LC-MS. r.t., retention time. Note: XCMS comparing the presence and absence of *Ch*CYP805A11 from *in vitro* experiments (sample conditions B vs A, see the legend for panel h) with substrate **3** and cofactor NADPH ($n$ = 3 independent replicates per condition) was also performed; no figure shown due to no unique *m/z* features ($P < 0.1$ and fold change > 5 between samples, see Methods) observed. **(h)** The data are from 4 h *in vitro* reactions and reported as mean ± SD of the extracted ion abundance ($n$ = 3) for the exact ion mass $[M + H]^+$ corresponding to each compound. The left panel shows consumption of compound **3**, while the middle and right panels show production of **5** and **5**-D respectively. All six conditions are labeled A to F (see legend). Orange bars indicate samples treated with $NaBD_3CN$ reducing agent after the reaction, while grey bars indicate samples without $NaBD_3CN$ treatment. **(i)** Isotope envelope of **5** in centroid mode, comparing sample B treated with $NaBD_3CN$ (red) and sample D without $NaBD_3CN$ treatment (black). Sample B shows a clear shift in the isotope envelope, with increased abundance of the M + 1 isotopologue consistent with deuterium incorporation during reduction of imine in **4'** by $NaBD_3CN$. **(j)** Comparison of MS/MS spectra of enzymatically produced and chemically reduced **5**-D from sample B treated with $NaBD_3CN$ (red traces), enzymatically produced **5**-$M_1$ (M + 1 isotopolgue) from sample D without $NaBD_3CN$ treatment (violet traces), and enzymatically produced **5** (parent isotopologue) from sample D without $NaBD_3CN$ treatment (black traces) at a collision energy of 30 V. All three spectra exhibit similar fragmentation patterns, with a mass shift of + 1 Da and corresponding fragment ion differences indicating the position of deuterium incorporation. **(k)** Putative structures for ion fragments generated from MS/MS analysis of **5**-D ($[M + H]^+$ = *m/z* 331.1763, r.t. = 7.14 min). The theoretical and experimental fragment ions support the proposed structure.

**Note regarding the rationale for selection of expression hosts:** *Ch*CYP805A11 was expressed in *S. cerevisiae* for microsome preparation because cytochrome P450 enzymes are ER-membrane-bound and generally require a eukaryotic expression system. Yeast-derived microsomes were also essential for distinguishing enzymatic products from those arising due to promiscuous plant reductases, clarifying that compound **4** is a plant-derived side product of CET biosynthesis, rather than a bona fide pathway intermediate. *Ch*NmrA-1 was expressed and purified from *E. coli* because there have been many precedence of soluble or non-membrane bound plant enzymes that have efficiently and successfully been expressed in and purified from bacterial systems. For this reason, *Ch*NmrA-1 was first attempted to be expressed in and purified from *E. coli* for *in vitro* assays, which was found to be successful as discussed in the manuscript. Heterologous host systems used for expression of CET biosynthetic enzymes from *C. harringtonia* are summarized in **Supplementary Table 4**.

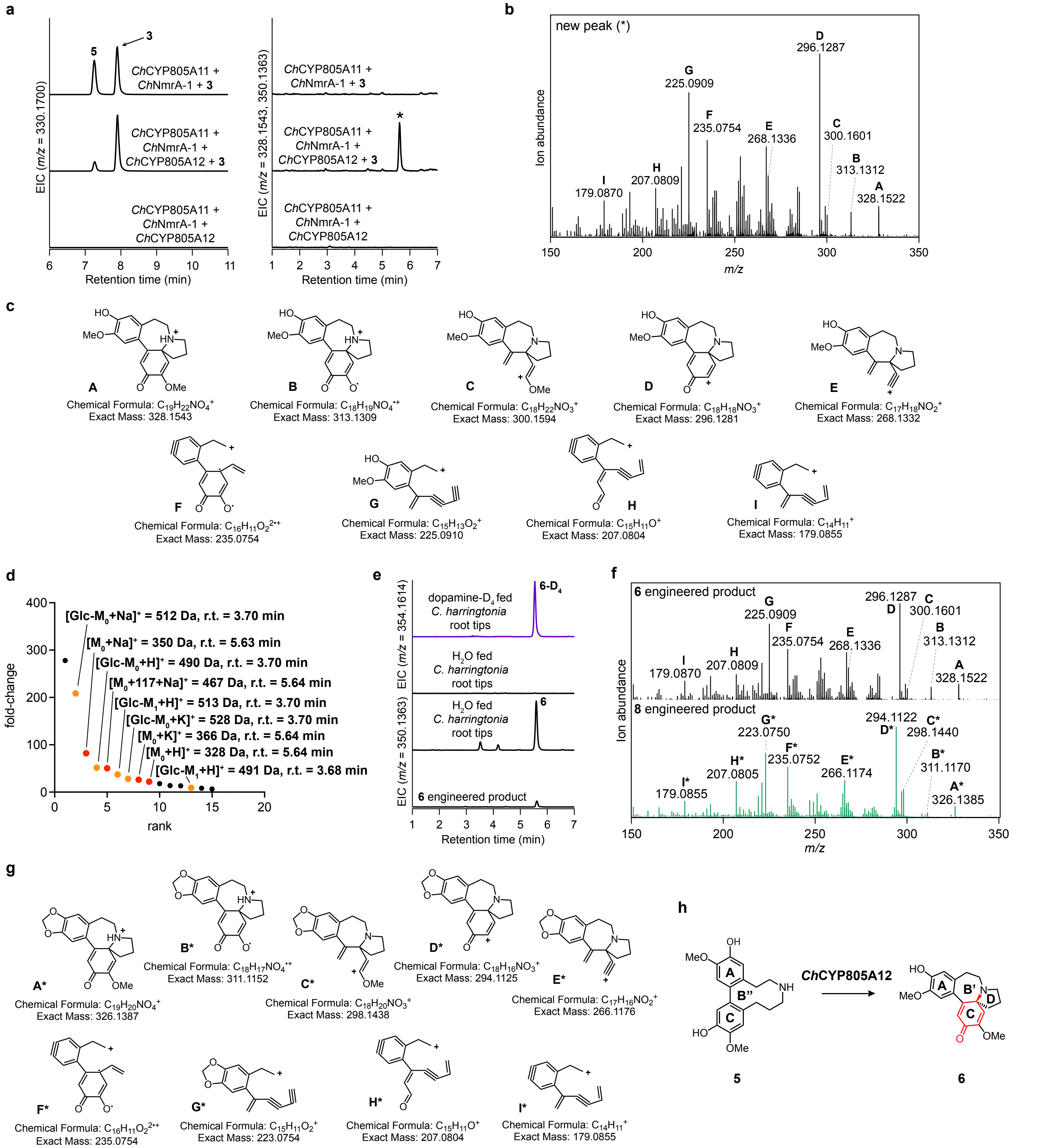


**Extended Data Figure 4. Characterization of *Ch*CYP805A12 and its resulting product (6). (a)** Addition of *Ch*CYP805A12 to the *N. benthamiana* transient expression system with co-infiltrated **3** leads to consumption of **5** ($[M + H]^+ = m/z$ 330.1700) and production of a new compound (*) that corresponds to a loss of two hydrogens ($[M + H]^+ = m/z$ 328.1543, $[M + Na]^+ = m/z$ 350.1363), as shown by the LC-MS chromatograms. The new compound is dependent on substrate (**5**)**.** This experiment was performed more than three times, with similar results observed each time. **(b)** MS/MS fragmentation spectrum of the generated *m/z* 328.1543 product (*) at a collision energy of 30 V, with key experimental fragment ions labeled. **(c)** Putative structures for ion fragments generated from MS/MS analysis of the new product ($[M + H]^+ = m/z$ 328.1543). The theoretical and experimental fragment ions support the proposed structure. **(d)** Untargeted metabolite analysis (XCMS) comparing the presence and absence of *Ch*CYP805A12 in the *N. benthamiana* transient co-expression system with co-infiltrated substrate **3** (*n* = 3 independent replicates per condition). The unique *m/z* features ($P < 0.15$ and fold change > 5 between samples, see Methods) are shown in ranked order based on their increasing fold change in ion abundance between the two conditions. Different adducts of the putative product (*m/z* 328.1543, r.t. of 5.6 min) are shown in red; the mass isotopologues and adducts of the glycosylated putative product (*m/z* 490.2070, r.t. of 3.7 min) are shown in yellow-orange. The rest in black are mostly below signal to noise threshold and cannot be identified in the raw LC-MS, independent of the initial substrate **3**, or non-native to *C. harringtonia*. r.t., retention time. **(e)** The new engineered product **6** ($[M + Na]^+$ = *m/z* 350.1363) corresponds to a metabolite found in the root tips of *C. harringtonia*, as shown by the LC-MS chromatograms. Deuterium-labeled analog of **6** (**6**-$D_4$) is observed in dopamine-$D_4$ fed *C. harringtonia* root tips ($[M + Na]^+$ = *m/z* 354.1614), shown in violet traces. **(f)** Comparison of the MS/MS spectrum of enzymatically produced **6** (black traces) with that of enzymatically produced **8** (green traces) in *N. benthamiana* at a collision energy of 30V. The two spectra have very similar fragmentation patterns with a major difference of two hydrogens in mass for certain fragments, suggesting that compounds **6** and **8** are structurally similar and most likely differ by the presence or absence of the methylenedioxy bridge. See Extended Data Fig. 6 **(g)** Putative structures for ion fragments generated from MS/MS analysis of compound **8** ($[M + H]^+$ = *m/z* 326.1387). The theoretical and experimental fragment ions support the proposed structure. **(h)** Proposed reaction catalyzed by *Ch*CYP805A12, as supported by MS/MS fragmentation pattern of the new product (**6**) and its comparison to that of the structurally related CET intermediate **8**. See Extended Data Fig. 6.

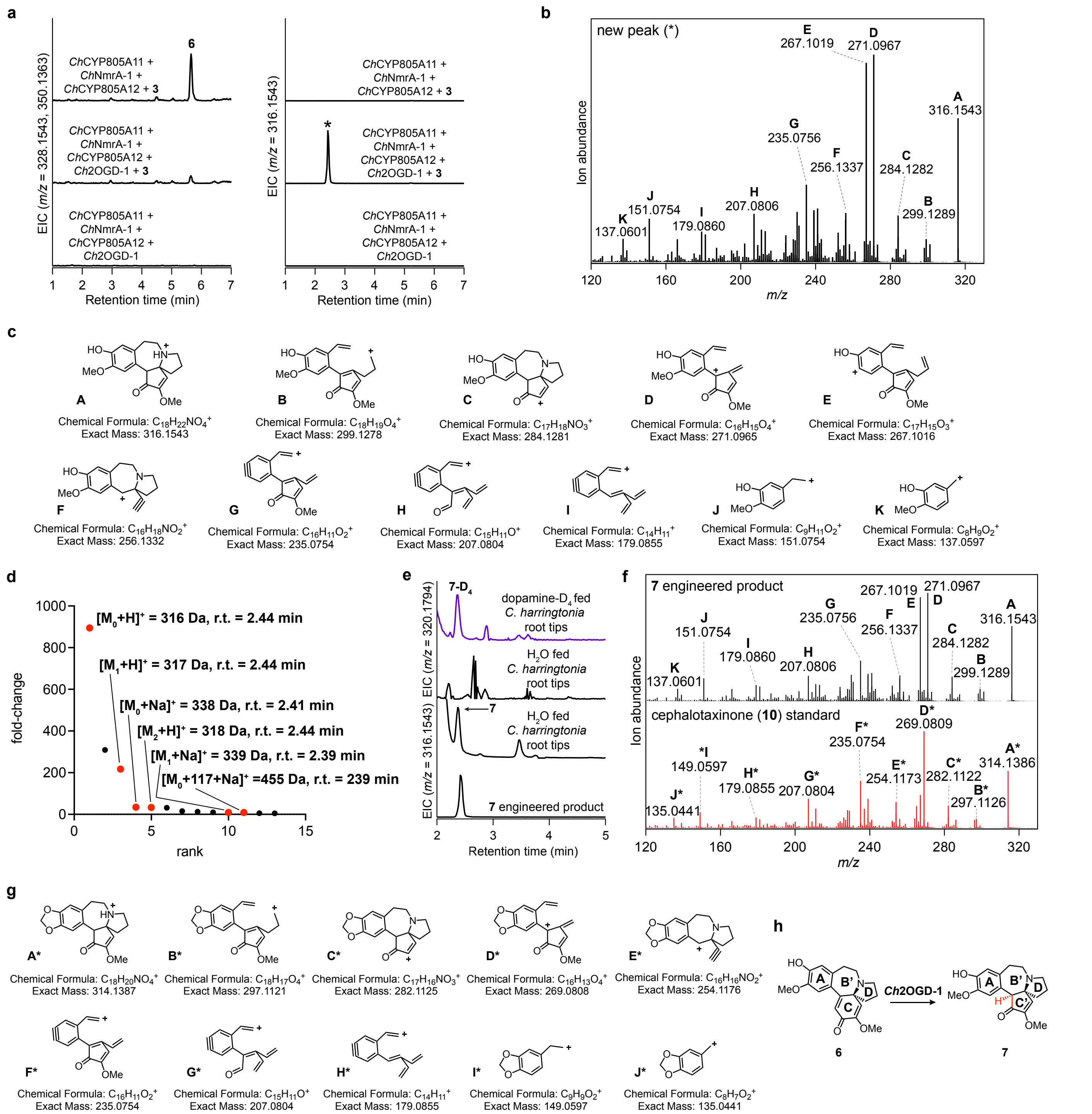


**Extended Data Figure 5. Characterization of *Ch*2OGD-1 and its resulting product (7). (a)** Addition of *Ch*2OGD-1 to the *N. benthamiana* transient expression system with co-infiltrated **3** leads to consumption of **6** ($[M + H]^+ = m/z$ 328.1543) and production of a new compound (*) that corresponds to a loss of carbon ($[M + H]^+ = m/z$ 316.1543), as shown by the LC-MS chromatograms. The new compound is dependent on substrate (**6**). This experiment was performed more than three times, with similar results observed each time. **(b)** MS/MS fragmentation spectrum of the generated *m/z* 316.1543 product (*) at a collision energy of 30 V, with key experimental fragment ions labeled. **(c)** Putative structures for ion fragments generated from MS/MS analysis of the new product ($[M + H]^+ = m/z$ 316.1543). The theoretical and experimental fragment ions support the proposed structure. **(d)** Untargeted metabolite analysis (XCMS) comparing the presence and absence of *Ch*2OGD-1 in the *N. benthamiana* transient co-expression system with co-infiltrated substrate **3** ($n = 3$ independent replicates per condition). The unique *m/z* features ($P < 0.1$ and fold change > 5 between samples, see Methods) are shown in ranked order based on their increasing fold change in ion abundance between the two conditions. The mass isotopologues and adducts of the putative product (*m/z* 316.1543, r.t. of 2.4 min) are shown in red. The rest in black are mostly below signal to noise threshold and cannot be identified in the raw LC-MS, independent of the initial substrate **3**, or non-native to *C. harringtonia*. r.t., retention time. **(e)** The new engineered product **7** ($[M + H]^+ = m/z$ 316.1543, r.t. = 2.4 min) corresponds to a metabolite found in the root tips of *C. harringtonia*, as shown by the LC-MS chromatograms. Deuterium-labeled analog of **7** (**7**-$D_4$) is observed in dopamine-$D_4$ fed *C. harringtonia* root tips ($[M + H]^+$ = *m/z* 320.1794, r.t. = 2.4 min), shown in violet traces. **(f)** Comparison of the MS/MS spectrum of enzymatically produced **7** in *N. benthamiana* (black traces) with that of cephalotaxinone standard (red traces) at a collision energy of 30V. The two spectra have very similar fragmentation patterns with a major difference of two hydrogens in mass for certain fragments, suggesting that compound **7** and cephalotaxinone are structurally similar and most likely differ by the presence or absence of the methylenedioxy bridge. See Extended Data Fig. 7 **(g)** Putative structures for ion fragments generated from MS/MS analysis of cephalotaxinone ($[M + H]^+$ = *m/z* 314.1387). **(h)** Proposed reaction catalyzed by *Ch*2OGD-1, as supported by MS/MS fragmentation pattern of the new product (**7**) and its comparison to that of the structurally related CET intermediate, cephalotaxinone. See Extended Data Fig. 7.

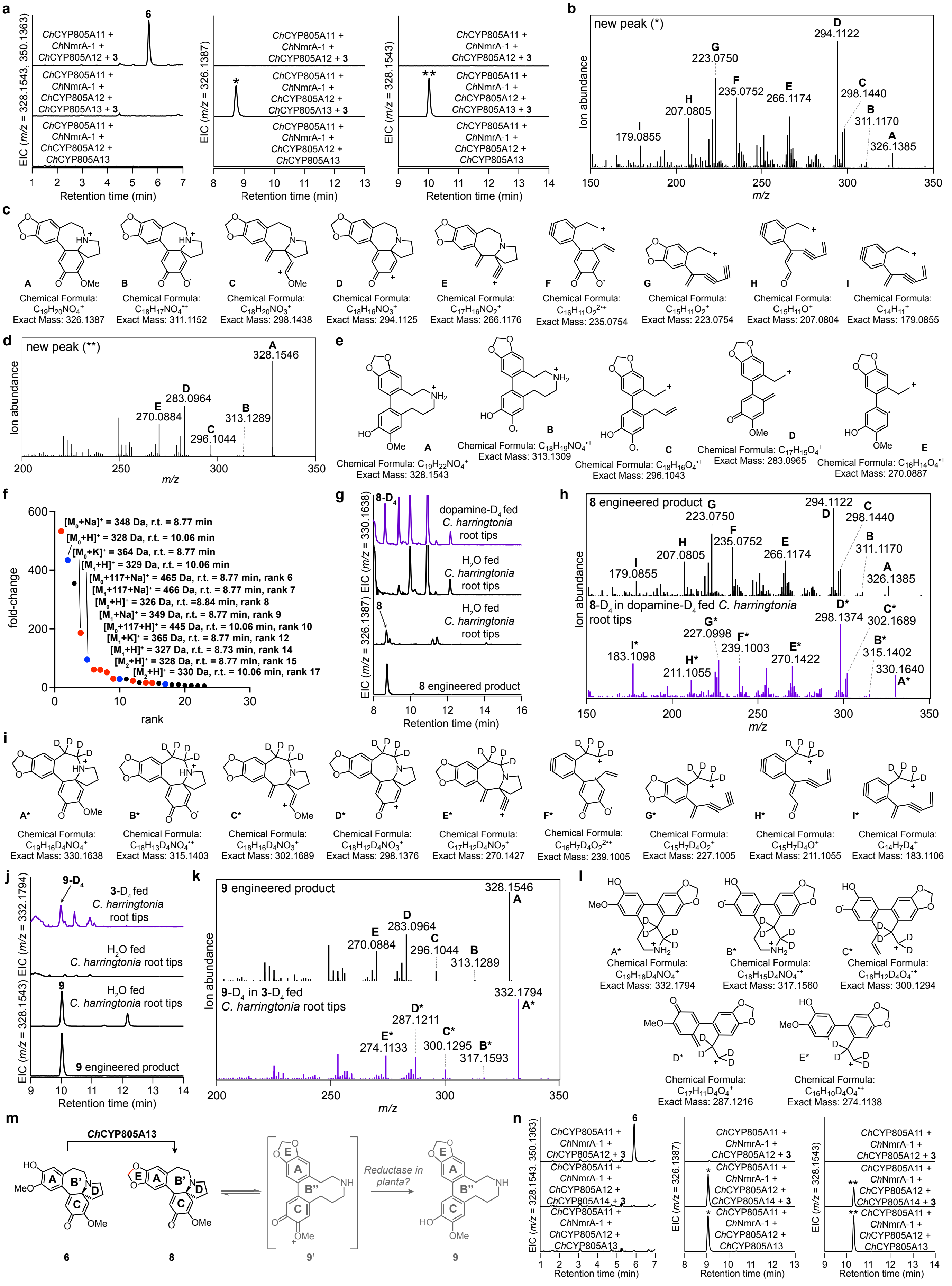

a
ChCYP805A11 + ChNmrA-1 + ChCYP805A12 + 3
ChCYP805A11 + ChNmrA-1 + ChCYP805A12 + ChCYP805A13 + 3
ChCYP805A11 + ChNmrA-1 + ChCYP805A12 + ChCYP805A13
Retention time (min)
b
new peak (*)
Ion abundance
m/z
c
d
new peak (**)
e
f
fold-change
rank
g
dopamine-D4 fed C. harringtonia root tips
H2O fed C. harringtonia root tips
8 engineered product
h
8 engineered product
8-D4 in dopamine-D4 fed C. harringtonia root tips
i
j
3-D4 fed C. harringtonia root tips
9 engineered product
k
9 engineered product
9-D4 in 3-D4 fed C. harringtonia root tips
l
m
ChCYP805A13
Reductase in planta?
n
ChCYP805A11 + ChNmrA-1 + ChCYP805A12 + ChCYP805A14 + 3

**Extended Data Figure 6. Characterization of *Ch*CYP805A13 and its resulting products (8) and (9). (a)** Addition of *Ch*CYP805A13 to the *N. benthamiana* transient expression system (all without *Ch*2OGD-1) with co-infiltrated **3** leads to consumption of **6** ($[M + H]^+$ = *m/z* 328.1543, $[M + Na]^+$ = *m/z* $^+$ = *m/z* 350.1363, r.t. = 5.6 min) and production of two new compounds (*) and (**) that correspond to a loss of two hydrogens ($[M + H]^+$ = *m/z* 326.1387, r.t. = 8.8 min) and no change in mass ($[M + H]^+$ = *m/z* 328.1543, r.t. = 10.1 min) respectively, as shown by the LC-MS chromatograms. The two new compounds are dependent on substrate (**6**). This experiment was performed more than three times, with similar results observed each time. **(b)** MS/MS fragmentation spectrum of the generated *m/z* 326.1387 product (*) at a collision energy of 30 V, with key experimental fragment ions labeled. **(c)** Putative structures for ion fragments generated from MS/MS analysis of the new product ($[M + H]^+$ = *m/z* 326.1387). The theoretical and experimental fragment ions support the proposed structure. **(d)** MS/MS fragmentation spectrum of the generated *m/z* 328.1387 product (**) at a collision energy of 30 V, with key experimental fragment ions labeled. **(e)** Putative structures for ion fragments generated from MS/MS analysis of another new product ($[M + H]^+$ = *m/z* 328.1543, r.t. = 10.1 min). The theoretical and experimental fragment ions support the proposed structure. **(f)** Untargeted metabolite analysis (XCMS) comparing the presence and absence of *Ch*CYP805A13 in the *N. benthamiana* transient co-expression system (all without *Ch*2OGD-1) with co-infiltrated substrate **3** ($n$ = 3 independent replicates per condition). The unique *m/z* features ($P < 0.1$ and fold change > 5 between samples, see Methods) are shown in ranked order based on their increasing fold change in ion abundance between the two conditions. The mass isotopologues and adducts of the putative product (*m/z* 326.1387, r.t. of 8.8 min) and the other product (*m/z* 328.1543, r.t. of 10.1 min) are shown in red and blue respectively. The rest in black are mostly below signal to noise threshold and cannot be identified in the raw LC-MS, independent of the initial substrate **3**, or non-native to *C. harringtonia*. **(g)** The new engineered product **8** ($[M + H]^+$ = *m/z* 326.1387, r.t. = 8.8 min) corresponds to a metabolite found in the root tips of *C. harringtonia*, as shown by the LC-MS chromatograms. Deuterium-labeled analog of **8** (**8**-$D_4$) is observed in dopamine-$D_4$ fed *C. harringtonia* root tips ($[M + H]^+$ = *m/z* 330.1638, r.t. = 8.8 min), shown in violet traces. **(h)** Comparison of the MS/MS spectrum of enzymatically produced **8** in *N. benthamiana* (black traces) with that of **8**-$D_4$ from dopamine-$D_4$ fed *C. harringtonia* root tips (violet traces) at a collision energy of 30V. The two spectra have very similar fragmentation patterns with a major difference of four hydrogens in mass. **(i)** Putative structures for ion fragments generated from MS/MS analysis of **8**-$D_4$ ($[M + H]^+$ = *m/z* 330.1638, r.t. = 8.8 min). **(j)** Another new engineered product **9** ($[M + H]^+$ = *m/z* 328.1543, r.t. = 10.1 min) corresponds to a metabolite found in the root tips of *C. harringtonia*, as shown by the LC-MS chromatograms. Deuterium-labeled analog of **9** (**9**-$D_4$) is observed in **3**-$D_4$ fed *C. harringtonia* root tips ($[M + H]^+$ = *m/z* 332.1794, r.t. = 10.1 min), shown in violet traces. **(k)** Comparison of the MS/MS spectrum of enzymatically produced **9** in *N. benthamiana* (black traces) with that of **9**-$D_4$ from **3**-$D_4$ fed *C. harringtonia* root tips (violet traces) at a collision energy of 30V. The two spectra have very similar fragmentation patterns with a major difference of four hydrogens in mass. **(l)** Putative structures for ion fragments generated from MS/MS analysis of **9**-$D_4$ ($[M + H]^+$ = *m/z* 332.1794, r.t. = 10.1 min). **(m)** Proposed reaction catalyzed by *Ch*CYP805A13, as supported by MS/MS fragmentation patterns of the new products **8** and **9** and their comparison to those of the corresponding four-deuterated forms (**8**-$D_4$ and **9**-$D_4$). **(n)** *Ch*CYP805A14, a truncated variant of *Ch*CYP805A13 lacking the N-terminal transmembrane domain, catalyzes the same reaction as *Ch*CYP805A13, as shown by the LC-MS chromatograms.

**Note regarding *Ch*CYP805A14:** *Ch*CYP805A14 in the candidate gene list (Fig. 3c) was found to catalyze the same reaction as *Ch*CYP805A13 (**Extended Data Fig. 6n**). Sequence comparison revealed that *Ch*CYP805A14 is a truncated variant of *Ch*CYP805A13, lacking the N-terminal transmembrane domain (approximately the first 49 amino acids) with a difference in 16 amino acid residues (**Supplementary Table 5**). This result suggested that the lack of N-terminal domain, which is often essential for cytochrome P450 trafficking and anchoring to endoplasmic reticulum (ER) membrane, does not affect the catalytic activity of *Ch*CYP805A14 in *N. benthamiana*.

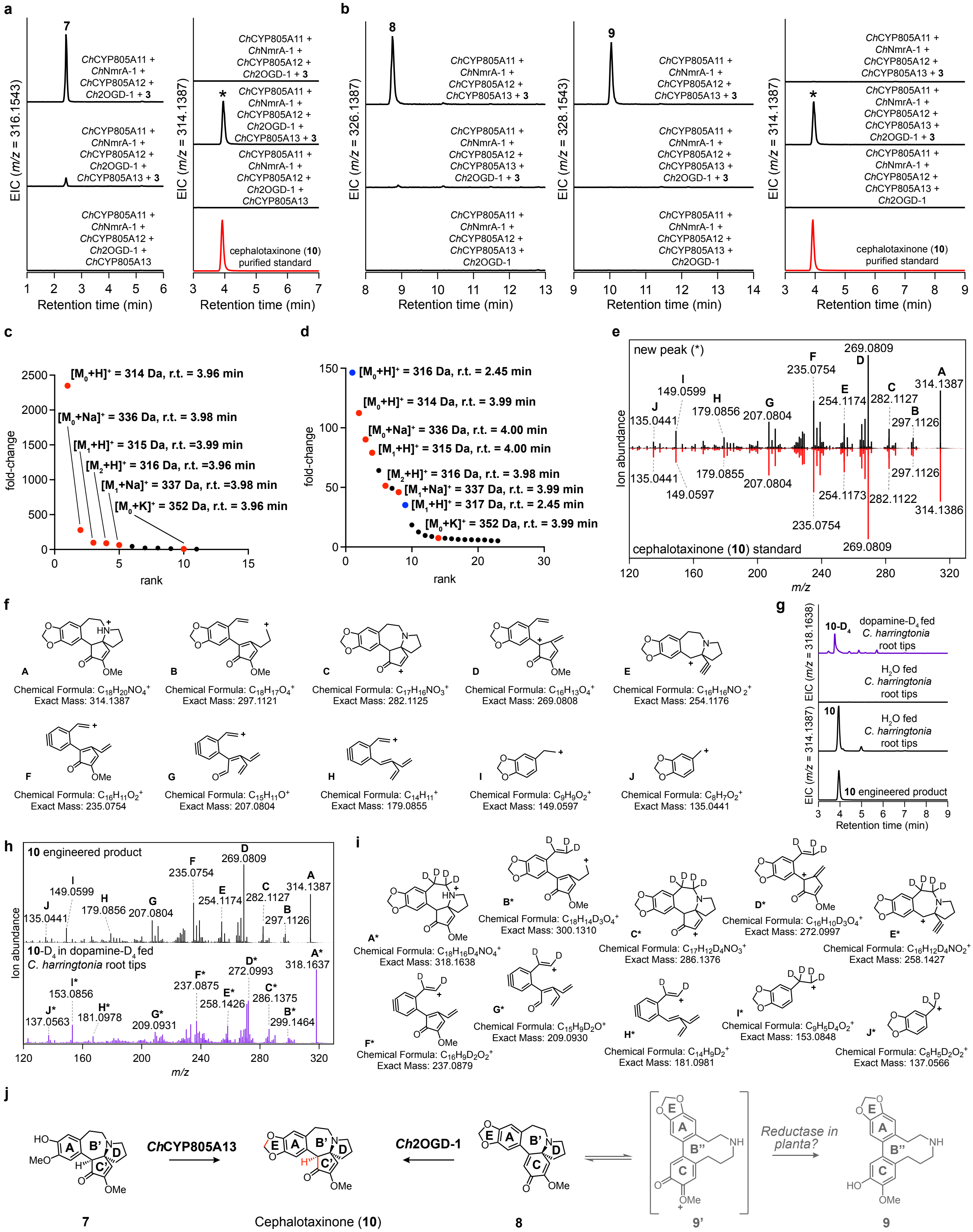


**Extended Data Figure 7. Characterization of *Ch*CYP805A13 and *Ch*2OGD-1 together and their resulting final product cephalotaxinone (10). (a)** Addition of *Ch*CYP805A13 to the *N. benthamiana* transient expression system (all four enzymes without *Ch*CYP805A13) with co-infiltrated **3** leads to consumption of **7** ($[M + H]^+$ = *m/z* 316.1543) and production of a new compound (*) that corresponds to a loss of two hydrogens ($[M + H]^+$ = *m/z* 314.1387), as shown by the LC-MS chromatograms. The new compound is dependent on substrate (**7**)**.** This experiment was performed more than three times, with similar results observed each time. **(b)**

Addition of *Ch*2OGD-1 to the *N. benthamiana* transient expression system (all four enzymes without *Ch*2OGD-1) with co-infiltrated **3** leads to consumption of **8** ($[M + H]^+$ = *m/z* 326.1387, r.t. = 8.8 min), decrease of **9** ($[M + H]^+$ = *m/z* 328.1543, r.t. = 10.1 min), and production of a new compound (*) that corresponds to a loss of carbon from compound **8** ($[M + H]^+$ = *m/z* 314.1387), as shown by the LC-MS chromatograms. The new compound is dependent on substrate (**8**). This experiment was performed more than three times, with similar results observed each time. **(c)** Untargeted metabolite analysis (XCMS) comparing the presence and absence of *Ch*CYP805A13 in the *N. benthamiana* transient co-expression system (all four enzymes without *Ch*CYP805A13) with co-infiltrated substrate **3** (*n* = 3 independent replicates per condition). The unique *m/z* features ($P < 0.1$ and fold change > 5 between samples, see Methods) are shown in ranked order based on their increasing fold change in ion abundance between the two conditions. The mass isotopologues and adducts of the putative product (*m/z* 314.1387, r.t. of 4.0 min) are shown in red. The rest in black are mostly below signal to noise threshold and cannot be identified in the raw LC-MS, independent of the initial substrate **3**, or non-native to *C. harringtonia*. **(d)** Untargeted metabolite analysis (XCMS) comparing the presence and absence of *Ch*2OGD-1 in the *N. benthamiana* transient co-expression system (all without *Ch*2OGD-1) with co-infiltrated substrate **3** (*n* = 3 independent replicates per condition). The unique *m/z* features ($P < 0.1$ and fold change > 5 between samples, see Methods) are shown in ranked order based on their increasing fold change in ion abundance between the two conditions. The mass isotopologues and adducts of the putative product (*m/z* 314.1387, r.t. of 4.0 min) are shown in red. The mass isotopologues of compound **7** (*m/z* 316.1543, r.t. of 2.4 min) are shown in blue; this arises from the remaining unreacted **7** produced from co-expression of *Ch*2OGD-1 with *Ch*CYP805A11, *Ch*NmrA-1, and *Ch*CYP805A12. The rest in black are mostly below signal to noise threshold and cannot be identified in the raw LC-MS, independent of the initial substrate **3**, or non-native to *C. harringtonia*. r.t., retention time. **(e)** Mirror plots compare the MS/MS fragmentation spectrum of the generated *m/z* 314.1387 product (*), shown in black traces, with that of an authentic purified cephalotaxinone (**10**), shown in red traces, at a collision energy of 30 V, with key experimental fragment ions labeled. **(f)** Putative structures for ion fragments generated from MS/MS analysis of the new product ($[M + H]^+$ = *m/z* 314.1387), cephalotaxinone. The theoretical and experimental fragment ions support the proposed structure. **(g)** The new engineered product **10** ($[M + H]^+$ = *m/z* 314.1387, r.t. = 4.0 min) corresponds to a metabolite found in the root tips of *C. harringtonia*, as shown by the LC-MS chromatograms. Deuterium-labeled analog of **10** (**10**-$D_4$) is observed in dopamine-$D_4$ fed *C. harringtonia* root tips ($[M + H]^+$ = *m/z* 318.1638, r.t. = 4.0 min), shown in violet traces. **(h)** Comparison of the MS/MS spectrum of enzymatically produced **10** in *N. benthamiana* (black traces) with that of **10**-$D_4$ from dopamine-$D_4$ fed *C. harringtonia* root tips (violet traces) at a collision energy of 30V. **(i)** Putative structures for ion fragments generated from MS/MS analysis of **10**-$D_4$ ($[M + H]^+$ = *m/z* 318.1638, r.t. = 4.0 min). **(j)** Proposed reactions catalyzed by *Ch*CYP805A13 and *Ch*2OGD-1 to form cephalotaxinone (**10**) from compounds **7** and **8** respectively, as supported by MS/MS fragmentation of the new engineered product and its comparison to an authentic **10** standard.

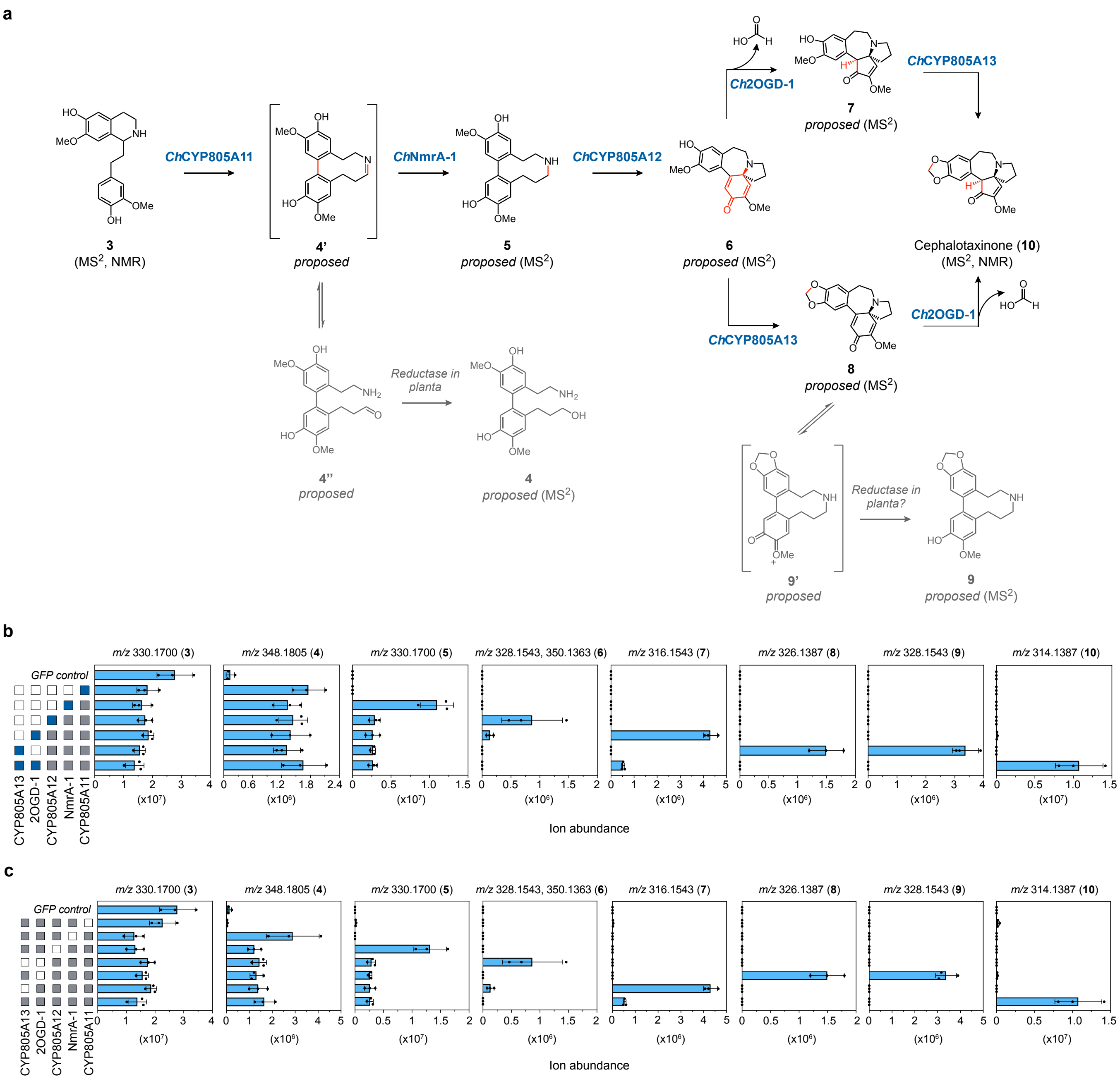

**Extended Data Figure 8. Discovery of a pathway for CET alkaloid biosynthesis, from compound 3 to cephalotaxinone. (a)** Proposed biosynthetic pathway of CET reconstituted in *N. benthamiana* through transient co-expression of five identified biosynthetic genes from *C. harringtonia*, illustrating the stepwise conversion from compound **3** to cephalotaxinone (**10**). All the structures are supported by LC-MS/MS analyses, with the first and final intermediates (**3** and cephalotaxinone (**10**)) further supported by authentic synthesized and purified standards, including NMR analyses. **(b)** Quantification of each CET biosynthetic intermediate produced from sequential addition of each enzyme in *N. benthamiana* transient expression system. Gray boxes to the left of the graphs indicate biosynthetic genes that are co-expressed in *N. benthamiana* leaves; blue box represents the final acting enzyme. **(c)** Accumulation of proposed CET pathway intermediates in dropout experiments, in which each individual enzyme was removed from the engineered pathway to cephalotaxinone (**10**) in *N. benthamiana*. Gray boxes to the left of the graphs indicate biosynthetic genes that are co-expressed in *N. benthamiana* leaves; white boxes indicate their absence. **(b-c)** For each intermediate, the data are reported as mean ± SD of the extracted ion abundance ($n$ = 3) for the exact ion mass $[M + H]^+$ (for **6**, both $[M + H]^+$ and $[M + Na]^+$) corresponding to each compound.

**Note regarding *Ch*2OGD-1 and *Ch*CYP805A13:** Steps from **6** to cephalotaxinone (**10**) may occur in bifurcated sequence (**Extended Data Figs. 5-8**). Both *Ch*2OGD-1 and *Ch*CYP805A13 can consume intermediate **6** to produce **7** and **8** respectively. Both routes then converge to form cephalotaxinone. This suggests metabolic flexibility in the late-stage tailoring steps of CET biosynthesis.

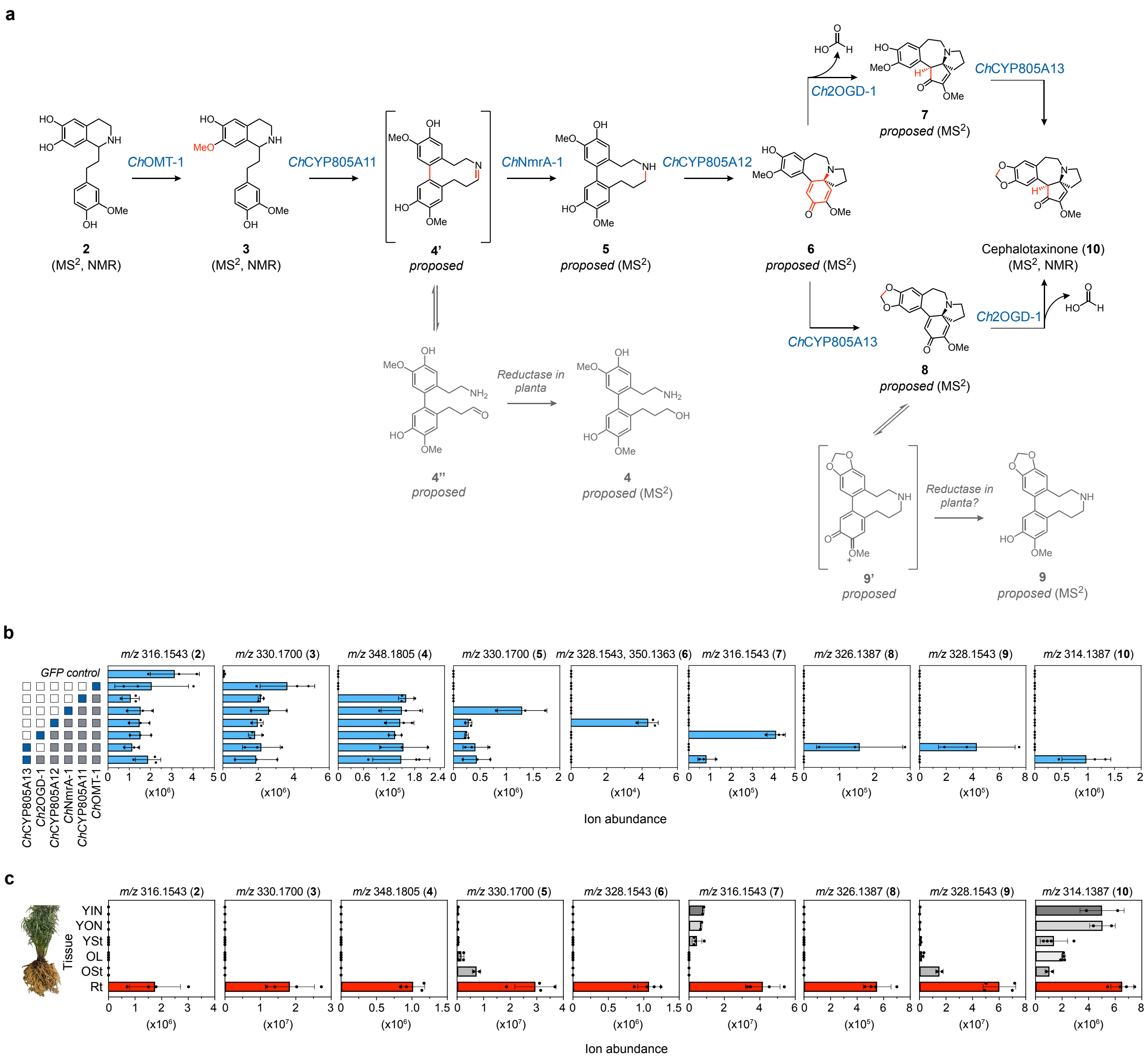


**Extended Data Figure 9. Reconstitution of CET biosynthetic pathway from the earliest phenethylisoquinoline (2) to cephalotaxinone (10) in *N. benthamiana*.** **(a)** Proposed biosynthetic pathway of CET reconstituted in *N. benthamiana* through transient co-expression of all six identified biosynthetic genes from *C. harringtonia*, illustrating the stepwise conversion from compound **2** to cephalotaxinone (**10**). All the structures are supported by LC-MS/MS analyses, with intermediates **2**, **3**, and **10** further supported by authentic synthesized or purified standards, including NMR analyses. **(b)** Gray boxes to the left of the graphs indicate biosynthetic genes that are co-expressed in *N. benthamiana* leaves; blue box represents the final acting enzyme. For each intermediate, the data are reported as mean ± SD of the extracted ion abundance ($n = 3$) for the exact ion mass $[M + H]^+$ (for **6**, both $[M + H]^+$ and $[M + Na]^+$) corresponding to each compound. **(c)** Natural accumulation of CET biosynthetic intermediates in the native plant, *C. harringtonia*, harvested in May 2023. The data are reported as mean ± SD of the extracted ion abundance ($n = 4$ biological replicates for YSt, ON, and Rt, and $n = 2$ for YIN, YON, and OSt) for the exact ion mass $[M + H]^+$ corresponding to each compound. For quantification of cephalotaxinone, a 200-fold dilution was performed to the extracted samples to avoid saturation of the parent isotope peaks in LC-MS.

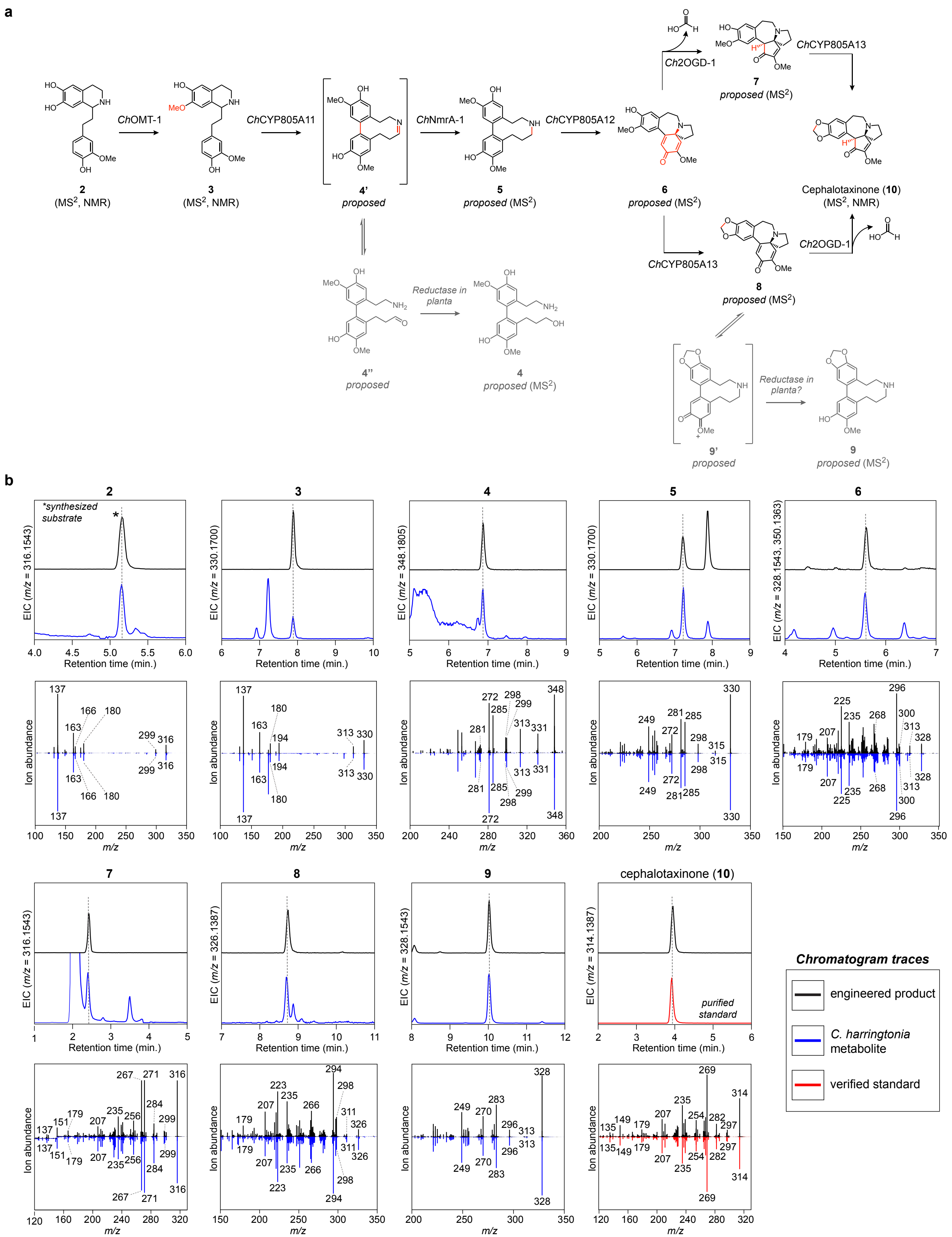


**Extended Data Figure 10. Comparison of intermediates produced in the *N. benthamiana* co-expression system to *C. harringtonia* metabolites. (a)** Proposed biosynthetic pathway of CET from compound **2** to cephalotaxinone (**10**). **(b)** LC-MS comparison of each downstream biosynthetic product of **2** produced heterologously in *N. benthamiana* co-expression system (black traces) with the co-eluting equivalent mass ion detected in *C. harringtonia* root tip extracts (blue traces) or with a verified purified standard (red traces). Mirror plots compare the MS/MS spectra of the co-eluting peaks, demonstrating structural similarity between these compounds. All these compounds produced in the *N. benthamiana* expression system are biologically relevant CET pathway intermediates or side-products. Collision energies for all shown MS/MS analyses were 30 V, except for compounds **2** and **3** (20 V) and compound **4** (15 V). These LC-MS comparisons were performed once with distinct biological replicates of *C. harringtonia* metabolite extractions ($n$ = 17 biological replicates of root tips). All these CET pathway intermediates generated by heterologous expression in *N. benthamiana* exhibited consistent retention times and MS/MS spectra across individual experiments.

**SUPPLEMENTARY INFORMATION**

### Discovery of cephalotaxinone enzymes reveals a whole plant model for homoharringtonine biosynthesis

Yaereen Dho[1] Kevin Smith,[2] & Elizabeth S. Sattely[2,3]*

[1]Department of Chemistry, Stanford University, Stanford, CA 94305

[2]Department of Chemical Engineering, Stanford University, Stanford, CA 94305

[3]HHMI, Stanford University, Stanford, CA 94305

*corresponding author

| | page(s) |
|---|---|
| **Supplementary Figures** | 2 |
| **Supplementary Figures – NMR spectra** | 16 |
| **Supplementary Tables** | 67 |
| **Supplementary References** | 88 |

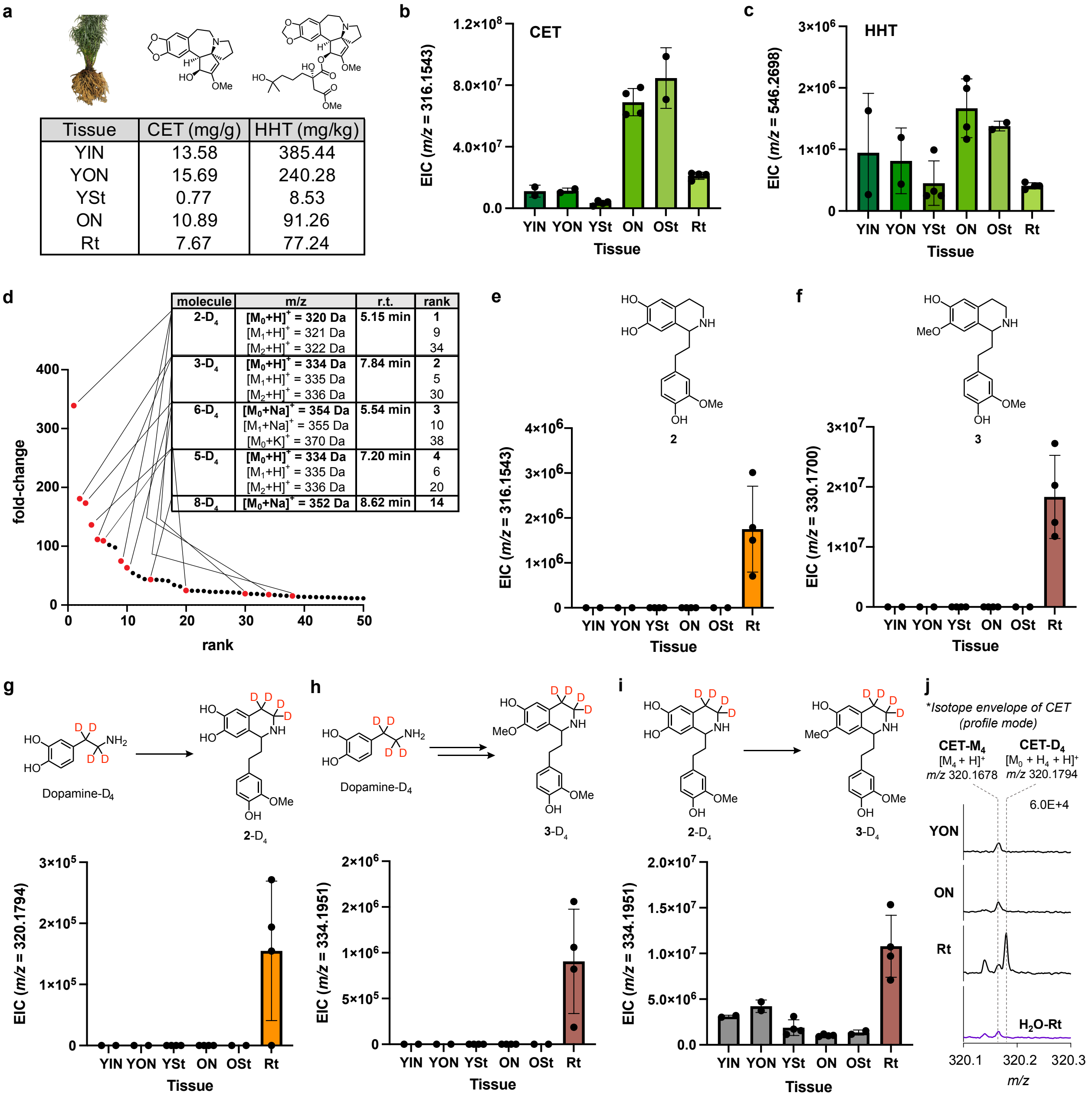


**Supplementary Figure 1. Natural abundance and stable-isotope enrichment of *Cephalotaxus* alkaloids and their intermediates in *C. harringtonia*. (a)** Quantification of CET and HHT per dry weight in different tissues of *C. harringtonia* collected in May 2022 (calculated from average extracted ion chromatogram (EIC) in Fig. 2B using standard curve). **(b-c)** Natural abundance of **(b)** CET ($[M + H]^+$ = *m/z* 316.1543, r.t. = 2 min) and **(c)** HHT ($[M + H]^+$ = *m/z* 546.2698, r.t. = 10.8 min) in the native plant, *C. harringtonia*, harvested in May 2023. **d)** Untargeted metabolite analysis (XCMS) comparing dopamine-$D_4$ fed root tips to water fed root tips of *C. harringtonia* ($n$ = 9 independent replicates for each experimental condition, from May 2022). The unique mass features ($P < 0.2$; fold change > 3.5; 200 < *m/z* < 600) are ranked based on their increasing fold change in abundance between the two conditions, with top 50 shown. Features confirmed to be above signal to noise and represent true mass signatures corresponding to potential intermediates (with four deuteriums, resulting from incorporation of dopamine-D4) in CET biosynthesis are shown in red and their isotopologues and/or adducts in orange. r.t., retention time. While not shown here, CET-$D_4$ ranked 127$^{th}$ out of 148 unique features identified by XCMS with increasing fold change. **(e-f)** Natural accumulation of **(e)** compound **2** ($[M + H]^+$ = *m/z* 316.1543, r.t. = 5.2 min) and **(f)** compound **3** ($[M + H]^+$ = *m/z* 330.1700, r.t. = 7.8 min) in *C. harringtonia*, harvested in May 2023. **(g-h)** Enrichment of **(g)** **2**-$D_4$ ($[M + H]^+$ = *m/z* 320.1794, r.t. = 7.8 min) and **(h)** **3**-$D_4$ ($[M + H]^+$ = *m/z* 334.1951, r.t. = 5.2 min) exclusively observed in the root tips from feeding dopamine-$D_4$. **(i)** O-methylation of **2**-$D_4$, forming **3**-$D_4$, predominantly occurs in the root tips. Feed-in experiments were conducted with tissues harvested in May 2023. **(j)** Isotope envelope of CET in **3**-$D_4$ fed tissues (YON, ON, and Rt) and in $H_2O$ fed Rt in profile mode, focusing on the fourth isotope peak. In the **3**-$D_4$ fed root tip (Rt) sample, an additional fourth isotope peak appears at *m/z* 320.1794, corresponding to CET-$D_4$ ($[M_0 + H_4 + H]^+$) due to four deuterium incorporation, which is distinct from the natural carbon isotope peak at *m/z* 320.1678, corresponding to CET-$M_4$ ($[M_4 + H]^+$). Feed-in experiment was performed in June 2024. All the bar graphs are reported as mean ± SD ($n$ = 4 biological replicates for YSt, ON, and Rt, and $n$ = 2 for YIN, YON, and OSt). Young inner needles (YIN), young outer needles (YON), Young stems (YSt), old needles (ON), old stems (OSt), and root tips (Rt).

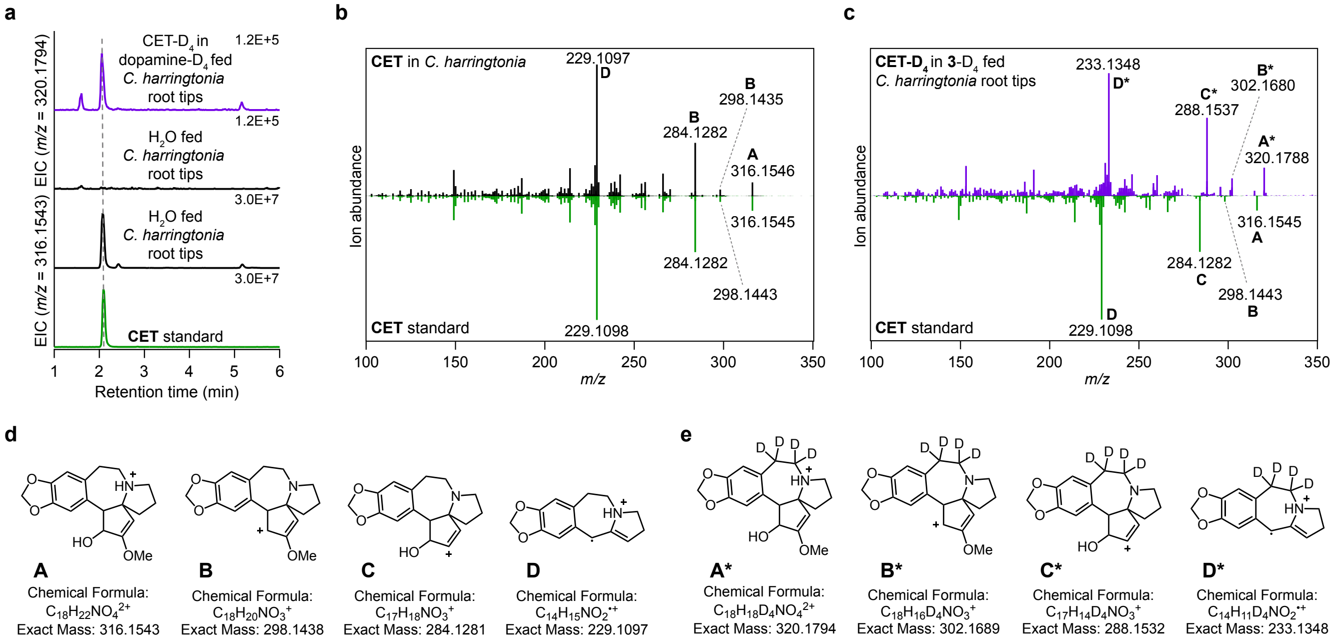


**Supplementary Figure 2. Verification of CET-D4 enrichment in *C. harringtonia* root tips from stable isotope labeled precursor studies. (a)** Deuterium-labeled analog of CET ($[M + H]^+$ = *m/z* 320.1794, r.t. = 2.1 min) is observed in dopamine-D4 fed *C. harringtonia* root tips (violet traces) and absent in water-fed root tip samples (black traces), as compared to CET found in *C. harringtonia* (black traces) and CET standard (green traces). The absence of CET-D4 in water control indicates that CET-D4 observed in dopamine-D4 treated root tips is derived truly from dopamine-D4. Similar result of enrichment of CET-D4 was observed with feeding **2**-D4 or **3**-D4 to *C. harringtonia* root tips, as shown in **Fig. 2** and **Supplementary Fig. 1**. **(b-c)** MS/MS fragmentation spectra of **(b)** CET in root tip extract of *C. harringtonia* (black traces) and **(c)** CET-D4 generated from feeding **3**-D4 (dopamine-D4 or **2**-D4) to *C. harringtonia* root tips (violet traces) and their comparison to the MS/MS fragmentation spectrum of authentic CET standard (green traces) at a collision energy of 35 V, with key experimental fragment ions labeled. **(d-e)** Putative structures for ion fragments generated from MS/MS analysis of **(d)** CET ($[M + H]^+$ = *m/z* 316.1543) and **(e)** CET-D4 ($[M + H]^+$ = *m/z* 320.1794) generated from feeding dopamine-D4, **2**-D4, or **3**-D4 to *C. harringtonia* root tips. Experimental and theoretical fragment ions support the proposed structures, including both unlabeled and deuterium-labeled derivatives.

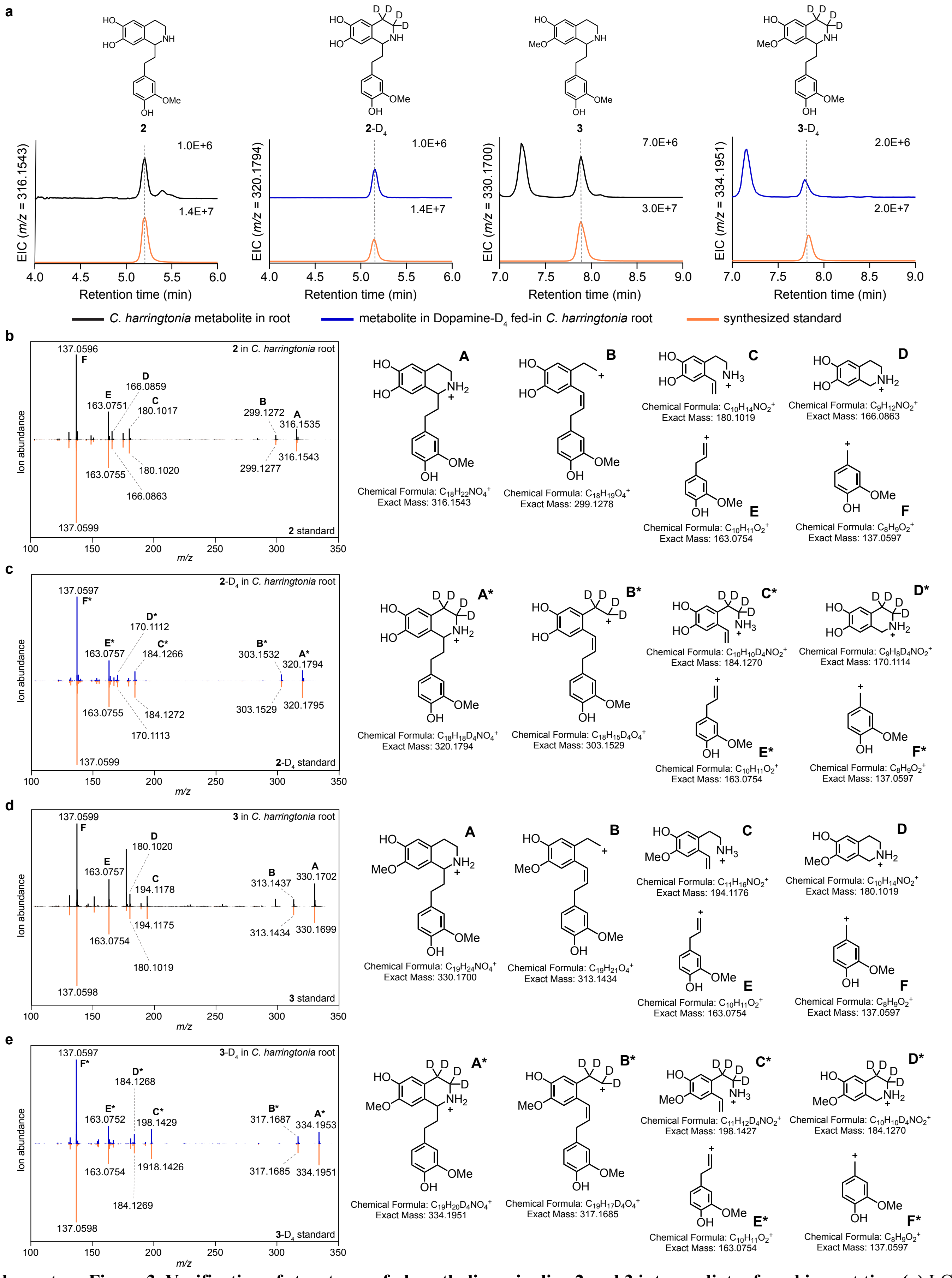


**Supplementary Figure 3. Verification of structures of phenethylisoquinoline 2 and 3 intermediates found in root tips. (a)** LC-MS chromatograms (EICs) showing detection of compound **2** ($[M + H]^+$ = *m/z* 316.1543, r.t. = 5.2 min), compound **2**-$D_4$ ($[M + H]^+$ =

*m/z* 320.1794, r.t. = 5.2 min), compound **3** ($[M + H]^+$ = *m/z* 330.1700, r.t. = 7.8 min), and compound **3**-$D_4$ ($[M + H]^+$ = *m/z* 334.1951, r.t. = 7.8 min) in *C. harringtonia* root-tip extract and comparison with synthesized standards. Deuterium-labeled analogs (**2**-$D_4$ and **3**-$D_4$) are from root-tip samples fed with dopamine-$D_4$. **(b-e)** MS/MS fragmentation spectra of **(b) 2**, **(c) 2**-$D_4$, **(d) 3**-$D_4$ in root-tip extract and comparison with synthesized standards at a collision energy of 20V, with putative structures for ion fragments for each compound. Experimental and theoretical fragment ions support the proposed structures, including both unlabeled and deuterium-labeled derivatives. For all the spectra, natural metabolites observed in *C. harringtonia* root tips are shown in black, metabolite in Dopamine-$D_4$ fed-in *C. harringtonia* root tips in blue, and synthesized standards in orange.

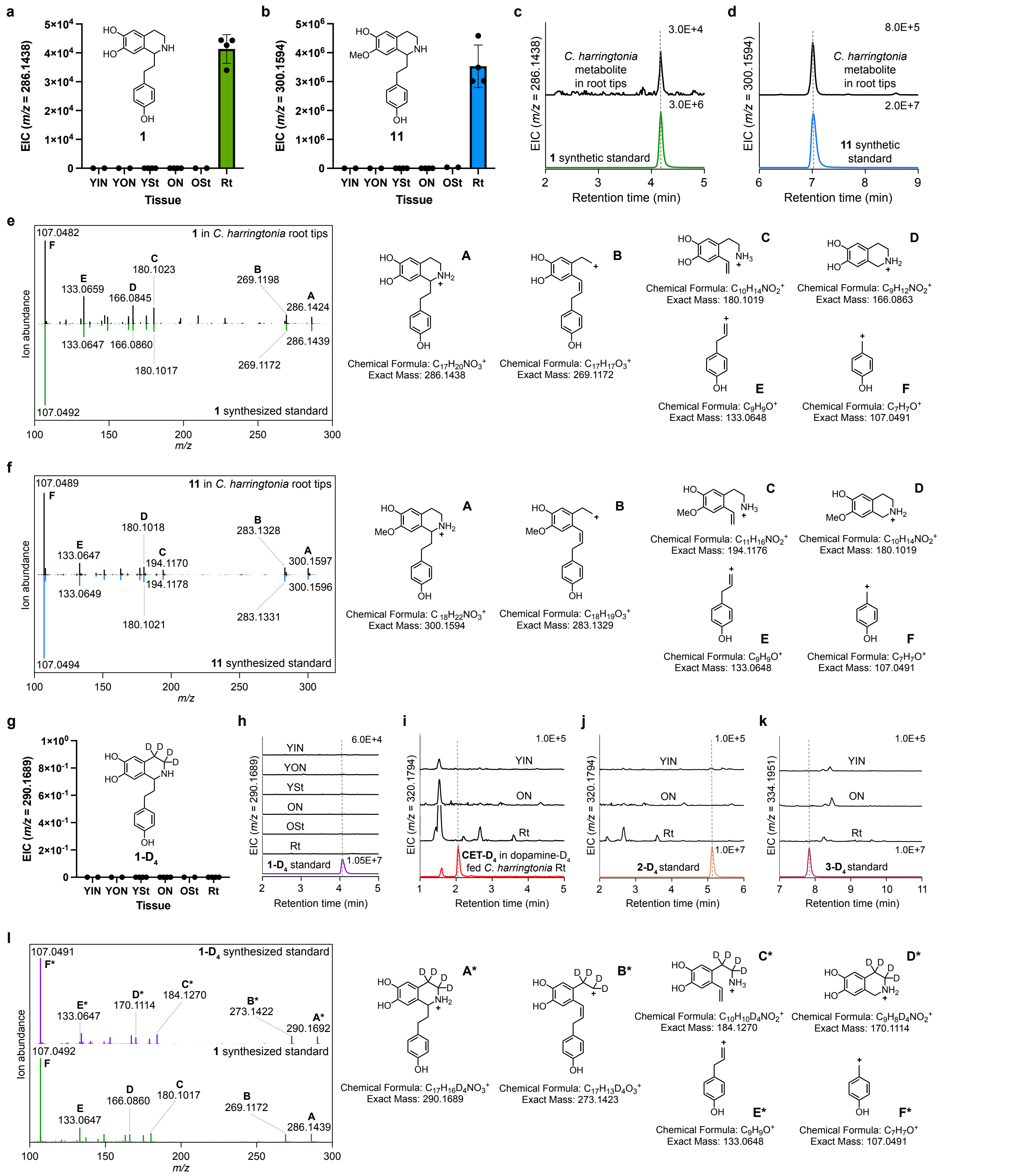


**Supplementary Figure 4. Characterization of other phenethylisoquinolines 1 and 11 in *C. harringtonia*.** **(a-b)** Natural accumulation of **(a)** compound **1** ($[M + H]^+$ = *m/z* 286.1438) and **(b)** compound **11** ($[M + H]^+$ = *m/z* 300.1594) in *C. harringtonia*. **(c-d)** LC-MS chromatograms (EICs) showing detection of **(c)** compound **1** ($[M + H]^+$ = *m/z* 286.1438, r.t. = 4.1 min) and **(d)** compound **11** ($[M + H]^+$ = *m/z* 300.1594, r.t. = 7.0 min) in *C. harringtonia* root-tip extract and comparison with authentic synthesized standards. **(e-f)** MS/MS fragmentation spectra of **(e)** compound **1** and **(f)** compound **11** in root-tip extract and comparison with synthesized standards at a collision energy of 20V, with putative structures for ion fragments for each compound. Experimental and theoretical fragment ions support the proposed structures. **(g)** No enrichment of **1**-$D_4$ ($[M + H]^+$ = *m/z* 290.1689) observed anywhere in different tissues of *C. harringtonia* from feeding dopamine-$D_4$. **(h)** LC-MS chromatograms (EICs) of different tissues of *C. harringtonia* fed with dopamine-$D_4$ showing no detection of **1**-$D_4$. Similarly, no compound **11**-$D_4$ was detected in different tissues of *C. harringtonia* fed with dopamine-$D_4$ (n.d., not detected). **(i-k)** LC-MS chromatograms (EICs) of different tissues of *C. harringtonia* fed with **1**-$D_4$ showing no enrichment of **(i)** CET-$D_4$, **(j)** **2**-$D_4$, and **(k)** **3-**$D_4$. At least 3 biological replicates of the study were conducted, with similar results observed. The negative results from feeding **1**-$D_4$ suggest that compound **1** does not serve as an intermediate in CET biosynthesis or that the negative results could be attributed to the failure of the feeding assay. **(l)** MS/MS fragmentation spectrum of **1**-$D_4$ synthesized standard (violet traces) in comparison to that of **1** synthesized standard (green traces) at a collision energy of 20V, with putative structures for ion fragments for each compound. Experimental and theoretical fragment ions support the proposed structures. All the experiments were conducted in May 2023. All the bar graphs are reported as mean ± SD (*n* = 4 biological replicates for YSt, ON, and Rt, and *n* = 2 for YIN, YON, and OSt). Young inner needles (YIN), young outer needles (YON), Young stems (YSt), old needles (ON), old stems (OSt), and root tips (Rt).

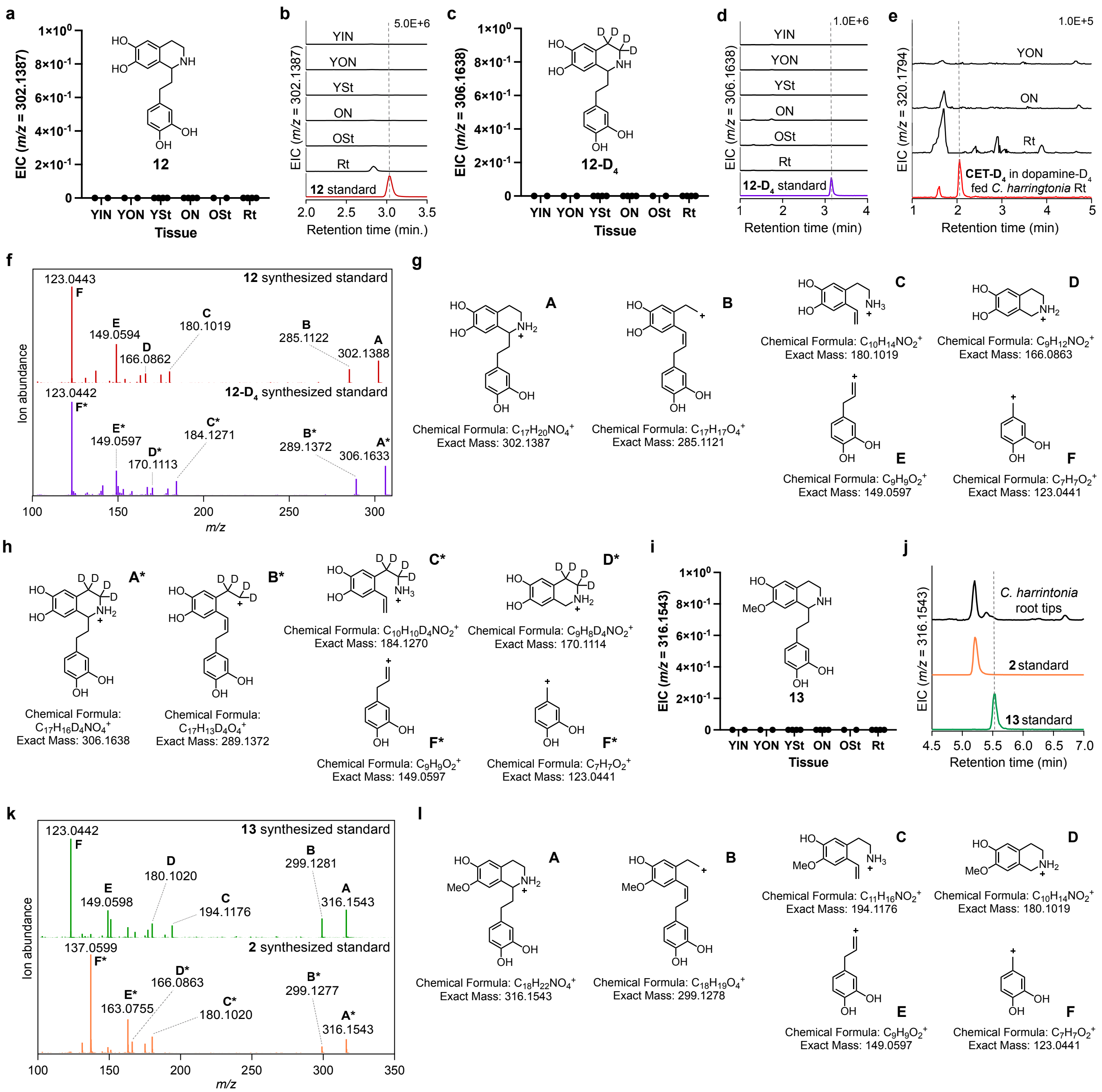


**Supplementary Figure 5. No detection of other phenethylisoquinolines 12 and 13 in *C. harringtonia*. (a-b)** No natural accumulation of compound **12** ($[M + H]^+$ = *m/z* 306.1387) detected in *C. harringtonia*, shown in **(a)** bar graphs of EICs and **(b)** LC-MS chromatograms (EICs). **(c-d)** No enrichment of **12**-$D_4$ ($[M + H]^+$ = *m/z* 306.1638) detected anywhere in different tissues of *C. harringtonia* from feeding dopamine-$D_4$, shown in **(c)** bar graphs of EICs and **(d)** LC-MS chromatograms (EICs). **(e)** LC-MS chromatograms (EICs) of different tissues of *C. harringtonia* fed with **12**-$D_4$ showing no enrichment of CET-$D_4$. At least 3 biological replicates of the study were conducted, with similar results observed. The negative results from feeding **12**-$D_4$ suggests that compound **12** does not serve as an intermediate in CET biosynthesis or that the negative results could be attributed to the failure of the feeding assay. But because compound **12** is not naturally observed in *C. harringtonia*, it is likely that compound **12** does not serve as an intermediate in CET biosynthesis. **(f)** MS/MS fragmentation spectrum of **12** synthesized standard (maroon traces) in comparison to that of **12**-$D_4$ synthesized standard (violet traces) at a collision energy of 20V. **(g-h)** Putative structures for ion fragments generated from MS/MS analysis of **(g)** compound **12** and **(g)** compound **12**-$D_4$. Experimental and theoretical fragment ions support the proposed structures. **(i)** No natural accumulation of compound **13** ($[M + H]^+$ = *m/z* 316.1543, r.t. = 5.6 min) observed in different tissues of *C. harringtonia*. **(j)** LC-MS chromatograms (EICs) show a peak for compound **2** ($[M + H]^+$ = *m/z* 316.1543, r.t. = 5.2 min), whereas compound **13** is not detected (n.d.) in *C. harringtonia* root tips. Similarly, no compound **13**-$D_4$ was detected in different tissues of *C. harringtonia* fed with dopamine-$D_4$. No presence of compound **13** in *C. harringtonia* suggests that **13** is not an intermediate of CET biosynthesis. **(k)** MS/MS fragmentation spectrum of **13** synthesized standard (green traces) in comparison to that of **2** synthesized standard (orange traces) at a collision energy of 20V. **(l)** Putative structures for ion fragments generated from MS/MS analysis of compound **13** and its comparison to compound **2**. Experimental and theoretical fragment ions support the proposed structures. See Supplementary Fig. 3 for MS/MS fragmentation of compound **2**. All the experiments were conducted in May 2023, except the feeding experiment of **12**-$D_4$ in **(e)** conducted in June 2024. All the bar graphs are reported as mean ± SD ($n$ = 4 biological replicates for YSt, ON, and Rt, and $n$ = 2 for YIN, YON, and OSt). Young inner needles (YIN), young outer needles (YON), Young stems (YSt), old needles (ON), old stems (OSt), and root tips (Rt).

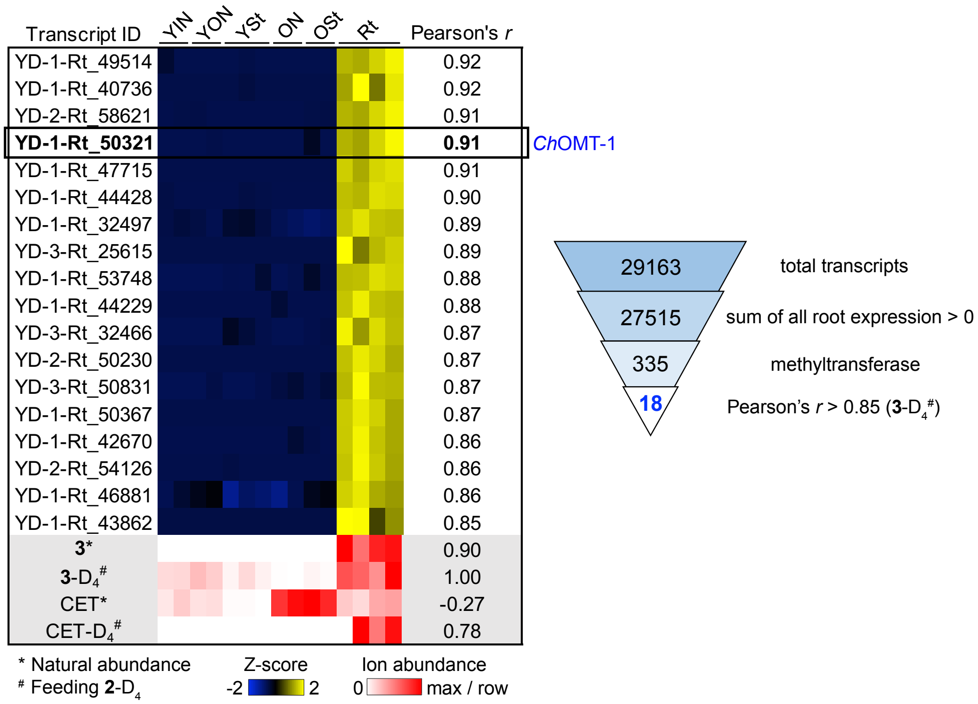


**Supplementary Figure 6. Identification of OMT responsible for O-methylation of 2 to 3.** The heatmap shows expression profiles (Z-score scaled) of the top 18 methyltransferase-annotated transcripts in 15 samples of six different tissues of *C. harringtonia* (from an independent experiment conducted in May 2023) selected based on expression in root tips (site of active biosynthesis) and strong correlation with **3**-$D_4$ enrichment ($m/z$ 334.1951) following **2**-$D_4$ feeding (Pearson's $r > 0.85$). Transcripts are ordered by decreasing correlation, with *Ch*OMT-1 (YD-1-Rt_50321) identified from the first set of five candidates cloned and functionally tested (highlighted in Fig. 3a). Below, a metabolomic heatmap (red scale) shows LC-MS ion abundances of **3** and CET at natural isotope abundance (*) and their four-deuterated analogs ($^{\#}$) after **2**-$D_4$ feeding, measured in the same *C. harringtonia* tissues used for transcriptomic analysis (each tissue was divided for parallel profiling). Right: Funnel plot showing the transcript selection process from 29,163 total transcripts to 18 candidate methyltransferases. Six different tissues (15 samples): 2 young inner needles (YIN), 2 young outer needles (YON), 3 young stems (YSt), 2 old needles (ON), 2 old stems (OSt), and 4 root tips (Rt)—harvested in early May 2023.

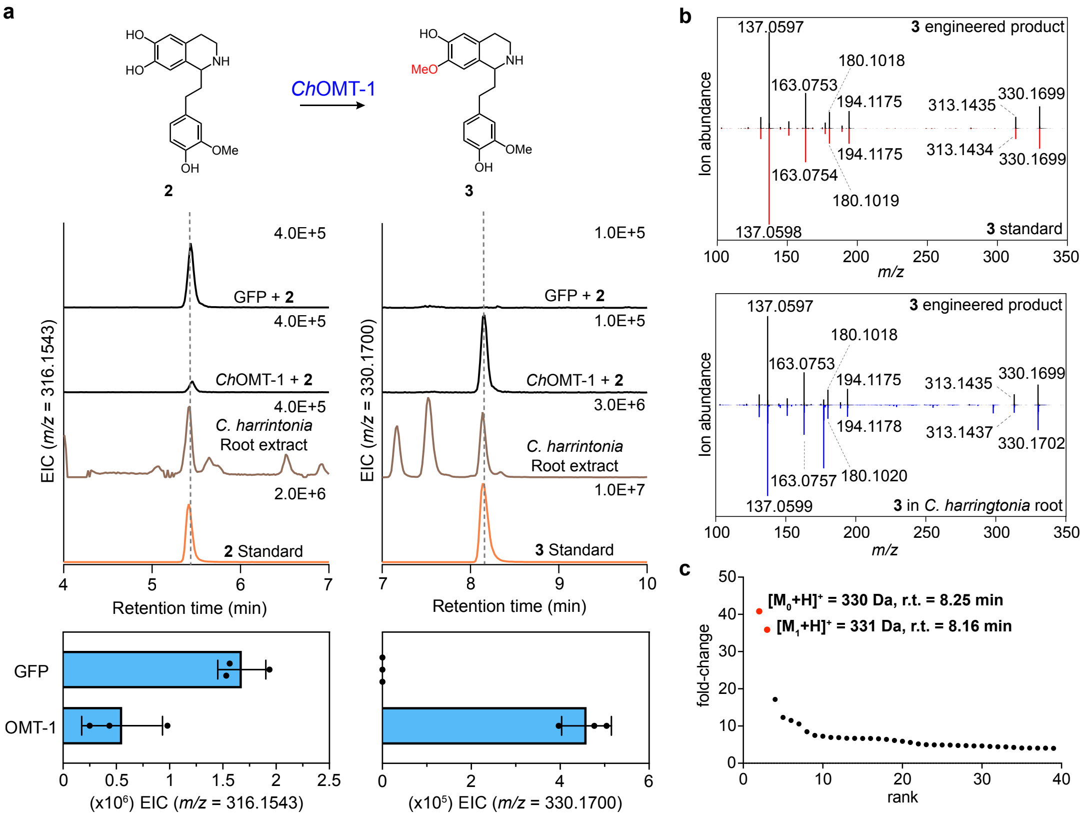


**Supplementary Figure 7. Characterization of *Ch*OMT-1. (a)** Expression of *Ch*OMT-1 in *N. benthamiana* with co-infiltrated **2** leads to consumption of **2** ($[M + H]^+$ = *m/z* 316.1543, r.t. of 5.4 min) and production of an O-methylated compound **3** ($[M + H]^+$ = *m/z* 330.1700, r.t of 8.1 min), as shown by the LC-MS chromatograms. Both compounds are true metabolites detected in *C. harringtonia* root tip extract (brown traces). Retention times of authentic standards (orange traces) are shown for comparison. Bottom shows quantification of **2** and **3** levels (mean ± SD, $n = 3$) in extracted ion abundance for the corresponding exact ion mass $[M + H]^+$. **(b)** Mirror plots compare the MS/MS spectra of enzymatically produced **3** in *N. benthamiana* with an authentic synthesized **3** standard (top) and **3** extracted from *C. harringtonia* root tips (bottom). Fragmentation patterns of the engineered **3** product (black traces) closely match those of both the standard (red traces) and the native compound (blue traces), confirming structural identity. Collision energies for all shown MS/MS analyses were 20 V. **(c)** Unique *m/z* features shown in ranked order based on their increasing fold change in ion abundance between transient expression of GFP (negative control) to that of *Ch*OMT-1 with co-infiltrated substrate **2** ($n$ = 3 independent replicates per condition), determined by untargeted metabolite analysis using XCMS ($P < 0.1$ and fold change > 5 between samples, see Methods). The mass isotopologues ($M_0$ and $M_1$) of the putative product (*m/z* 330.1700) are shown in red. The rest in black are mostly below signal to noise threshold and cannot be identified in the raw LC-MS, independent of the substrate **2**, or non-native to *C. harringtonia*. r.t., retention time.

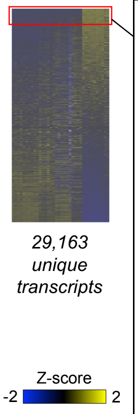


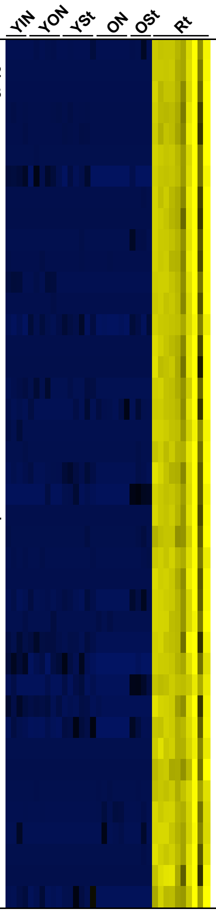


| gene | Best Pfam hit | Pearson's *r* (all tissues) | Pearson's *r* (only roots) | rank | Best BLAST hit against old data | pident |
|---|---|---|---|---|---|---|
| **OMT-1** | O-methyltransferase | 1.0000 | 1.00 | 1 | Gene.501593::Needles_204133 | 70.37 |
| **CYP805A12** | Cytochrome P450 | 0.9977 | 0.99 | 2 | Gene.389593::Needles_119791 | 60.531 |
| **CYP805A13** | Cytochrome P450 | 0.9945 | 0.99 | 3 | Gene.389593::Needles_119791 | 56.856 |
| CYP | Cytochrome P450 | 0.9862 | 0.99 | 4 | Gene.389593::Needles_119791 | 58.256 |
| **PPO-1** | Common central domain of tyrosinase | 0.9848 | 0.99 | 5 | NA | NA |
| 2OGD | 2OG-Fe(II) oxygenase superfamily | 0.9968 | 0.98 | 6 | Gene.477585::Needles_1837202 | 50.311 |
| - | Stress responsive A/B Barrel Domain | 0.9855 | 0.98 | 7 | Gene.456602::Needles_167329 | 33.478 |
| - | UDP-glucoronosyl and UDP-glucosyl transferase | 0.9901 | 0.98 | 8 | Gene.465984::Needles_174593 | 76.173 |
| - | Helix-loop-helix DNA-binding domain | 0.9842 | 0.98 | 9 | Gene.498174::Needles_201031 | 29.67 |
| - | Stress responsive A/B Barrel Domain | 0.9934 | 0.98 | 10 | Gene.456602::Needles_167329 | 34.821 |
| **CYP805A11** | Cytochrome P450 | 0.9910 | 0.98 | 11 | Gene.389593::Needles_119791 | 57.721 |
| BetV | Pathogenesis-related protein Bet v 1 family | 0.9948 | 0.97 | 12 | Gene.509023::Needles_211076 | 80.864 |
| FMN | NADH:flavin oxidoreductase / NADH oxidase family | 0.9927 | 0.97 | 13 | Gene.471099::Needles_1786122 | 81.543 |
| - | Stress responsive A/B Barrel Domain | 0.9919 | 0.97 | 14 | Gene.456602::Needles_167329 | 33.478 |
| - | Stress responsive A/B Barrel Domain | 0.9909 | 0.97 | 15 | Gene.456602::Needles_167329 | 33.624 |
| - | Stress responsive A/B Barrel Domain | 0.9803 | 0.97 | 16 | Gene.456602::Needles_167329 | 33.036 |
| ZDH | Zinc-binding dehydrogenase | 0.9925 | 0.97 | 17 | Gene.472862::Needles_179994 | 81.303 |
| - | Stress responsive A/B Barrel Domain | 0.9835 | 0.96 | 18 | Gene.456602::Needles_167329 | 34.419 |
| BetV | Pathogenesis-related protein Bet v 1 family | 0.9970 | 0.96 | 19 | Gene.509023::Needles_211076 | 77.778 |
| - | UDP-glucoronosyl and UDP-glucosyl transferase | 0.9967 | 0.96 | 20 | Gene.413024::Needles_135730 | 51.927 |
| **NmrA-1** | NmrA-like family | 0.9907 | 0.96 | 21 | Gene.449032::Needles_161618 | 88.715 |
| - | Stress responsive A/B Barrel Domain | 0.9863 | 0.96 | 22 | Gene.456602::Needles_167329 | 34.821 |
| **CYP805A14** | Cytochrome P450 | 0.9927 | 0.96 | 23 | Gene.389593::Needles_119791 | 60.944 |
| CYP | Cytochrome P450 | 0.9897 | 0.96 | 24 | Gene.390407::Needles_120342 | 75.944 |
| OMT | O-methyltransferase | 0.9975 | 0.96 | 25 | Gene.501593::Needles_204133 | 71.616 |
| CYP | Cytochrome P450 | 0.9939 | 0.96 | 26 | Gene.389593::Needles_119791 | 59.287 |
| - | Stress responsive A/B Barrel Domain | 0.9950 | 0.95 | 27 | Gene.456602::Needles_167329 | 34.821 |
| BetV | Pathogenesis-related protein Bet v 1 family | 0.9963 | 0.95 | 28 | Gene.509023::Needles_211076 | 87.654 |
| **TyDC-1** | Pyridoxal-dependent decarboxylase conserved domain | 0.9814 | 0.94 | 29 | Gene.330462::Needles_81944 | 87.001 |
| SDR | short chain dehydrogenase | 0.9812 | 0.94 | 30 | Gene.477334::Needles_183524 | 100 |
| 2OGD | 2OG-Fe(II) oxygenase superfamily | 0.9908 | 0.93 | 31 | Gene.477585::Needles_1837202 | 53.153 |
| ADH | Alcohol dehydrogenase GroES-like domain | 0.9841 | 0.93 | 32 | Gene.459106::Needles_169249 | 93.49 |
| CYP | Cytochrome P450 | 0.9862 | 0.92 | 33 | Gene.424990::Needles_144130 | 92.194 |
| - | NA | 0.9967 | 0.92 | 34 | Gene.517584::Needles_219908 | 72.897 |
| CYP | Cytochrome P450 | 0.9894 | 0.91 | 35 | Gene.390407::Needles_120342 | 75.547 |
| - | Protein kinase domain | 0.9964 | 0.91 | 36 | Gene.241866::Needles_32206 | 36.394 |
| SDR | short chain dehydrogenase | 0.9960 | 0.91 | 37 | Gene.461803::Needles_171344 | 74.359 |
| - | NA | 0.9929 | 0.91 | 38 | NA | NA |
| **2OGD-1** | 2OG-Fe(II) oxygenase superfamily | 0.9947 | 0.91 | 39 | Gene.486674::Needles_191161 | 55.247 |
| CYP | Cytochrome P450 | 0.9826 | 0.91 | 40 | Gene.389593::Needles_119791 | 69.799 |
| FAD | FAD binding domain | 0.9836 | 0.90 | 41 | Gene.425978::Needles_144829 | 50.407 |

**Supplementary Figure 8. Co-expression analysis identifies candidate genes involved in CET biosynthesis, captured only in the new in-house transcriptome.** RNA-seq analyses across various tissue types—young inner needles (YIN), young outer needles (YON), young stems (YSt), old needles (ON), old stems (OSt), and root tips (Rt)—combined with Pearson's correlation using *Ch*OMT-1 as the bait gene, identify a set of strongly co-expressing biosynthetic gene candidates from 29,163 unique transcripts. Initial Pearson correlation analysis across all tissues using *Ch*OMT-1 as a query gene identified 316 transcripts ($r > 0.98$). As many of these showed expression only in root tips, a second round of correlation analysis was conducted using Rt samples alone, yielding 41 top candidates ($r > 0.90$) as shown. Genes involved in CET biosynthesis are highlighted in bold. Expression values within the heat map are $log_2$-transformed values of each transcript calculated as Trimmed mean of M (TMM)-normalized counts per million (CPM) in Z-score scale. "Best BLAST hit against old data" refers to matches against an earlier in-house transcriptome generated from tissues harvested in August 2018, whereas the transcriptome used for all analyses in this study was assembled from tissues harvested in mid-May 2023. "Pident" indicates percent identity of protein sequences between the new and old transcriptomes. Notably, CET biosynthetic genes are absent from the old August 2018 dataset, suggesting seasonal regulation of pathway expression. The whole heatmap is composed of 36 unique samples—YIN (4), YON (6), YSt (6), ON (6), OSt (4), and Rt (10)—harvested from the same plant but at two different times (one set harvested in early May 2023 and the other two weeks after).

**Note regarding transcriptome data:** CET biosynthetic genes found in the new May 2023 transcriptome dataset are absent from the old August 2018 dataset, suggesting seasonal regulation of pathway expression.

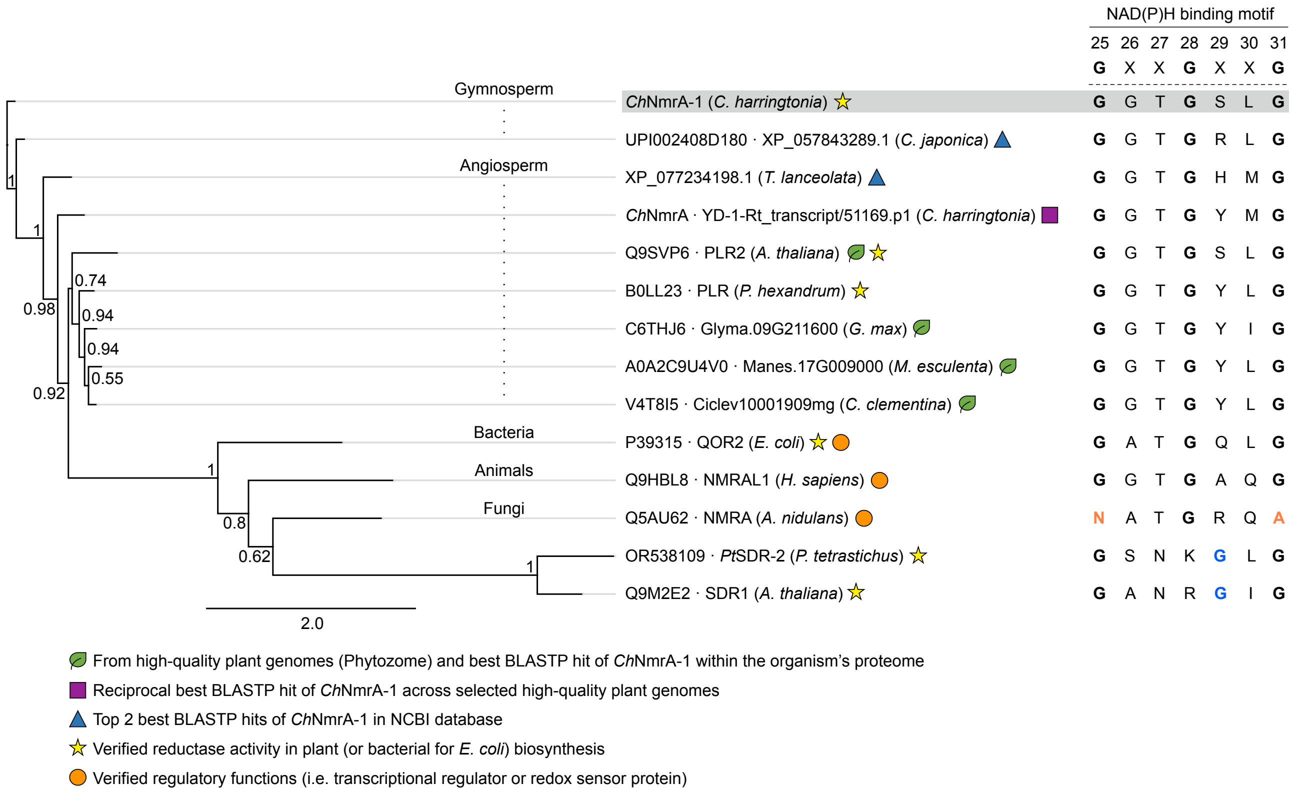


**Supplementary Figure 9. Phylogenetic tree and sequence analyses of *Ch*NmrA-1 and other NmrA-like family (atypical short-chain dehydrogenases, SDRs) and classical SDRs.** A phylogenetic tree (MAFFT alignment[1], FastTree[2]) of NmrA-like family (PF05368) from multiple kingdoms of life (bacteria, animals, fungi, and mainly plants, including angiosperm and gymnosperm lineages) and classical SDRs (PF00106) is shown. Proteins from high-quality plant genomes (Phytozome) and the best BLASTP hit of *Ch*NmrA-1 within each organism's proteome are marked with green leaf symbols; reciprocal best BLASTP hit across selected high-quality plant genomes is indicated by a purple square; the top two BLASTP hits in the NCBI database are indicated by blue triangles (first *C. japonica*, a gymnosperm, and second *T. lanceolata*, an angiosperm). Proteins with verified reductase activity in plant (or bacterial for *E. coli*) biosynthesis are marked with yellow stars, including the pinoresinol-lariciresinol reductases (PLR) in lignan biosynthesis, whereas proteins with verified regulatory functions (i.e. transcriptional regulators or redox sensor proteins) are indicated by orange circles. Proteins shown in the tree, except the last two classical SDRs (PF00106) at the bottom, belong to the NmrA-like family (PF05368). Node values indicate Shimodaira-Hasegawa (SH)-like local support values calculated by FastTree[2], and branch lengths (black lines) are drawn to scale (grey lines, for alignment). Also shown on the right is the conserved Rossmann-like $NAD(P)^+$/NAD(P)H-binding motif (GXXGXXG) for each aligned protein, with numbering corresponding to *Ch*NmrA-1 (highlighted in grey) as it represents the focal NmrA-like family protein investigated in this study. This highlights conservation of the glycine-rich cofactor-binding loop in plant homologs. Classical SDRs, including SDR1 from *A. thaliana* (UniProt ID: Q9M2E2), are included as canonical enzymatic representatives to highlight conservation of the Rossmann-like NAD(P)H-binding glycine-rich motif, which appears as a GXXXGXG sequence in classical SDRs (with second glycine highlighted in blue) and a GXXGXXG variant in NmrA-like family proteins[3,4]. NmrA from *A. nidulans* has the first and last glycine residues in the motif replaced with Asn and Ala respectively (highlighted in orange)[3].

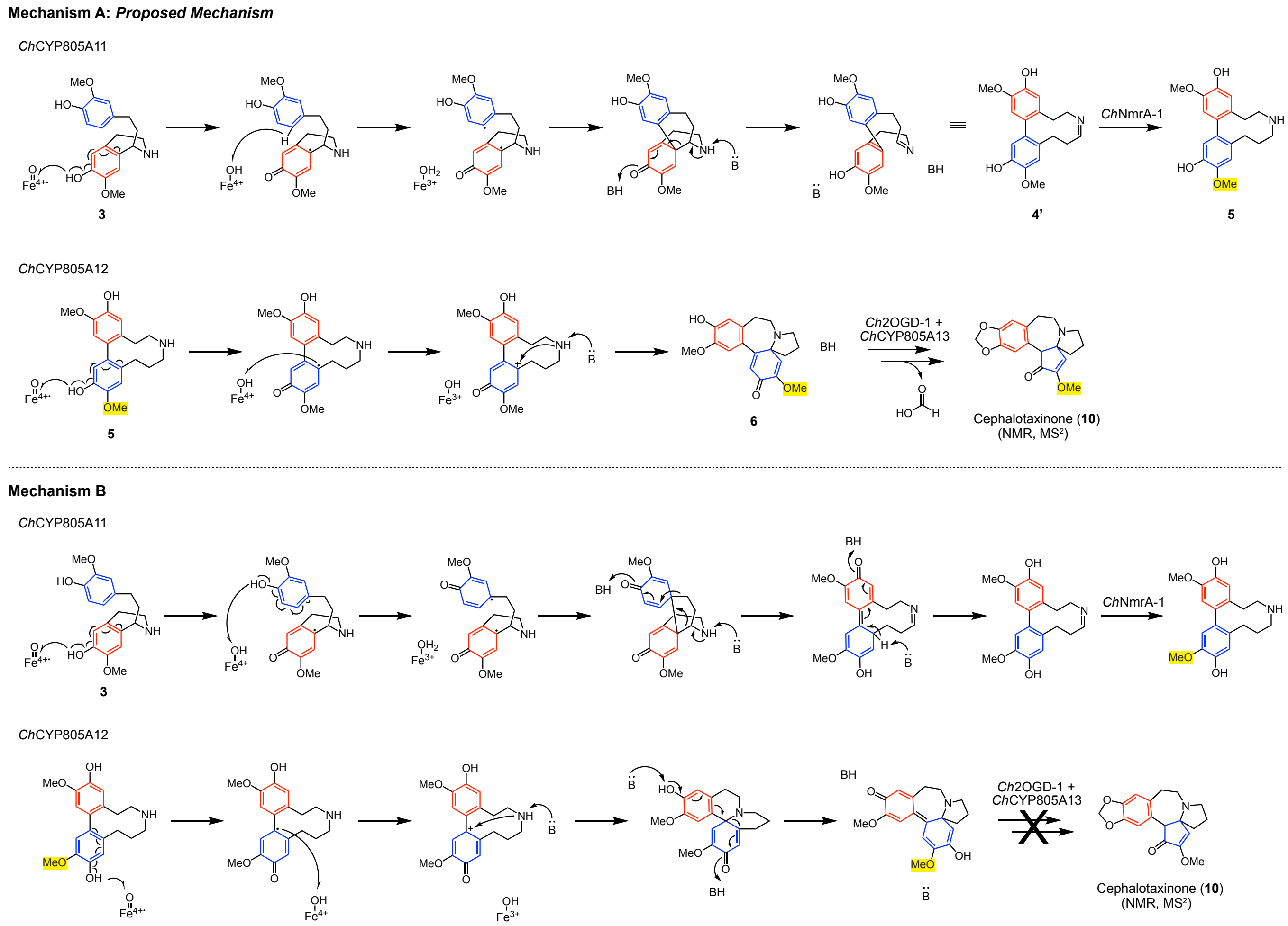


**Supplementary Figure 10. Two proposed catalytic mechanisms for *Ch*CYP805A11 and *Ch*CYP805A12.** Mechanism A involves *Ch*CYP805A11 mediated para-meta oxidative coupling of two methoxyphenols of **3** to form **4'**, which is subsequently reduced by *Ch*NmrA-1 to produce **5**. Oxidation of **5** to **6** by *Ch*CYP805A12, followed by catalysis by *Ch*2OGD-1 and *Ch*CYP805A13, enables formation of cephalotaxinone. In contrast, Mechanism B involves *Ch*CYP805A11 mediated para-para oxidative coupling of two methoxyphenols of **3** to generate a structural isomer of **4'**, which is reduced by *Ch*NmrA-1 to yield a structural isomer of **5**, differing only in the position of an O-methyl group. Oxidation of this isomer of **5** by *Ch*CYP805A12 does not lead to formation of **6** and the final product cephalotaxinone. As shown, the position of the O-methyl group (highlighted in yellow) is critical for cephalotaxinone biosynthesis. Only Mechanism A appears to be productive, supporting Mechanism A as the more likely pathway for *Ch*CYP805A11 and *Ch*CYP805A12.

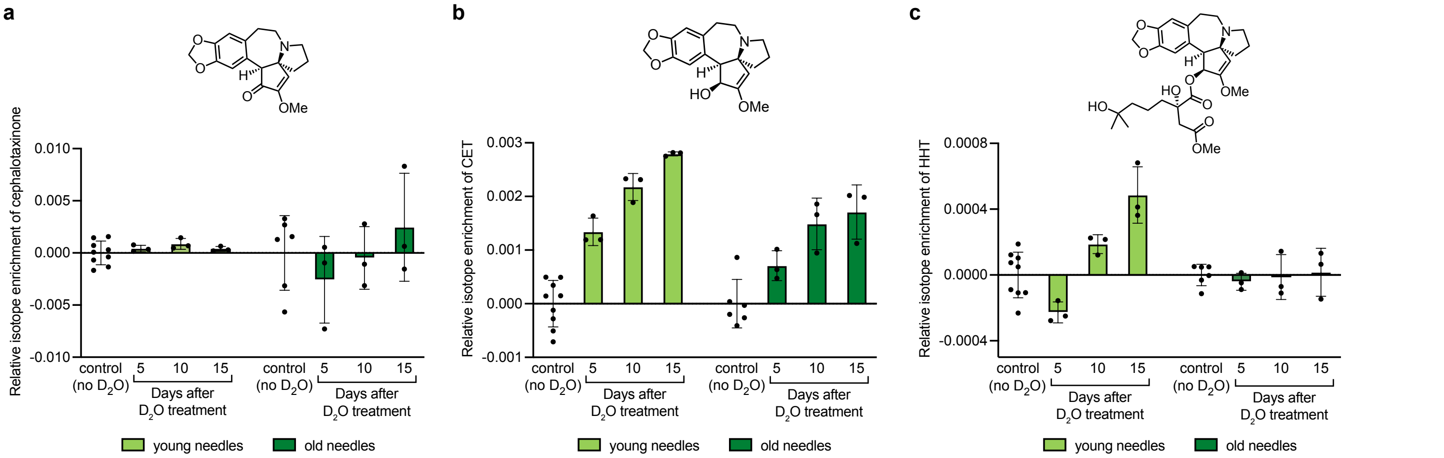


**Supplementary Figure 11. Analysis of biosynthetic location of *Cephalotaxus* alkaloids using $D_2O$ labeling. (a)** Relative isotope enrichment of cephalotaxinone in young and old needles over time. No detectable deuterated cephalotaxinone is observed in either tissue type. **(b)** Relative isotope enrichment of CET in young and old needles over time. Increase of deuterated CET is observed over time across excised branch tissues, including both young and old needles, supporting active reduction of cephalotaxinone to CET in aerial tissues. **(c)** Relative isotope enrichment of HHT over time in young and old needles. Deuterated HHT is detected at days 10 and 15 after $D_2O$ treatment with increase over time, indicating that the side-chain biosynthesis leading to HHT occurs in aerial part of the plant, especially young needles. For each metabolite, data are reported as mean ± SD for each time point ($n$ = 3 biological replicates for $D_2O$-treated samples, $n$ = 9 biological replicates for young needle controls (3 replicates each at 5, 10, and 15 days), and $n$ = 6 biological replicates for old needle controls (2 replicates each at 5, 10, and 15 days)). Young needles are shown in light green, while old needles in dark green.

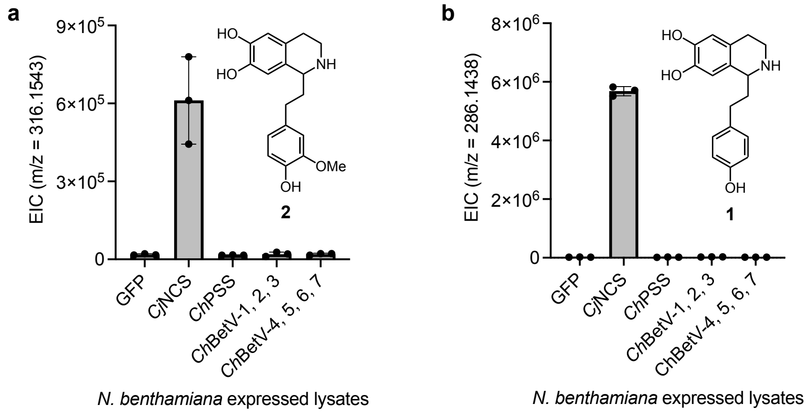


**Supplementary Figure 12. Functional screening of candidate *C. harringtonia* Pictet-Spenglerases using *N. benthamiana*-expressed lysates.** The candidate *C. harringtonia* Pictet-Spenglerases were selected from the Bet v 1 protein family (PF00407) because *Cj*NCS (a norcoclaurine synthase from *C. japonica* known to catalyze condensation of 4-hydroxyphenylacetaldehyde (4HPAA) with dopamine in benzylisoquinoline alkaloid biosynthesis[5] and reported to condense dopamine with 3-(4-hydroxyphenyl)propanal to form **1**[6]) and *Ch*PSS (a Pictet-Spenglerase from *C. hainanensis* reported to catalyze condensation of dopamine with 3-(4-hydroxyphenyl)propanal to form compound **1** in heterologous expression in *E. coli*[7], corresponding to YD-1-Rt_transcript/53915.p1 in our transcriptome with two aa difference) are members of this family. Candidates were further prioritized based on high co-expression with *Ch*OMT-1 across all tissues (Pearson's $r > 0.90$). This resulted in top seven candidates, including *Ch*BetV-1, -2, and -3 that also appear in the top candidate gene list containing the six CET biosynthetic enzymes (see Supplementary Fig. 8), which were transiently expressed in *N. benthamiana*. Crude leaf lysates were incubated with dopamine and the corresponding aldehyde substrates, and reaction products were analyzed by LC-MS. GFP-expressing *N. benthamiana* crude protein lysates are included as a negative control. The functionality of *N. benthamiana* crude leaf protein lysates expressing *Cj*NCS (GenBank accession: AB267399.2, previously cloned in our laboratory[6]) or *Ch*PSS (YD-1-Rt_transcript/53915.p1) was also each examined. **(a)** *In vitro* assay of Pictet-Spengler condensation of dopamine with 3-(4-hydroxy-3-methoxyphenyl)propanal to form compound **2**. **(b)** *In vitro* Pictet-Spengler condensation of dopamine with 3-(4-hydroxyphenyl)propanal to form compound **1**. The data are reported as mean ± SD of the extracted ion abundance ($n = 3$) for the exact ion mass $[M + H]^+$ corresponding to each compound. The *C. harringtonia* candidate enzymes were co-expressed for batch testing. None of the tested *C. harringtonia* candidates exhibit detectable Pictet-Spenglerase activity under the conditions tested (see Methods). *Ch*BetV-1: YD-1-Rt_transcript/53876.p1; *Ch*BetV-2: YD-3-Rt_transcript/61329.p1; *Ch*BetV-3: YD-1-Rt_transcript/54093.p1; *Ch*BetV-4: YD-2-Rt_transcript/59141.p1; *Ch*BetV-5: YD-1-Rt_transcript/53150.p1; *Ch*BetV-6: YD-2-Rt_transcript/59530.p1; *Ch*BetV-7: YD-1-Rt_transcript/54278.p1.

**Note:** *Cj*NCS-expressing *N. benthamiana* leaf lysate was found to catalyze both the formation of **1** and **2** through condensation of dopamine with 3-(4-hydroxyphenyl)propanal and 3-(4-hydroxy-3-methoxyphenyl)propanal respectively. In contrast, *Ch*PSS-expressing *N. benthamiana* leaf lysate did not result in condensation of both dopamine with 3-(4-hydroxyphenyl)propanal and 3-(4-hydroxy-3-methoxyphenyl)propanal to form **1** and **2** in a rate above background, respectively. None of the tested seven *C. harringtonia* candidate enzymes catalyzed a Pictet-Spengler reaction to produce **1** or **2**. The trace amounts of both **1** and **2** in all samples are most likely a result of non-enzymatic condensation reaction.

#### Supplementary Figures – NMR spectra

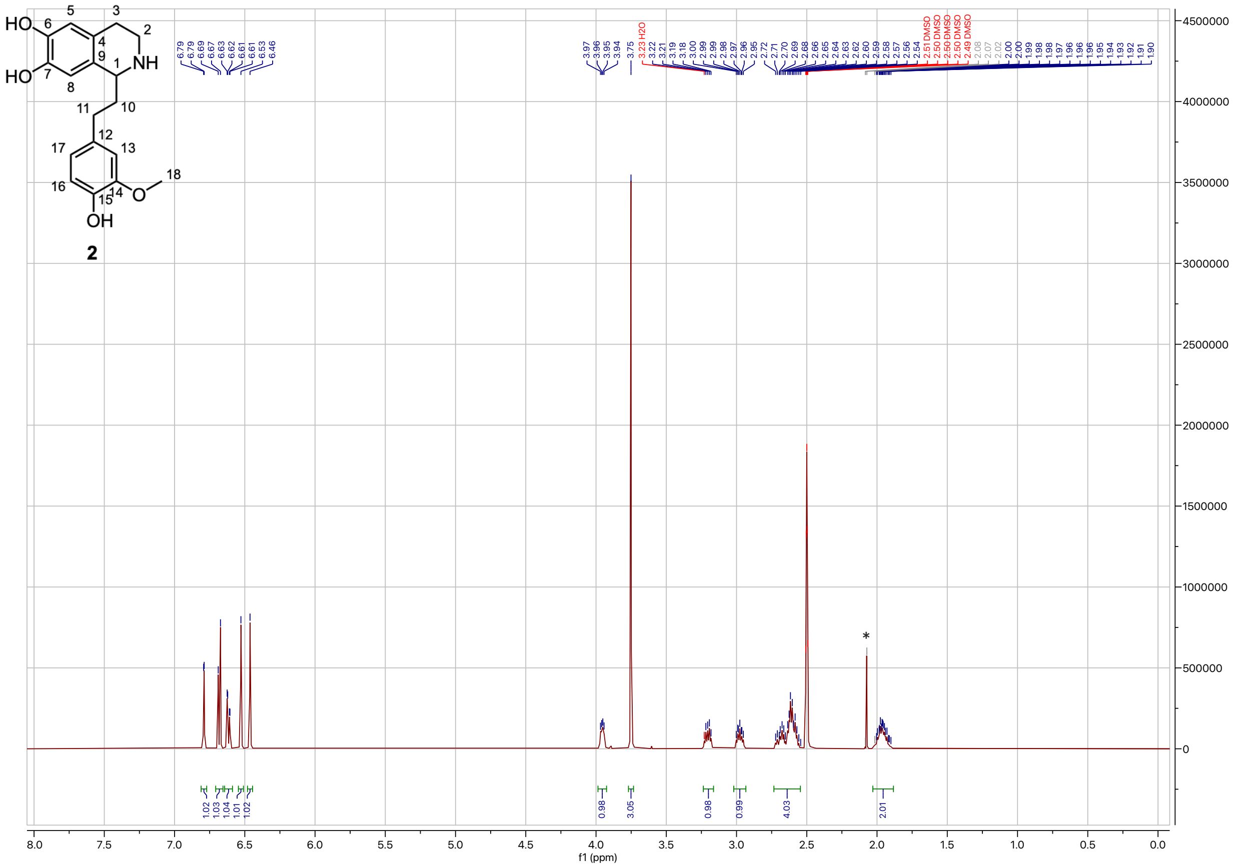

**Supplementary Figure. 13. $^1$H-NMR spectrum of 1-(4-hydroxy-3-methoxyphenethyl)-1,2,3,4-tetrahydroisoquinoline-6,7-diol (2) (DMSO-d$_6$, 500 MHz, 298 K).** $^1$H NMR (500 MHz, DMSO) δ 6.79 (d, $J$ = 1.9 Hz, 1H), 6.68 (d, $J$ = 8.0 Hz, 1H), 6.62 (dd, $J$ = 8.0, 1.9 Hz, 1H), 6.53 (s, 1H), 6.46 (s, 1H), 3.96 (dd, $J$ = 8.3, 4.2 Hz, 1H), 3.75 (s, 3H), 3.24 – 3.16 (m, 1H), 2.98 (ddd, $J$ = 12.5, 7.4, 5.2 Hz, 1H), 2.73 – 2.55 (m, 4H), 2.03 – 1.88 (m, 2H). Asterisk at 2.08 ppm corresponds to acetone that was used to clean the spatula for weighing the compound.

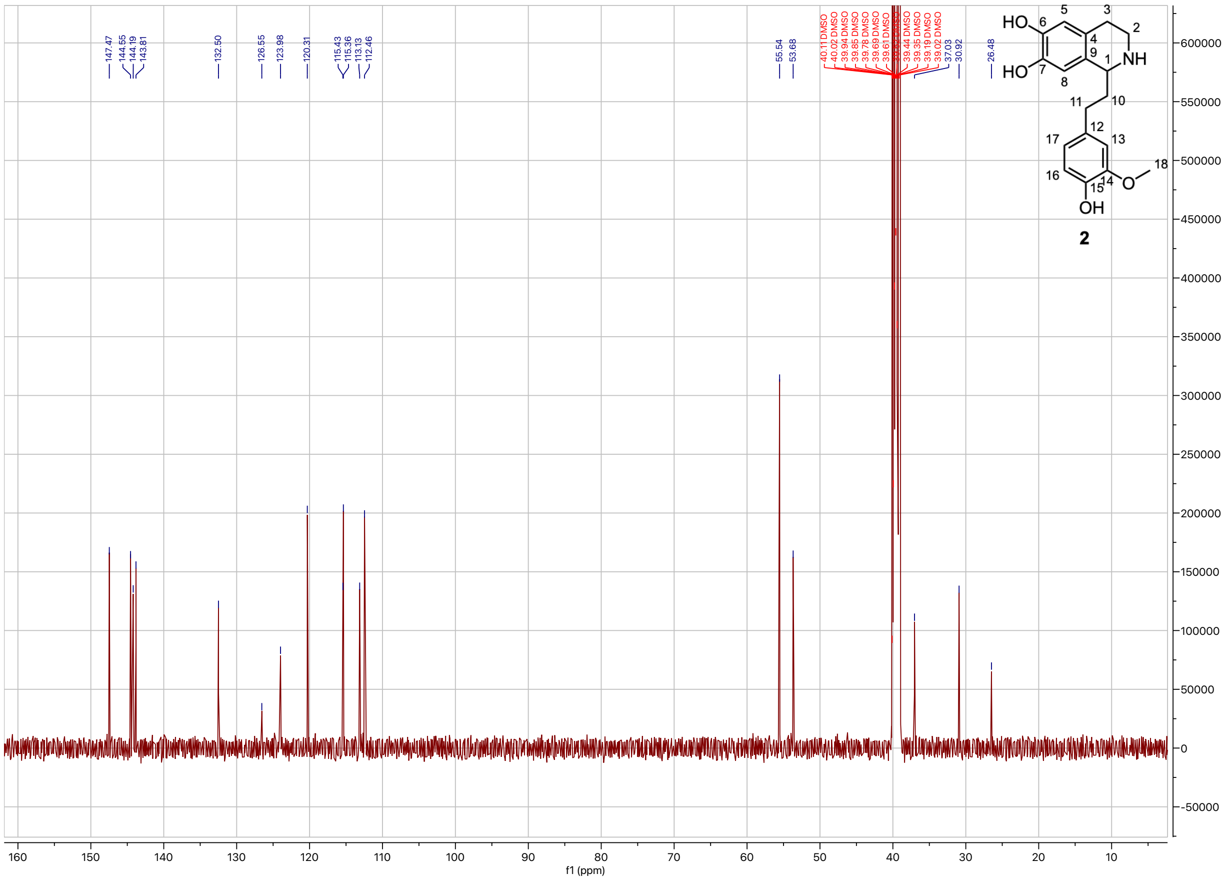


**Supplementary Figure. 14. $^{13}$C-NMR spectrum of 1-(4-hydroxy-3-methoxyphenethyl)-1,2,3,4-tetrahydroisoquinoline-6,7-diol (2) (DMSO-d$_6$, 500 MHz, 298 K).**

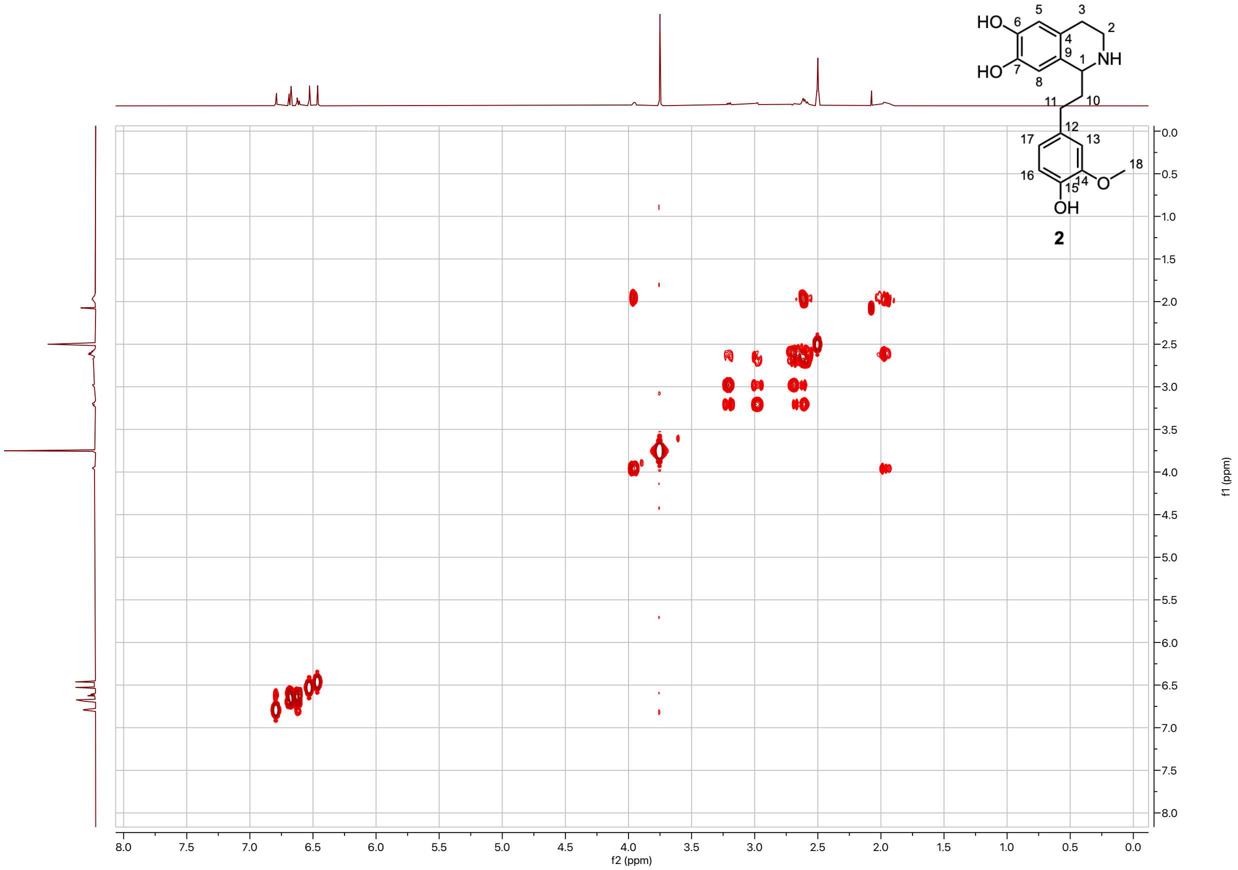


**Supplementary Figure. 15. COSY spectrum of 1-(4-hydroxy-3-methoxyphenethyl)-1,2,3,4-tetrahydroisoquinoline-6,7-diol (2) (DMSO-$d_6$, 500 MHz, 298 K).**

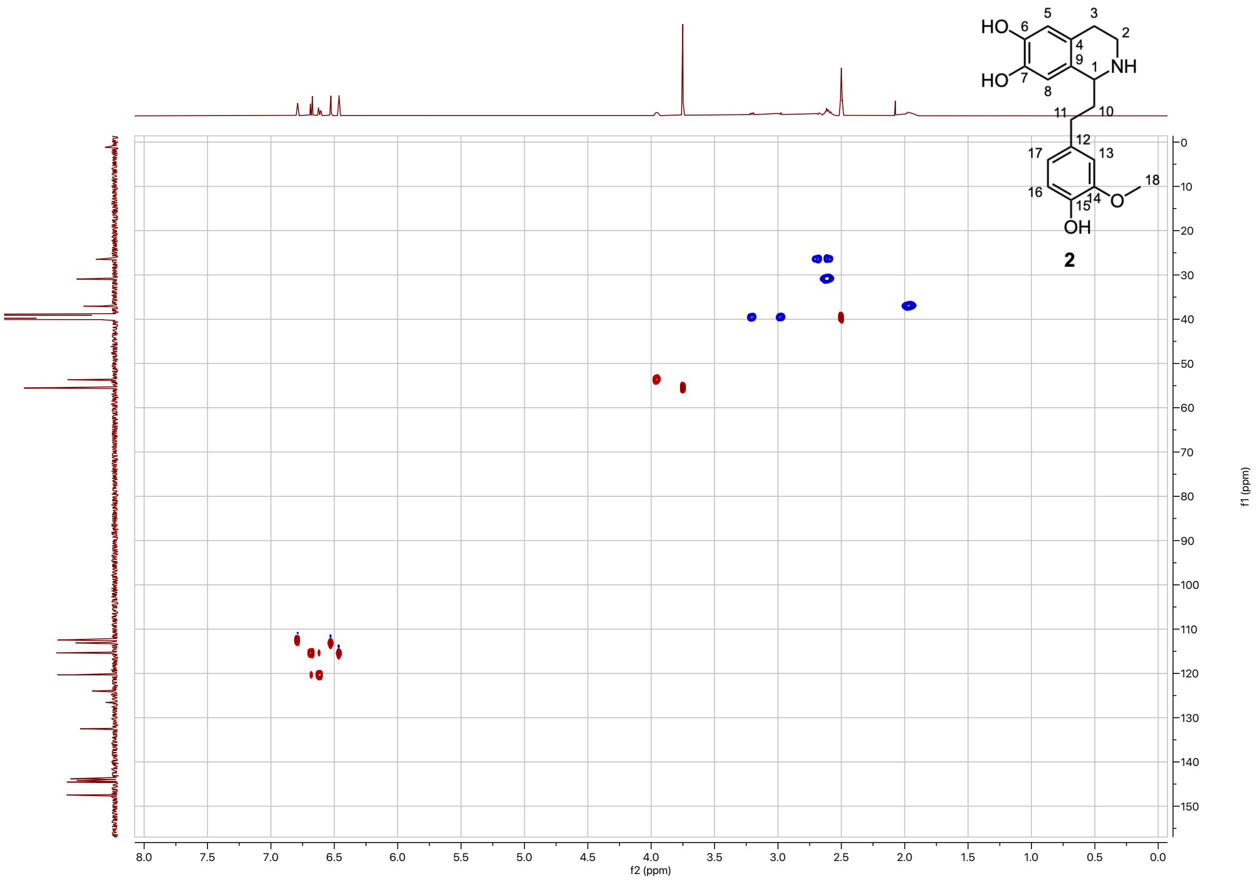


**Supplementary Figure. 16. HSQC spectrum of 1-(4-hydroxy-3-methoxyphenethyl)-1,2,3,4-tetrahydroisoquinoline-6,7-diol (2) (DMSO-d$_6$, 500 MHz, 298 K).**

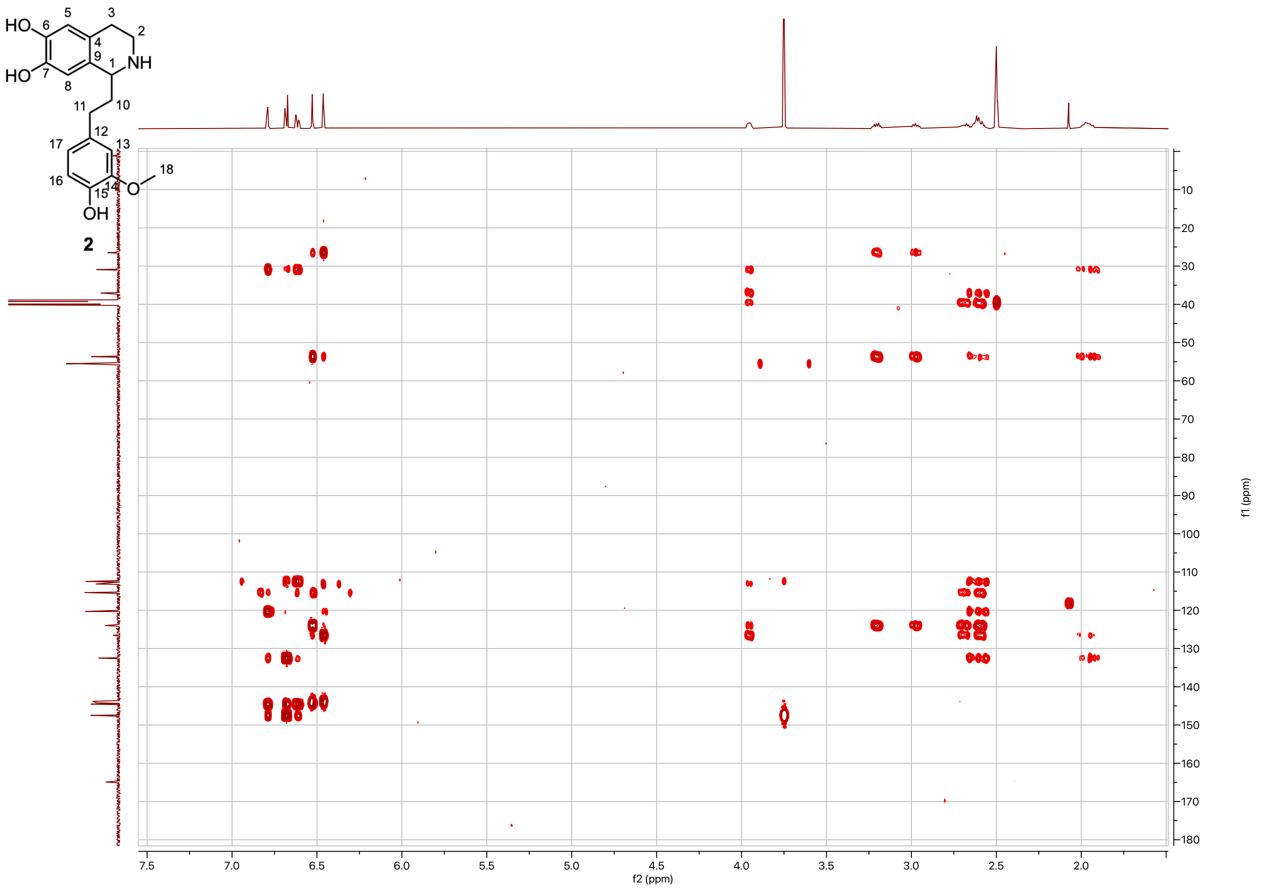


**Supplementary Figure. 17. HMBC spectrum of 1-(4-hydroxy-3-methoxyphenethyl)-1,2,3,4-tetrahydroisoquinoline-6,7-diol (2) (DMSO-$d_6$, 500 MHz, 298 K).**

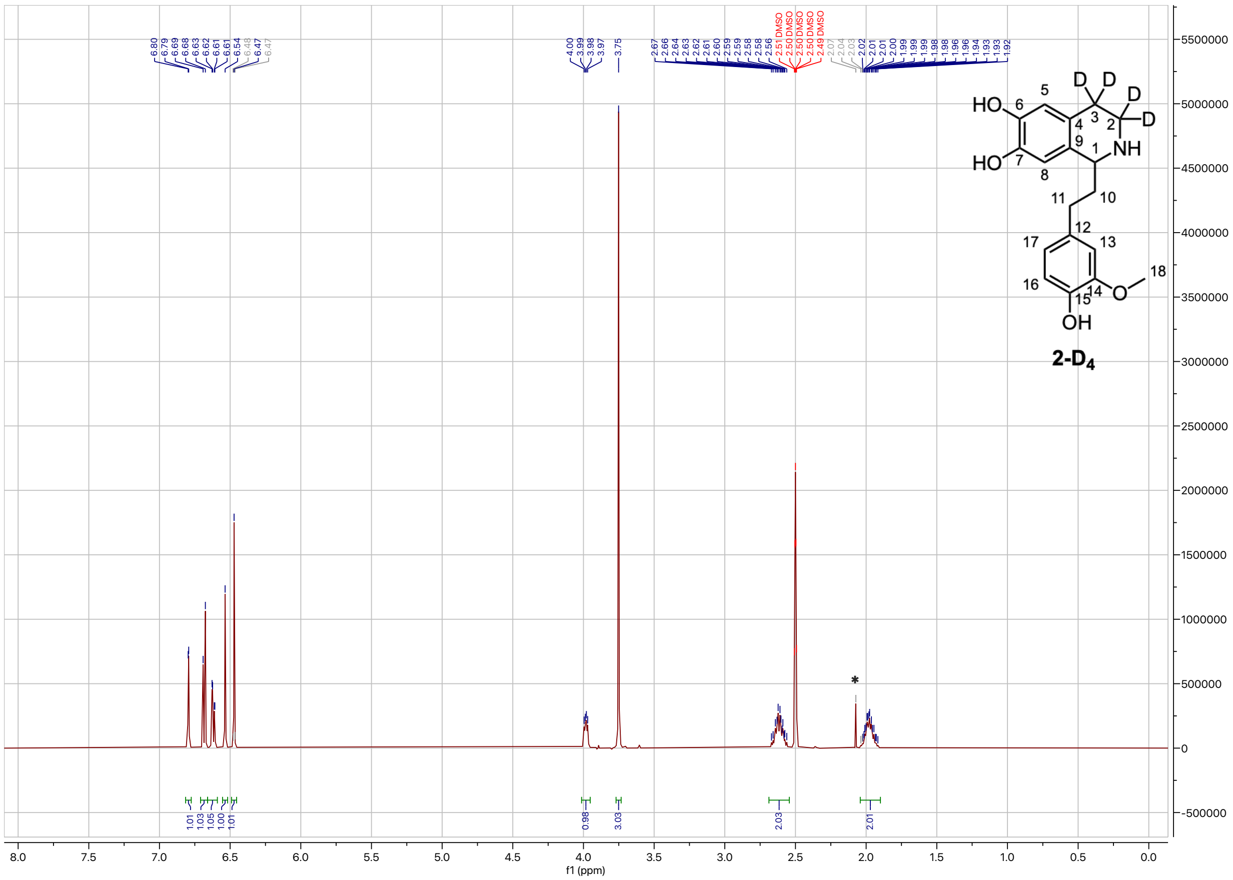

**Supplementary Figure. 18. [1]H-NMR spectrum of 1-(4-hydroxy-3-methoxyphenethyl)-1,2,3,4-tetrahydroisoquinoline-3,3,4,4-$d_4$-6,7-diol (2-D₄) (DMSO-d₆, 500 MHz, 298 K).** $^1$H NMR (500 MHz, DMSO) δ 6.79 (d, $J$ = 1.9 Hz, 1H), 6.68 (d, $J$ = 8.0 Hz, 1H), 6.62 (dd, $J$ = 7.9, 2.0 Hz, 1H), 6.54 (s, 1H), 6.47 (s, 1H), 3.98 (dd, $J$ = 8.0, 4.5 Hz, 1H), 3.75 (s, 3H), 2.68 – 2.55 (m, 2H), 2.04 – 1.90 (m, 2H). Asterisk at 2.07 ppm corresponds to acetone that was used to clean the spatula for weighing the compound.

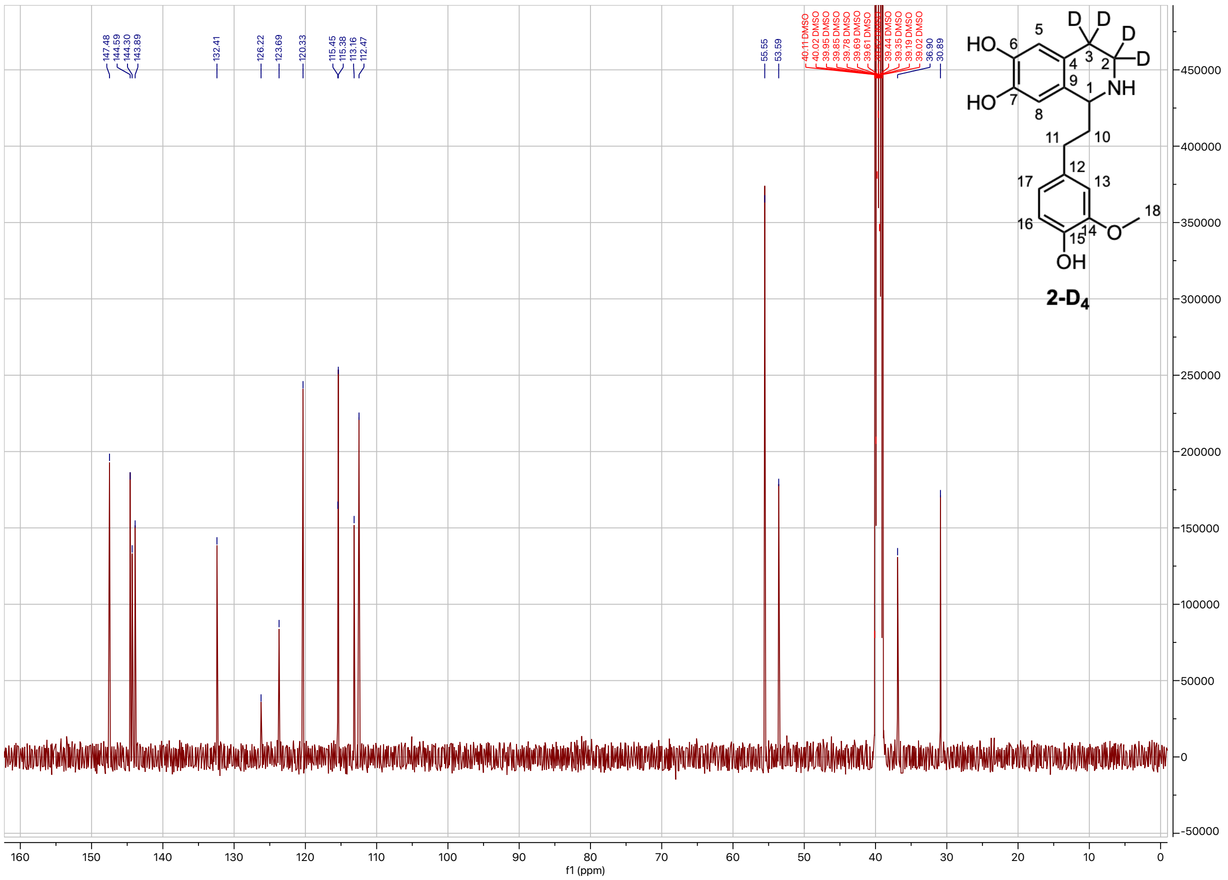

**Supplementary Figure. 19. $^{13}$C-NMR spectrum of 1-(4-hydroxy-3-methoxyphenethyl)-1,2,3,4-tetrahydroisoquinoline-3,3,4,4-$d_4$-6,7-diol (2-D$_4$) (DMSO-d$_6$, 500 MHz, 298 K).**

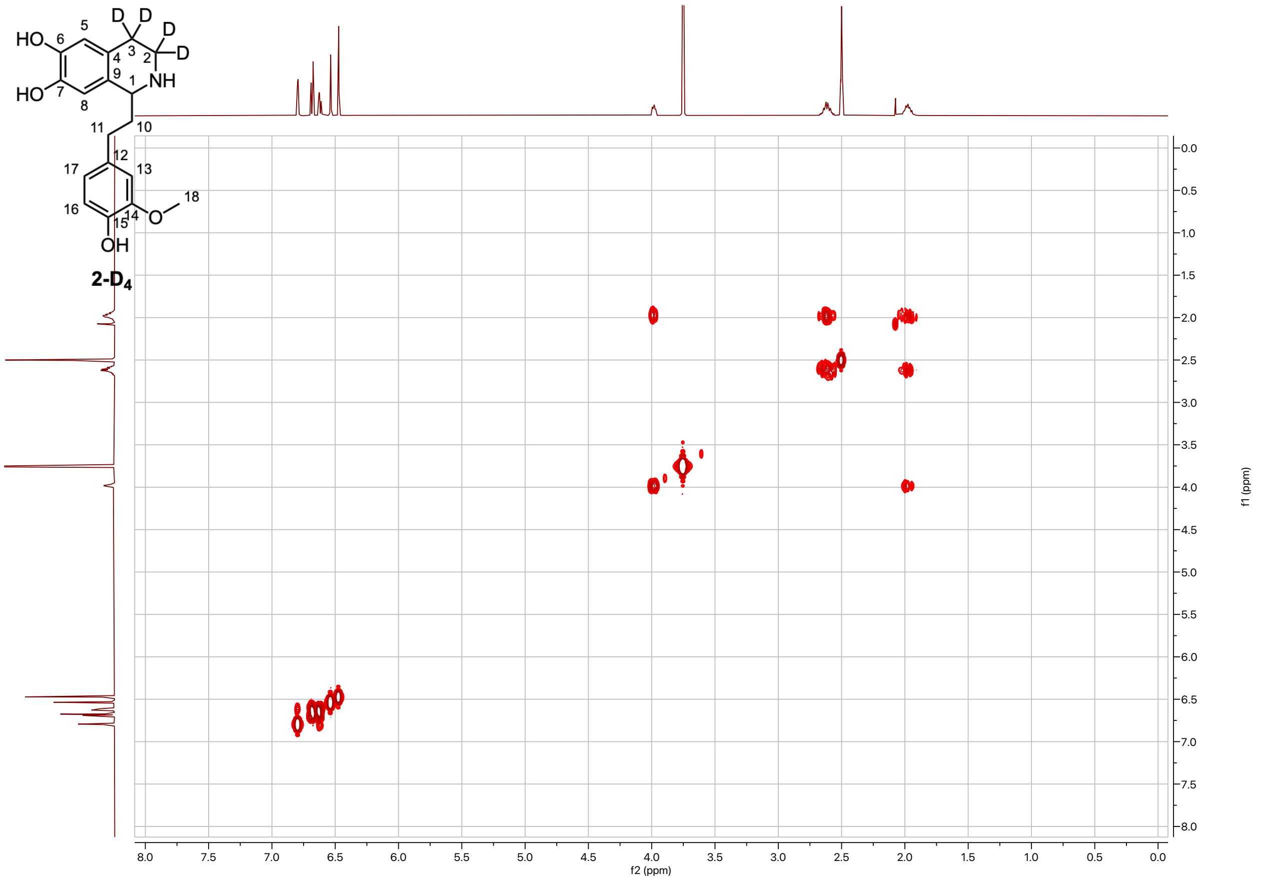


**Supplementary Figure. 20. COSY spectrum of 1-(4-hydroxy-3-methoxyphenethyl)-1,2,3,4-tetrahydroisoquinoline-3,3,4,4-$d_4$-6,7-diol (2-D₄) (DMSO-d₆, 500 MHz, 298 K).**

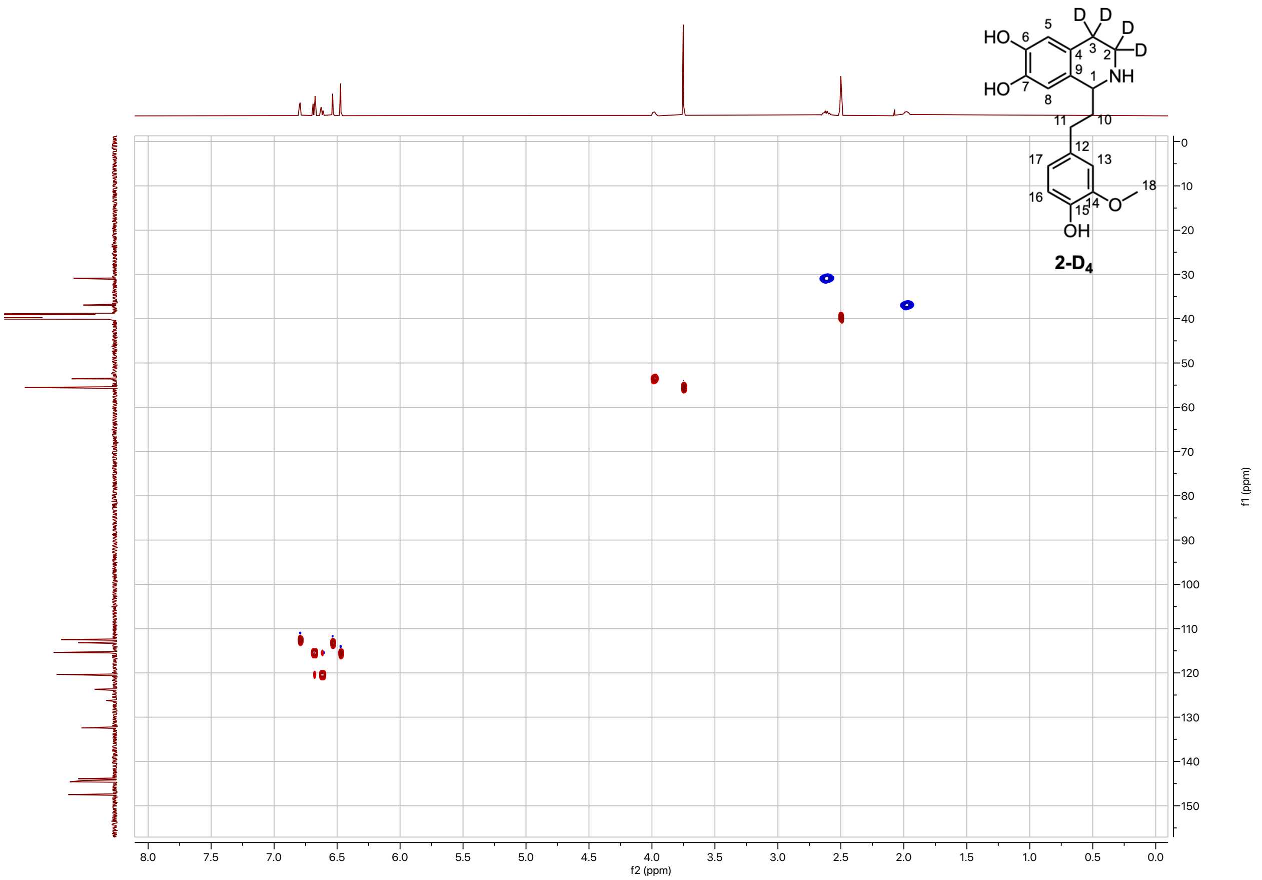


**Supplementary Figure. 21. HSQC spectrum of 1-(4-hydroxy-3-methoxyphenethyl)-1,2,3,4-tetrahydroisoquinoline-3,3,4,4-*d*$_4$-6,7-diol (2-D$_4$) (DMSO-d$_6$, 500 MHz, 298 K).**

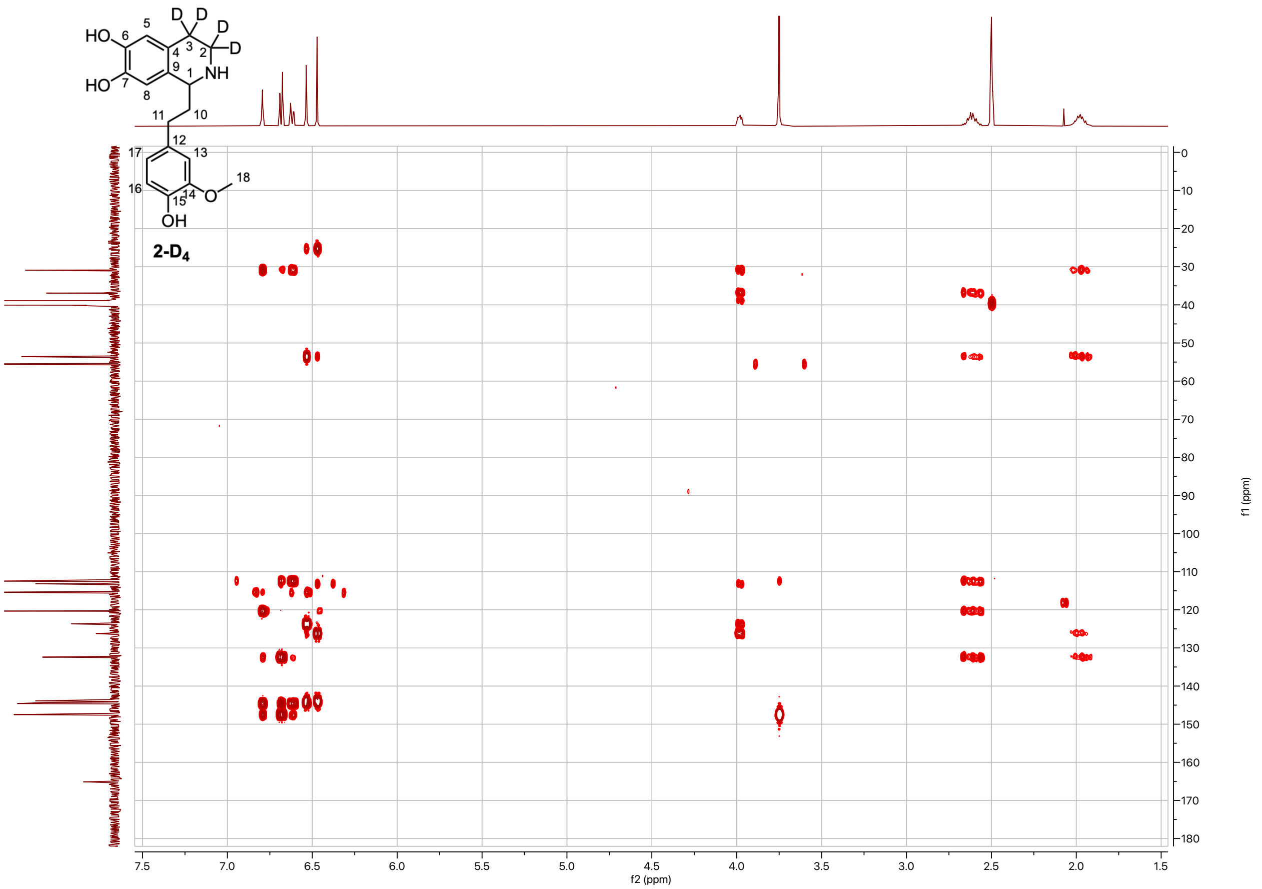


**Supplementary Figure. 22. HMBC spectrum of 1-(4-hydroxy-3-methoxyphenethyl)-1,2,3,4-tetrahydroisoquinoline-3,3,4,4-*d*$_4$-6,7-diol (2-D$_4$) (DMSO-d$_6$, 500 MHz, 298 K).**

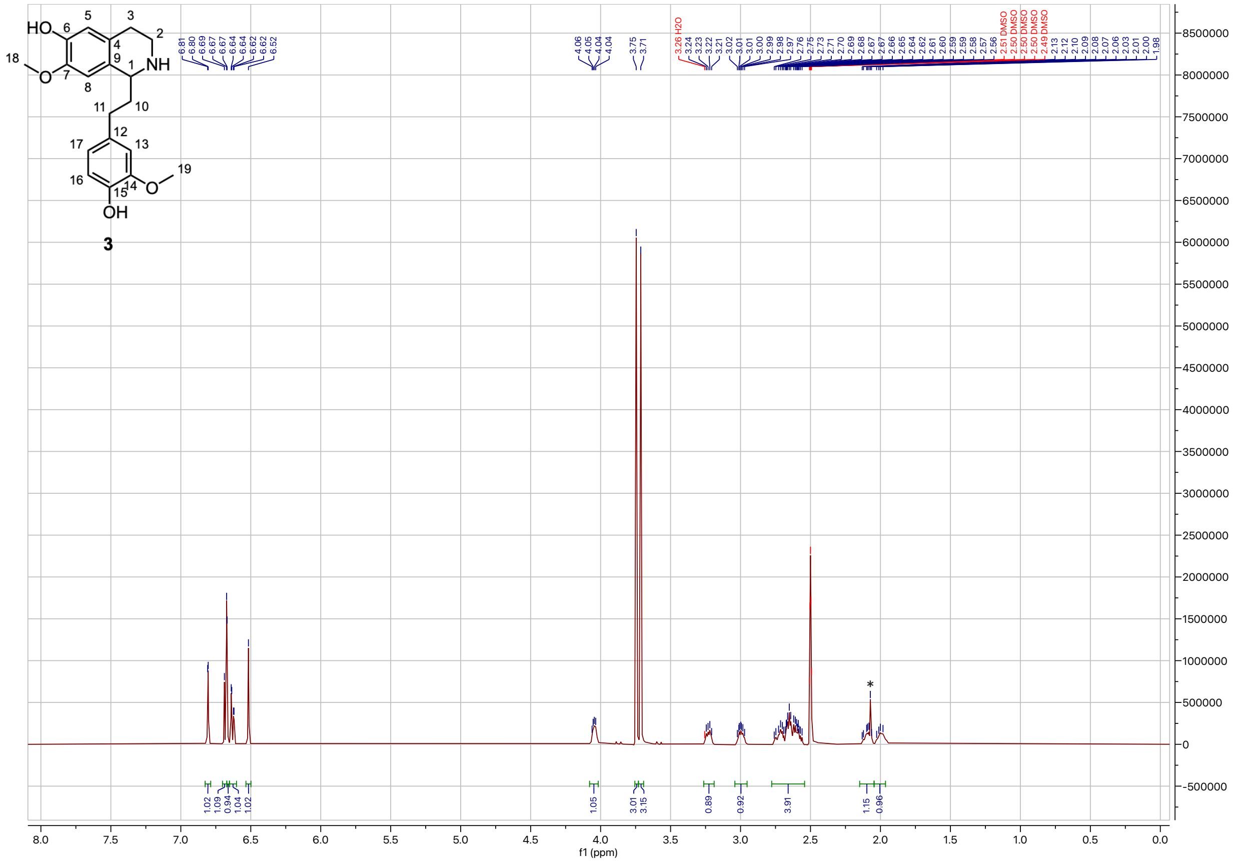

**Supplementary Figure. 23. ¹H-NMR spectrum of 1-(4-hydroxy-3-methoxyphenethyl)-7-methoxy-1,2,3,4-tetrahydroisoquinolin-6-ol (3) (DMSO-$d_6$, 500 MHz, 298 K).** $^1$H NMR (500 MHz, DMSO) δ 6.81 (d, $J$ = 1.9 Hz, 1H), 6.68 (d, $J$ = 8.0 Hz, 1H), 6.67 (s, 1H), 6.63 (dd, $J$ = 8.0, 2.0 Hz, 1H), 6.52 (s, 1H), 4.05 (dd, $J$ = 8.6, 4.3 Hz, 1H), 3.75 (s, 3H), 3.71 (s, 3H), 3.26 – 3.19 (m, 1H), 2.99 (dt, $J$ = 11.5, 4.0 Hz, 1H), 2.78 – 2.54 (m, 4H), 2.14 – 2.05 (m, 1H), 2.05 – 1.94 (m, 1H). Asterisk at 2.08 ppm corresponds to acetone that was used to clean the spatula for weighing the compound.

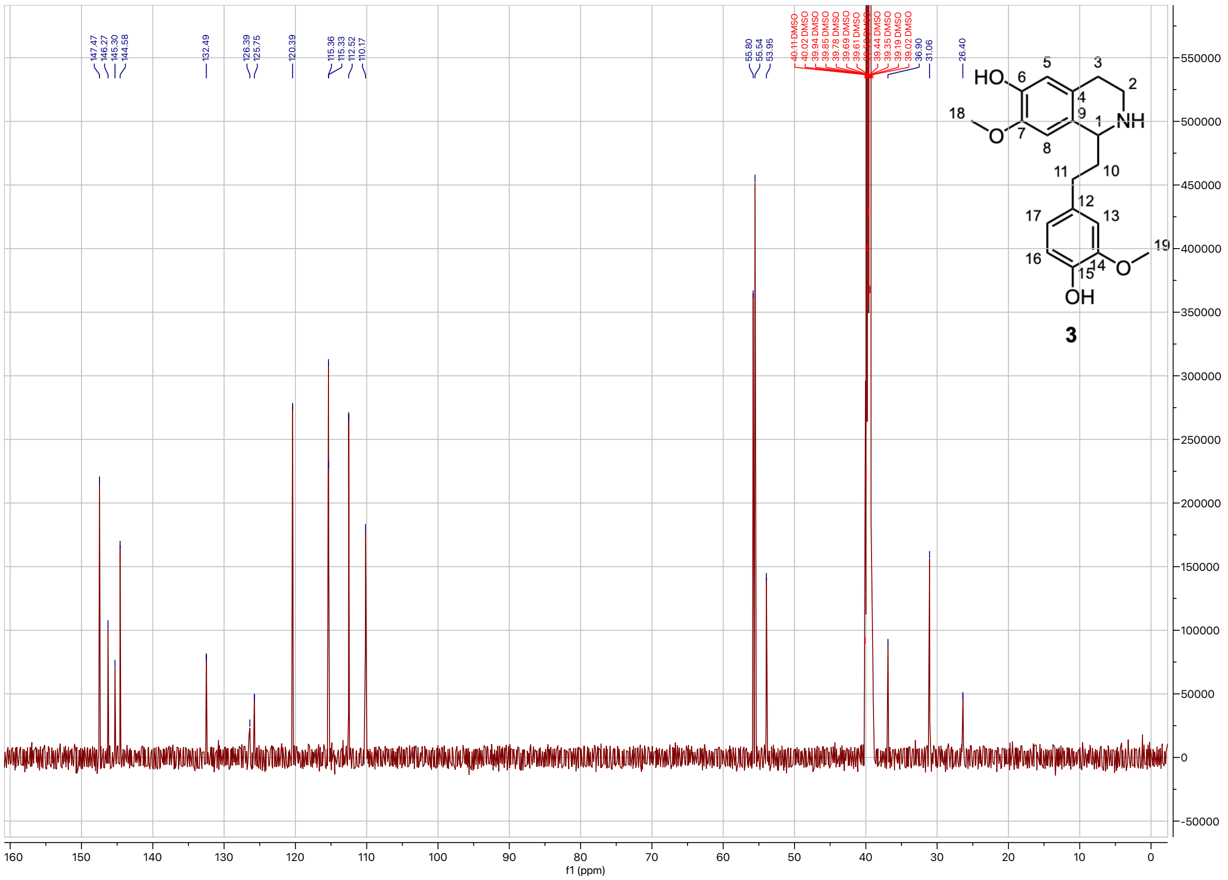

**Supplementary Figure. 24. $^{13}$C-NMR spectrum of 1-(4-hydroxy-3-methoxyphenethyl)-7-methoxy-1,2,3,4-tetrahydroisoquinolin-6-ol (3) (DMSO-d$_6$, 500 MHz, 298 K).**

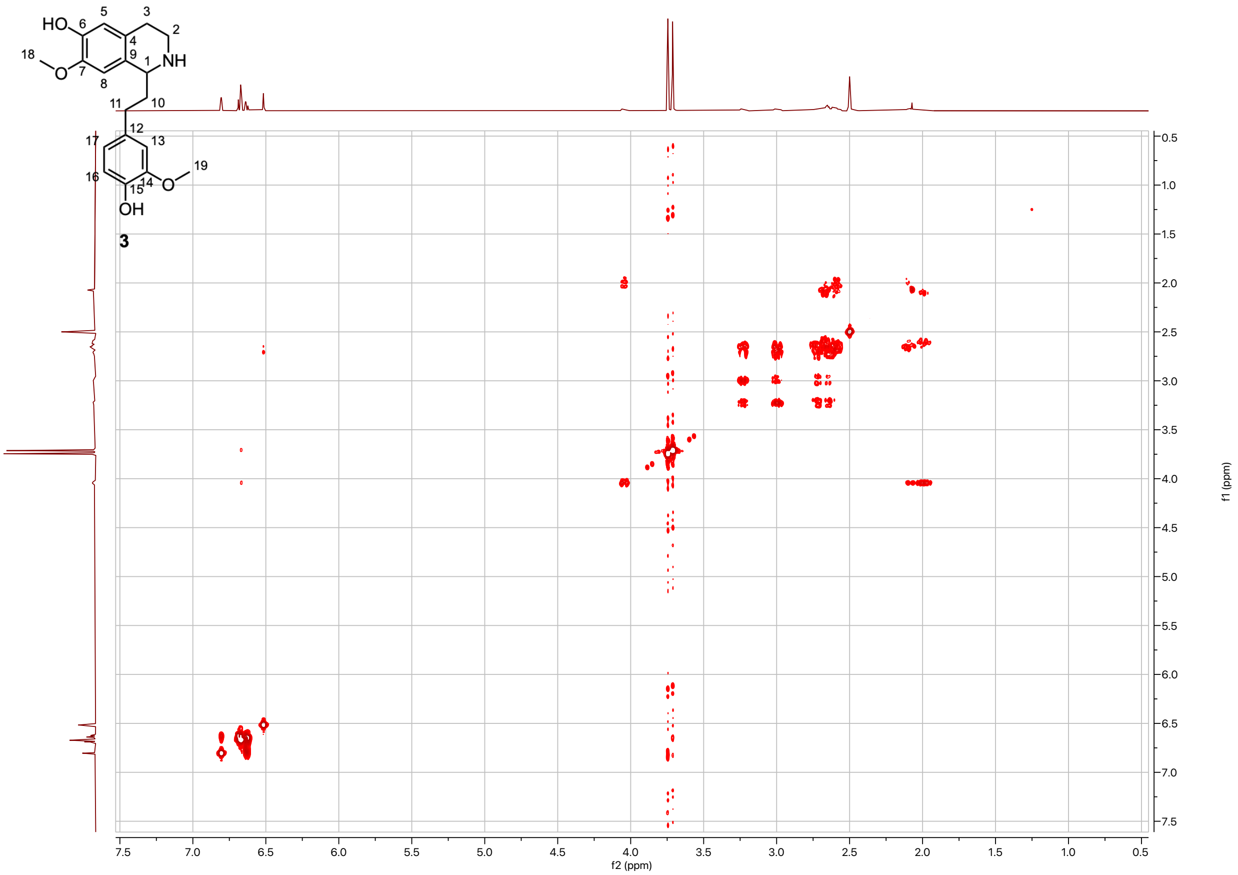


**Supplementary Figure. 25. COSY spectrum of 1-(4-hydroxy-3-methoxyphenethyl)-7-methoxy-1,2,3,4-tetrahydroisoquinolin-6-ol (3) (DMSO-d$_6$, 500 MHz, 298 K).**

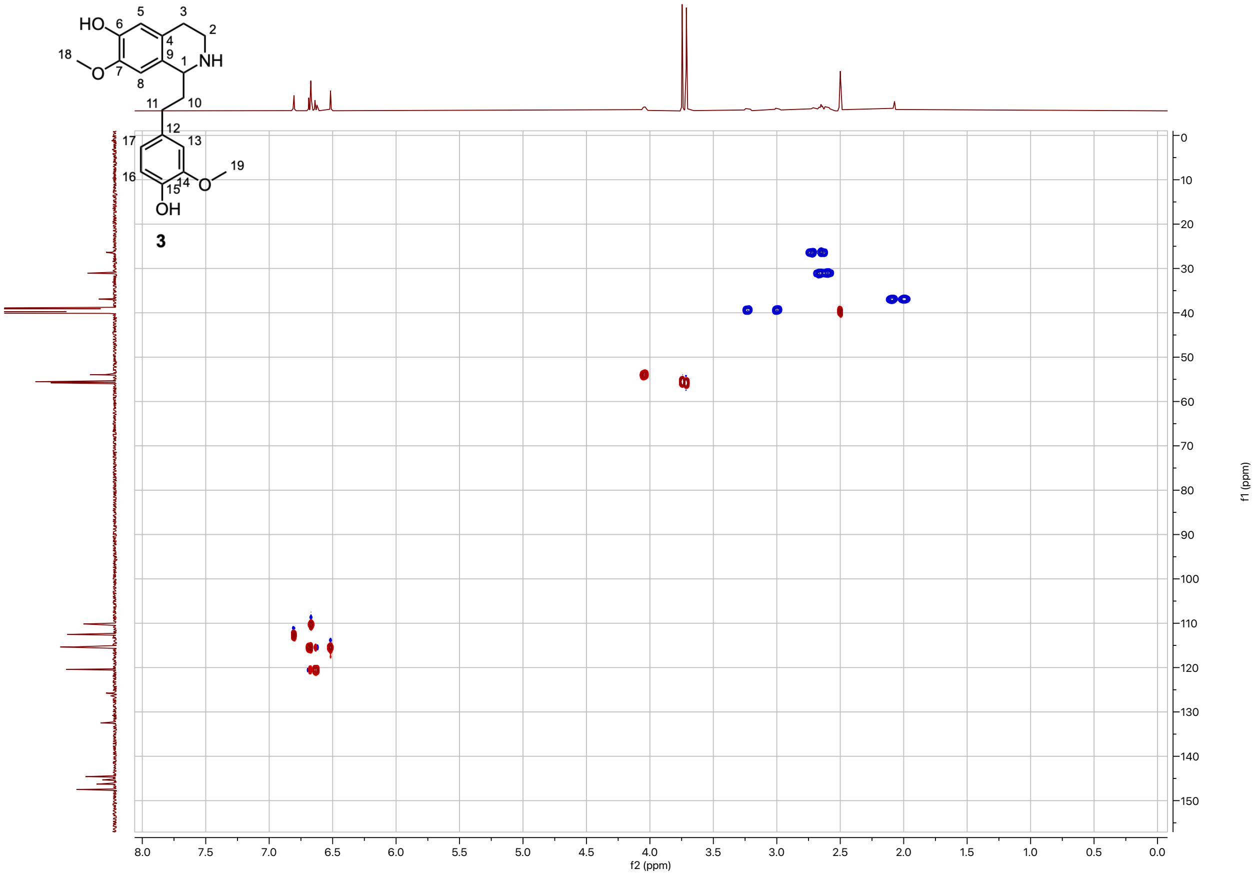


**Supplementary Figure. 26. HSQC spectrum of 1-(4-hydroxy-3-methoxyphenethyl)-7-methoxy-1,2,3,4-tetrahydroisoquinolin-6-ol (3) (DMSO-$d_6$, 500 MHz, 298 K).**

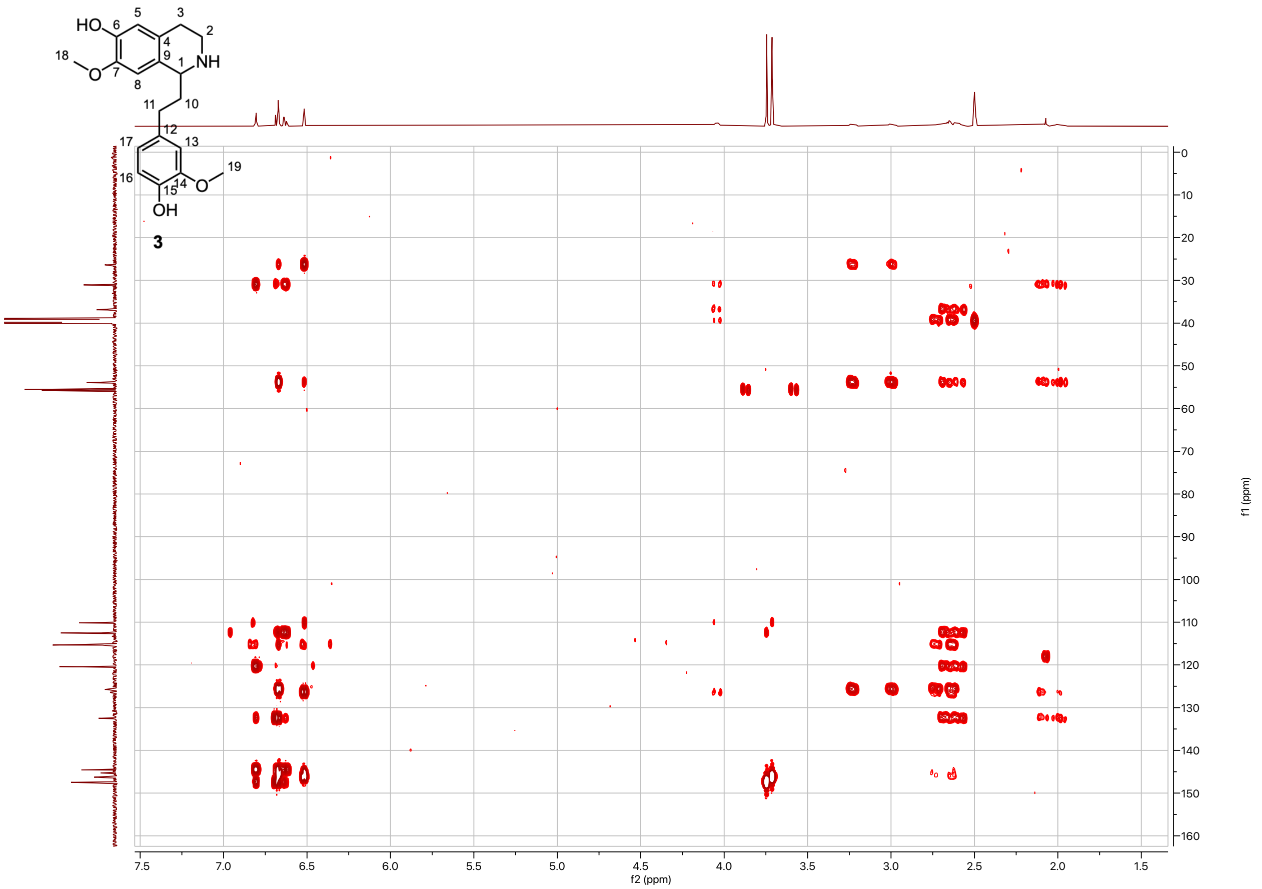


**Supplementary Figure. 27. HMBC spectrum of 1-(4-hydroxy-3-methoxyphenethyl)-7-methoxy-1,2,3,4-tetrahydroisoquinolin-6-ol (3) (DMSO-d$_6$, 500 MHz, 298 K).**

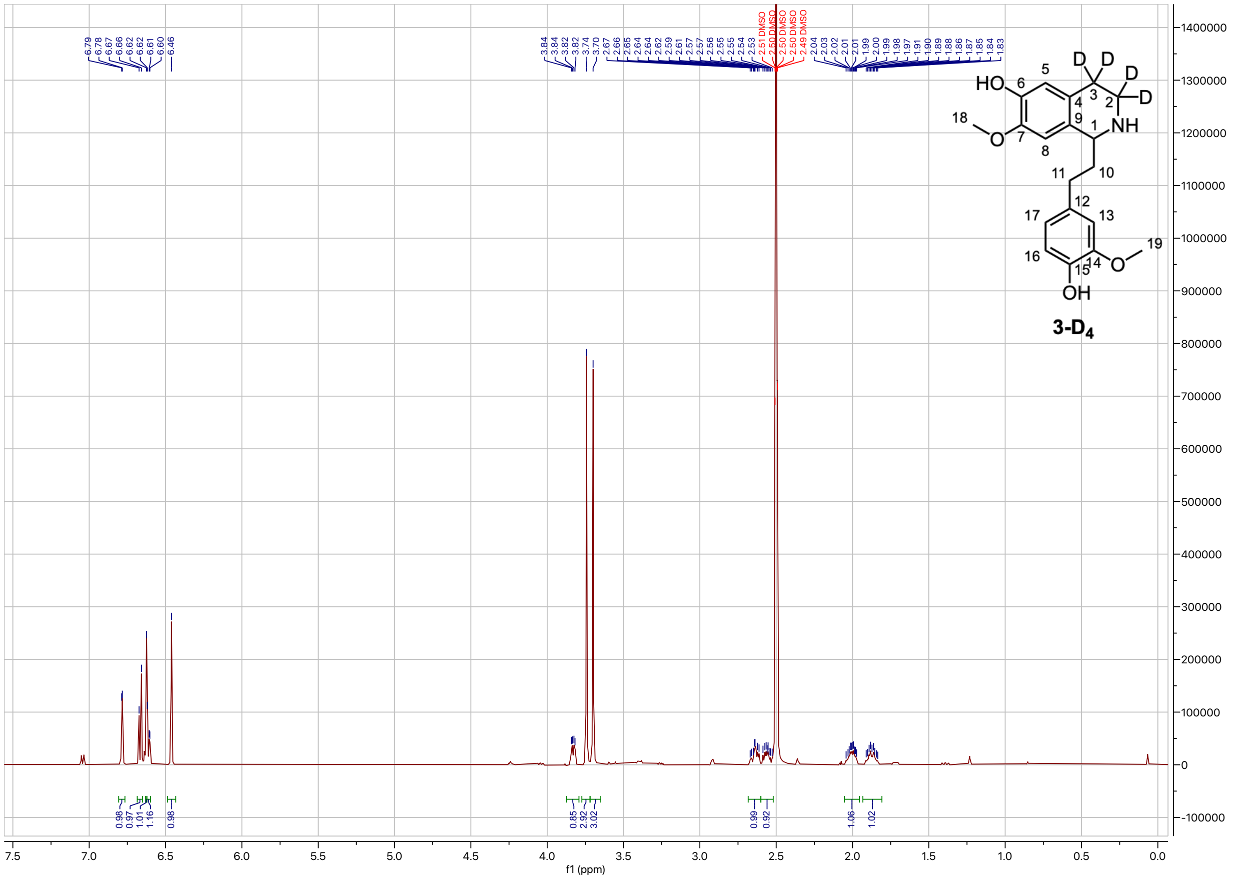

**Supplementary Figure. 28. $^1$H-NMR spectrum of 1-(4-hydroxy-3-methoxyphenethyl)-7-methoxy-1,2,3,4-tetrahydroisoquinolin-3,3,4,4-$d_4$-6-ol (3-D$_4$) (DMSO-d$_6$, 500 MHz, 298 K).** $^1$H NMR (500 MHz, DMSO) δ 6.79 (d, *J* = 2.1 Hz, 1H), 6.67 (d, *J* = 7.9 Hz, 1H), 6.62 (s, 1H), 6.6 (dd, *J* = 8.0, 1.9 Hz, 1H), 6.46 (s, 1H), 3.83 (dd, *J* = 8.8, 3.3 Hz, 1H), 3.74 (s, 3H), 3.70 (s, 3H), 2.68 – 2.60 (m, 1H), 2.60 – 2.52 (m, 1H), 2.05 – 1.95 (m, 1H), 1.93 – 1.81 (m, 1H).

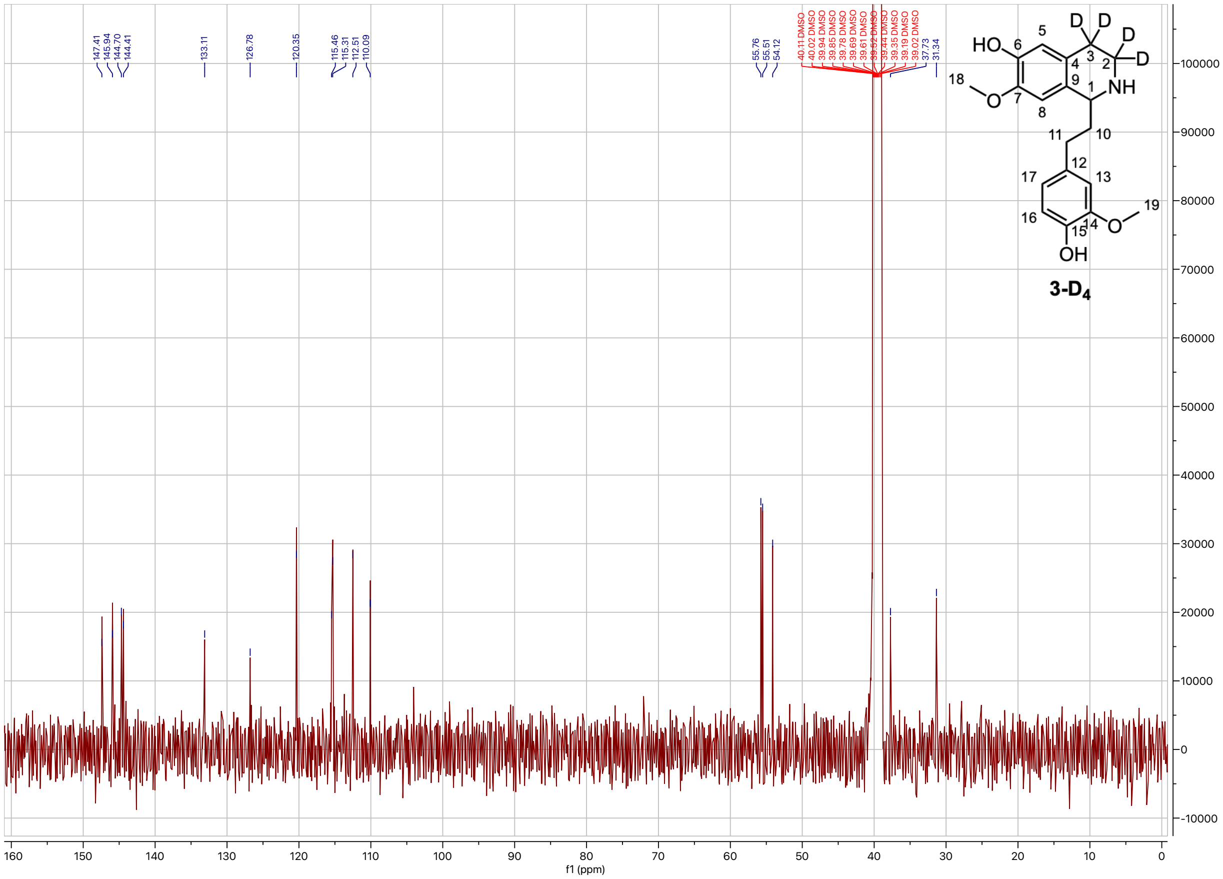

**Supplementary Figure. 29. $^{13}$C-NMR spectrum of 1-(4-hydroxy-3-methoxyphenethyl)-7-methoxy-1,2,3,4-tetrahydroisoquinolin-3,3,4,4-$d_4$-6-ol (3-D$_4$) (DMSO-d$_6$, 500 MHz, 298 K).**

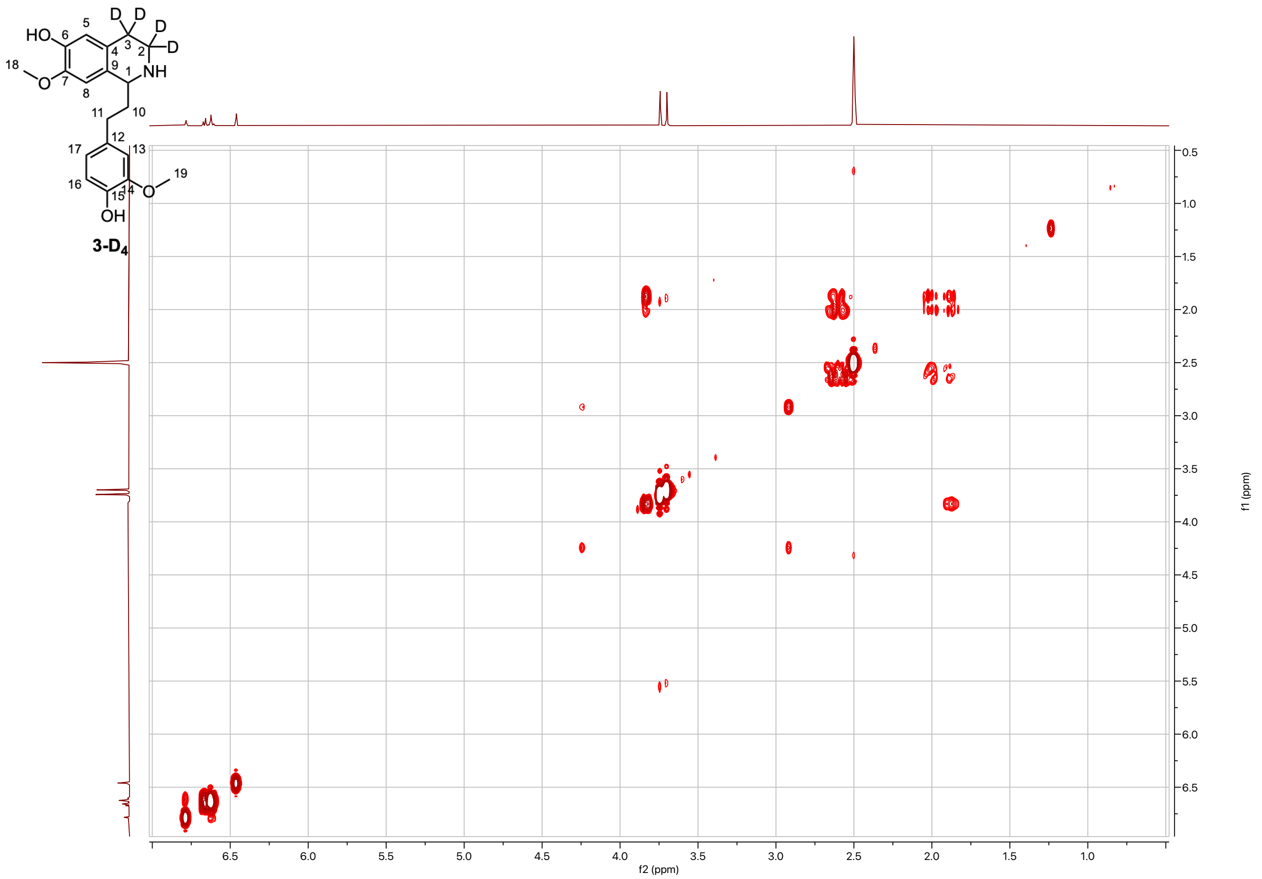


**Supplementary Figure. 30. COSY spectrum of 1-(4-hydroxy-3-methoxyphenethyl)-7-methoxy-1,2,3,4-tetrahydroisoquinolin-3,3,4,4-$d_4$-6-ol (3-D$_4$) (DMSO-d$_6$, 500 MHz, 298 K).**

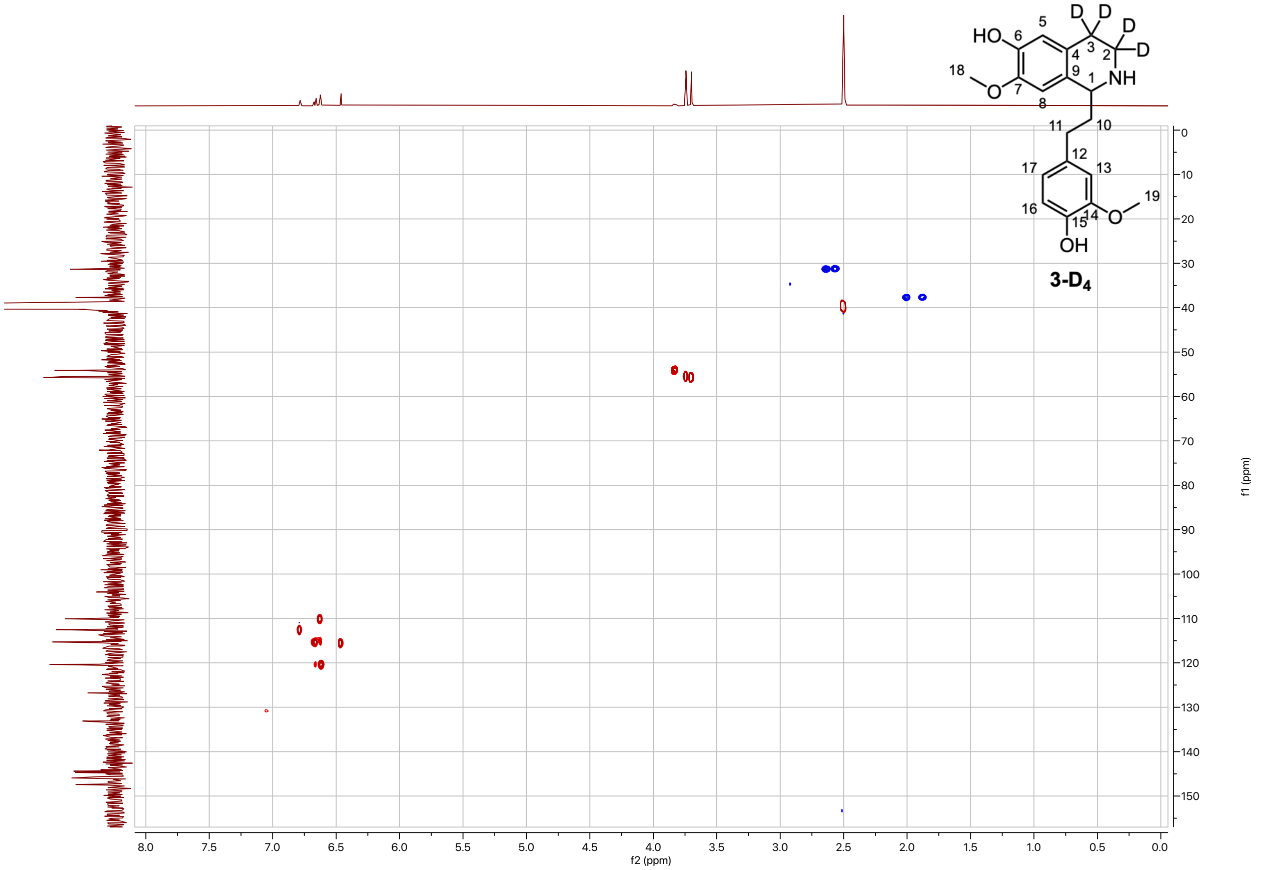


**Supplementary Figure. 31. HSQC spectrum of 1-(4-hydroxy-3-methoxyphenethyl)-7-methoxy-1,2,3,4-tetrahydroisoquinolin-3,3,4,4-*d*$_4$-6-ol (3-D$_4$) (DMSO-d$_6$, 500 MHz, 298 K).**

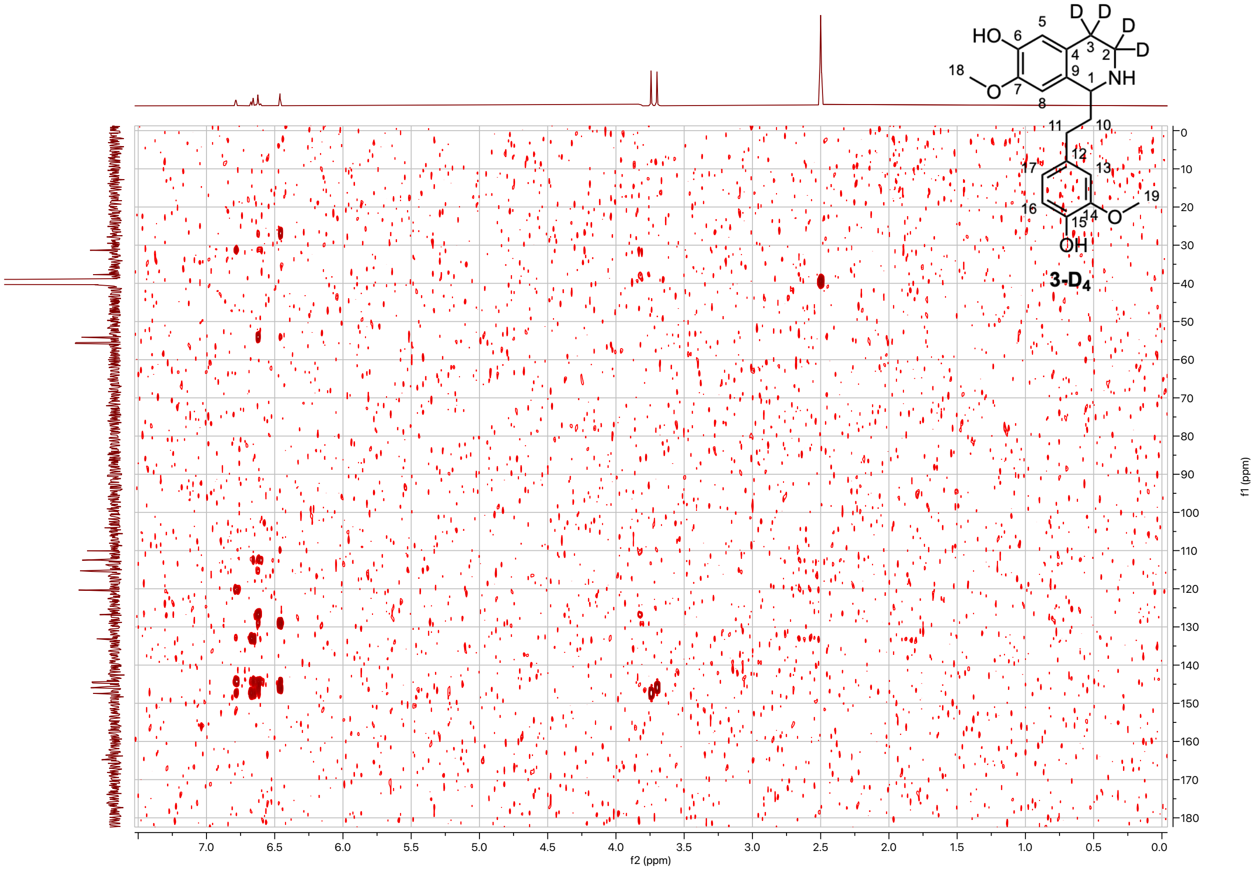


**Supplementary Figure. 32. HMBC spectrum of 1-(4-hydroxy-3-methoxyphenethyl)-7-methoxy-1,2,3,4-tetrahydroisoquinolin-3,3,4,4-$d_4$-6-ol (3-D$_4$) (DMSO-d$_6$, 500 MHz, 298 K).**

*

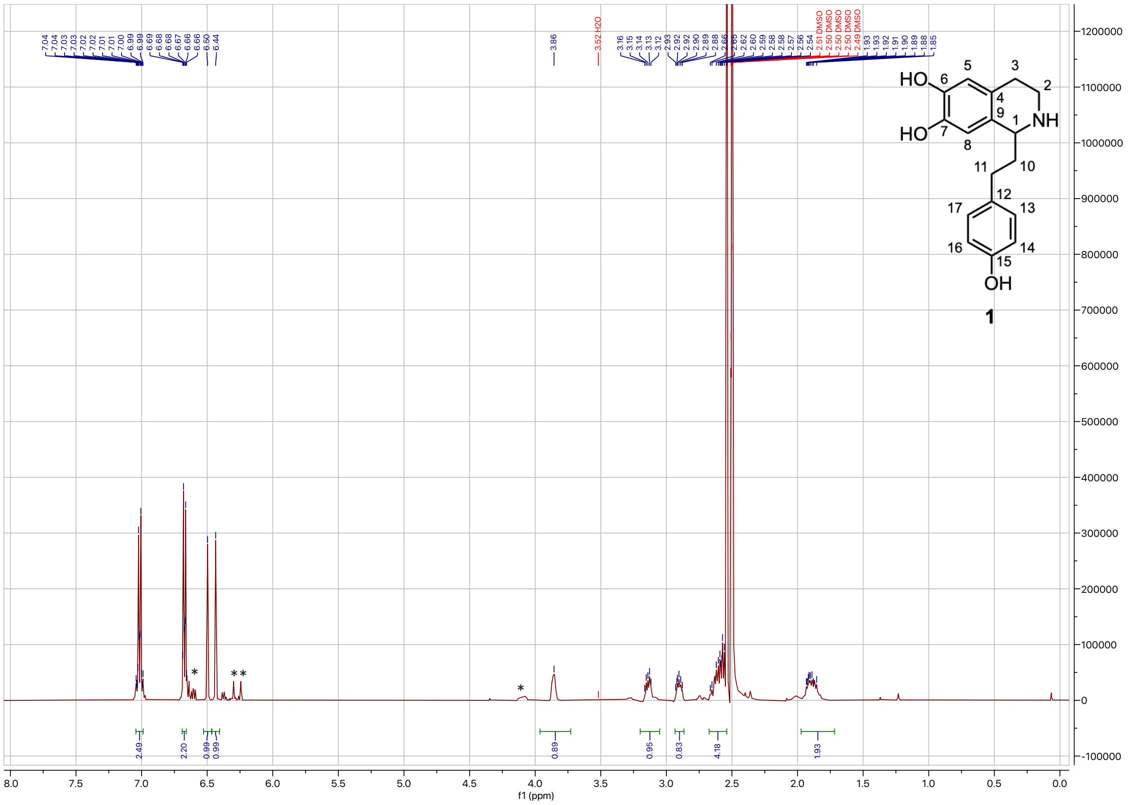


**Supplementary Figure. 33. $^{1}$H-NMR spectrum of 1-(4-hydroxyphenethyl)-1,2,3,4-tetrahydroisoquinoline-6,7-diol (1) (DMSO-$d_6$, 500 MHz, 298 K).** $^{1}$H NMR (500 MHz, DMSO) δ 7.04 – 6.99 (m, 2H), 6.69 – 6.66 (m, 2H), 6.50 (s, 1H), 6.44 (s, 1H), 3.90 – 3.81 (m, 1H), 3.14 (dt, $J$ = 11.7, 5.6 Hz, 1H), 2.90 (dt, $J$ = 12.8, 6.4 Hz, 1H), 2.67 – 2.54 (m, 4H), 1.97 – 1.78 (m, 2H). The sharp intense peak at 2.54ppm corresponds to protonated DMSO. Asterisk indicates impurities.

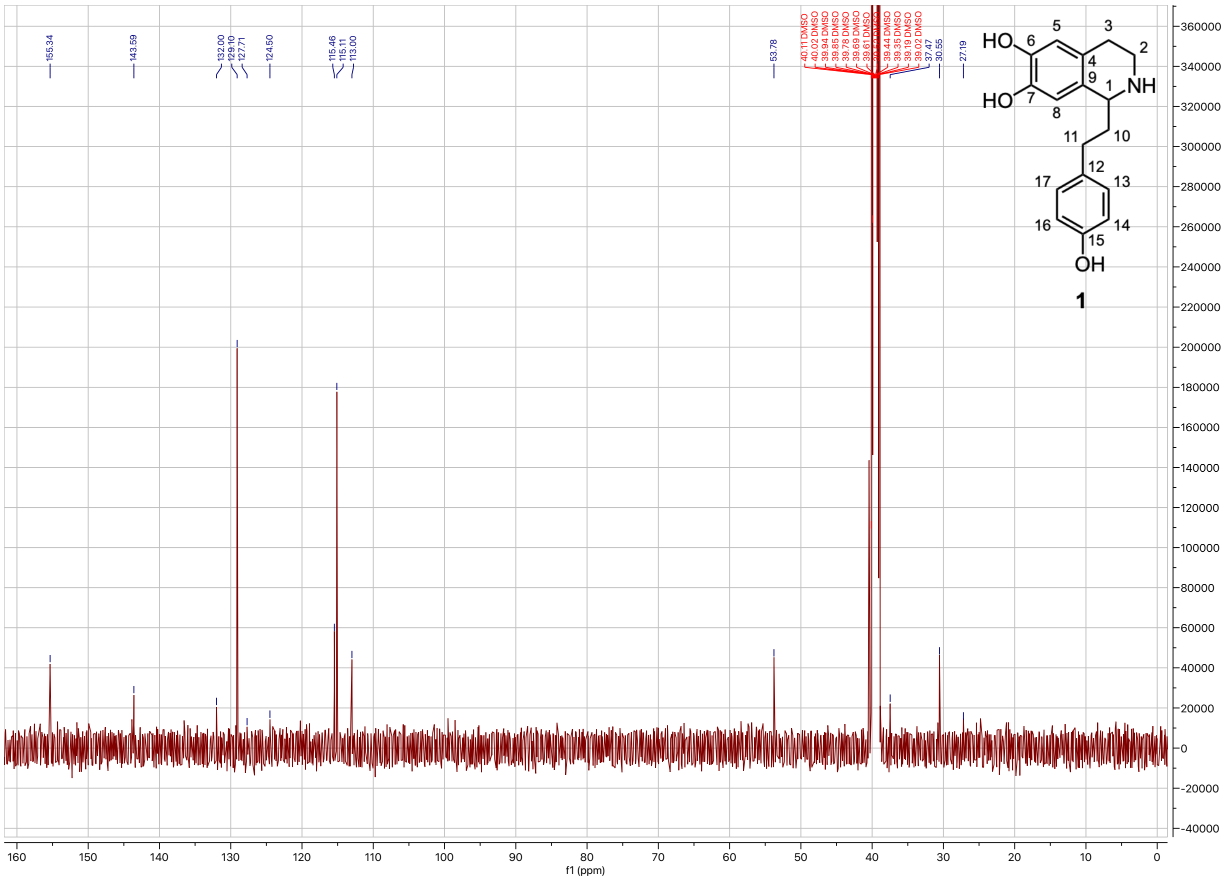

**Supplementary Figure. 34. $^{13}$C-NMR spectrum of 1-(4-hydroxyphenethyl)-1,2,3,4-tetrahydroisoquinoline-6,7-diol (1) (DMSO-$d_6$, 500 MHz, 298 K).**

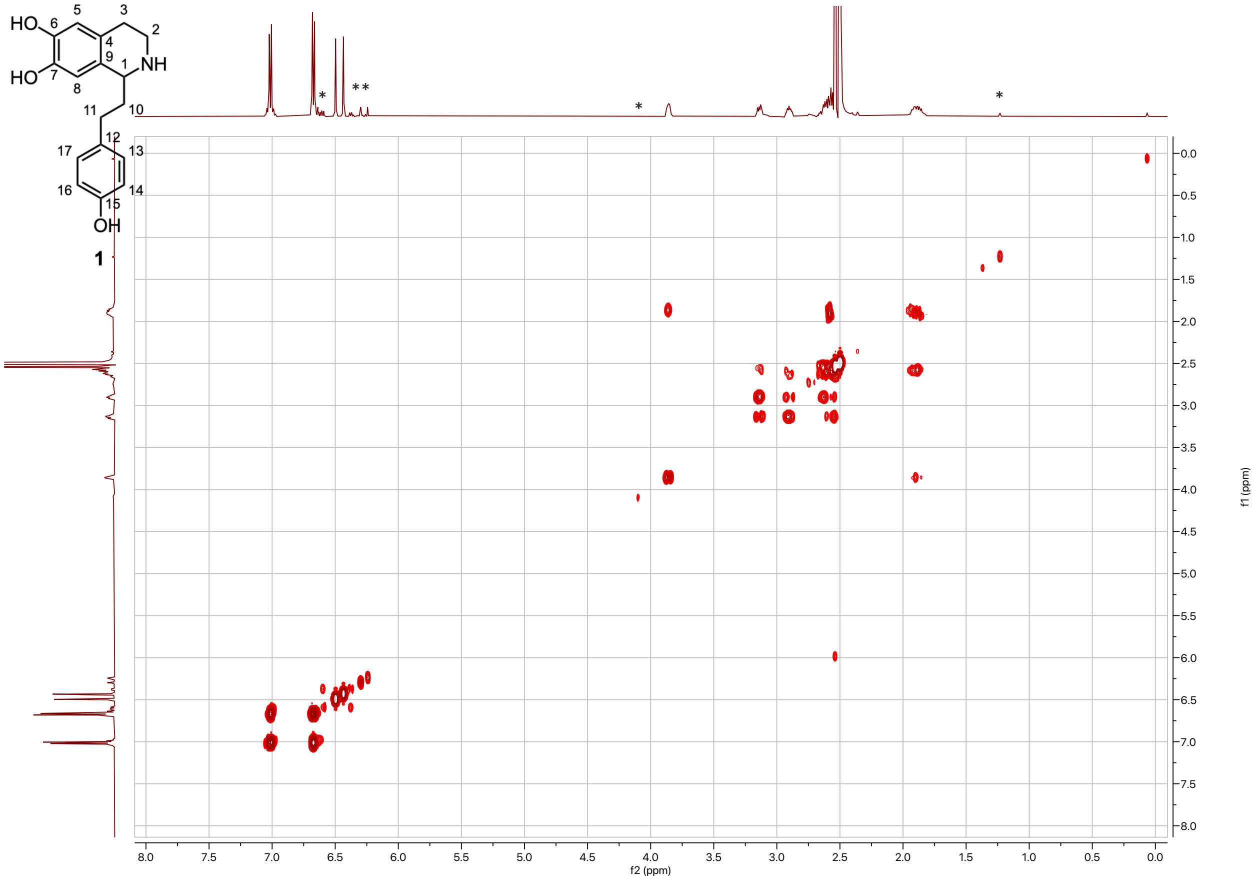


**Supplementary Figure. 35. COSY spectrum of 1-(4-hydroxyphenethyl)-1,2,3,4-tetrahydroisoquinoline-6,7-diol (1) (DMSO-$d_6$, 500 MHz, 298 K).** Asterisk indicates impurities.

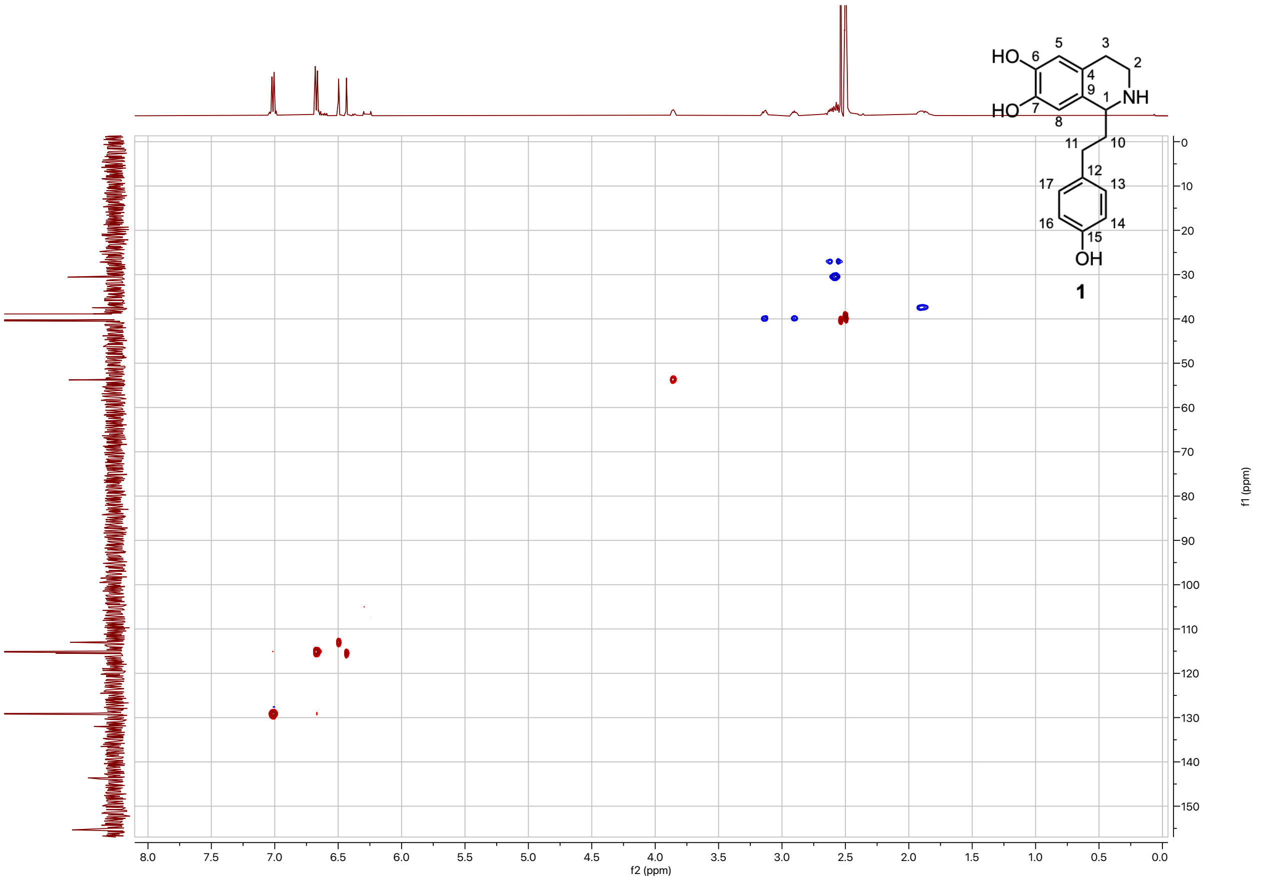


**Supplementary Figure. 36. HSQC spectrum of 1-(4-hydroxyphenethyl)-1,2,3,4-tetrahydroisoquinoline-6,7-diol (1) (DMSO-$d_6$, 500 MHz, 298 K).**

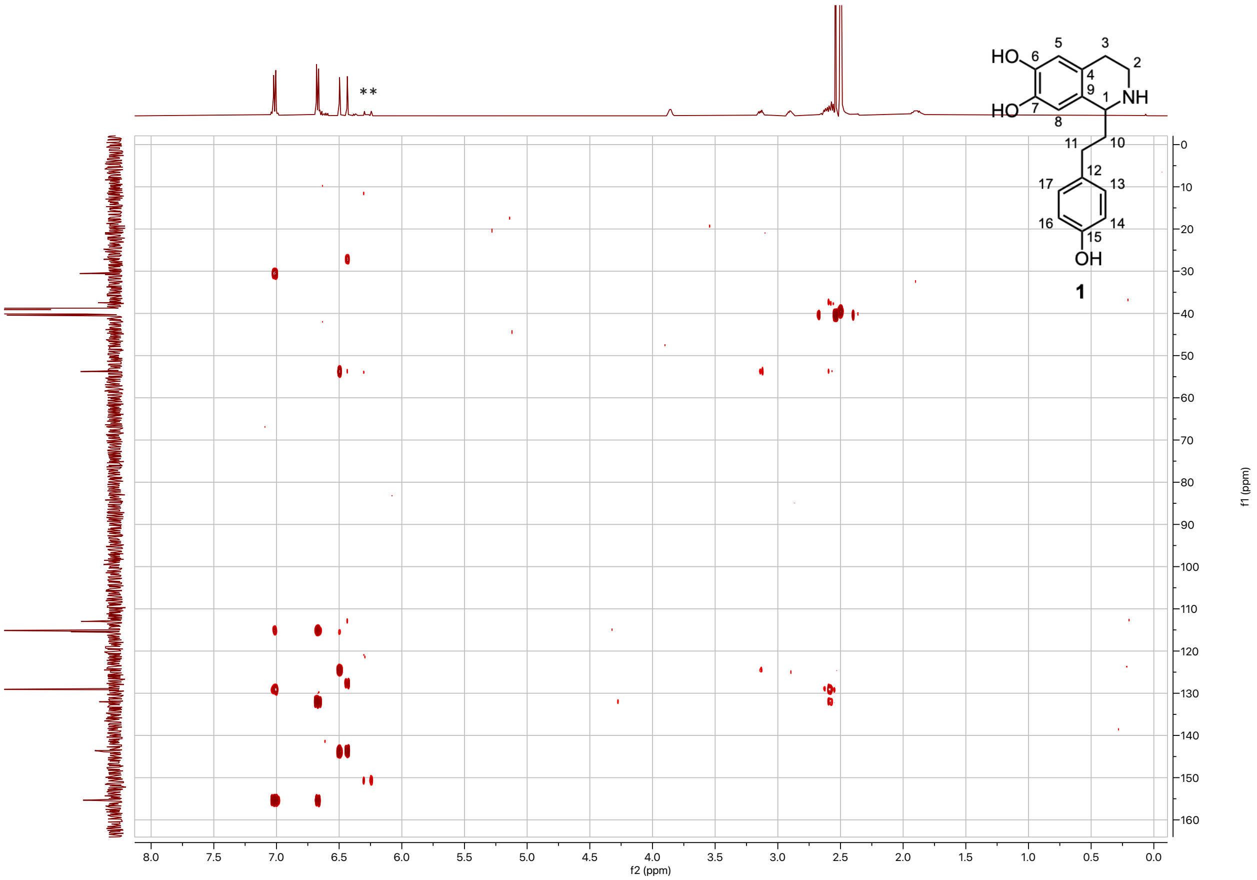


**Supplementary Figure. 37. HMBC spectrum of 1-(4-hydroxyphenethyl)-1,2,3,4-tetrahydroisoquinoline-6,7-diol (1) (DMSO-d₆, 500 MHz, 298 K).**

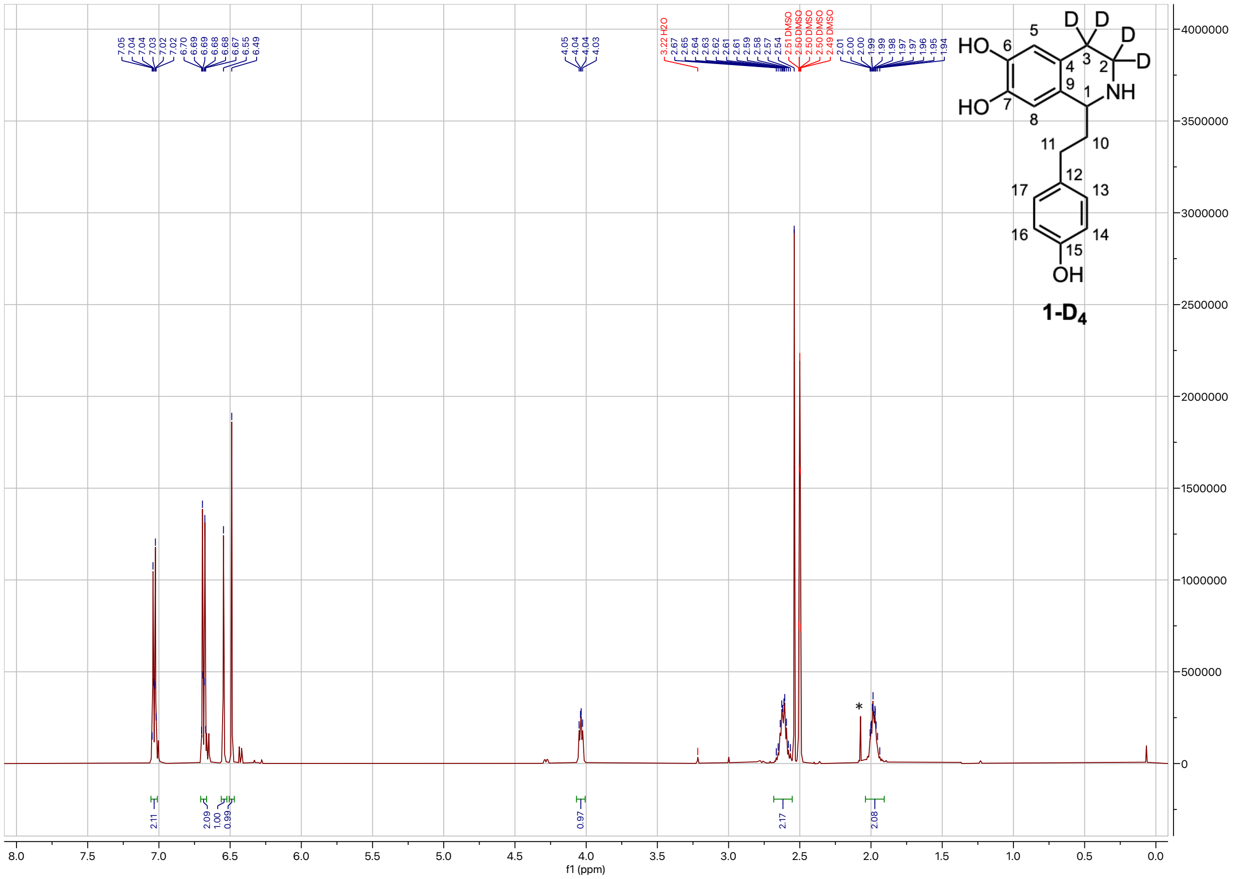

**Supplementary Figure. 38. $^{1}$H-NMR spectrum of 1-(4-hydroxyphenethyl)-1,2,3,4-tetrahydroisoquinoline-3,3,4,4-*d*$_4$-6,7-diol (1-D$_4$) (DMSO-d$_6$, 500 MHz, 298 K).** $^{1}$H NMR (500 MHz, DMSO) δ 7.06 – 7.01 (m, 2H), 6.71 – 6.67 (m, 2H), 6.55 (s, 1H), 6.49 (s, 1H), 4.04 (dd, *J* = 7.3, 5.0 Hz, 1H), 2.68 – 2.55 (m, 2H), 2.04 – 1.91 (m, 2H). Asterisk at 2.08 ppm corresponds to acetone that was used to clean the spatula for weighing the compound. The sharp intense peak at 2.54ppm corresponds to protonated DMSO.

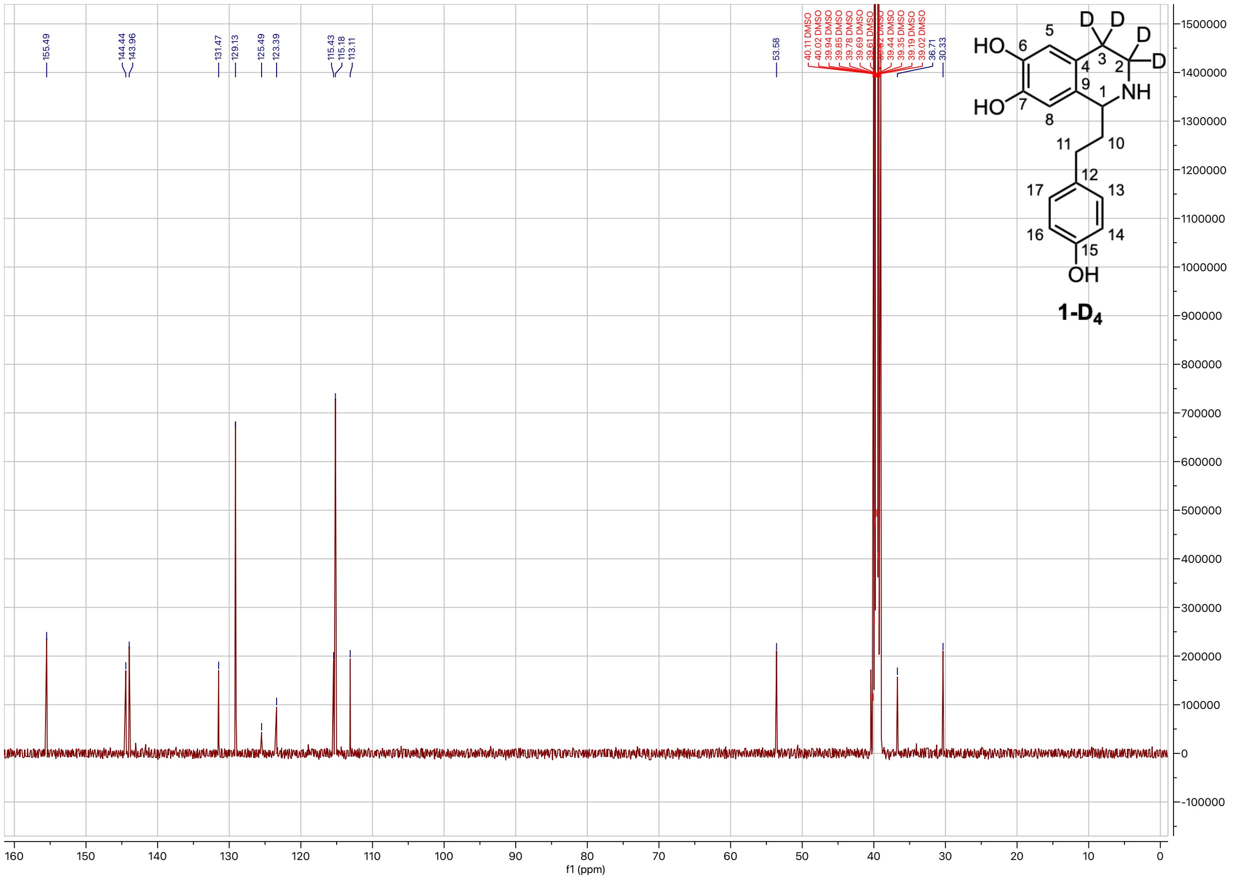

**Supplementary Figure. 39.** $^{13}$**C-NMR spectrum of 1-(4-hydroxyphenethyl)-1,2,3,4-tetrahydroisoquinoline-3,3,4,4-*d*$_4$-6,7-diol (1-D$_4$) (DMSO-d$_6$, 500 MHz, 298 K).**

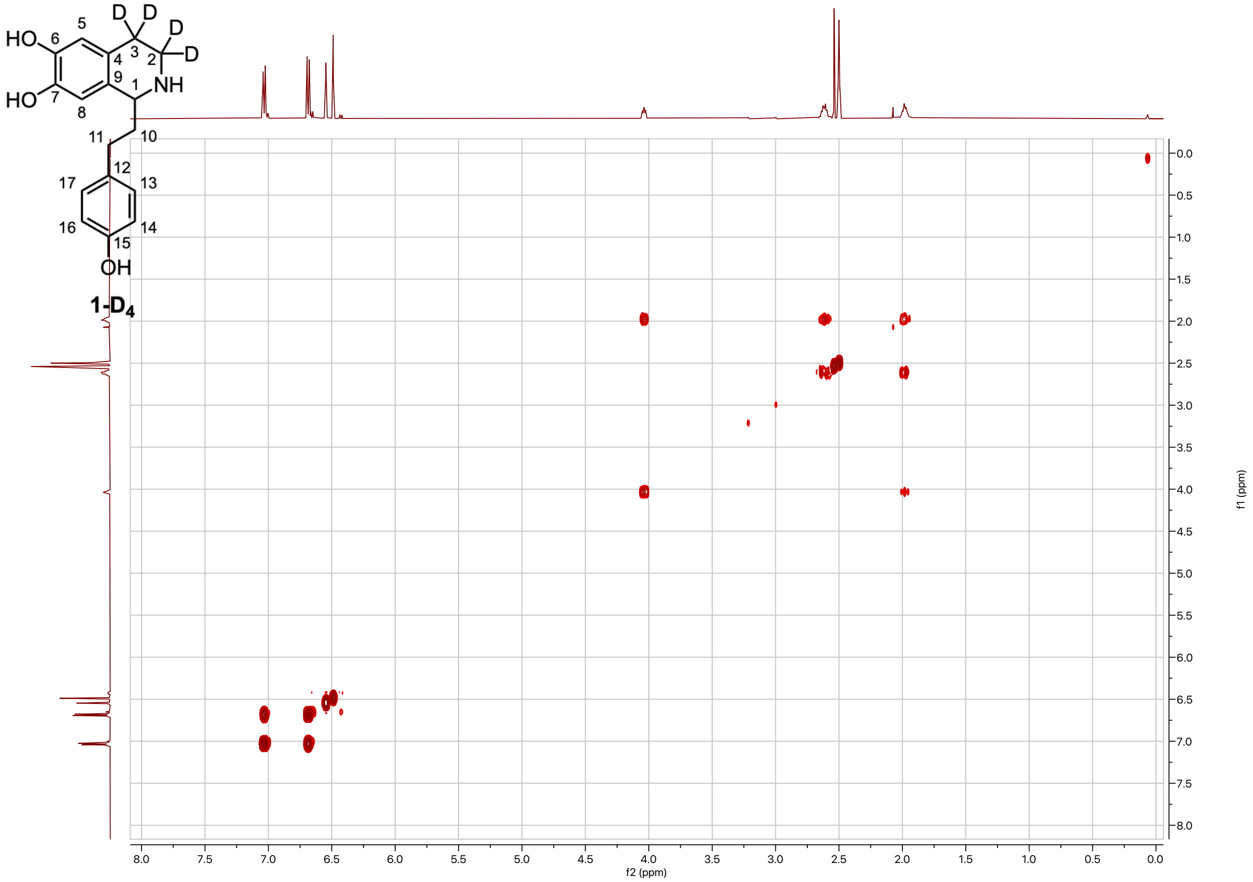


**Supplementary Figure. 40. COSY spectrum of 1-(4-hydroxyphenethyl)-1,2,3,4-tetrahydroisoquinoline-3,3,4,4-$d_4$-6,7-diol (1-D$_4$) (DMSO-d$_6$, 500 MHz, 298 K).**

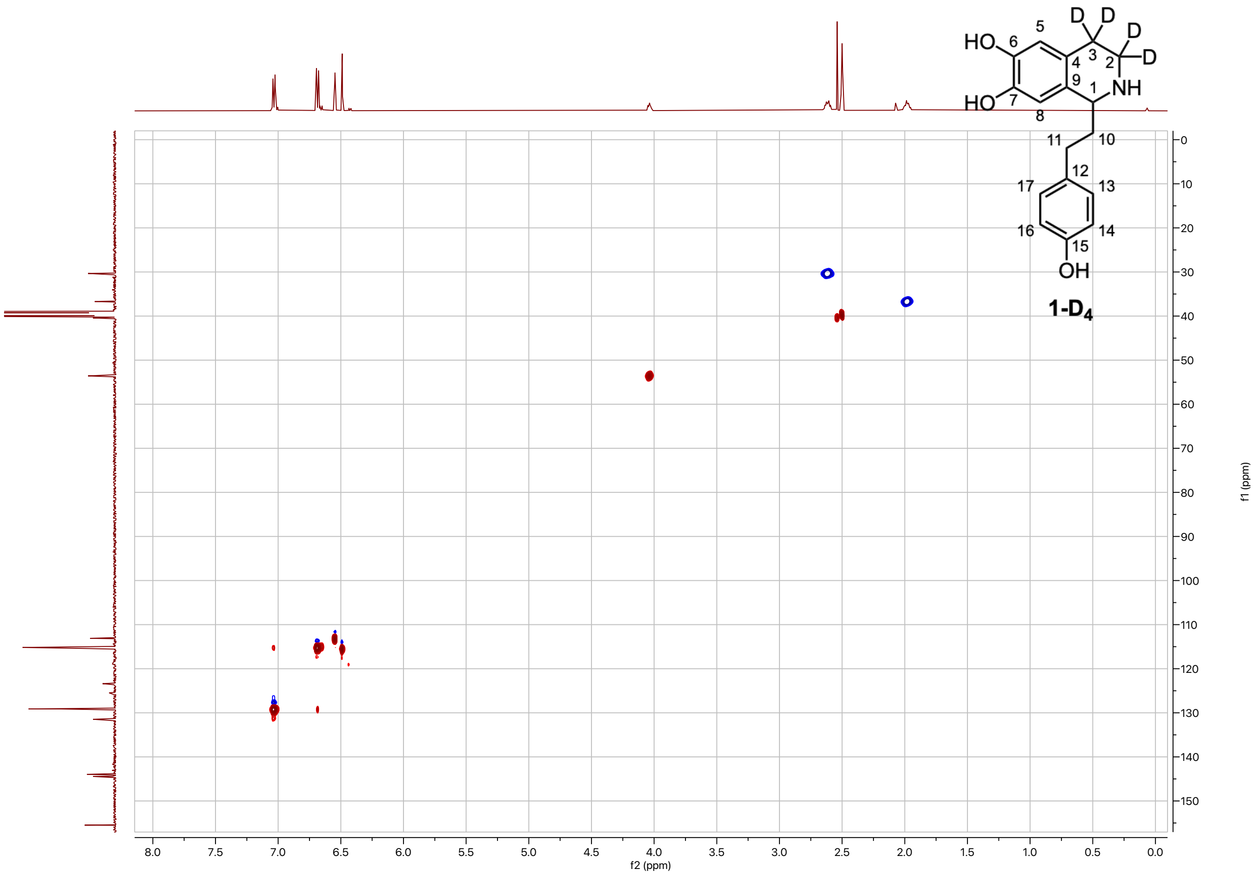


**Supplementary Figure. 41. HSQC spectrum of 1-(4-hydroxyphenethyl)-1,2,3,4-tetrahydroisoquinoline-3,3,4,4-*d*$_4$-6,7-diol (1-D$_4$) (DMSO-d$_6$, 500 MHz, 298 K).**

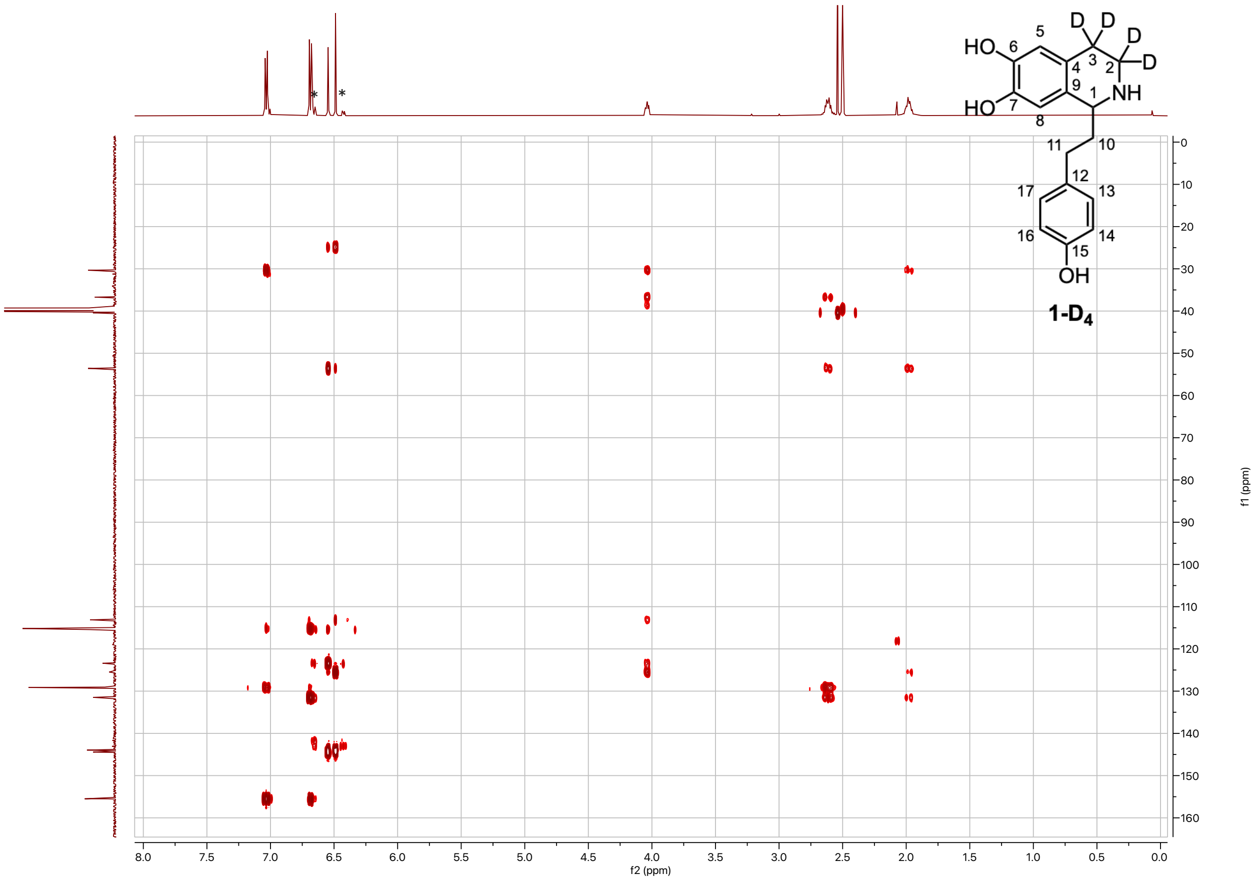


**Supplementary Figure. 42. HMBC spectrum of 1-(4-hydroxyphenethyl)-1,2,3,4-tetrahydroisoquinoline-3,3,4,4-$d_4$-6,7-diol (1-D4) (DMSO-d6, 500 MHz, 298 K).** Asterisk indicates impurities.

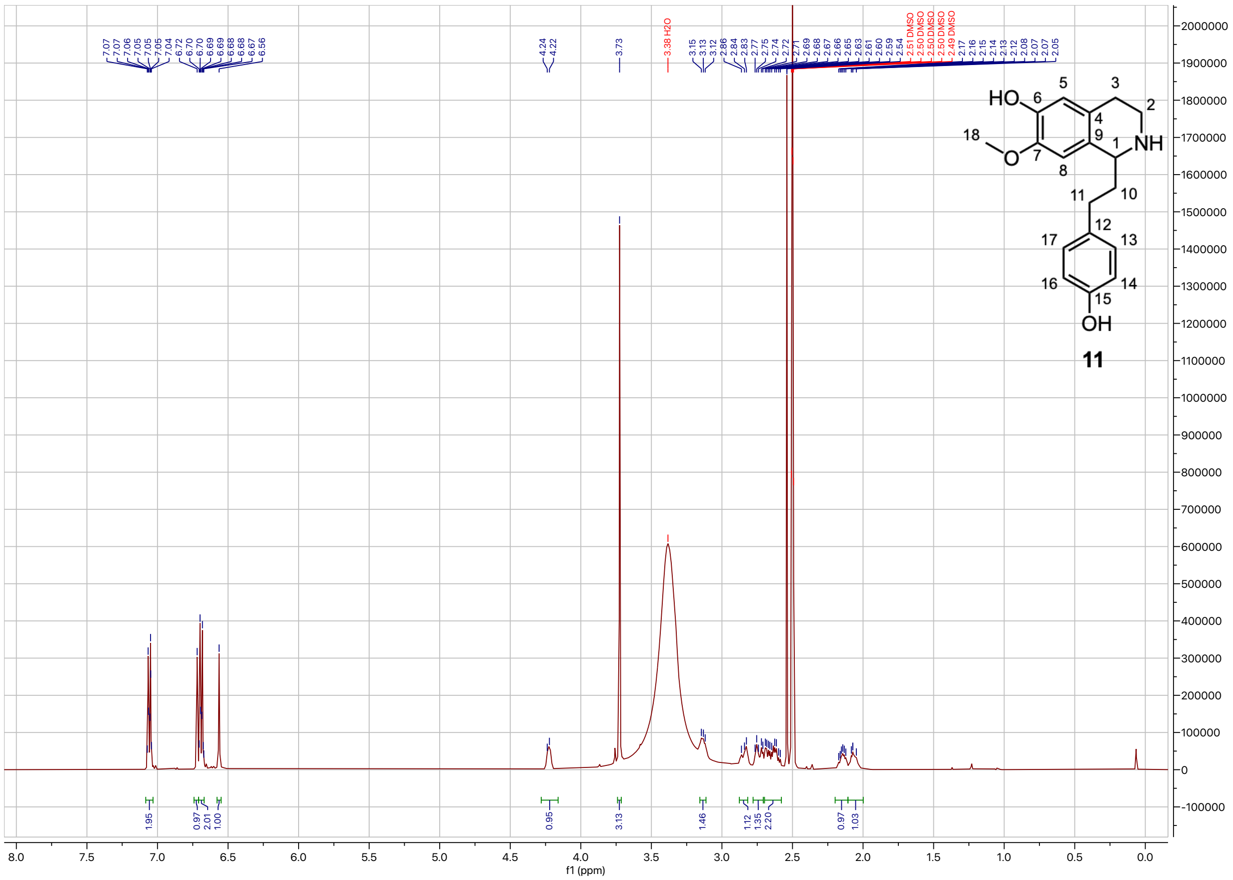

**Supplementary Figure. 43. $^{1}$H-NMR spectrum of 11-(4-hydroxyphenethyl)-7-methoxy-1,2,3,4-tetrahydroisoquinolin-6-ol (11) (DMSO-$d_6$, 500 MHz, 298 K).** $^{1}$H NMR (500 MHz, DMSO) δ 7.08 – 7.03 (m, 2H), 6.72 (s, 1H), 6.71 – 6.67 (m, 2H), 6.56 (s, 1H), 4.27 – 4.18 (m, 1H), 3.73 (s, 3H), 3.16 – 3.11 (m, 1H), 2.88 – 2.82 (m, 1H), 2.78 – 2.71 (m, 1H), 2.70 – 2.58 (m, 2H), 2.20 – 2.11 (m, 1H), 2.10 – 2.00 (m, 1H). The broad large peak at 3.38ppm corresponds to water. The sharp intense peak at 2.54ppm corresponds to protonated DMSO.

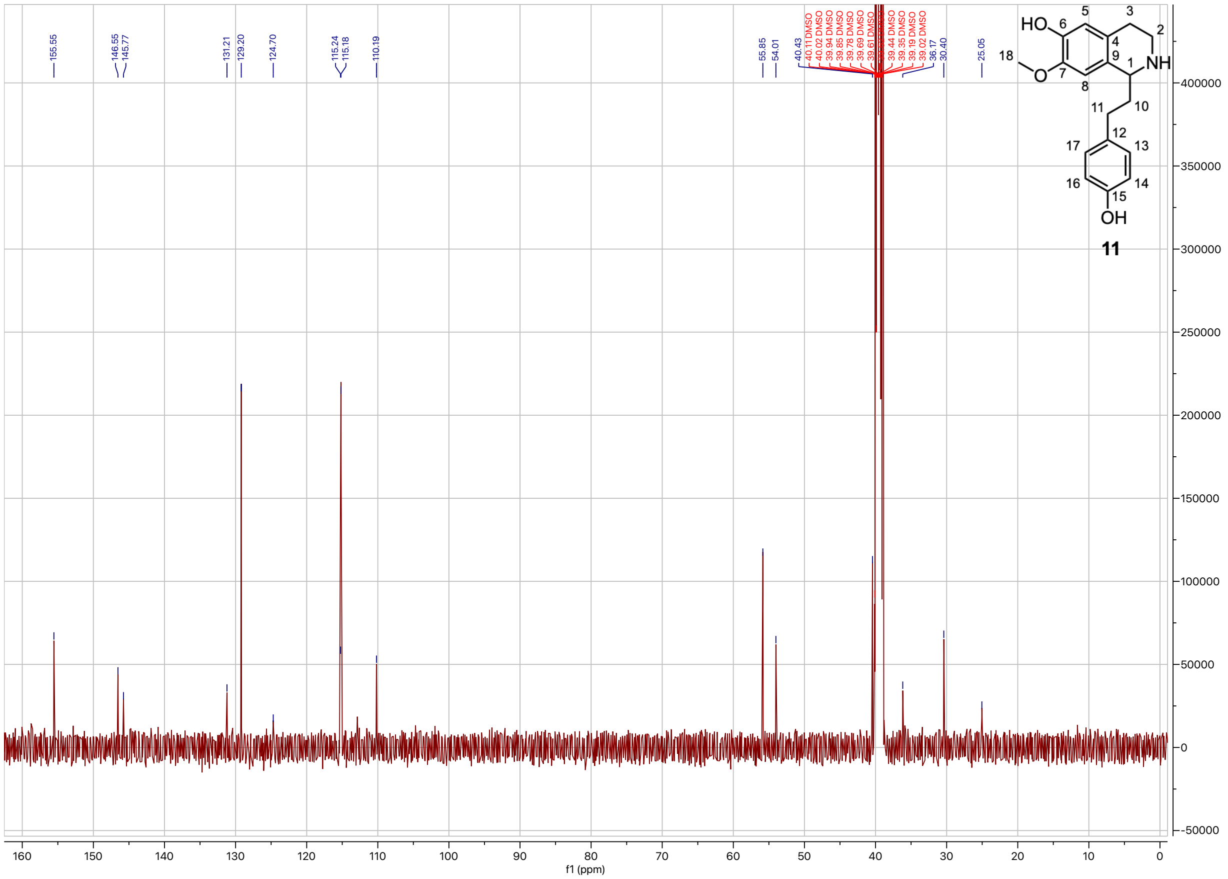

**Supplementary Figure. 44. $^{13}$C-NMR spectrum of 1-(4-hydroxyphenethyl)-7-methoxy-1,2,3,4-tetrahydroisoquinolin-6-ol (11) (DMSO-$d_6$, 500 MHz, 298 K).**

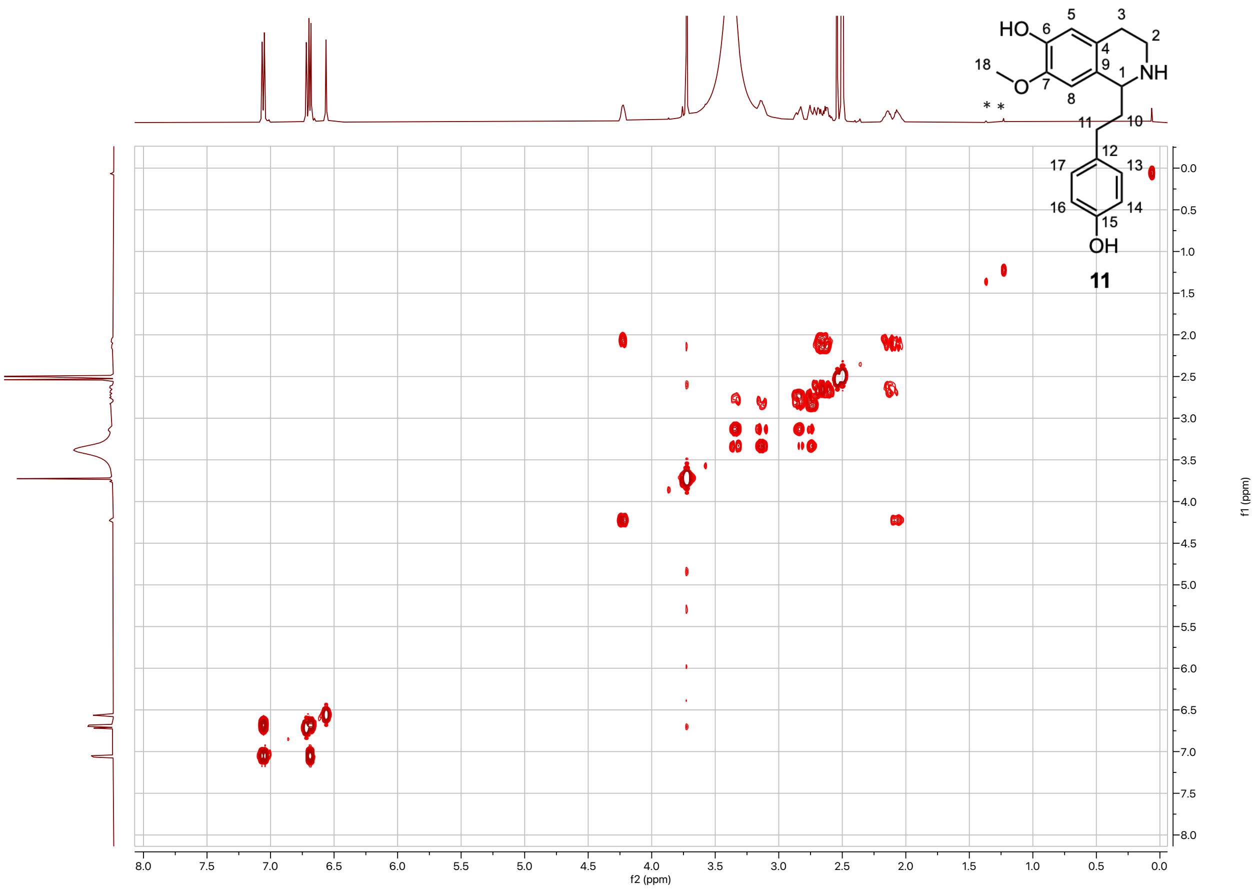


**Supplementary Figure. 45. COSY spectrum of 1-(4-hydroxyphenethyl)-7-methoxy-1,2,3,4-tetrahydroisoquinolin-6-ol (11) (DMSO-d$_6$, 500 MHz, 298 K).** The broad large peak at 3.38ppm corresponds to water. The sharp intense peak at 2.54ppm corresponds to protonated DMSO. Asterisk indicates impurities.

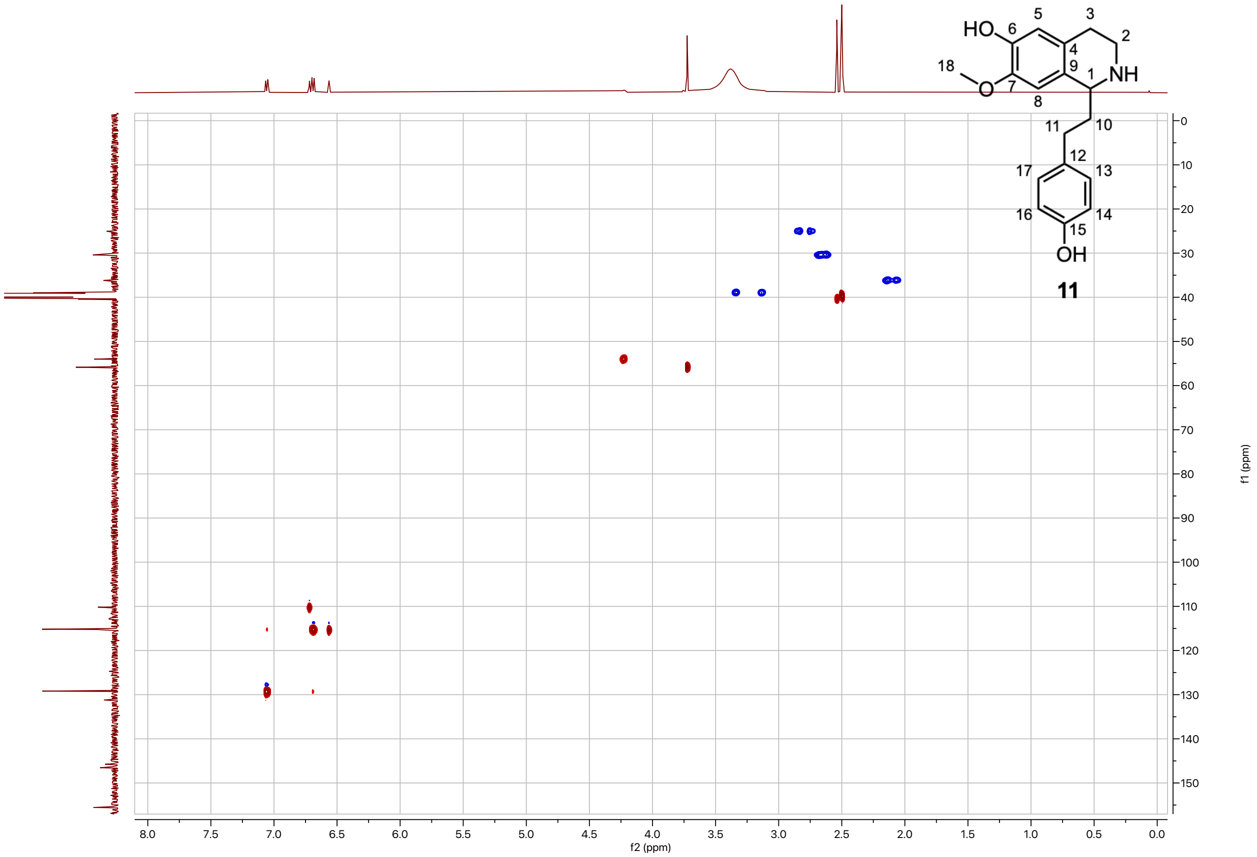

**Supplementary Figure. 46. HSQC spectrum of 1-(4-hydroxyphenethyl)-7-methoxy-1,2,3,4-tetrahydroisoquinolin-6-ol (11) (DMSO-$d_6$, 500 MHz, 298 K).**

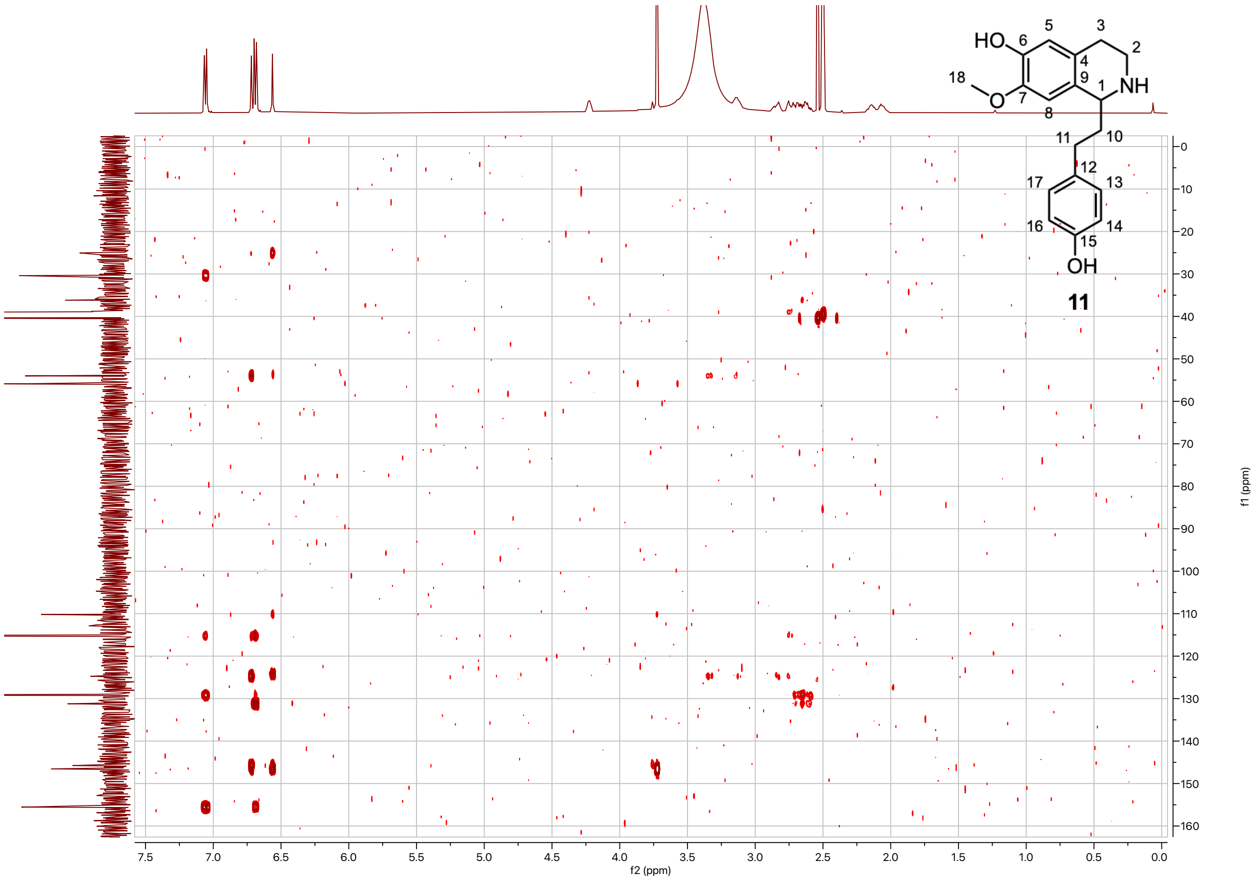


**Supplementary Figure. 47. HMBC spectrum of 1-(4-hydroxyphenethyl)-7-methoxy-1,2,3,4-tetrahydroisoquinolin-6-ol (11) (DMSO-d₆, 500 MHz, 298 K).**

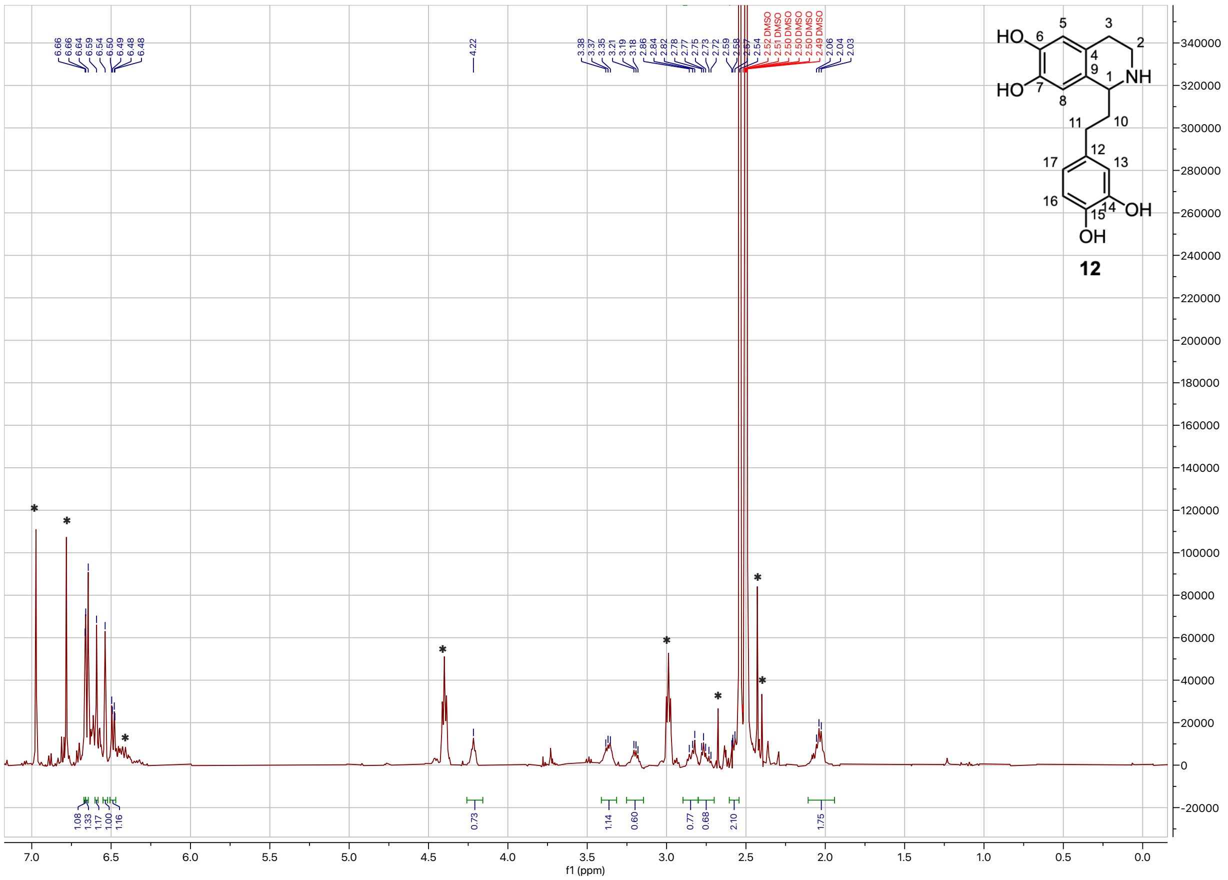

**Supplementary Figure. 48. [1]H-NMR spectrum of 1-(3,4-dihydroxyphenethyl)-1,2,3,4-tetrahydroisoquinoline-6,7-diol (12) (DMSO-$d_6$, 500 MHz, 298 K).** $^1$H NMR (500 MHz, DMSO) δ 6.66 (d, *J* = 1.7 Hz, 1H), 6.65 (d, *J* = 8.0 Hz, 1H), 6.59 (s, 1H), 6.54 (s, 1H), 6.49 (dd, *J* = 8.1, 1.9 Hz, 1H), 4.22 (t, *J* = 5.7 Hz, 1H), 3.41 – 3.31 (m, 1H), 3.21 (m, 1H), 2.90 – 2.80 (m, 1H), 2.80 – 2.70 (m, 1H), 2.60 – 2.54 (m, 2H), 2.04 (m, 2H). Asterisk indicates impurities. The sharp intense peak at 2.54ppm corresponds to protonated DMSO.

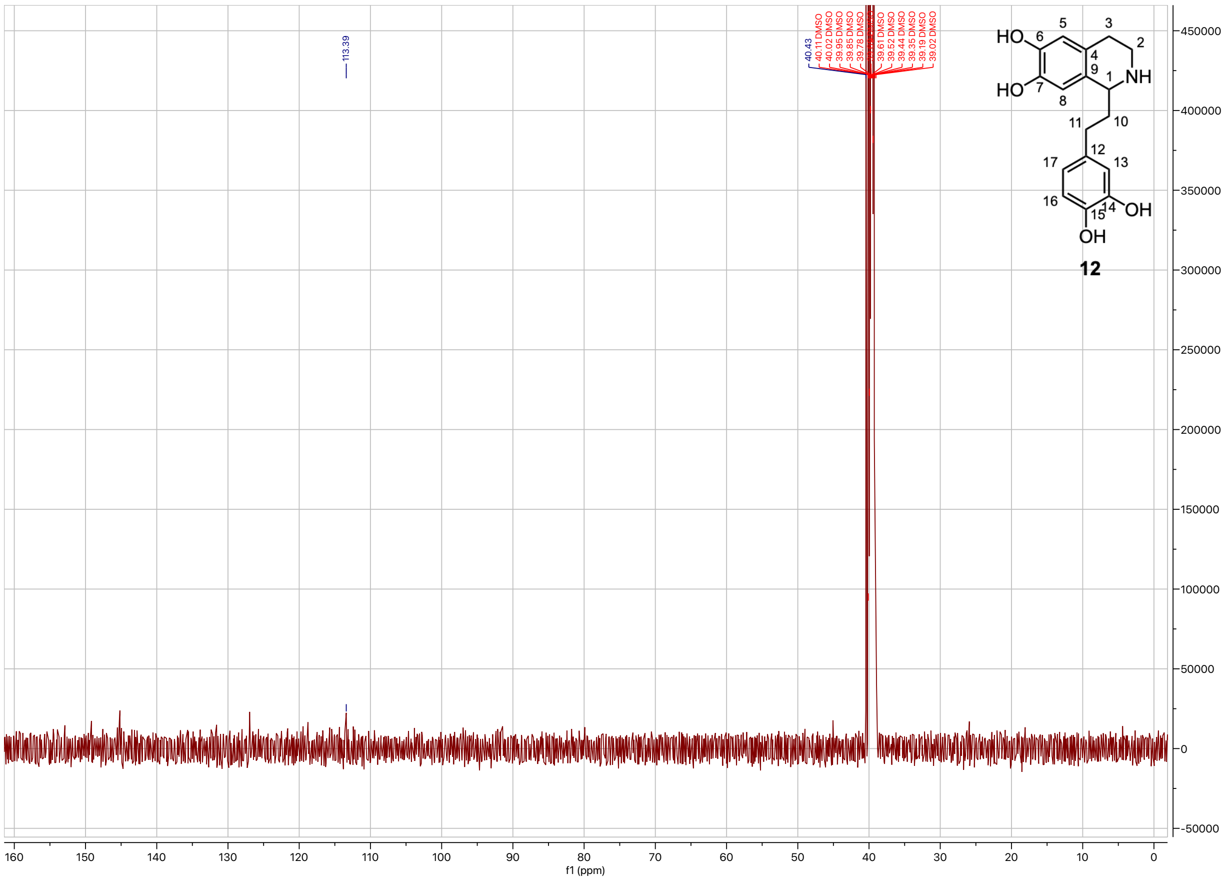

**Supplementary Figure. 49. $^{13}$C-NMR spectrum of 1-(3,4-dihydroxyphenethyl)-1,2,3,4-tetrahydroisoquinoline-6,7-diol (12) (DMSO-d$_6$, 500 MHz, 298 K).**

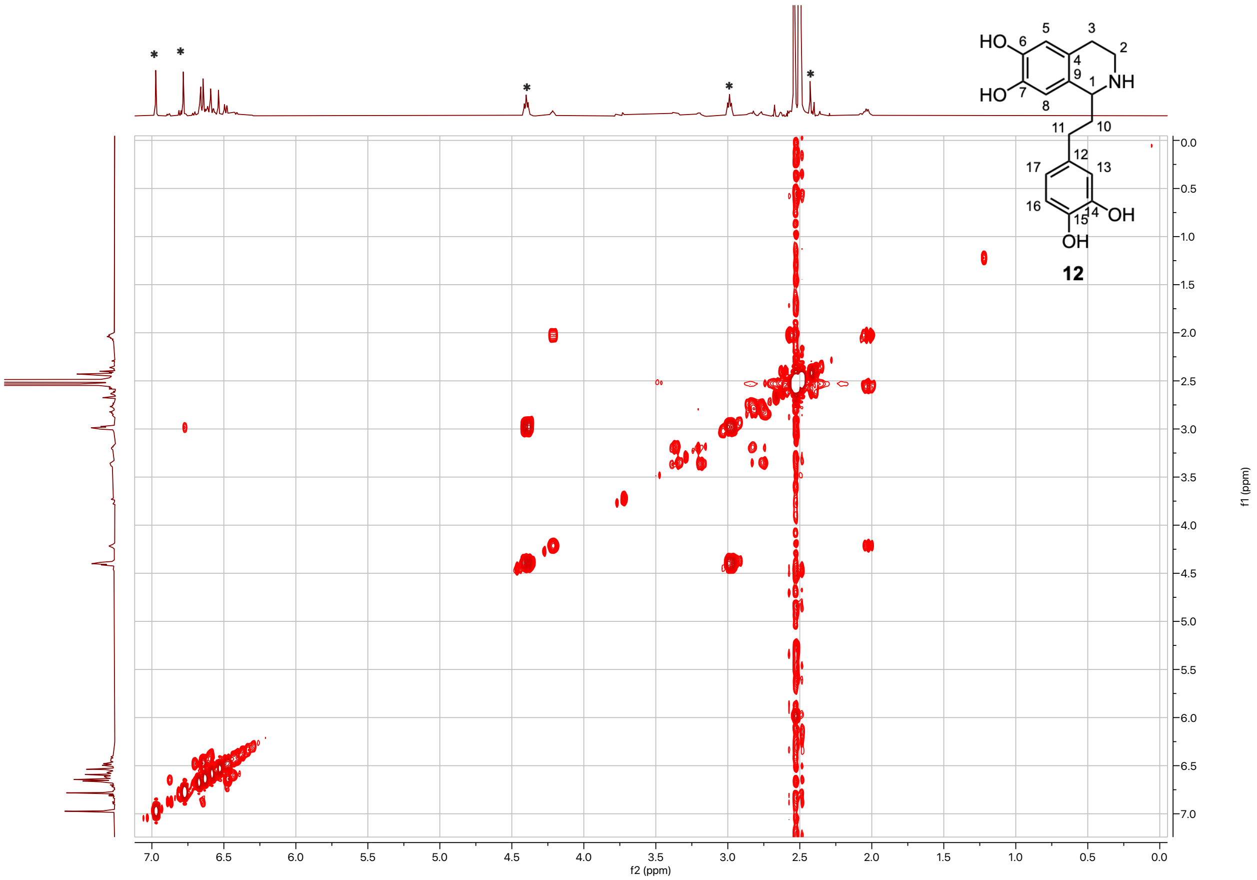


**Supplementary Figure. 50. COSY spectrum of 1-(3,4-dihydroxyphenethyl)-1,2,3,4-tetrahydroisoquinoline-6,7-diol (12) (DMSO-d$_6$, 500 MHz, 298 K).** Asterisk indicates impurities. The sharp intense peak at 2.54ppm corresponds to protonated DMSO.

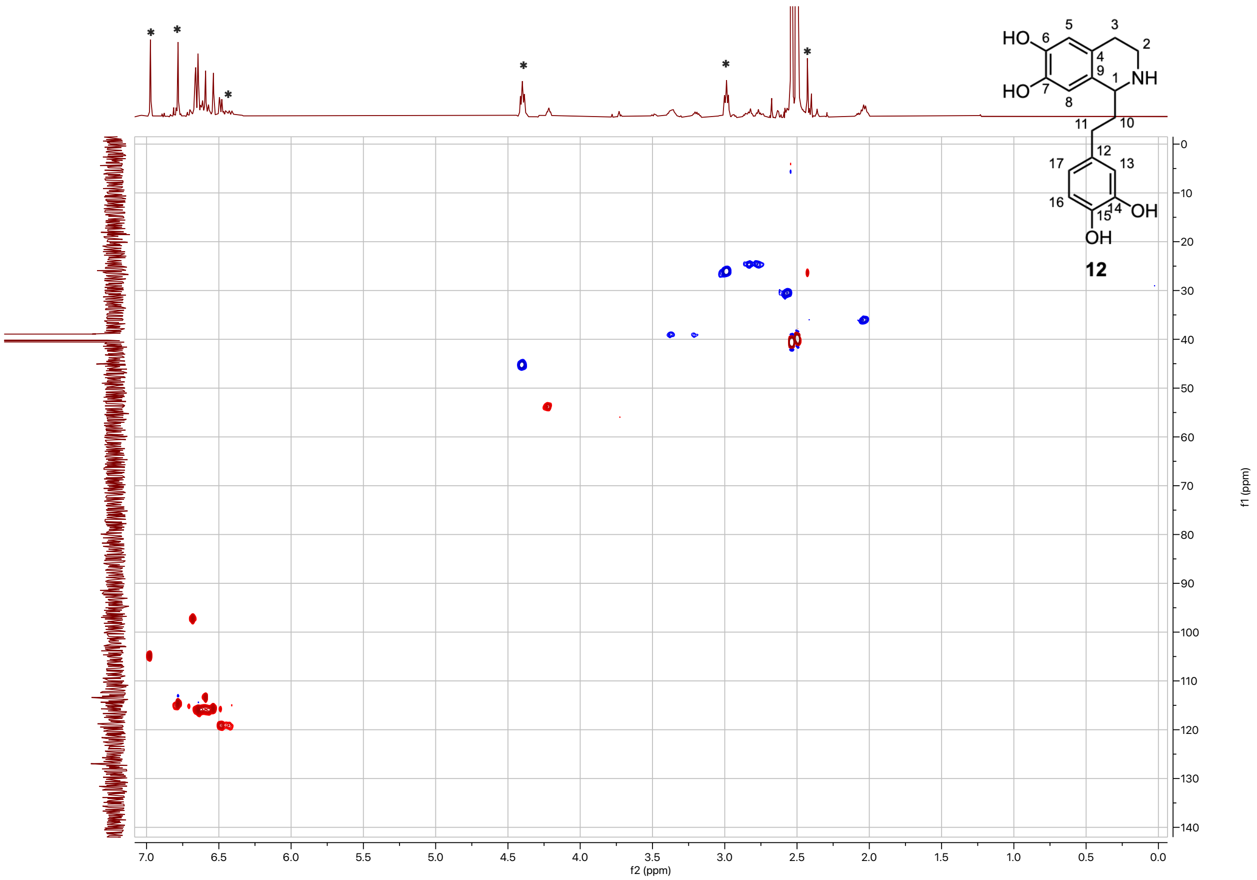


**Supplementary Figure. 51. HSQC spectrum of 1-(3,4-dihydroxyphenethyl)-1,2,3,4-tetrahydroisoquinoline-6,7-diol (12) (DMSO-d$_6$, 500 MHz, 298 K).** Asterisk indicates impurities.

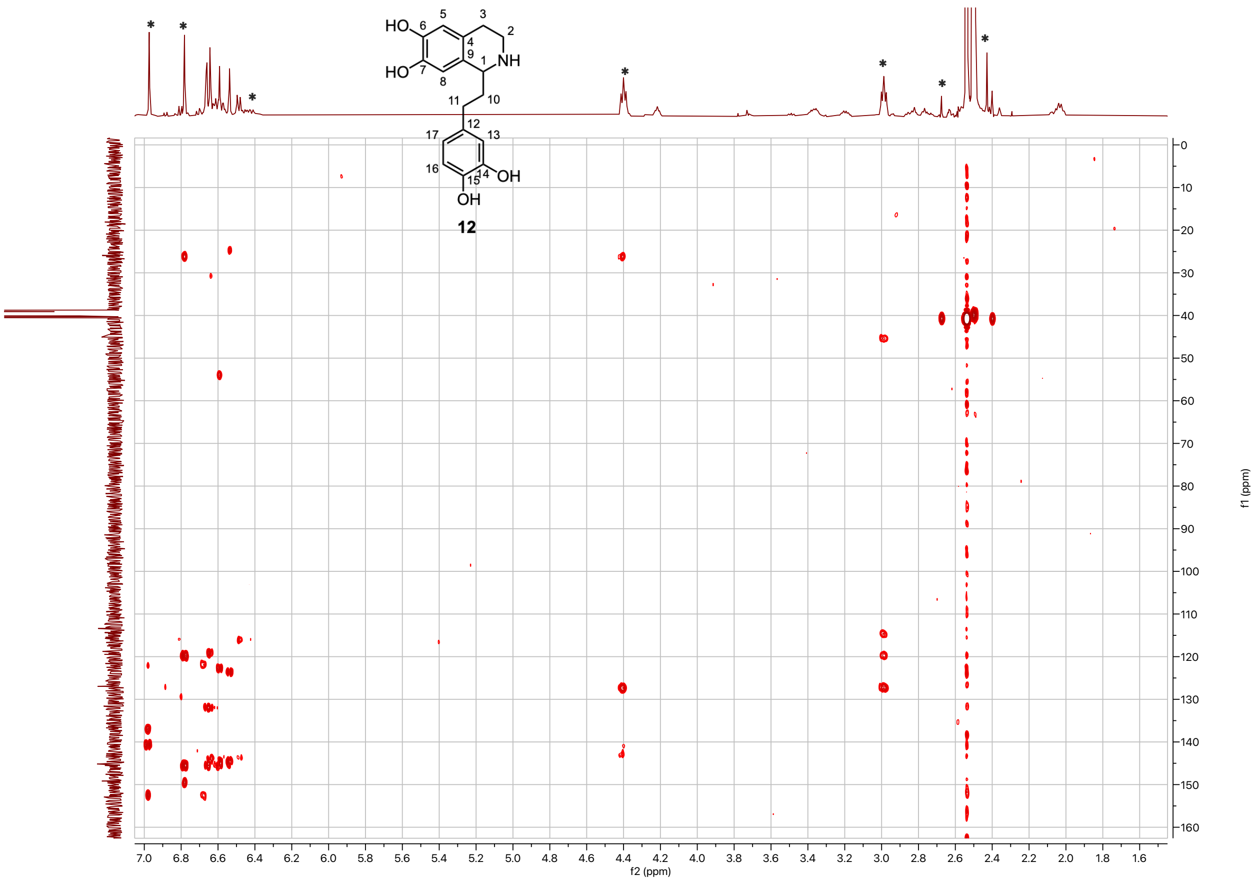


**Supplementary Figure. 52. HMBC spectrum of 1-(3,4-dihydroxyphenethyl)-1,2,3,4-tetrahydroisoquinoline-6,7-diol (12) (DMSO-$d_6$, 500 MHz, 298 K).** Asterisk indicates impurities.

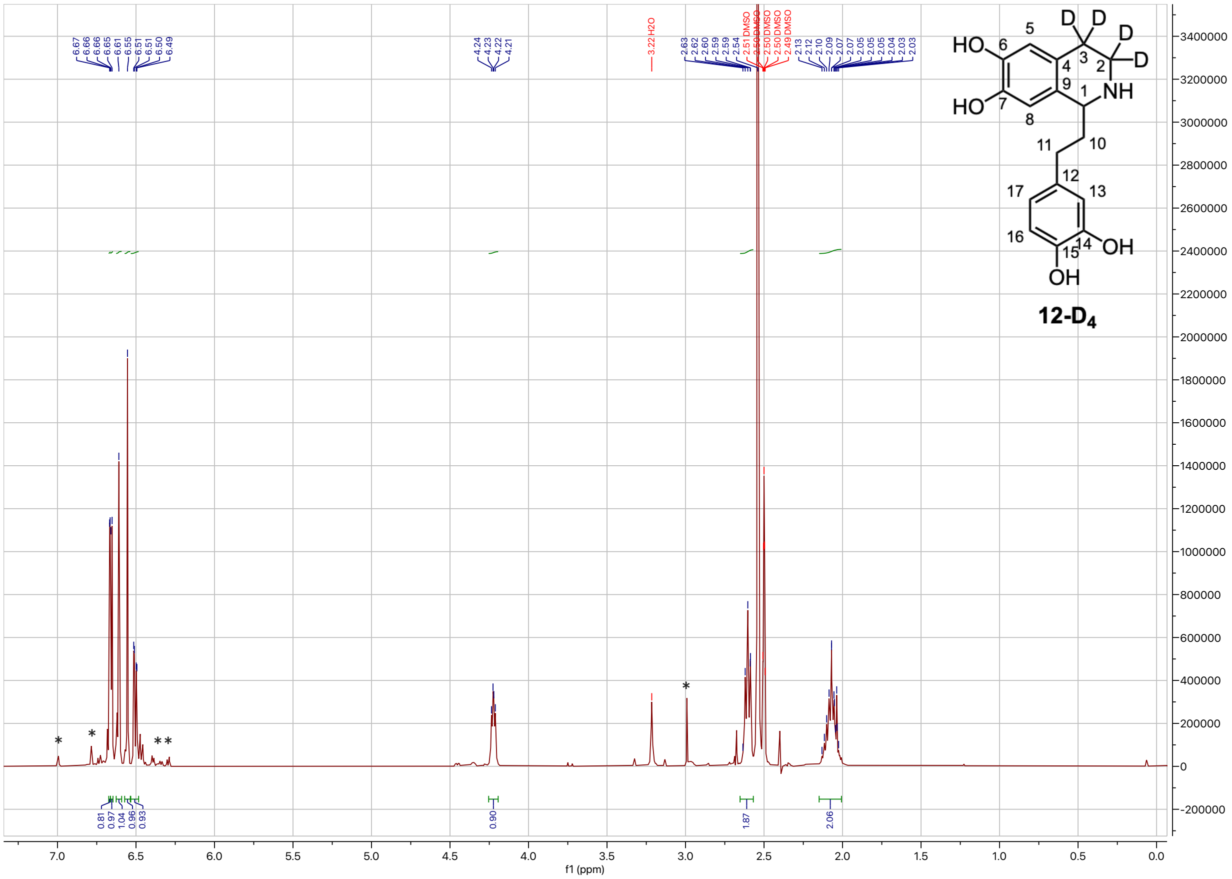

**Supplementary Figure. 53. ¹H-NMR spectrum of 1-(3,4-dihydroxyphenethyl)-1,2,3,4-tetrahydroisoquinoline-3,3,4,4-$d_4$-6,7-diol (12-$D_4$) (DMSO-$d_6$, 500 MHz, 298 K).** ¹H NMR (500 MHz, DMSO) δ 6.67 (d, $J$ = 1.7 Hz, 1H), 6.66 (d, $J$ = 4.0 Hz, 1H), 6.61 (s, 1H), 6.55 (s, 1H), 6.50 (dd, $J$ = 8.0, 2.1 Hz, 1H), 4.22 (dd, $J$ = 7.4, 4.9 Hz, 1H), 2.60 (t, $J$ = 8.1 Hz, 2H), 2.15 – 2.01 (m, 2H). Asterisk indicates impurities. The sharp intense peak at 2.54ppm corresponds to protonated DMSO.

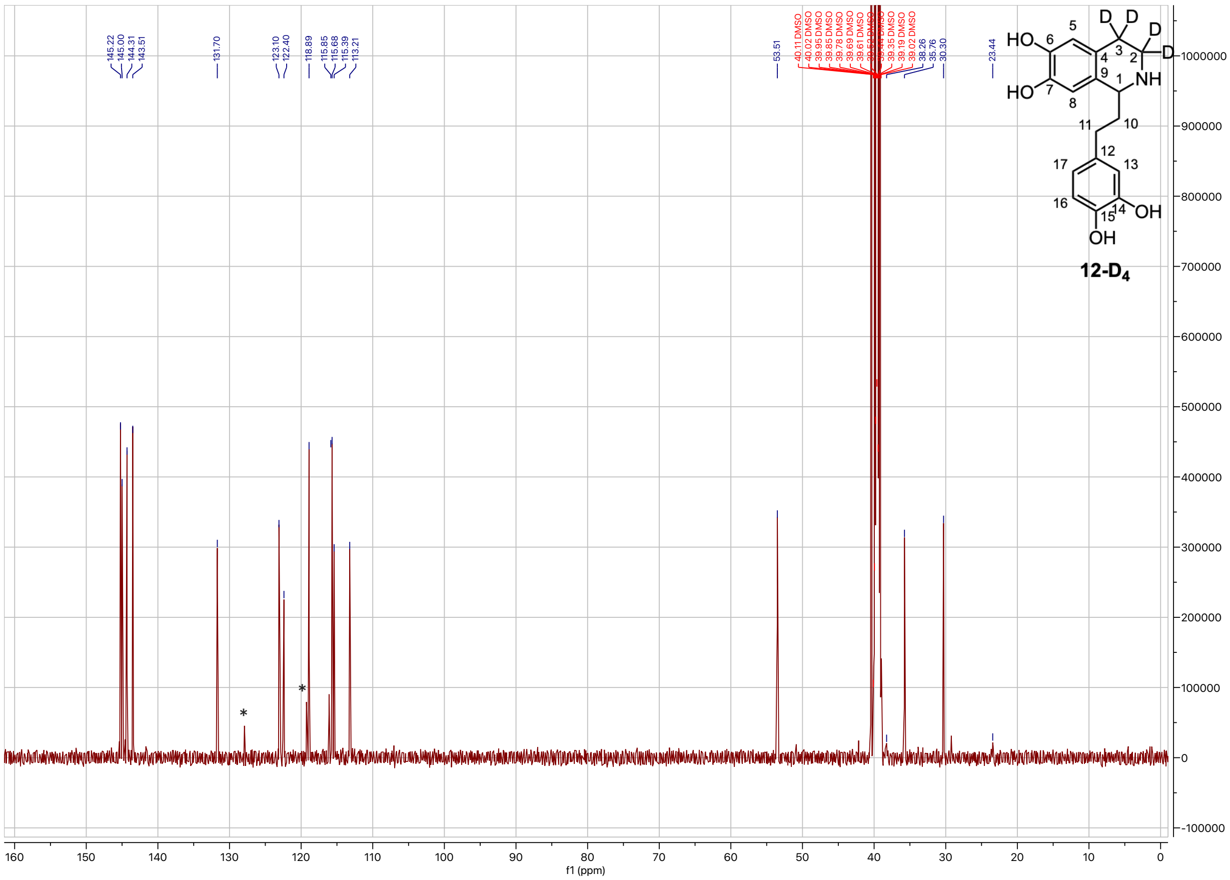

**Supplementary Figure. 54. $^{13}$C-NMR spectrum of 1-(3,4-dihydroxyphenethyl)-1,2,3,4-tetrahydroisoquinoline-3,3,4,4-*d*$_4$-6,7-diol (12-D$_4$) (DMSO-d$_6$, 500 MHz, 298 K).** Asterisk indicates co-eluting impurities.

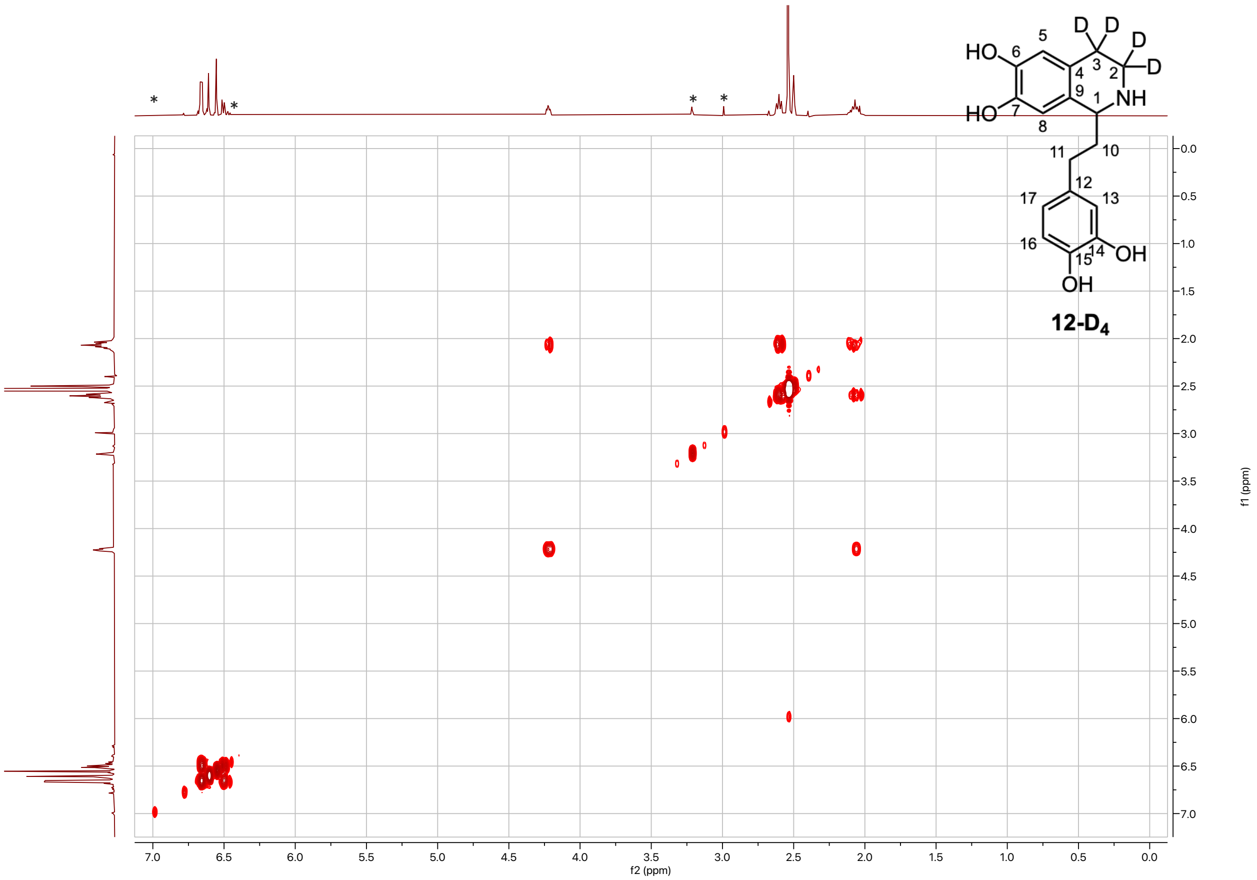


**Supplementary Figure. 55. COSY spectrum of 1-(3,4-dihydroxyphenethyl)-1,2,3,4-tetrahydroisoquinoline-3,3,4,4-*d*$_4$-6,7-diol (12-D$_4$) (DMSO-d$_6$, 500 MHz, 298 K).** Asterisk indicates impurities. The sharp intense peak at 2.54ppm corresponds to protonated DMSO.

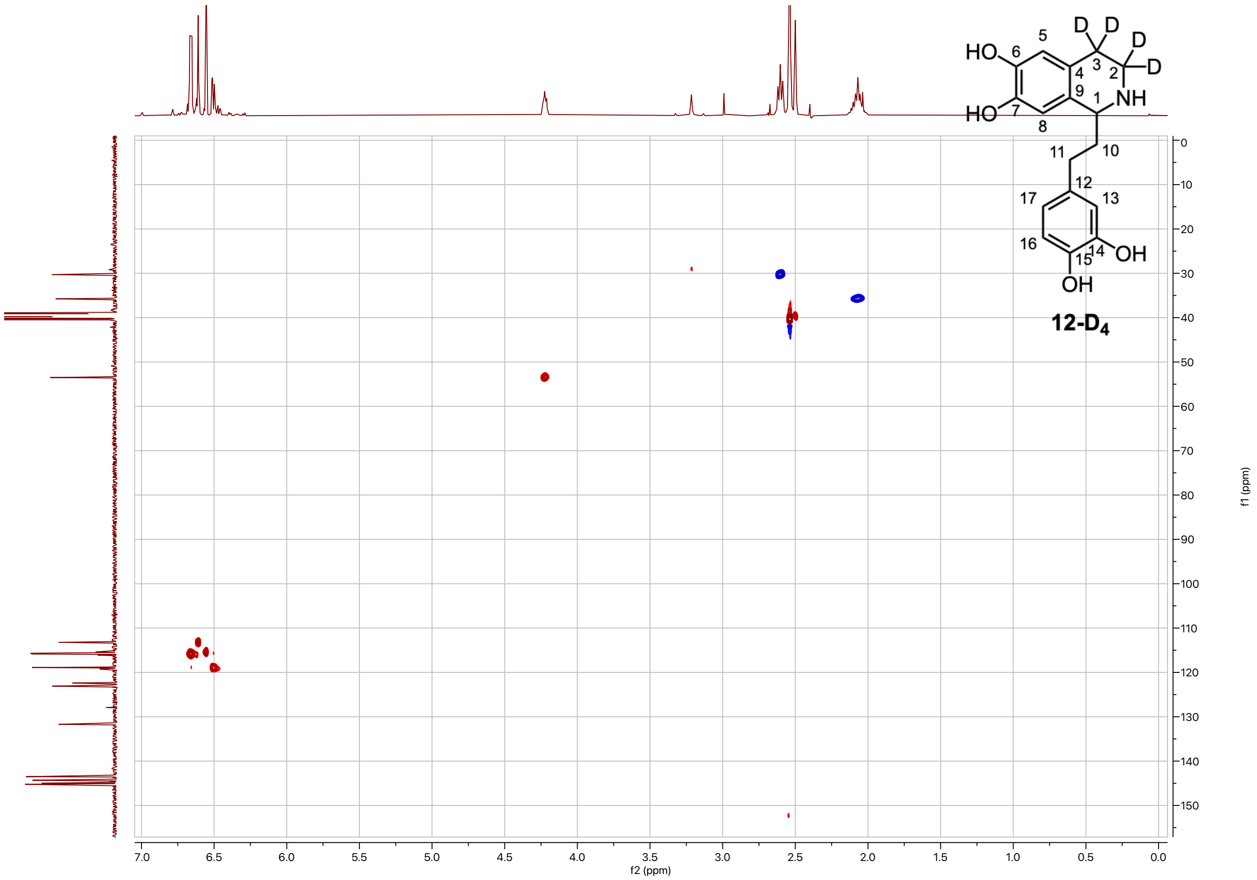


**Supplementary Figure. 56. HSQC spectrum of 1-(3,4-dihydroxyphenethyl)-1,2,3,4-tetrahydroisoquinoline-3,3,4,4-$d_4$-6,7-diol (12-D$_4$) (DMSO-d$_6$, 500 MHz, 298 K).**

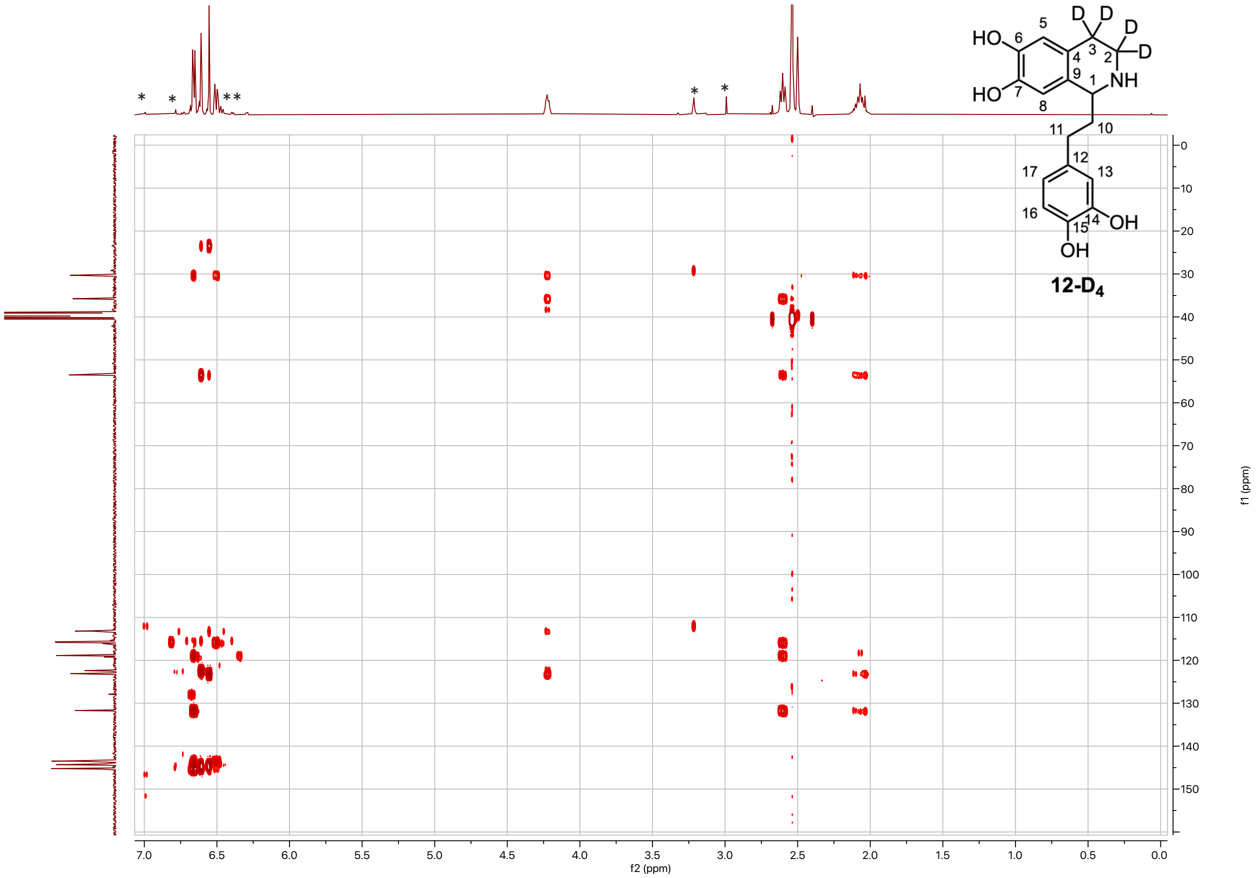


**Supplementary Figure. 57. HMBC spectrum of 1-(3,4-dihydroxyphenethyl)-1,2,3,4-tetrahydroisoquinoline-3,3,4,4-$d_4$-6,7-diol (12-D$_4$) (DMSO-d$_6$, 500 MHz, 298 K).** Asterisk indicates impurities. The sharp intense peak at 2.54ppm corresponds to protonated DMSO.

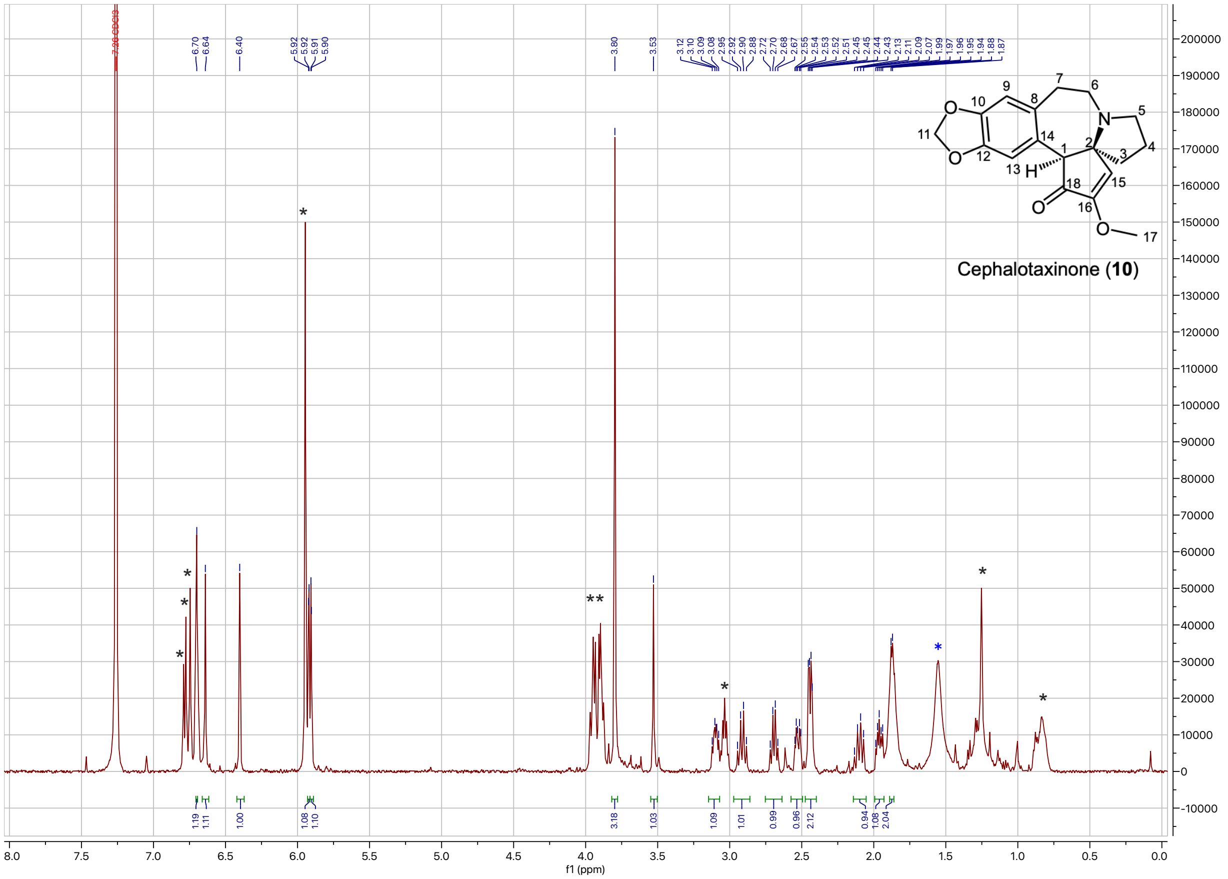

**Supplementary Figure. 58. $^1$H-NMR spectrum of cephalotaxinone (10) ($CDCl_3$, 500 MHz, 298 K).** $^1$H NMR (500 MHz, $CDCl_3$) δ 6.70 (s, 1H), 6.64 (s, 1H), 6.40 (s, 1H), 5.92 (d, $J$ = 1.5 Hz, 1H), 5.91 (d, $J$ = 1.5 Hz, 1H), 3.80 (s, 3H), 3.53 (s, 1H), 3.15 – 3.07 (m, 1H), 2.91 (q, $J$ = 10.3 Hz, 1H), 2.69 (q, $J$ = 8.8 Hz, 1H), 2.57 – 2.49 (m, 1H), 2.47 – 2.41 (m, 2H), 2.14 – 2.05 (m, 1H), 1.99 – 1.93 (m, 1H), 1.89 – 1.85 (m, 2 H). Black asterisk indicates impurities, most likely from degradation of the molecule. The broad peak at 1.55 ppm corresponds to water (blue asterisk).

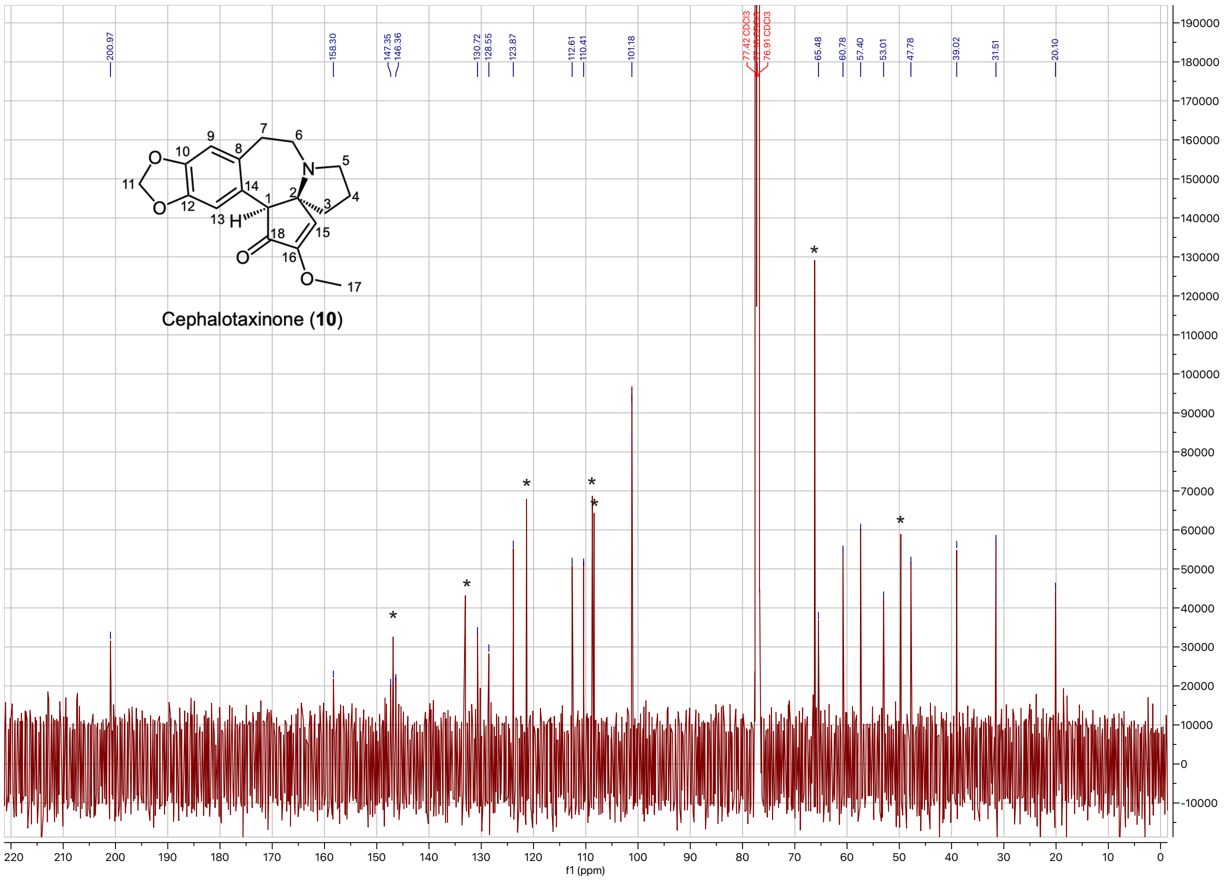


**Supplementary Figure. 59. $^{13}$C-NMR spectrum of cephalotaxinone (10) ($CDCl_3$, 500 MHz, 298 K).** Black asterisk indicates impurities, most likely from degradation of the molecule.

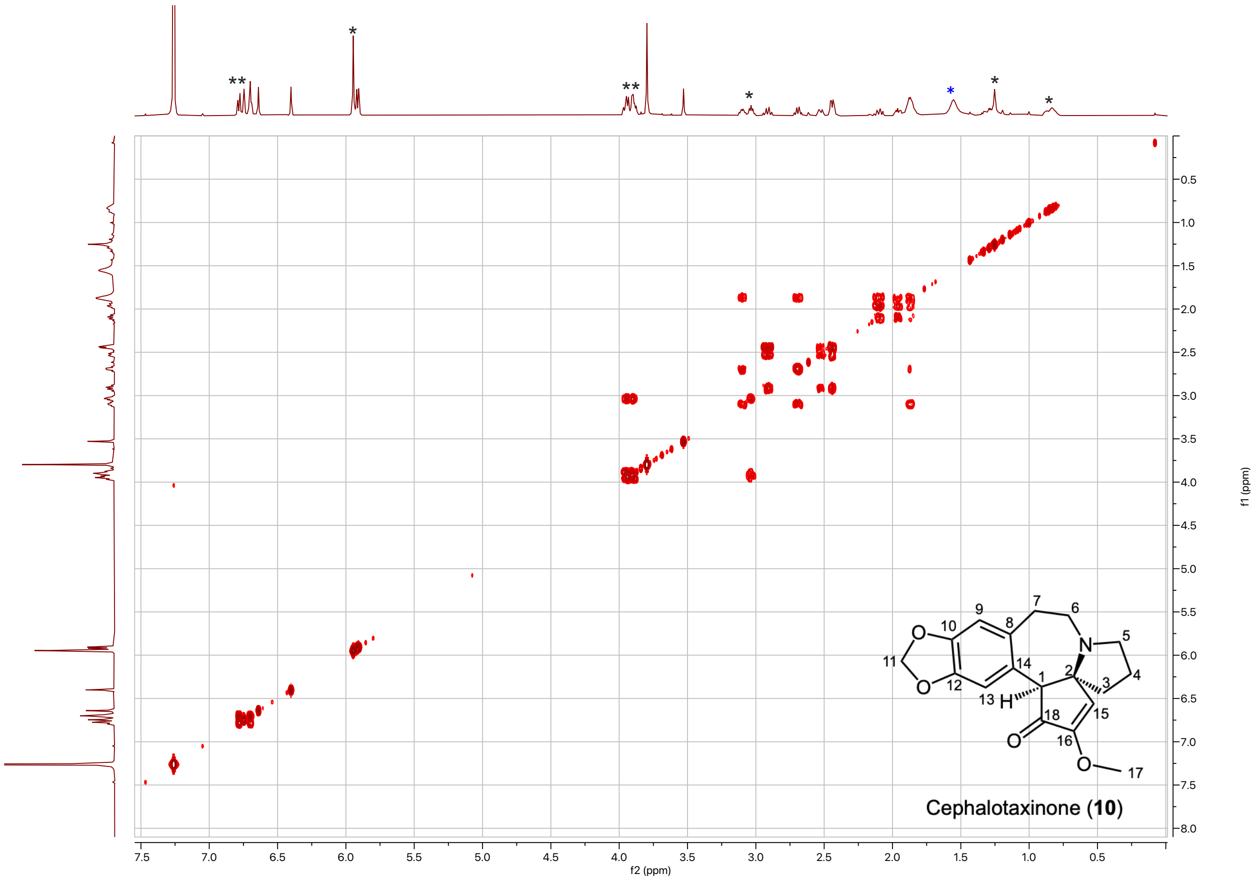


**Supplementary Figure. 60. COSY spectrum of cephalotaxinone (10) ($CDCl_3$, 500 MHz, 298 K).** Black asterisk indicates impurities, most likely from degradation of the molecule. The broad peak at 1.55 ppm corresponds to water (blue asterisk).

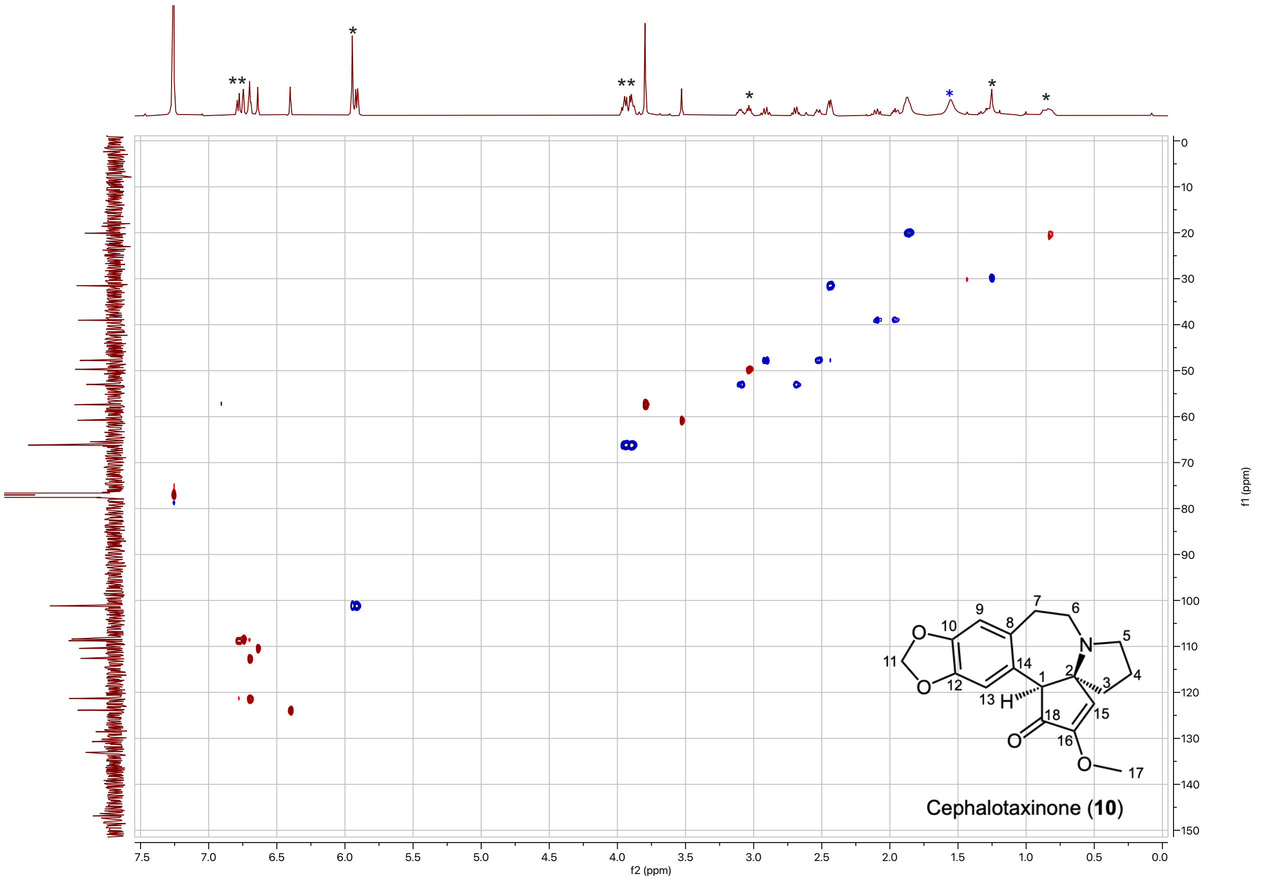


**Supplementary Figure. 61. HSQC spectrum of cephalotaxinone (10) ($CDCl_3$, 500 MHz, 298 K).** Black asterisk indicates impurities, most likely from degradation of the molecule. The broad peak at 1.55 ppm corresponds to water (blue asterisk).

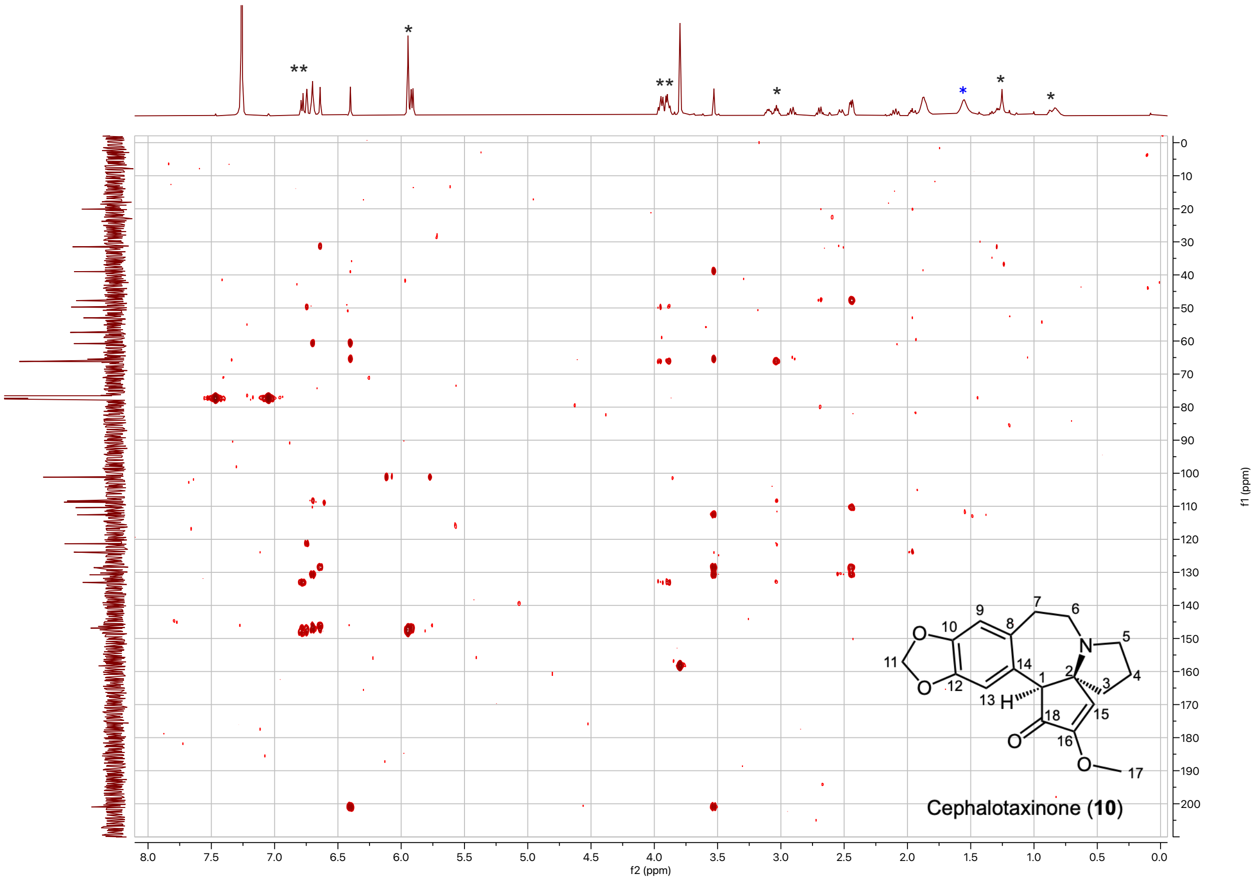


**Supplementary Figure. 62. HMBC spectrum of cephalotaxinone (10) ($CDCl_3$, 500 MHz, 298 K).** Black asterisk indicates impurities, most likely from degradation of the molecule. The broad peak at 1.55 ppm corresponds to water (blue asterisk).

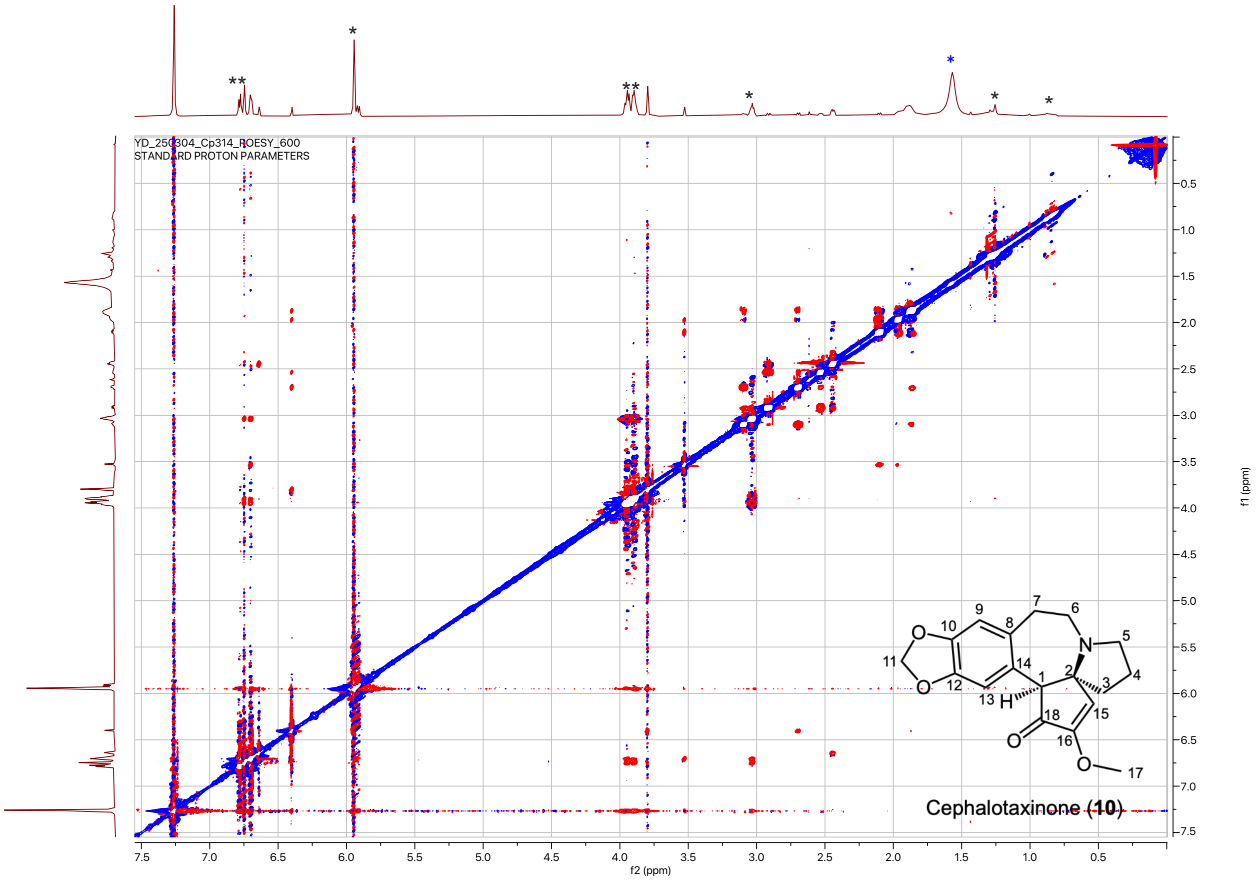


**Supplementary Figure. 63. ROESY spectrum of cephalotaxinone (10) ($CDCl_3$, 600 MHz, 298 K).** Asterisk indicates impurities. Black asterisk indicates impurities, most likely from degradation of the molecule. The broad peak at 1.55 ppm corresponds to water (blue asterisk).

#### SUPPLEMENTARY TABLES

**Supplementary Table 1.** Summary of natural accumulation and isotope-labeled precursor feeding experiments of different phenethylisoquinolines.

| Chemical structure | 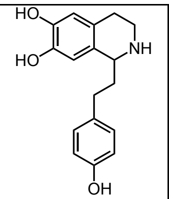 | 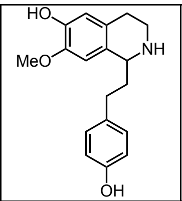 | 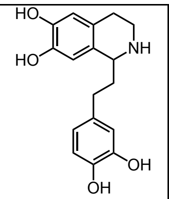 | 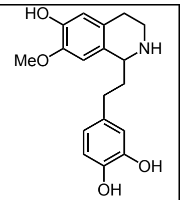 | 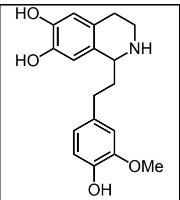 | 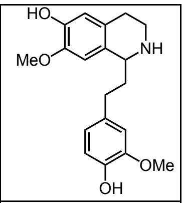 |
|---|---|---|---|---|---|---|
| Compound number | **1** | **11** | **12** | **13** | **2** | **3** |
| Metabolite detection in any tissues of *C. harringtonia* | Yes, in root tips | Yes, in root tips | None | None | Yes, in root tips | Yes, in root tips |
| Enrichment of its four-deuterated form from dopamine-D4 feeding assay | None | None | None | None | Yes, in root tips | Yes, in root tips |
| CET-D4 enrichment from feeding its four deuterated form | None | None | None | None | Yes, in root tips | Yes, in root tips |

**Supplementary Table 2.** Summary of tissue-specific detection of deuterium-labeled *Cephalotaxus* alkaloids using chopped-tissue isotope-labeled precursor feeding and excised aerial branch $D_2O$ labeling assays

| Compound | Isotope-labeled precursor feeding assay (chopped-tissue plate assay – 5 days) * Includes data for roots | $D_2O$ labeling assay (excised aerial branch assay – 5, 10, 15 days) * Includes no data for roots |
|---|---|---|
| Cephalotaxinone | Root tips only | Not observed |
| CET | Root tips only | Young and old needles (all 5, 10, 15 days) |
| HHT | Not observed | Young needles (10 and 15 days) |

**Supplementary Table 3.** Seasonal feed-in Experiments with *C. harringtonia* (var. Fastigiata).

| **Exp. #** | **Time Experiment Conducted** | **Substrates** | **Enrichment of CET-D4** |
|---|---|---|---|
| 1 | May 2022 | dopamine-D4 | Detected |
| 2 | Nov. 2022 | dopamine-D4<br>**1**-D4 | Not detected |
| 3 | May 2023 | dopamine-D4<br>**1**-D4<br>**2**-D4 | Detected<br>(only dopamine-D4 & compound **2**-D4) |
| 4 | Dec. 2023 | dopamine-D4<br>**1**-D4<br>**2**-D4 | Not detected |
| 5 | June 2024 | **3**-D4<br>**12**-D4 | Detected<br>(only compound **3**-D4) |

**Supplementary Table 4.** Summary of heterologous host systems used for expression of CET biosynthetic enzymes from *C. harringtonia*.

| Enzyme | *N. benthamiana* | *S. cerevisiae* | *E. coli* |
|---|---|---|---|
| *Ch*OMT-1 | Yes | No | Yes |
| *Ch*CYP805A11 | Yes | Yes | No |
| *Ch*NmrA-1 | Yes | No | Yes |
| *Ch*CYP805A12 | Yes | No | No |
| *Ch*2OGD-1 | Yes | No | No |
| *Ch*CYP805A13 | Yes | No | No |

*Ch*OMT-1 was expressed in *E. coli* to prepare compound **3**-D4 from **2**-D4 for stable-isotope labeled precursor feeding experiments.

**Supplementary Table 5.** Sequence comparison between *Ch*CYP805A13 and *Ch*CYP805A14.

| Gene | Transcript ID | Sequence |
|---|---|---|
| CYP805A13 | YD-1-Rt_transcript/50321.p1 | MENFVVNYIIEAFQHYFFDMIPIFGGASVVLFSFVLTVVAAHYLLSDKK<br>**KNDEKRMS**WPPSPP**TM**PLLGNLHMLNKGGNFLMAAYDIAKSYGPVM<br>TLWMGTSPAVVLTGQTAIWEALVNQASNFADRPFMNTNRFFSSGDITT<br>MTSDCNENWMKLRKILHNNVISRFNIASHSSSHHRNVNGLIKKLVEEMN<br>ANNGVVRPFHSFKIM**G**ISFMAKLCFGPDFEDEIFAIK**V**ENLIAEDIALIGK<br>GETLLESIPLARYWNPSTF**L**TKKNQKAISASILELLLPIVKYGRSYRAYLK<br>QSAPNSYLNCLLSISEEEDDELKLKLSDEEIAFNIFELLILSVDSTSTALEW<br>AVAYLINNPHIQNKVYQEVNEALPGGKERLLRVEDLEKLPYLQAVVKE<br>TLRKEAVAPFALPHQTANECKVMGVNIPAKATVLFNLFSVNNDPQLWS<br>NPDEFAPERFLGNNVDVRSCYLPFGAGRRICPGMDMAYIHVP**V**TLGSL<br>**M**KYFEWGCVKEGSPPDLSREERSMIMYMKHPL**E**ARITYRT |
| CYP805A14 | YD-3-Rt_transcript/38063.p1 | **MMRKGISL**WPPSPP**RL**PLLGNLHMLNKGGNFLMAAYDIAKSYGPVMT<br>LWMGTSPAVVLTGQTAIWEALVNQASNFADRPFMNTNRFFSSGDITTM<br>TSDCNENWMKLRKILHNNVISRFNIASHSSSHHRNVNGLIKKLVEEMNA<br>NNGVVRPFHSFKIM**A**ISFMAKLCFGPDFEDEIFAIK**A**ENLIAEDIALIGKG<br>ETLLESIPLARYWNPSTF**I**TKKNQKAISASILELLLPIVKYGRSYRAYLKQ<br>SAPNSYLNCLLSISEEEDDELKLKLSDEEIAFNIFELLILSVDSTSTALEWA<br>VAYLINNPHIQNKVYQEVNEALPGGKERLLRVEDLEKLPYLQAVVKET<br>LRKEAVAPFALPHQTANECKVMGVNIPAKATVLFNLFSVNNDPQLWSN<br>PDEFAPERFLGNNVDVRSCYLPFGAGRRICPGMDMAYIHVP**I**TLGSL**I**KY<br>FEWGCVKEGSPPDLSREERSMIMYMKHPL**Q**ARITYRT |

The 49 amino acid residues that correspond to a N-terminal transmembrane domain of a cytochrome P450 are in blue. The 16 amino acid residues that differ between *Ch*CYP805A13 and *Ch*CYP805A14 are in bold.

**Supplementary Table 6.** $^{13}$C and $^{1}$H δ assignments as well as 2D-NMR correlations of 1-(4-hydroxy-3-methoxyphenethyl)-1,2,3,4-tetrahydroisoquinoline-6,7-diol (**2**) recorded in DMSO-d6.

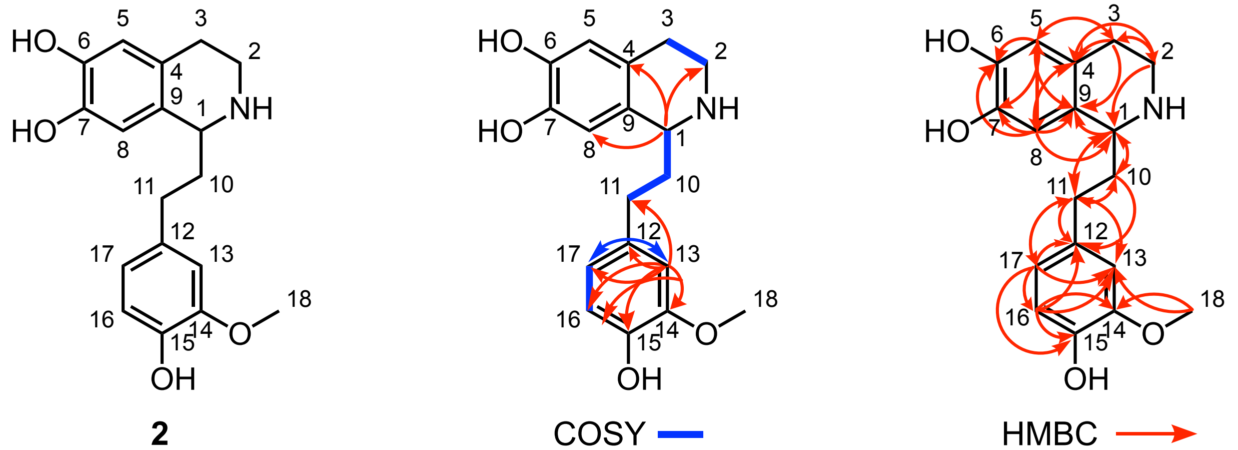


| C-# | δ $^{13}$C (ppm) | δ $^{1}$H (mult.; *J* in Hz) | COSY | HMBC |
|---|---|---|---|---|
| 1 | 53.68 | 3.96 (dd; 8.3, 4.2) | 10 | 2, 4, 8, 9, 10, 11 |
| 2a | 39.48 | 2.98 (ddd; 12.5, 7.4, 5.2) | 2b, 3a, 3b | 1, 3, 4 |
| 2b | 39.48 | 3.20 (m) | 2a, 3a, 3b | 1, 3, 4 |
| 3a | 26.48 | 2.61 (m) | 2a, 2b, 3b | 2, 4, 5, 9 |
| 3b | 26.48 | 2.68 (m) | 2a, 2b, 3a | 2, 4, 5, 9 |
| 4 | 123.98 | -- | -- | -- |
| 5 | 115.43 | 6.46 (s) | -- | 1, 3, 6, 7, 8, 9 |
| 6 | 143.81 | -- | -- | -- |
| 7 | 144.19 | -- | -- | -- |
| 8 | 113.13 | 6.53 (s) | -- | 1, 3, 4, 5, 6, 7, 9 |
| 9 | 126.55 | -- | -- | -- |
| 10 | 37.03 | 1.96 (m; 2H) | 1, 11 | 1, 9, 11, 12 |
| 11 | 30.92 | 2.61 (m; 2H) | 10 | 1, 10, 12, 13, 17 |
| 12 | 132.50 | -- | -- | -- |
| 13 | 112.46 | 6.79 (d; 1.9) | 17 | 11, 12, 14, 15, 16, 17 |
| 14 | 147.47 | -- | -- | -- |
| 15 | 144.55 | -- | -- | -- |
| 16 | 115.36 | 6.68 (d; 8.0) | 17 | 11, 12, 13, 14, 15 |
| 17 | 120.31 | 6.62 (dd; 8.0, 1.9) | 13, 16 | 11, 12, 13, 14, 15, 16 |
| 18 | 55.54 | 3.75 (s) | -- | 13, 14 |

NMR spectra are shown in **Supplementary Figs. 13-17**. s = singlet, d = doublet, dd = doublet of doublets, ddd = doublet of doublets of doublets, m = multiplet.

**Supplementary Table 7.** $^{13}C$ and $^{1}H$ δ assignments as well as 2D-NMR correlations of 1-(4-hydroxy-3-methoxyphenethyl)-1,2,3,4-tetrahydroisoquinoline-3,3,4,4-$d_4$-6,7-diol (**2-D₄**) recorded in DMSO-d₆.

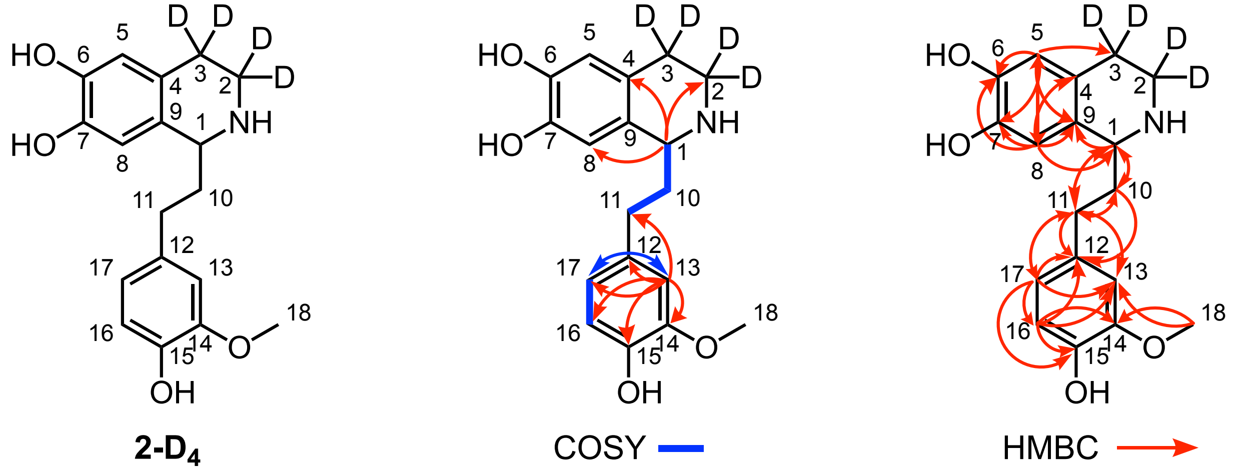


| C-# | δ $^{13}C$ (ppm) | δ $^{1}H$ (mult.; *J* in Hz) | COSY | HMBC |
|---|---|---|---|---|
| 1 | 53.59 | 3.98 (dd; 8.0, 4.5) | 10 | 2, 4, 8, 9, 10, 11 |
| 2 | 39.48 | -- | -- | -- |
| 3 | 25.26 | -- | -- | -- |
| 4 | 123.69 | -- | -- | -- |
| 5 | 115.45 | 6.47 (s) | -- | 1, 3, 6, 7, 8, 9 |
| 6 | 143.89 | -- | -- | -- |
| 7 | 144.30 | -- | -- | -- |
| 8 | 113.16 | 6.54 (s) | -- | 1, 3, 4, 5, 6, 7, 9 |
| 9 | 126.22 | -- | -- | -- |
| 10 | 36.90 | 1.97 (m; 2H) | 1, 11 | 1, 9, 11, 12 |
| 11 | 30.89 | 2.61 (m; 2H) | 10 | 1, 10, 12, 13, 17 |
| 12 | 132.41 | -- | -- | -- |
| 13 | 112.47 | 6.79 (d; 1.9) | 17 | 11, 12, 14, 15, 16, 17 |
| 14 | 147.48 | -- | -- | -- |
| 15 | 144.59 | -- | -- | -- |
| 16 | 115.38 | 6.68 (d; 8.0) | 17 | 11, 12, 13, 14, 15 |
| 17 | 120.33 | 6.62 (dd; 7.9, 2.0) | 13, 16 | 11, 12, 13, 14, 15, 16 |
| 18 | 55.55 | 3.75 (s) | -- | 13, 14 |

NMR spectra are shown in **Supplementary Figs. 18-22**. s = singlet, d = doublet, dd = doublet of doublets, m = multiplet.

**Supplementary Table 8.** $^{13}C$ and $^{1}H$ δ assignments as well as 2D-NMR correlations of 1-(4-hydroxy-3-methoxyphenethyl)-7-methoxy-1,2,3,4-tetrahydroisoquinolin-6-ol (**3**) recorded in DMSO-d₆.

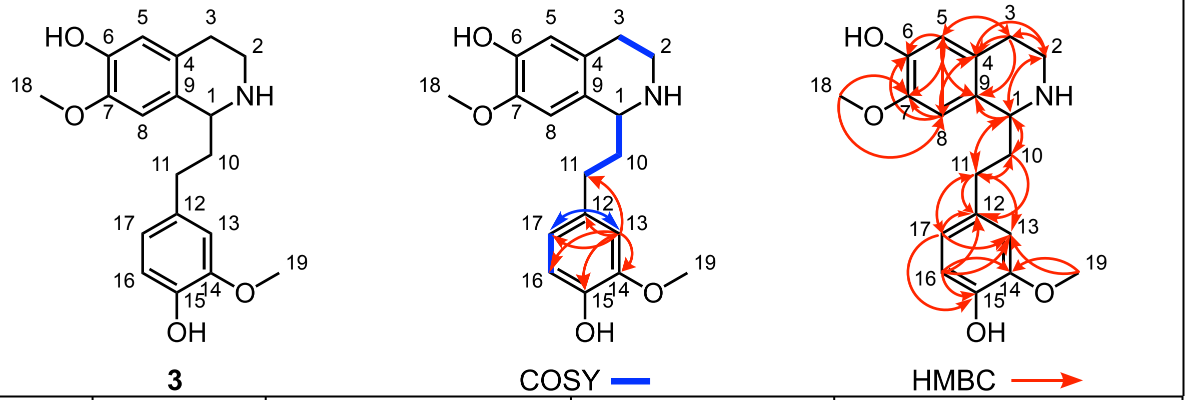


| C-# | δ $^{13}C$ (ppm) | δ $^{1}H$ (mult.; *J* in Hz) | COSY | HMBC |
|---|---|---|---|---|
| 1 | 53.95 | 4.05 (dd; 8.6, 4.3) | 10a, 10b | 2, 9, 10, 11 |
| 2a | 39.36 | 2.99 (dt; 11.5, 4.0) | 2b, 3a, 3b | 1, 3, 4 |
| 2b | 39.36 | 3.23 (m) | 2a, 3a, 3b | 1, 3, 4 |
| 3a | 26.40 | 2.65 (m) | 2a, 2b, 3b | 2, 4, 5, 9 |
| 3b | 26.40 | 2.72 (m) | 2a, 2b, 3a | 2, 4, 5, 9 |
| 4 | 125.75 | -- | -- | -- |
| 5 | 115.36 | 6.52 (s) | -- | 1, 3, 6, 7, 8, 9 |
| 6 | 145.30 | -- | -- | -- |
| 7 | 146.27 | -- | -- | -- |
| 8 | 110.17 | 6.67 (s) | -- | 1, 3, 4, 5, 6, 7, 9 |
| 9 | 126.39 | -- | -- | -- |
| 10a | 36.90 | 2.00 (m) | 1, 10b, 11a, 11b | 1, 9, 11, 12 |
| 10b | 36.90 | 2.08 (m) | 1, 10a, 11a, 11b | 1, 9, 11, 12 |
| 11a | 31.06 | 2.60 (m) | 10a, 10b, 11b | 1, 10, 12, 13, 17 |
| 11b | 31.06 | 2.67 (m) | 10a, 10b, 11a | 1, 10, 12, 13, 17 |
| 12 | 132.49 | -- | -- | -- |
| 13 | 112.52 | 6.81 (d; 1.9) | 17 | 11, 12, 14, 15, 16, 17 |
| 14 | 147.47 | -- | -- | -- |
| 15 | 144.58 | -- | -- | -- |
| 16 | 115.33 | 6.68 (d; 8.0) | 17 | 11, 12, 13, 14, 15 |
| 17 | 120.39 | 6.63 (dd; 8.0, 2.0) | 13, 16 | 11, 12, 13, 14, 15 |
| 18 | 55.80 | 3.71 (s) | -- | 7, 8 |
| 19 | 55.54 | 3.75 (s) | -- | 13, 14 |

NMR spectra are shown in **Supplementary Figs. 23-27**. s = singlet, d = doublet, dd = doublet of doublets, dt = doublet of triplets, t = triplet, m = multiplet.

**Supplementary Table 9.** $^{13}$C and $^{1}$H δ assignments as well as 2D-NMR correlations of 1-(4-hydroxy-3-methoxyphenethyl)-7-methoxy-1,2,3,4-tetrahydroisoquinolin-3,3,4,4-$d_4$-6-ol (**3-D**$_4$) recorded in DMSO-d$_6$.

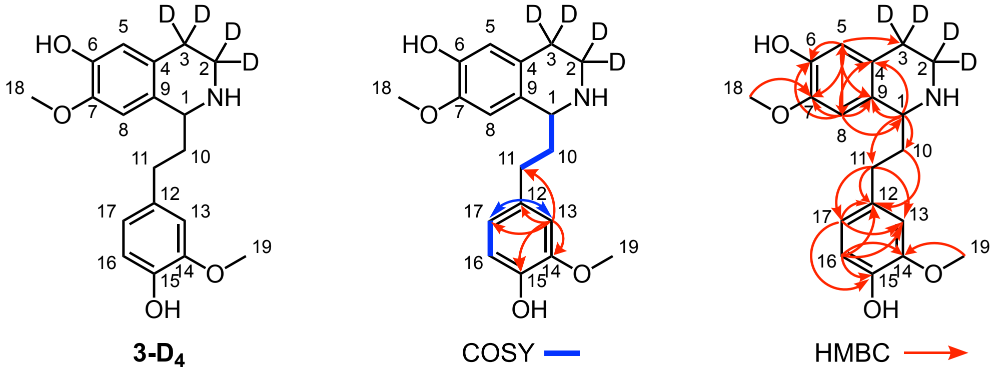


| C-# | δ $^{13}$C (ppm) | δ $^{1}$H (mult.; *J* in Hz) | COSY | HMBC |
|---|---|---|---|---|
| 1 | 54.12 | 3.83 (dd; 8.8, 3.3) | 10a, 10b | 4, 9, 10, 11 |
| 2 | (N.A.; overlapping with DMSO) | -- | -- | -- |
| 3 | 27.00 | -- | -- | -- |
| 4 | 126.78 | -- | -- | -- |
| 5 | 115.46 | 6.46 (s) | -- | 1, 3, 6, 7, 8, 9 |
| 6 | 144.70 | -- | -- | -- |
| 7 | 145.94 | -- | -- | -- |
| 8 | 110.09 | 6.62 (s) | -- | 1, 3, 4, 5, 6, 7, 9 |
| 9 | 129.03 | -- | -- | -- |
| 10a | 37.73 | 1.87 (m) | 1, 10b, 11a, 11b | 12 |
| 10b | 37.73 | 2.01 (m) | 10a, 11a, 11b | 12 |
| 11a | 31.34 | 2.56 (m) | 10a, 10b, 11b | 12, 13, 17 |
| 11b | 31.34 | 2.64 (m) | 10a, 10b, 11a | 12, 13, 17 |
| 12 | 133.11 | -- | -- | -- |
| 13 | 112.51 | 6.79 (d; 2.1) | 17 | 11, 12, 14, 15, 17 |
| 14 | 147.41 | -- | -- | -- |
| 15 | 144.41 | -- | -- | -- |
| 16 | 115.31 | 6.67 (d; 7.9) | 17 | 12, 13, 14, 15 |
| 17 | 120.35 | 6.60 (dd; 8.0, 1.9) | 13, 16 | 13, 14, 15 |
| 18 | 55.76 | 3.70 (s) | -- | 7 |
| 19 | 55.51 | 3.74 (s) | -- | 14 |

NMR spectra are shown in **Supplementary Figs. 28-32**. N.A. not assigned (due to weak signal intensity or overlapping signals). s = singlet, d = doublet, dd = doublet of doublets, m = multiplet.

**Supplementary Table 10.** $^{13}$C and $^{1}$H δ assignments as well as 2D-NMR correlations of 1-(4-hydroxyphenethyl)-1,2,3,4-tetrahydroisoquinoline-6,7-diol (**1**) recorded in DMSO-$d_6$.

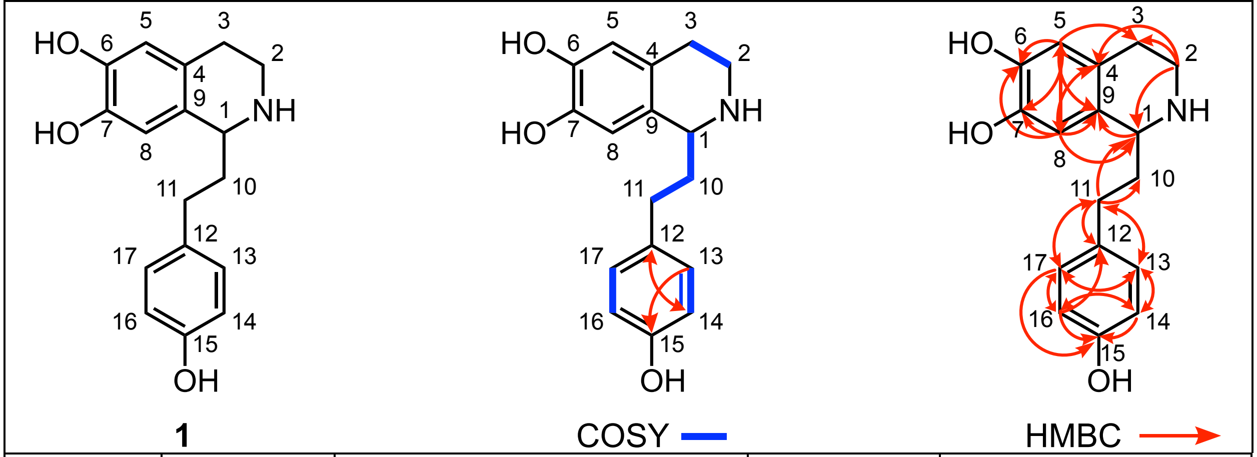


| C-# | δ $^{13}$C (ppm) | δ $^{1}$H (mult.; *J* in Hz) | COSY | HMBC |
|---|---|---|---|---|
| 1 | 53.78 | 3.86 (m) | 10 | -- |
| 2a | 39.88 | 2.90 (dt; 12.8, 6.4) | 2b, 3a, 3b | 1, 3, 4 |
| 2b | 39.88 | 3.14 (dt; 11.7, 5.6) | 2a, 3a, 3b | 1, 3, 4 |
| 3a | 27.19 | 2.56 (m) | 2a, 2b, 3b | -- |
| 3b | 27.19 | 2.62 (m) | 2a, 2b, 3a | -- |
| 4 | 124.50 | -- | -- | -- |
| 5 | 115.46 | 6.44 (s) | -- | 1, 3, 6, 7, 8, 9 |
| 6 | 143.59 | -- | -- | -- |
| 7 | 143.88 | -- | -- | -- |
| 8 | 113.00 | 6.50 (s) | -- | 1, 3, 4, 5, 6, 7 |
| 9 | 127.71 | -- | -- | -- |
| 10 | 37.47 | 1.89 (m; 2H) | 1, 11 | -- |
| 11 | 30.55 | 2.61 (m; 2H) | 10 | 1, 10, 12, 13, 17 |
| 12 | 132.00 | -- | -- | -- |
| 13 | 129.10 | 7.02 (m) | 14 | 11, 14, 15, 17 |
| 14 | 115.11 | 6.67 (m) | 13 | 12, 13, 15, 16 |
| 15 | 155.34 | -- | -- | -- |
| 16 | 115.11 | 6.67 (m) | 17 | 12, 14, 15, 17 |
| 17 | 129.10 | 7.02 (m) | 16 | 11, 13, 15, 16 |

NMR spectra are shown in **Supplementary Figs. 33-37**. s = singlet, d = doublet, dd = doublet of doublets, dt = doublet of triplets, t = triplet, m = multiplet.

**Supplementary Table 11.** $^{13}C$ and $^{1}H$ δ assignments as well as 2D-NMR correlations of 1-(4-hydroxyphenethyl)-1,2,3,4-tetrahydroisoquinoline-3,3,4,4-$d_4$-6,7-diol (**1-D₄**) recorded in DMSO-$d_6$.

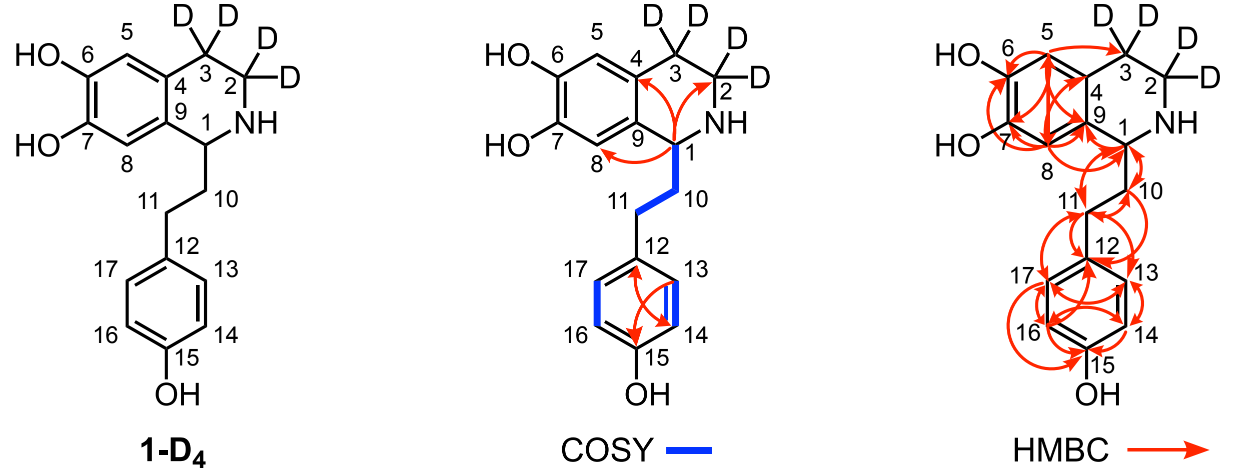


| C-# | δ $^{13}C$ (ppm) | δ $^{1}H$ (mult.; *J* in Hz) | COSY | HMBC |
|---|---|---|---|---|
| 1 | 53.58 | 4.04 (dd; 7.3, 5.0) | 10 | 2, 4, 8, 9, 10, 11 |
| 2 | 38.56 | -- | -- | -- |
| 3 | 24.86 | -- | -- | -- |
| 4 | 123.39 | -- | -- | -- |
| 5 | 115.43 | 6.49 (s) | -- | 1, 3, 6, 7, 8, 9 |
| 6 | 143.96 | -- | -- | -- |
| 7 | 144.44 | -- | -- | -- |
| 8 | 113.11 | 6.55 (s) | -- | 1, 3, 4, 5, 6, 7 |
| 9 | 125.49 | -- | -- | -- |
| 10 | 36.71 | 1.98 (m; 2H) | 1, 11 | 1, 9, 11, 12 |
| 11 | 30.33 | 2.62 (m; 2H) | 10 | 1, 10, 12, 13, 17 |
| 12 | 131.47 | -- | -- | -- |
| 13 | 129.13 | 7.03 (m) | 14 | 11, 14, 15, 17 |
| 14 | 115.18 | 6.69 (m) | 13 | 12, 13, 15, 16 |
| 15 | 155.49 | -- | -- | -- |
| 16 | 115.18 | 6.69 (m) | 17 | 12, 14, 15, 17 |
| 17 | 129.13 | 7.03 (m) | 16 | 11, 13, 15, 16 |

NMR spectra are shown in **Supplementary Figs. 38-42**. s = singlet, d = doublet, dd = doublet of doublets, m = multiplet.

**Supplementary Table 12.** $^{13}C$ and $^{1}H$ δ assignments as well as 2D-NMR correlations of 1-(4-hydroxyphenethyl)-7-methoxy-1,2,3,4-tetrahydroisoquinolin-6-ol (**11**) recorded in DMSO-$d_6$.

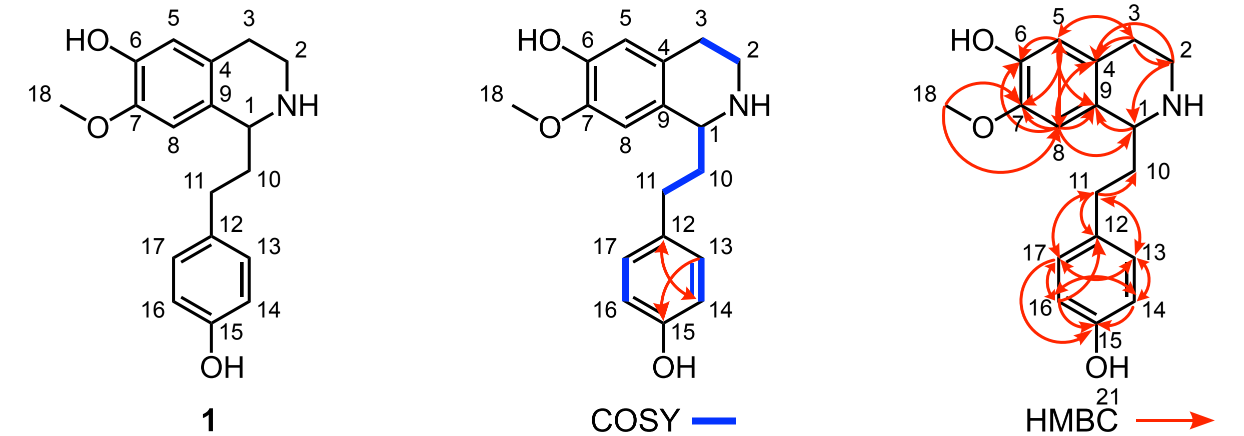


| C-# | δ $^{13}C$ (ppm) | δ $^{1}H$ (mult.; *J* in Hz) | COSY | HMBC |
|---|---|---|---|---|
| 1 | 54.01 | 4.23 (m) | 10 | -- |
| 2a | 38.92 | 3.13 (m) | 2b, 3a, 3b | 1, 4 |
| 2b | 38.92 | 3.33 (overlapping with $H_2O$) | 2a, 3a, 3b | 1, 4 |
| 3a | 25.05 | 2.74 (m) | 2a, 2b, 3b | 2, 4, 5 |
| 3b | 25.05 | 2.84 (m) | 2a, 2b, 3a | 4 |
| 4 | 124.70 | -- | -- | -- |
| 5 | 115.24 | 6.56 (s) | -- | 1, 3, 6, 7, 8, 9 |
| 6 | 145.77 | -- | -- | -- |
| 7 | 146.55 | -- | -- | -- |
| 8 | 110.19 | 6.72 (s) | -- | 1, 3, 4, 5, 6, 7 |
| 9 | 124.22 | -- | -- | -- |
| 10a | 36.17 | 2.07 (m) | 1, 11 | -- |
| 10b | 36.17 | 2.15 (m) | 1, 11 | -- |
| 11a | 30.40 | 2.62 (m) | 10 | 10, 12, 13, 17 |
| 11b | 30.40 | 2.68 (m) | 10 | 10, 12, 13, 17 |
| 12 | 131.12 | -- | -- | -- |
| 13 | 129.20 | 7.06 (m) | 14 | 11, 14, 15, 17 |
| 14 | 115.18 | 6.69 (m) | 13 | 12, 13, 15, 16 |
| 15 | 155.55 | -- | -- | -- |
| 16 | 115.18 | 6.69 (m) | 17 | 12, 14, 15, 17 |
| 17 | 129.20 | 7.06 (m) | 16 | 11, 13, 15, 16 |
| 18 | 55.85 | 3.73 (s) | -- | 7, 8 |

NMR spectra are shown in **Supplementary Figs. 43-47**. s = singlet, m = multiplet.

**Supplementary Table 13.** $^{13}C$ and $^{1}H$ δ assignments as well as 2D-NMR correlations of 1-(3,4-dihydroxyphenethyl)-1,2,3,4-tetrahydroisoquinoline-6,7-diol (**12**) recorded in DMSO-d₆.

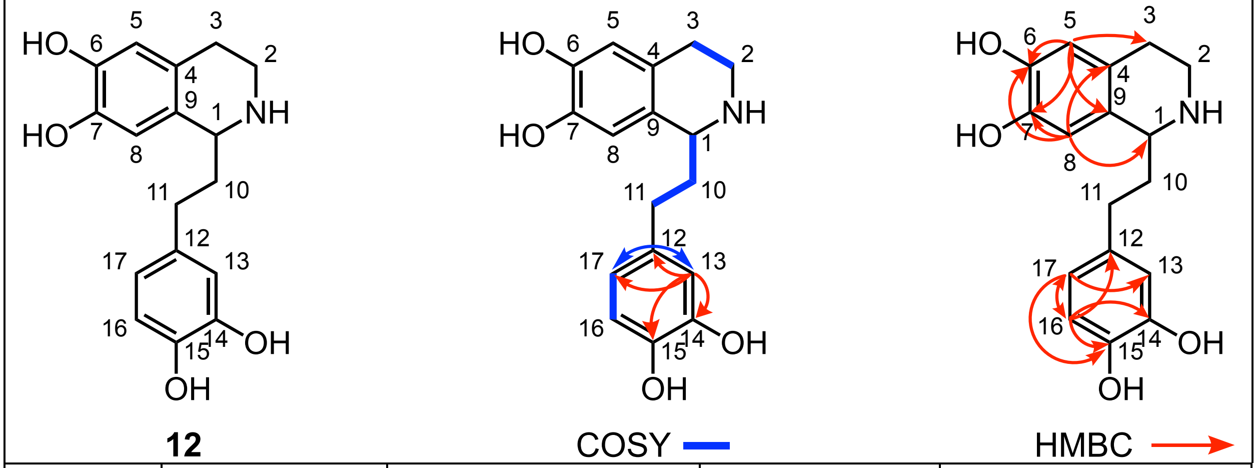


| C-# | δ $^{13}$C (ppm) | δ $^{1}$H (mult.; *J* in Hz) | COSY | HMBC |
|---|---|---|---|---|
| 1 | 53.83 | 4.22 (t; 5.7) | 10 | -- |
| 2a | 39.05 | 3.21 (m) | 2b, 3a, 3b | -- |
| 2b | 39.05 | 3.37 (m) | 2a, 3a, 3b | -- |
| 3a | 24.66 | 2.75 (m) | 2a, 2b, 3b | -- |
| 3b | 24.66 | 2.84 (m) | 2a, 2b, 3a | -- |
| 4 | 122.72 | -- | -- | -- |
| 5 | 115.58 | 6.54 (s) | -- | 3, 6, 7, 9 |
| 6 | 144.30 | -- | -- | -- |
| 7 | 144.71 | -- | -- | -- |
| 8 | 113.39 | 6.59 (s) | -- | 1, 4, 6, 7 |
| 9 | 123.59 | -- | -- | -- |
| 10 | 36.01 | 2.04 (m; 2H) | 1, 11 | -- |
| 11 | 30.47 | 2.58 (m; 2H) | 10 | -- |
| 12 | 131.91 | -- | -- | -- |
| 13 | 115.95 | 6.66 (d; 1.7) | 17 | 12, 14, 15, 17 |
| 14 | 145.47 | -- | -- | -- |
| 15 | 143.72 | -- | -- | -- |
| 16 | 115.88 | 6.65 (d; 8.0) | 17 | 12, 14, 15, 17 |
| 17 | 119.11 | 6.49 (dd; 8.1, 1.9) | 13, 16 | 13, 15, 16 |

NMR spectra are shown in **Supplementary Figs. 48-52**. s = singlet, d = doublet, dd = doublet of doublets, t = triplet, m = multiplet.

**Supplementary Table 14.** $^{13}C$ and $^{1}H$ δ assignments as well as 2D-NMR correlations of 1-(3,4-dihydroxyphenethyl)-1,2,3,4-tetrahydroisoquinoline-3,3,4,4-$d_4$-6,7-diol (**12-D₄**) recorded in DMSO-d₆.

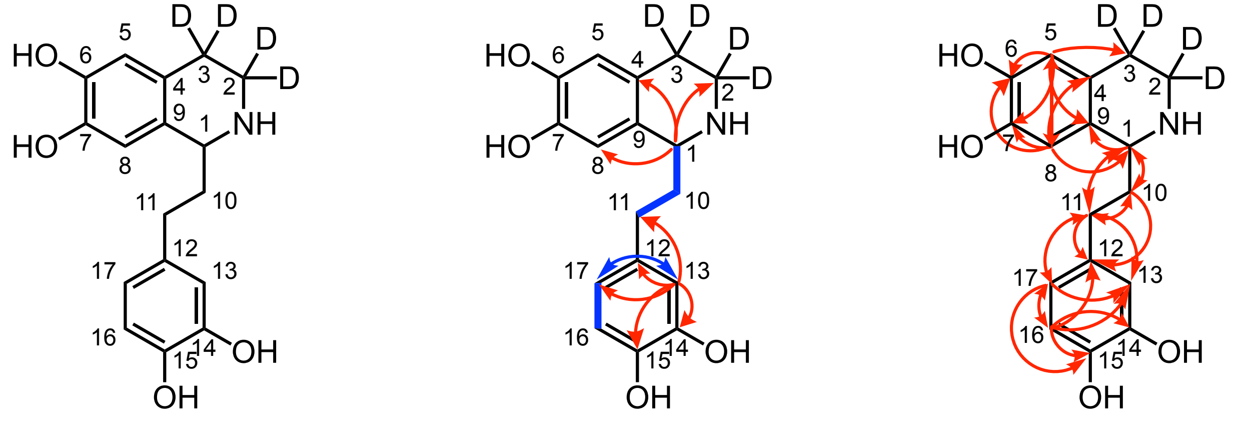


| C-# | δ $^{13}C$ (ppm) | δ $^{1}H$ (mult.; *J* in Hz) | COSY | HMBC |
|---|---|---|---|---|
| 1 | 53.51 | 4.22 (dd; 7.4, 4.9) | 10 | 2, 4, 8, 9, 10, 11 |
| 2 | 38.28 | -- | -- | -- |
| 3 | 23.44 | -- | -- | -- |
| 4 | 122.40 | -- | -- | -- |
| 5 | 115.39 | 6.55 (s) | -- | 1, 3, 6, 7, 8, 9 |
| 6 | 144.31 | -- | -- | -- |
| 7 | 145.00 | -- | -- | -- |
| 8 | 113.21 | 6.61 (s) | -- | 1, 3, 4, 5, 6, 7 |
| 9 | 123.10 | -- | -- | -- |
| 10 | 35.76 | 2.07 (m; 2H) | 1, 11 | 1, 9, 11, 12 |
| 11 | 30.30 | 2.60 (t; 8.1; 2H) | 10 | 1, 10, 12, 13, 17 |
| 12 | 131.70 | -- | -- | -- |
| 13 | 115.85 | 6.67 (d; 1.7) | 17 | 11, 12, 14, 15, 17 |
| 14 | 145.22 | -- | -- | -- |
| 15 | 144.51 | -- | -- | -- |
| 16 | 115.68 | 6.66 (d; 4.0) | 17 | 12, 13, 14, 15, 17 |
| 17 | 118.89 | 6.50 (dd; 8.0, 2.1) | 13, 16 | 11, 13, 15, 16 |

NMR spectra are shown in **Supplementary Figs. 53-57**. s = singlet, d = doublet, dd = doublet of doublets, t = triplet, m = multiplet.

**Supplementary Table 15.** $^{13}$C and $^{1}$H δ assignments as well as 2D-NMR correlations of cephalotaxinone (**10**) recorded in $CDCl_3$.

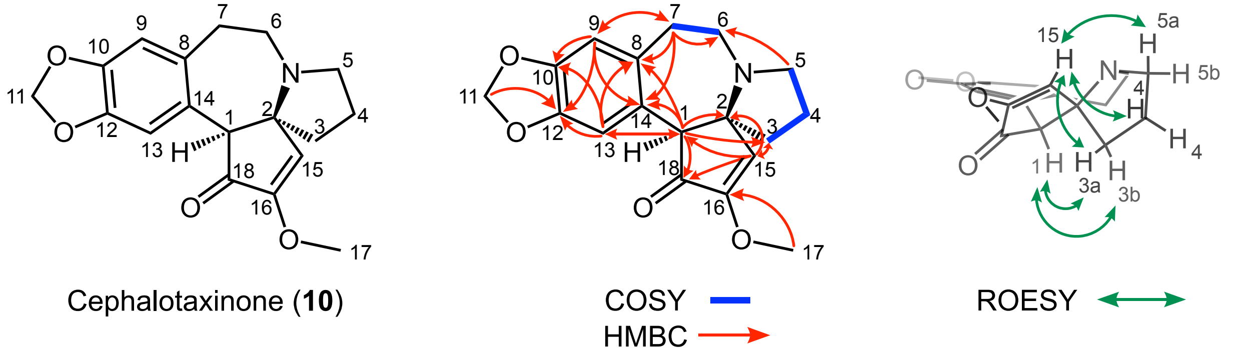


| C-# | δ $^{13}$C (ppm) | δ $^{1}$H (mult.; *J* in Hz) | COSY | HMBC | key ROESY |
|---|---|---|---|---|---|
| 1 | 60.78 | 3.53 (s) | -- | 2, 3, 8, 13, 14, 18 | 3a, 3b |
| 2 | 65.48 | -- | -- | -- | |
| 3a | 39.02 | 1.96 (m) | 3b, 4 | 15 | 1, 15 |
| 3b | 39.02 | 2.10 (m) | 3a, 4 | -- | 1 |
| 4 | 20.10 | 1.87 (m; 2H) | 3a, 3b, 5a, 5b | -- | 15 |
| 5a | 53.01 | 2.69 (q; 8.8) | 4, 5b | 6 | 15 |
| 5b | 53.01 | 3.10 (m) | 4, 5a | -- | |
| 6a | 47.78 | 2.53 (m) | 6b, 7 | 8 | |
| 6b | 47.78 | 2.91 (q; 10.3) | 6a, 7 | -- | |
| 7 | 31.51 | 2.44 (m; 2H) | 6a, 6b | 6, 8, 9, 14 | |
| 8 | 130.72 | -- | -- | -- | |
| 9 | 110.41 | 6.64 (s) | -- | 7, 10, 12, 14 | |
| 10 | 147.35 | -- | -- | -- | |
| 11a | 101.18 | 5.91 (d; 1.5) | -- | 12 | |
| 11b | 101.18 | 5.92 (d; 1.5) | -- | 12 | |
| 12 | 146.36 | -- | -- | -- | |
| 13 | 112.61 | 6.70 (s) | -- | 1, 8, 10, 12 | |
| 14 | 128.55 | -- | -- | -- | |
| 15 | 123.87 | 6.40 (s) | -- | 1, 2, 18 | 3a, 4, 5a |
| 16 | 158.30 | -- | -- | -- | |
| 17 | 57.40 | 3.80 (s) | -- | 16 | |
| 18 | 200.97 | -- | -- | -- | |

NMR spectra are shown in **Supplementary Figs. 58-63**. s = singlet, d = doublet, dd = doublet of doublets, t = triplet, q = quartet, m = multiplet.

**Supplementary Table 16.** Comparison of the $^1$H δ assignments of cephalotaxinone (**10**) recorded in $CDCl_3$ to previously published spectrum[8].

| δ $^1$H (mult., *J* in Hz) in $CDCl_3$ | Published (ref. 8) δ $^1$H (mult., *J* in Hz) in $CDCl_3$ |
|---|---|
| 6.70 (s, 1H) | 6.70 (s, 1H) |
| 6.64 (s, 1H) | 6.64 (s, 1H) |
| 6.40 (s, 1H) | 6.40 (s, 1H) |
| 5.92 (d, *J* = 1.5 Hz, 1H) | 5.92 (d, *J* = 1.5 Hz, 1H) |
| 5.91 (d, *J* = 1.4 Hz, 1H) | 5.91 (d, *J* = 1.4 Hz, 1H) |
| 3.80 (s, 3H) | 3.80 (s, 3H) |
| 3.53 (s, 1H) | 3.53 (s, 1H) |
| 3.13 – 3.07 (m, 1H) | 3.11 – 3.08 (m, 1H) |
| 2.91 (q, *J* = 10.3 Hz, 1H) | 2.93 – 2.89 (m, 1H) |
| 2.69 (q, *J* = 8.8 Hz, 1H) | 2.71 – 2.67 (m, 1H) |
| 2.56 – 2.49 (m, 1H) | 2.54 – 2.51 (m, 1H) |
| 2.47 – 2.41 (m, 2H) | 2.45 – 2.43 (m, 2H) |
| 2.14 – 2.05 (m, 1H) | 2.11 – 2.08 (m, 1H) |
| 1.99 – 1.93 (m, 1H) | 1.98 – 1.95 (m, 1H) |
| 1.89 – 1.85 (m, 2 H) | 1.89 – 1.85 (m, 2H) |

**Supplementary Table 17.** Comparison of the $^{13}$C δ assignments of cephalotaxinone (**10**) recorded in $CDCl_3$ to previously published spectrum[8].

| δ $^{13}$C (ppm) in $CDCl_3$ | Published (ref. 8) δ $^{13}$C (ppm) in $CDCl_3$ |
|---|---|
| 200.97 | 201.0 |
| 158.30 | 158.3 |
| 147.35 | 147.4 |
| 146.36 | 146.4 |
| 130.72 | 130.7 |
| 128.55 | 128.6 |
| 123.87 | 123.9 |
| 112.61 | 112.6 |
| 110.41 | 110.4 |
| 101.18 | 101.2 |
| 65.48 | 65.6 |
| 60.78 | 60.8 |
| 57.40 | 57.4 |
| 53.01 | 53.0 |
| 47.78 | 47.8 |
| 39.02 | 39.0 |
| 31.51 | 31.5 |
| 20.10 | 20.1 |

**Supplementary Table 18.** Primers used for cloning CET biosynthetic genes in this study.

| Gene | Primer Name | Plasmid | Forward /Reverse | Sequences |
|---|---|---|---|---|
| ***Nicotiana benthamiana expression*** | | | | |
| OMT-1 | OMT-1_F | pEAQ-HT | Forward | **ATT CTG CCC AAA TTC GCG ACC GGT** ATG GCT CTC GCA TCT CTG TCC |
| OMT-1 | OMT-1_R | pEAQ-HT | Reverse | **GAA ACC AGA GTT AAA GGC CTC GAG** CTA TAA CTG AGA AGG TAC TTT AAA AGT TG |
| CYP805A11 | CYP805A11_F | pEAQ-HT | Forward | **ATT CTG CCC AAA TTC GCG ACC GGT** ATG GAA AAC TTT CTT GTG AAT CAT ATT ATT G |
| CYP805A11 | CYP805A11_R | pEAQ-HT | Reverse | **GAA ACC AGA GTT AAA GGC CTC GAG** TTA ATT ACG GCG TGT GAC ACG AG |
| NmrA-1 | NmrA-1_F | pEAQ-HT | Forward | **ATT CTG CCC AAA TTC GCG ACC GGT** ATG GCA GAA ATT GTT GAT ATT CAG AGG |
| NmrA-1 | NmrA-1_R | pEAQ-HT | Reverse | **GAA ACC AGA GTT AAA GGC CTC GAG** TTA TAC ATA TCG CTT CAA AAA TAA TTC TAC AG |
| CYP805A12 | CYP805A12_F | pEAQ-HT | Forward | **ATT CTG CCC AAA TTC GCG ACC GGT** ATG GAT ACC TTT CTT GTG AAT TCT ATT AT |
| CYP805A12 | CYP805A12_R | pEAQ-HT | Reverse | **GAA ACC AGA GTT AAA GGC CTC GAG** CTA AGT ACG GCA TCT GAT ACG AG |
| CYP805A13 | CYP805A13_F | pEAQ-HT | Forward | **ATT CTG CCC AAA TTC GCG ACC GGT** ATG GAA AAC TTT GTG GTG AAT TAT ATT ATT G |
| CYP805A13 | CYP805A13_R | pEAQ-HT | Reverse | **GAA ACC AGA GTT AAA GGC CTC GAG** TCA AGT ACG GTA AGT GAT ACG AGC |
| CYP805A14 | CYP805A14_F | pEAQ-HT | Forward | **ATT CTG CCC AAA TTC GCG ACC GGT** ATG ATG AGA AAA GGA ATA TCA TTG TGG |
| CYP805A14 | CYP805A14_R | pEAQ-HT | Reverse | **GAA ACC AGA GTT AAA GGC CTC GAG** TCA AGT ACG GTA AGT GAT ACG AG |
| 2OGD-1 | 2OGD-1_F | pEAQ-HT | Forward | **ATT CTG CCC AAA TTC GCG ACC GGT** ATG GCT TCC TCG CTG GAA AAT GA |
| 2OGD-1 | 2OGD-1_R | pEAQ-HT | Reverse | **GAA ACC AGA GTT AAA GGC CTC GAG** TCA AAC AAC GCT AGA AAT TCC GGC |
| ***Yeast expression*** | | | | |
| CYP805A11 | CYP805A11-Y_F | pYeDP60 | Forward | **TAC ACA CAC TAA ATT ACC GGA TCC** ATG GAA AAC TTT CTT GTG AAT CAT ATT ATT G |
| CYP805A11 | CYP805A11-Y_R | pYeDP60 | Reverse | **CAT GGG AGA TCC CCC GCG GAA TTC** TCA GTG GTG GTG GTG GTG GTG ATT ACG GCG TGT GAC ACG AGC |
| ***E. coli expression*** | | | | |
| NmrA-1 | NmrA-1-B_F | pET-24b | Forward | **TAA CTT TAA GAA GGA GAT ATA CAT** ATG GCA GAA ATT GTT GAT ATT CAG AGG |
| NmrA-1 | NmrA-1-B_R | pET-24b | Reverse | **TCA GTG GTG GTG GTG GTG GTG CTC GAG** TAC ATA TCG CTT CAA AAA TAA TTC TAC AGT G |

Primer sequences that overlap with the destination plasmid for DNA assembly are in **bold**.

**Supplementary Table 19.** Nucleotide sequences for CET biosynthetic genes cloned and described in this manuscript.

| Gene | Transcript ID | Sequence |
|---|---|---|
| OMT-1 | YD-1-Rt_transcript/50321.p1 | ATGGCTCTCGCATCTCTGTCCACCGCCTTTTCTTTC CTGAAAGCAATGATGAACGAAAAGGAAGCGATTA AGACGGGCCAAACAGGGGACGGCAGAGAGAAAGC ATTGCTCGACCATGTCCTCAAAACGGCTAAACAGG GGGATCCGCTCTCCGTCTTGAATGCGATGGACACT TATGCAAAGACGGTTAGCTGGTTGATGAACTTCGG CGACGAGAAGGGTCCCATTTTAGACAACGCTCTGA AAAAATATAATCCAAAAGCCACTCTGGAGATCGGA ACCTACTGCGGGTATTCTGCAGTGAGAATTGCGTC GCAGTTGCAGAGGGAAGGCAGTAAGCTTCTCGCTT TCGAGATGGGCGCCGATAACTGCAATATCGCCAGG CAAATTATCGACCACGCAGGGCTGTCGTCGAAGGT GGATGTGGTGGAAGGAAAATTCGACGAGCGTTTAG ATGTGGCCAGGCAGTTTTGAAGGAGATGAAGGCT TCCTGCTTTGATTTTGTTTTCCTCGACCACTGGAAA GACCGCTATTTGGCAGATTACCTGATCTTGAAGGA AAACGGTATGCTGGGGAATGGAACCGGTATATGTG CAGACAATATGGGGAGATTCGAGGGACCCGTCGA CTTCAAGAAATATATAAAAACGCATCCGGAAGAGC TTGAGAGCTTCGAACATAAAAGCCATATCGAGTAC CTTTGGTGGCTTCCAGACAGCGTCGTCGTCTCAACT TTTAAAGTACCTTCTCAGTTATAG |
| CYP805A11 | YD-3-Rt_transcript/31277.p1 | ATGGAAAACTTTCTTGTGAATCATATTATTGAGGC CCTTCATTTCTTTCACACCCTCATCTTTGAAGGCGC TGCTATTTCACTTCTTCTTTTTTCAATTGTCCTTACA CTTTTAGCAGCTCACTATTTGTTATCAGCCAACAAG AAAGGGAACAGGATGTGGCCTCCATCCCCACCCAC ACTGCCTTTAATTGGTAACATGCATCAGATGAACA AAGGCGACACTTTCCTTTCGGCAATAACTGATATG GCGAAGTCGTATGGTCCAGTCATTACCCTGTGGTT GGGAACATCGCCTCTGGTAATTCTCACCGGCCAGA CTGCAATTTGGGAGGCACTGGTTAACCAGGCCTCC AATTTCGCAGGTCGTCCATTCTTTCCCACAAATCAA TTTCTCTCCGCCGACAGTGAGCAAACCACCCTGAC TTCAAGCAGCGAGCGTTGGGTTAAGCTCAGGAAAA TTCTTCACAACAACGTTCTCAGTCCCTTCCACATTG CAAGGCATAGCTCTTTCCACCAGAGAAACGTGCAT GATCTCATTAGCAAAATAGAGGCGGAGATGAAAG CAAGCAAGGGCGTCGTGAGGCCTCTTCATTTCCTG AAAATAATGGCAATAATTTTTTTTGCCCAATTATGC TTCGGCCCTGATTTTCATGACGAAATTTTCGCCAAC AAGATTGAACAATTCCTAGCAGAAGGTATTATCCT CAGTGGAAGAGGAGCAACGCTGCTGGAGTCGCTTC CTTTGGCTCGCTACTGGCATCCTTCGTCCTACATAA CAGAGAAGAAAGTAAATGCTATGAGAGCTGGCGT TATGGAGCTCTTCTTACCTCTCATCCAATTTGGCCG GCGTTACAGAGCATACTTGAAGCGAAGTGCTCCCA ACTGTTTCTTGAATTGTCTGCTGTCCATCGGCGAAG GAAAAGAAGATGACGTCAAGTCAAAGCTATCCGA TGTAGAAATAACTTTCAACCTGTTTGAGTTATTTCT TCTTAGCGTTGACAGCACTTCCACGTCAATAGAAT GGGCTCTGGCTTATCTCGTAAACAATCCCCACATA CAAGATAAGGTTTATCAAGAGGTAACTGAAGCTCT GCCGGAAAGAAAAGGGGAACTTATTCGTTTGGAG GACTTGGGGAAGCTGCCATATTTGCAAGCTGTGGT GAAAGAAACACTGAGAAAAGAAGCCATTGCACCT ATTACTGTGCCTCACCAAACTCTCAATGAGTGTAA |

| | | |
|---|---|---|
| | | GGTGATGGGCATCAACATACCCGCGAAGACAGTA<br>GTGATTTGCAATCTGCATTCTGTGTCCAACGATCCC<br>CAACTGTGGAGCGATCCTGATGAGTTCCGGCCAGA<br>AAGATTTGTGGGCAATAATGATGTCAAATCACGTT<br>ATCTTGCGTTTGGAGCAGGAAGGCGCATGTGTGCA<br>GGAATGGACATGGCTTACGCCTACATTCCTATCAC<br>TCTTGCCAACTTACTCAAAAACTTCGAATGGGGGT<br>GTGTTAAGGAAGGGAGTCCTCCAGATCTCTCGCGC<br>GAGGAACGTTCAATGATCATGTTTATGAAACATCC<br>TCTGGAAGCTCGTGTCACACGCCGTAATTAA |
| NmrA-1 | YD-1-Rt_transcript/46483.p1 | ATGGCAGAAATTGTTGATATTCAGAGGAAGATGGC<br>AGATATGGGTAACTCTAACAGCAAAGTTTTGGTAA<br>TAGGAGGTACAGGAAGTTTAGGACAAAGGCTTGTT<br>AGGGCAAGCCTGGCTCTTAACCATCCTACTTATGTT<br>TTATTTCGTCGTCAAGCAACTCATGATATTGCCAAA<br>GTGGATCTACTCATTTCTTTCAAGCAATCCGGTGCT<br>CACCTTCTCCAAGGCTCATTGGATGACCATGAGAG<br>CTTGGTGGCTGCTCTCAAGCAAGTGGATGTTGTCA<br>TATCAGCTCTGGCAGGAACTGTTACAAAAAATGAC<br>ATAAGTAAGCAACTGAAATTATTGGATGCCATTAA<br>GGAAGCTGGCAATATCAAGCGTTTCCTCCCTTCAG<br>AATTTTGGGTGGATCCAGATAGATTGGTGGATTCT<br>GTGCATCCACCAAATCCAGTTGTTATTGATAAGAG<br>AAATCTGAGGCGGGCTATAGAGGCAGCCAATGTCC<br>CATACACTTACATCATAGCCAACTGCTTTGCTGGA<br>ATCTTTTTAGGAAGCATTGGACAGGTTGACCAAGG<br>TTTTATTCCTTCCAGAGAAAAAGTGGCCTTATATGG<br>TGATGGCAATGCTAAAGTAATATGGGTGGATGAGA<br>ATGACATTGCAACATATACAATCAAAGCAATAGAT<br>GATCCTCGGACCTTGAACAGAGCAGTTCATATTAG<br>GCCACCTACAAATATCCTGTCACAGAAGGAGGTTG<br>TAGAAATATGGGAAAAACTCATAGGTCACCCTTTG<br>GTGAAAACTCAAATTTCAGGGGAGGATTGGCTCAA<br>GACCATGGAAGGTAAACCTTTTGAAATACAAGCTA<br>TGATTGCACATGCCCATGAGCTGTTCTATGATGGTT<br>GTACTGCCAATTTTGAGATTAGGCCTGATGAAAAA<br>GAAGCTTCTCAACTGTATCCAGATGTGAAGTATAC<br>CACTGTAGAATTATTTTTGAAGCGATATGTATAA |
| CYP805A12 | YD-1-Rt_transcript/28576.p1 | ATGGATACCTTTCTTGTGAATTCTATTATCGAGGCC<br>CTTCATTTCTTTCACATCCCCATCTTTGGAGGCGCT<br>GCTATTTCACTTCTTCTATTTTCATTTGTTCTCACAG<br>TTGTGGCGGCTCACTATTTATTATCACCCAACAAG<br>AATGAGAAAAAGATGTGGCCTCCATCTCCACCCAC<br>ACTGCCTTTATTGGGTAACTTGCTCCAGTTGAACAA<br>TGGCGGCAGTTTTCTTTCGGCGGCAAATCACATGG<br>CGAAGTCGTATGGTCCAGTCATTACCCTGTGGATG<br>GGAACATCGCCTCTGGTACTCCTCACCGGCCAAAC<br>TGCAATTTGGGAGGCGCTGGTTAACCAGGCCTCCA<br>ATTTCGCAGATCGTCCATACATGCAGAGAAATAGA<br>TTTTTCGCCTTCGGTGACATAACCACCATGACTTCT<br>GACTGCAACGAGAGTTGGATTAAGCTGAGGAAAA<br>TTCTTCACAGCAATGTTGTCAGTCGCTTCCACATTG<br>CAAGCCAGAGCTCTTCCCATCAGAGAAATGTGAAT<br>GATCTTATTAAGAAATTAGTGGAGGAAATGAAAGC<br>AAACAATGGCGTTGTGAGGCCTTTTCATTCCTTTAA<br>AATAATGGCAATAAGTTTTGTTGCGCAATTATGCTT<br>CGGCCCTGATTTTCAAGACGAAAACTTCGCCAACA<br>AGGTTGAAAATTTTATTGCTGAAGATATTGTCGTG<br>AGTGCAAGAGGAGAAACGCTGCTGGAGACGATTC<br>CTTTCGCTAGCTACTGGAATCCTTCGTCTTATATCA<br>CCCAAAAAAAAATAAAGACTTTGGGAGCTGGCGTT |

| | | |
|---|---|---|
| | | CTGGAGCTCTTGTTACCTATCATCCAATCTGCCCGG<br>AGCTACAGAGCATATTTGAAGCAAAGTGCTCCCAA<br>CTGTTTGTTGAATTGTTTACTGTCGATTTGCGAAGG<br>AGAAAATGATGAATTAAAGTTAAAGCTCTCCGATG<br>AAGCAATAGCTTTCAACATATTTGAGCTGTTTATTC<br>TCAGCGTGGACAGCACCTCCACGGCATTAGAATGG<br>GCTCTGGCTTATCTGATAAACAATCCCCACATACA<br>AAATAGGGTTTATCAAGAGGTAATTGAAGCTTCGC<br>CTGGAGGAAAAGAGCAGCTTCTTCGCGTAGAGGAC<br>CTGGGGAAGCTGCCATATTTGCAAGCTGTGGTGAA<br>AGAAACACTGAGAAAAGAGGCCATCGCACCTTTTG<br>CTGTGCCTCACCGAACTGCAAACGAGTGCAAGGTG<br>ATGGGTATCAACATACCCGCGAATGCACCAGTGTT<br>TTTCAATCTGTCTTCTGCGCACAACGATCCCCAACT<br>GTGGAGCGATCCTGATGAGTTCCAGCCGGAAAGAT<br>TTCTCGGCAATAATGATGTGCGATCATGTTTTCTTC<br>CGTTCGGAGCAGGAAGGCGCATATGTCCTGGAATG<br>GACATGACGTACACGCACGTTCCGATCACCCTCGG<br>CAACTTAATAAAAACCTTCGAATGGGAGTGTGTTA<br>AGGAAGGGAGTCCTCCAGATCTTACGCGCCAACAG<br>CGTGCACTGCTTATGTTTATGAAATATCCTCTGGAA<br>GCTCGTATCAGATGCCGTACTTAG |
| CYP805A13 | YD-2-Rt_transcript/33273.p1 | ATGGAAAACTTTGTGGTGAATTATATTATTGAGGC<br>GTTTCAACATTATTTCTTTGACATGATCCCCATCTT<br>TGGAGGCGCTTCAGTTGTTCTATTTTCATTTGTTCT<br>CACAGTTGTGGCGGCTCACTATTTGTTATCAGACA<br>AGAAGAAGAATGATGAGAAAAGGATGTCATGGCC<br>TCCATCTCCACCCACAATGCCTTTATTGGGTAACTT<br>GCACATGTTGAACAAAGGAGGCAATTTTCTTATGG<br>CGGCGTATGATATAGCGAAGTCGTATGGTCCAGTG<br>ATGACCCTGTGGATGGGAACATCACCTGCGGTCGT<br>CCTCACTGGCCAAACTGCAATTTGGGAGGCGCTGG<br>TTAACCAGGCCTCCAATTTCGCAGATCGTCCATTCA<br>TGAACACAAACCGATTTTTCTCCTCCGGTGACATA<br>ACCACCATGACTTCTGACTGCAACGAGAATTGGAT<br>GAAGCTGAGGAAAATTCTTCACAACAACGTTATCA<br>GTCGCTTCAACATTGCAAGTCACAGCTCTTCCCACC<br>ACAGAAATGTGAATGGTCTTATTAAGAAGTTAGTG<br>GAGGAAATGAACGCAAACAATGGCGTTGTGAGGC<br>CTTTCCATTCCTTCAAAATAATGGGAATAAGTTTTA<br>TGGCCAAATTATGCTTCGGCCCTGATTTTGAAGAC<br>GAAATTTTCGCCATCAAGGTTGAAAATTTAATAGC<br>TGAAGATATTGCCTTGATTGGAAAAGGAGAAACGC<br>TGCTGGAGTCGATTCCTTTGGCTCGCTACTGGAATC<br>CTTCTACTTTCTTAACAAAAAAGAATCAAAAGGCT<br>ATCAGCGCTAGCATTCTGGAGCTCTTGTTACCTATC<br>GTCAAATATGGGCGGAGTTATAGAGCATACTTGAA<br>GCAAAGTGCTCCCAACTCTTATTTGAATTGTTTGCT<br>GTCAATTTCCGAAGAAGAAGACGATGAATTGAAGT<br>TAAAGCTCTCCGATGAAGAAATAGCTTTCAACATA<br>TTTGAGCTGCTTATTCTTAGCGTGGACAGCACCTCC<br>ACGGCATTAGAATGGGCTGTGGCTTATCTCATCAA<br>CAATCCCCACATACAAAATAAGGTTTATCAAGAGG<br>TAAATGAAGCTTTGCCAGGAGGAAAAGAGCGGCTT<br>CTTCGTGTAGAGGACTTGGAGAAGCTGCCATATTT<br>GCAAGCAGTAGTAAAAGAAACACTGAGAAAAGAA<br>GCCGTTGCACCTTTTGCTTTGCCCCACCAAACTGCA<br>AATGAGTGCAAGGTGATGGGCGTCAACATACCCGC<br>CAAGGCAACAGTGCTTTTCAATCTGTTTTCTGTGAA<br>CAACGATCCCCAACTGTGGAGCAATCCTGATGAGT<br>TCGCGCCAGAAAGATTTCTGGGGAATAATGTCGAT |

| | | |
|---|---|---|
| | | GTGCGATCATGTTATCTTCCGTTCGGAGCAGGAAG GCGCATATGTCCAGGAATGGACATGGCGTACATTC ACGTTCCAGTCACCCTTGGCAGCTTAATGAAATAC TTCGAATGGGGGTGTGTTAAGGAAGGGAGTCCTCC AGATCTCTCCCGCGAGGAACGTTCAATGATCATGT ATATGAAACATCCTCTGGAAGCTCGTATCACTTAC CGTACTTGA |
| CYP805A14 | YD-3-Rt_transcript/38063.p1 | ATGATGAGAAAAGGAATATCATTGTGGCCTCCATC TCCACCCAGACTGCCTTTATTGGGTAACTTGCACAT GTTGAACAAAGGAGGCAATTTTCTTATGGCGGCGT ATGATATAGCGAAGTCGTATGGTCCAGTGATGACC CTGTGGATGGGAACATCACCTGCGGTCGTCCTCAC TGGCCAAACTGCAATTTGGGAGGCGCTGGTTAACC AGGCCTCCAATTTCGCAGATCGTCCATTCATGAAC ACAAACCGATTTTTCTCCTCCGGTGACATAACCAC CATGACTTCTGACTGCAACGAGAATTGGATGAAGC TGAGGAAAATTCTTCACAACAACGTTATCAGTCGC TTCAACATTGCAAGTCACAGCTCTTCCCACCACAG AAATGTGAATGGTCTTATTAAGAAGTTAGTGGAGG AAATGAACGCAAACAATGGCGTTGTGAGGCCTTTC CATTCCTTCAAAATAATGGCAATAAGTTTTATGGC CAAATTATGCTTCGGCCCTGATTTTGAAGACGAAA TTTTCGCCATCAAGGCTGAAAATTTAATAGCTGAA GATATTGCCTTGATTGGAAAAGGAGAAACGCTGCT GGAGTCGATTCCTTTGGCTCGCTACTGGAATCCTTC TACTTTCATAACAAAAAAGAATCAAAAGGCTATCA GCGCTAGCATTCTGGAGCTCTTGTTACCTATCGTCA AATATGGGCGGAGTTATAGAGCATACTTGAAGCAA AGTGCTCCCAACTCTTATTTGAATTGCTTGCTGTCG ATTTCCGAAGAAGAAGATGATGAATTGAAGTTAAA GCTCTCCGATGAAGAAATAGCTTTCAACATATTTG AGCTGCTCATTCTTAGCGTGGACAGCACCTCCACG GCATTAGAATGGGCTGTGGCTTATCTCATCAACAA TCCCCACATACAAAATAAGGTTTATCAAGAGGTAA ATGAAGCTTTGCCAGGAGGAAAAGAGCGGCTTCTT CGTGTAGAGGACTTGGAGAAGCTGCCATATTTGCA AGCAGTGGTAAAAGAAACGCTGAGAAAAGAAGCC GTTGCACCTTTTGCTTTGCCCCACCAAACTGCAAAT GAGTGCAAGGTGATGGGCGTCAACATACCCGCCAA GGCAACAGTGCTTTTCAATCTGTTTTCTGTGAACAA CGATCCCCAACTGTGGAGCAATCCTGATGAGTTCG CGCCAGAAAGATTTCTGGGGAATAATGTCGATGTG CGATCATGTTATCTTCCGTTCGGAGCAGGAAGGCG CATATGTCCAGGAATGGACATGGCGTACATTCACG TTCCAATCACCCTTGGCAGTTTAATAAAATACTTCG AATGGGGGTGTGTTAAGGAAGGGAGTCCTCCAGAT CTCTCCCGCGAGGAACGTTCAATGATCATGTATAT GAAACATCCTCTCCAAGCTCGTATCACTTACCGTA CTTGA |
| 2OGD-1 | YD-1-Rt_transcript/46979.p1 | ATGGCTTCCTCGCTGGAAAATGAGGTTGATCTCCC TCTCATTGAACTTTCACAAATTCCGTTGGAGTTTGA CGGAGTAAACAACCTTCAATTCCATCCGGCGGTAG CCAGAGTCGGAGAAGCCTGCAAAGAATGGGGATT CTTCCGTGTTGTGGACTCTGGAATTCCAACAGATCT TCTCCAGAATTTAGAGTCCGTTAGCCGCCAAATAT GTGACATGCCTCCAGAGGTCAAAGACAGAGCCATC GGATCTATCCCCTATGAGAGCTACTTTCGCTATCCT TTGAGGGAAAGCTTCTGCATCCAAAATTTGCCTCA ATCGGATTCAGTTCAAGCATTGTCAAATAAGATAT GGCCGAATAATGAAAACCCTAAATTCAGCGAGACT GTAGGAGCATATGCAGATTTCCTTGCAGGTCTGCA |

| | | |
|---|---|---|
| | | ACGCAAGATCTTTATCATCATTCTTGCCAGCTTGGG TTTGGACGTGAAGACTTTCTACCATTCTGATTTCCA GGAATGCACAGCTTACTTGCATATAAACCACTACT ACGGTAGTGATGGAAAATCCTCAAAGCAGGAAGC CCTTGAAACGCATACAGATATCAGTTGCTTTACTAT CCTTTACCAAGACAACGAGGGGGGGCTCCAGATTC GATCAAAGGAAGGGGAGATGTTTAACGTCAAATCT GTCTCCAATTCATTTATTGTCAATATAGGCAACTGC TTTAAGGCATGGAGCAACGGCAGATACCGCAGCTC AGAGCACCGGGTTGTTTTAAAGGCTGGACGGATC GCATATCTCTTGGGTGGTTTGTGCTTTTTCCTAATG ATCAACAAATATTGGCTCCGGCGGAGCTTGTGGAC GACGATCATCCAAGACGTTACAAGCCTTTCACCTT CTACCAGTACAAGCAGGGTGCCTGGAATAATTTTC TGTCCAAAAAAGAGACGTATACGACATTCATGGAA GATTATGCCGGAATTTCTAGCGTTGTTTGA |

Note: 2OGD-1 is 100% identical to YD-3-Rt_transcript/53545.p1 and differs by a single SNP from YD-1-Rt_transcript/46979.p1 (99.9% identity). Because redundant transcripts were collapsed using CD-HIT-EST at a 99% identity cut-off (see Methods), YD-1-Rt_transcript/46979.p1 was retained in the final transcriptome and used for gene expression analysis; the 2OGD-1 sequence that we cloned and functionally characterized corresponds to YD-3-Rt_transcript/53545.p1. Similarly, CYP805A11 shares 99.9% nucleotide identity with YD-3-Rt_transcript/31277.p1, differing by a single SNP.

**Supplementary Table 20.** Nucleotide sequences for genes cloned and assayed for Pictet-Spengler like condensation between dopamine and the corresponding aldehydes to form compounds **1** or **2**, but which showed no detectable activity in this study (Supplementary Fig. 12).

| Gene | Transcript ID | Sequence |
|---|---|---|
| *Ch*PSS | YD-1-Rt_transcript/53915.p1 | ATGGTGGCGAAAAGTATAACAGTAGAAATTGATTC TCCAGTGGAGGCAAAGAGATTTTGGGCTGCAATTG TTAAGGACTATGATCTCCTCCCTAGGCAAATGCCA GGGGTATGTTCAGGCGTCACACTTGTCAAAGGCGA CGGAGGTGTCGGCACCATCATACAAATTGACTTCA CCCCTGTGAACAAGGATTTTAGTTACGTAAAGGAG CGAGTGGATGAAGTAGATGAGGGAAACTTTGTTTA CAGCTTCAGCTATGTGGAAGGGGGAGAGGTGGGA ACGAAATGGGCATCTGCTAAGTTTAAGATGCAATT GACACCGAAAACAGAGGGTGGATGCGTTTTGAAG CATACATGCGAGTATGACACCCTGCCCGGTATTCC CCTTGACGAAGCCAAACTGGAAGAGATGGAGAGG AGCGCCGCTGGTCTCTTAAAAGCCATAGAAGCATA CCTCGTCTCTAACCCTACCGTATATTGTTAA |
| *Ch*BetV-1 | YD-1-Rt_transcript/53876.p1 | ATGGTAGCAGGCAGCTTCAATATAGAAGTAGACTC TCCAGTGCAGGCAAAAAGACTGTGGAATGCGATG GTGAAGGATGGGCATAATCTACTCCCAAAGCAAGT TCCACACATAATCGAAAGCATCGCCTATCTTCAAG GCGACGGAGGAGTGGGCACCGTCATGCAAGTGAA CTTCACCTCTGCAAACAATTTTTTGAAGGAACTCGT GGAGGAAATAGACGACGAGAACTTTGCATTTAGTT GCAGTATTTTGGAGGGTGGACAGGTGGGAAAGAT ATTGGCTTCTGAGAAACATGTGGAGAAATACGCCG CTAAAGCAGGAGGTGGAACCCTGTGGACGGGCAC TGTGCACTATGAAACTCTGCCTGGTATTTCCCCTGA TGAATACAACACGCAGGAGATTAAAGACGTTTACA CCAGTATGTTCAAGCTCATTGAGGCATATCTCCTTG CTAACCCCACTCTCTACTGTTAA |
| *Ch*BetV-2 | YD-3-Rt_transcript/61329.p1 | ATGGTAGCAGGCAGCTTCATTATAGAAGTAGATTC TCCAGTGGAGGCAAAGAGACTGTGGAATGCGTCG GTGAAGGATGGGCATAATCTCGTCCCCAAGCAAGC TCCACACATAATCGAAAGCATCACCTTTCTCCAAG GCGACGGAGGCGTGGGCACCGTCAAGCAAGTGAT |

| | | |
|---|---|---|
| | | CTTCACCTCTGCAAACAAAGTTTTAGCTATGCGA AGGAACGCGTGGAGGAAATGGATGATGAGAACTT TGTATACAGTTGCAGTGTTTTGGAGGGAGGACAGG TGGGAAAGATATTGGCTTCTGAGAAACAGGTGGAG AAATACATCCCTAAAGCAGGGGGTGGAACCCTATG GACGGCCACGGTACACTATGAAACTCTCCCTGGTA TTTTGCCTGATGAATACAACATGCAGGAGATGAAA GACGGGTTCATGGGCATGTTCAAGCTCATCGAGGC ATATCTCCTCGCCAACCCCACTCTCTACTGTTAA |
| *Ch*BetV-3 | YD-1-Rt_transcript/54093.p1 | ATGGTAGCAGGCAGCTTCAATATAGAAGTAGATTC TCCAGTGCTGGCAAAAAGACTGTGGAATGCCTTCG TGAAGGATGGGCATAATCTCGTCCCCAAGCAACTT CCAGACCTAATCGAAAGCATCACCTTTCTCCAAGG CGACGGAGGCGTGGGCACCGTCAGGCAAGTTAACT TCACCTCTGCAAACACAGATTTTAGTTATGTGAAG GAGCGCGTGGAGGAAATGGATGAGGAAAATTTTG TATACAGTTGCAGTGATGTGGAGGGAGGACAGCTG GGAAAGATATTGGCTTCTTCGAAACAAGTGGAGAA ATACACCCCTAAAGCCGGGGGTGGATCCGTGTGGA CGGCCACATTTCACTATGAAATACTCCCTGGTATTT CCCCTGATGAATACAACATTCAGGAGATGAAAGAC GGCTTCATCGGTATGTTCAAGCTCATCGAGGCATA TCTCCTCGCTAACCCCACTCTCTACTGTTAA |
| *Ch*BetV-4 | YD-2-Rt_transcript/59141.p1 | ATGGTGGCAGGAAGCATCAGTCATGAATACGAGTC CACAGTGGACGTAAAGAGACTTTGGAATGCATTTG TGAAGGACGGCCATAATCTTATCCCAAAGCAAGTG CCGGAGGTTTTCGCAAGCATCACAATCCTCCAAGG CGATGGAGGTGTCGGCACCATCAGGCAAATAAACT TCACCCCTGGTATGTTAATGTACCTGCACATTTATT TCTTAATATTACGATTATCTTTACTGTTTATGCAAC ACTACTTAATCTGTAGTACGCTTTCAGCGAACGAG TATTTTAGCTACGTGAAGGAGCGCGTAGACGAAGT CGATGAGGAGAATTTAGTATACCGCTTCAGTCACG TGGAGGGAGGAATGTTGGGAAAGAAATTCGCTTCT GCGAAGTACGAGTACAGATACACCCGCAACGCCG GGGGCGGAAGCGTTGTCCACTTCGTAATGCATTTT GACAGCCTGCCTGGTGTTCCCGAGGACGAAGGCGA TGTCATGGAGATGAAAGAGAAGACCACCGCTTACT TTAAAAAGATTGAGGCACATCTCCTTGCCAACCCC ACCAGCTACTGTTAA |
| *Ch*BetV-5 | YD-1-Rt_transcript/53150.p1 | ATGGTGGCAGGAAGCATCAGTCATGAATACGAGTC CACAGTGGACGTAAAGAGACTTTGGAATGCATTTG TGAAGGACGGCCATAATCTTATCCCAAAGCAAGTG CCGGAGGTTTTCGCAAGCATCACAATCCTCCAAGG CGATGGAGGTGTCGGCACCATCAGGCAAATAAACT TCACACCTGCGAACGAGTATTTTAGCTACGTGAAG GAGCGCGTAGACGAAGTCGATGAGGAGAATTTAG TATACCGCTTCAGTCACGTGGAGGGAGGAATGTTG GGAAAGAAATTCGCTTCTGCGAAGTACGAGTACAA ATACACCCGCAAGGCCGGGGGCGGAAGCGTTGTCC ACTTCGTAATGCATTTTGACAGCCTGCCTGGTGTTC CCGAGGACGAAGGCGATGTCATGGAGATGAAAGA GAAGACCACCGCTTACTTTAAAAAGATTGAGGCAC ATCTCCTTGCCAACCCCACCAGCTACTGTTAA |
| *Ch*BetV-6 | YD-2-Rt_transcript/59530.p1 | ATGGTGGCAGGAAGCATCAGTCATGAATACGAGTC CACAGTGGACGTAAAGAGACTTTGGCATGCATTTG TGAAGGACGGCCATAATCTTATCCCAAAGCAAGTG CCGGAGGTTTTCGCAAGCATCACAATCCTCCAAGG CGATGGAGGTGTCGGCACCATCAGGCAAATAAACT TCACCCCTGGTATATTAAGGTACCTGCACATTTATT TCTTAATATTACGATTATCTTTACTGTTTATGCAAC |

| | | |
|---|---|---|
| | | ACTACTTAATCTGTAGTACGCTTTCAGCGAACGAG<br>TATTTTAGCTACGTGAAGGAGCGCGTAGACGAAGT<br>CGATGAGGAGAATTTAGTATACCGCTTCAGTCACG<br>TGGAGGGAGGAATGTTGGGAAAGAAATTCGCTTCT<br>GCGAAGTACGAGTACAAATACACCCGCAAGGCCG<br>GGGGCGGAAGCGTTGTCCACTTCGTAATGCATTTT<br>GACAGCCTGCCTGGTGTTCCCGAGGACGAAGGCGA<br>TGTCATGGAGATGAAAGAGAAGACCACCGCTTACT<br>TTAAAAAGATTGAGGCACATCTCCTTGCCAGCCCC<br>ACCAGCTACTGTTAA |
| *Ch*BetV-7 | YD-1-Rt_transcript/54278.p1 | ATGGTAGCAGGCAGCTTCATTATAGAAGTAGACTC<br>CCCCGTGGAGGCAAAGAGACTGTGGAATGCAATG<br>GTGAAGGATGGGCATAATCTCATCCCCAAGCAAGT<br>TCCACACATATTCGAAAGCGTCACCATTCTCCAAG<br>GAGACGGAGGCGTGGGCACAGTCAGACAAGTCAA<br>CTTCACCTCTGCAAATAAGGATTTTAGCTATGTGA<br>AGGAGCGCGTGGATGCAATTGACGAGGAGAAGTT<br>TTATTACTGTTACAGTGATGTGGAGGGTGGAGAGC<br>TGGGGAAGAAATTGGCTTCCGCGAAACAGGAGGT<br>GAAATACACTCCTAAAGCAGGGGGTGGATCCGTGT<br>GGACGTCCACGATTCACTATGAAACACTCCCTGGT<br>ATTTCCCCTGATGAAGGCAAATTGCAGGAGATGAA<br>AGACAGTTTCACCACTTTGTTCAAGCTCATCGAGG<br>CATATCTCCTCGCCAACCCCACTCTCTACTGTTAA |
